# Somatic LINE-1 retrotransposition in cortical neurons and non-brain tissues of Rett patients and healthy individuals

**DOI:** 10.1101/506758

**Authors:** Boxun Zhao, Qixi Wu, Adam Yongxin Ye, Jing Guo, Xianing Zheng, Xiaoxu Yang, Linlin Yan, Qing-Rong Liu, Thomas M. Hyde, Liping Wei, August Yue Huang

**Affiliations:** National Institute of Biological Sciences, Beijing, 102206, China; Graduate School of Peking Union Medical College, Beijing, 100730, China; School of Life Sciences, Peking University, Beijing, 100871, China; Peking-Tsinghua Center for Life Sciences, Beijing, 100871, China; Center for Bioinformatics, State Key Laboratory of Protein and Plant Gene Research, School of Life Sciences, Peking University, Beijing, 100871, China; Academy for Advanced Interdisciplinary Studies, Peking University, Beijing, 100871, China; College of Life Sciences, Beijing Normal University, Beijing, 100875, China; Laboratory of Clinical Investigation, National Institute on Aging, Baltimore, MD 21224, USA; Lieber Institute for Brain Development, Baltimore, MD 21205, USA; Departments of Psychiatry & Behavioral Sciences and Neurology, Johns Hopkins University School of Medicine, Baltimore, MD 21205, USA

**Keywords:** mosaicism, retrotransposition, somatic insertion, Rett syndrome

## Abstract

Mounting evidence supports that LINE-1 (L1) retrotransposition can occur postzygotically in healthy and diseased human tissues, contributing to genomic mosaicism in the brain and other somatic tissues of an individual. However, the genomic distribution of somatic L1Hs (Human-specific LINE-1) insertions and their potential impact on carrier cells remain unclear. Here, using a PCR-based targeted bulk sequencing approach, we profiled 9,181 somatic insertions from 20 postmortem tissues from five Rett patients and their matched healthy controls. We identified and validated somatic L1Hs insertions in both cortical neurons and non-brain tissues. In Rett patients, somatic insertions were significantly depleted in exons—mainly contributed by long genes—than healthy controls, implying that cells carrying *MECP2* mutations might be defenseless against a second exonic L1Hs insertion. We observed a significant increase of somatic L1Hs insertions in the brain compared with non-brain tissues from the same individual. Compared to germline insertions, somatic insertions were less sense-depleted to transcripts, indicating that they underwent weaker selective pressure on the orientation of insertion. Our observations demonstrate that somatic L1Hs insertions contribute to genomic diversity and MECP2 dysfunction alters their genomic patterns in Rett patients.

**Author Summary:** Human-specific LINE-1 (L1Hs) is the most active autonomous retrotransposon family in the human genome. Mounting evidence supports that L1Hs retrotransposition occurs postzygotically in the human brain cells, contributing to neuronal genomic diversity, but the extent of L1Hs-driven mosaicism in the brain is debated. In this study, we profiled genome-wide L1Hs insertions among 20 postmortem tissues from Rett patients and matched controls. We identified and validated somatic L1Hs insertions in both cortical neurons and non-brain tissues, with a higher jumping activity in the brain. We further found that MECP2 dysfunction might alter the genomic pattern of somatic L1Hs in Rett patients.

## Introduction

The term “somatic mosaicism” describes the genomic variations that occur in the somatic cells that make up the body of an individual. These variations contribute to intra-individual genetic diversity among different cells (Campbell *et al.*, 2015). In addition to various types of cancers, somatic mosaicisms reportedly contribute to a variety of neurological disorders, including epilepsy, neurodegeneration, and hemimegalencephaly (Poduri *et al.*, 2013). The human-specific LINE-1 (L1Hs) retrotransposon family is the only known family of active autonomous transposons in the human genome (Hancks and Kazazian, 2012; Kazazian and Moran, 2017). L1s retrotranspose through a process called target-primed reverse transcription (TPRT), with the capacity for *de novo* insertion into new genomic locations in both germline and somatic cells (Cost *et al.*, 2002; Luan *et al.*, 1993). Mounting evidence supports that L1Hs elements, with increased copy number in the brain relative to other tissues, contribute to neuronal diversity via somatic retrotransposition (Coufal *et al.*, 2009; Erwin *et al.*, 2016; Evrony *et al.*, 2012; Evrony *et al.*, 2015; Muotri *et al.*, 2010; Upton *et al.*, 2015).

Recent studies reported the occurrence of somatic L1Hs insertions during neurogenesis and in non-dividing mature neurons (Coufal *et al.*, 2009; Macia *et al.*, 2017). Other studies have observed dysregulated L1Hs copy number in patients with Rett syndrome (Muotri *et al.*, 2010) and schizophrenia (Bundo *et al.*, 2014). Methyl-CpG binding protein 2 (*MECP2*) is the major disease-causing gene of Rett syndrome (Amir *et al.*, 1999). Its gene product, MeCP2, can bind to the 5’ UTR of L1 elements and represses their expression and retrotransposition (Yu *et al.*, 2001). While it is known that L1 expression and copy number are elevated in the brains of *Mecp2* knockout mice as well as in patients with Rett syndrome (Muotri *et al.*, 2010; Skene *et al.*, 2010), little is known about the genomic distribution patterns of somatic L1Hs insertions in Rett patients and healthy individuals.

In contrast to germline insertions, the effects of somatic transposon insertions depend not only on their genomic location. Rather, the specific timing, tissue, and cell lineage at which they occur profoundly influence the impact of somatic insertions (Frank, 2010). Single-cell targeted sequencing approaches have been used to identify somatic insertions (Erwin *et al.*, 2016; Evrony *et al.*, 2012; Upton *et al.*, 2015). However, such methods typically require a large number of cells and demand considerable sequencing depth for unbiased profiling of human tissues (Grun and van Oudenaarden, 2015; Navin, 2015). Furthermore, owing to the rarity of somatic insertions, investigations of the clonal diversity of somatic insertions would require the sequencing of even larger numbers of cells (Navin, 2015).

Another limitation of single-cell sequencing approaches is that errors of allelic dropout and locus dropout, which frequently occur during the whole genome amplification (WGA) step of library construction, can reduce the sensitivity and specificity of somatic insertion detection. Estimates of the rate of somatic L1Hs insertions vary widely in single-cell genomics studies (Faulkner and Garcia-Perez, 2017). Bulk sequencing approach can potentially overcome these limitations and enable the genome-wide identification and quantification of somatic L1Hs insertions, but their low allele frequency in cell populations poses a great challenge to distinguishing true insertion events from technical artifacts (Evrony *et al.*, 2016).

Here, we introduced a PCR-based multiplex bulk sequencing method for sensitive enrichment and specific identification of L1Hs insertions from various types of human tissues. We used this method to perform genome-wide L1Hs insertion profiling of 20 postmortem tissues from five patients with Rett syndrome and their matched healthy controls. The aims of this study were to explore the genomic patterns of somatic L1Hs insertions in neuronal and non-neuronal samples, and to investigate whether MECP2 dysfunction could alter the distribution of L1Hs retrotransposition in patients with Rett syndrome.

## Results

### A bulk sequencing method to identify L1Hs insertions

Systematic genome-wide profiling of somatic L1Hs insertions requires effective enrichment of insertion signals and specific identification of true signals from background noise. Enriching neuronal nuclei from bulk brain tissue facilitates the accurate deciphering of cell type-specific characteristic and increases the chance of identifying clonal somatic insertions that are derived from the same progenitor cell and shared by multiple neurons. Therefore, we labeled prefrontal cortex (PFC) neuronal nuclei using an antibody against neuron-specific marker NeuN (Mullen *et al.*, 1992), and subsequently purified NeuN^+^ nuclei from postmortem human PFC by fluorescence-activated cell sorting (FACS) (Fig 1A; S1A–D Fig; Appendix 1). All initially sorted nuclei were re-analyzed with a second round of FACS, and the purity of the initial sorting was found to be > 96% (S1E–F Fig; Appendix 1). The integrity and purity of sorted nuclei were confirmed by fluorescence microscopy (S2A–C Fig).

**Fig 1.**
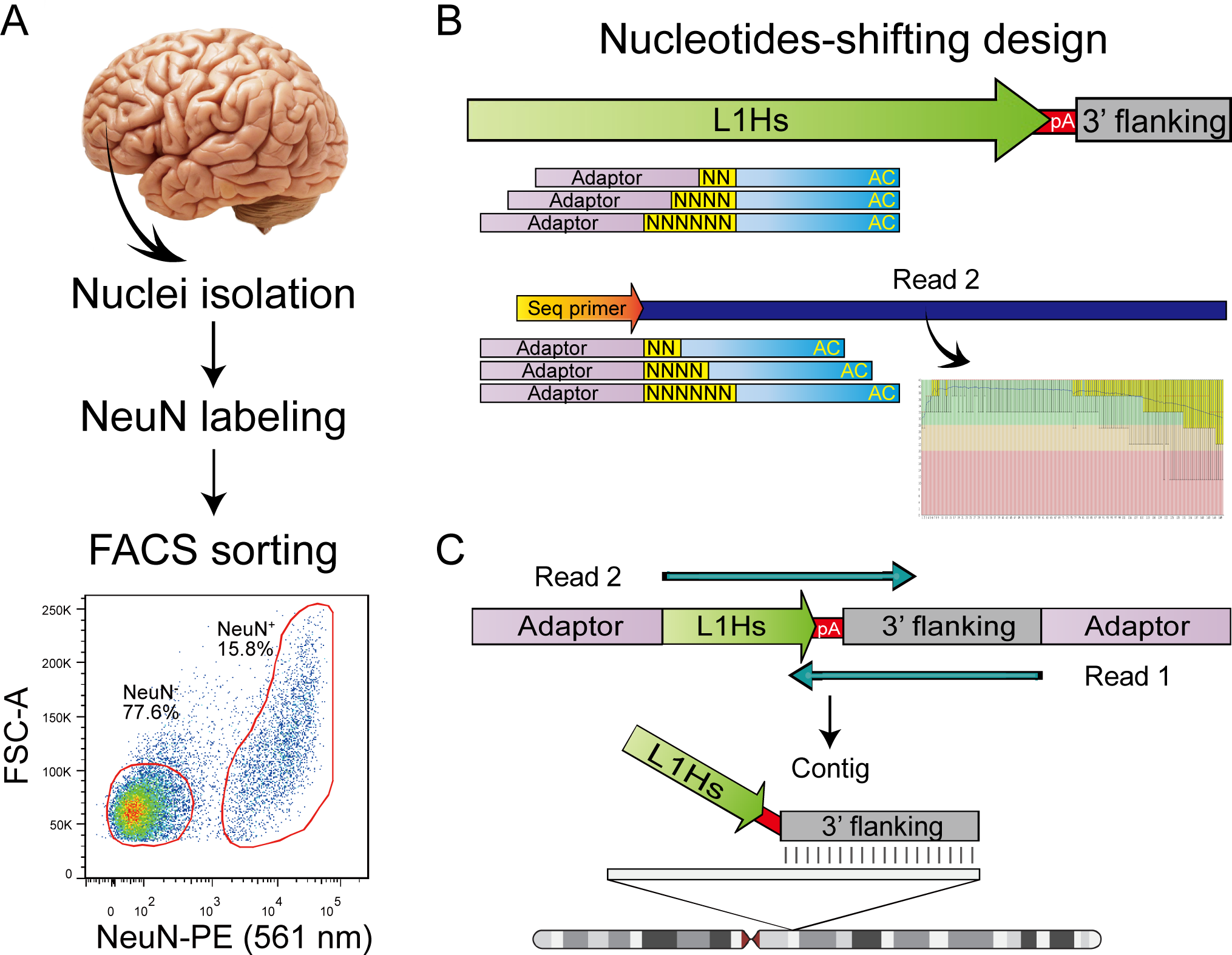
Overview of human active transposon sequencing. (**HAT-seq).** (A) Fluorescence-activated cell sorting (FACS) of prefrontal cortex (PFC) nuclei labeled with NeuN. Two populations (NeuN^+^ and NeuN^−^) were sorted. (B) Schematic of the nucleotides-shifting design of the HAT-seq method. By adding two, four, or six random nucleotides upstream of L1Hs-specific primer (L1Hs-AC-28), we transformed the library from a uniform phase-0 amplicon library to a mixed library with phase-2, phase-4, and phase-6 amplicons, which remarkedly improved the base calling accuracy in Read 2. (C) HAT-seq libraries were sequenced with paired-end 150-bp reads. After merging paired reads into contigs that fully spanned the L1Hs-genome 3’ junction, genomic locations of each L1Hs insertion were determined by the alignments of their 3’ flanking genomic sequences.

To distinguish the signals of active L1Hs elements from other transposon families that are typically inactive in human, we developed a method called human active transposon sequencing (HAT-seq) (Fig 1B; S3A Fig; S1 Table) based on ATLAS (Badge *et al.*, 2003) and several versions of high-throughput sequencing-based L1 amplification methods (Erwin *et al.*, 2016; Ewing and Kazazian, 2010; Philippe and Cristofari, 2016; Tang *et al.*, 2017). Firstly, L1Hs insertions were specifically enriched and amplified using a primer targeting the diagnostic “AC” motif of L1Hs (Hancks and Kazazian, 2012; Ovchinnikov *et al.*, 2001). To ameliorate the poor performance of Illumina sequencing platform for low-diversity libraries, we employed a nucleotides-shifting design by adding two, four, or six random nucleotides upstream of the L1Hs-specific primer, which greatly increased the diversity of the structure-transformed semi-amplicon library and markedly improved the sequencing quality of L1Hs 3’ end. The constructed libraries preserved information regarding the insertion direction and were sequenced by multiplexed 150 bp paired-end reads. This approach provided sequence information fully spanning the 3’ L1Hs-genome junction of each of L1Hs insertions, which enabled the identification of integration sites and facilitated *in silico* false-positive filtering based on both sequence features and read-count.

Genomic position of each L1Hs insertion was determined by the alignment of its 3’ flanking sequence (Fig 1C). A custom data analysis pipeline classified putative insertions into one of the following four categories: known reference (KR) germline insertions, known non-reference (KNR) germline insertions, unknown (UNK) germline insertions, and putative somatic insertions (S3B Fig). To further remove technical artifacts induced by non-specific or chimeric amplification and read misalignment in next-generation sequencing, we designed a series of stringent error filters to remove them in different aspects (Table 1): 1) read pairs with non-specific amplification signals and incorrect 3’ truncation were removed based on the sequence of L1Hs 3’ end (Read 2); 2) after merging paired-end reads into contigs, chimeric molecules with abnormal contig structures were identified by BLAST and filtered out; 3) reads with inconsistencies in BWA-MEM and BLAT alignments were defined as mapping errors; and 4) putative somatic insertion signals without multiple PCR duplicates or those present in different individuals were removed, as they were deemed likely to have resulted from sequencing errors. After applying these error filters, the remaining insertions were annotated with peak features to facilitate downstream analysis.

**Table 1.** Error filters used in the computational pipeline.

| **Filter name** | **Definition** |
| --- | --- |
| **Improper alignment** | We rejected reads with less than 30-bp alignment or more than 3 mismatches to the reference genome. |
| **Diagnostic motif** | We rejected reads without L1Hs diagnostic G motif (position 6012 relative to the L1Hs Repbase consensus) (Bao et al., 2015). |
| **Chimera within L1 segment** | We rejected reads with less than 95% identity (> 4 mismatches) to the L1Hs 3' end consensus sequence. |
| **Chimera within poly-A tail** | We rejected reads at risk of being chimeric (Upton et al., 2015). Read was re-aligned to hg19 using BLAST to find the corresponding best alignments for the non-retrotransposon and retrotransposon segments. Read was removed as a putative chimera when the overlap of the two best segments was > 10 bp and A% ≥ 50% or 6–10 bp and A% < 50%. |
| **Subfamily filter** | We rejected putative somatic insertion sites that overlapped with L1 young subfamilies (L1Hs and L1PA2–4) reference insertions. |
| **Known non-reference filter** | We rejected putative somatic insertion sites that overlapped with known non-reference L1 insertions in eul1db (Mir et al., 2015). |
| **Misaligned reads** | We rejected reads at risk of being misaligned, defined as inconsistent BWA and BLAT alignment. |
| **Local SV** | We rejected reads at risk of being derived from a nearby reference L1Hs (Upton et al., 2015). We extracted 2 kb from the reference genome extending downstream from an aligned non-retrotransposon section and aligned the full read contig against this region with BLAT to exclude genomic rearrangements. |
| **Observed in common** | We rejected putative somatic insertion sites observed in two or more individuals. |
| **PCR duplicate** | We rejected somatic insertion sites without supporting PCR duplicates. |

### Performance evaluation of the HAT-seq method using a positive control

To benchmark the performance of HAT-seq for detecting somatic L1Hs insertions, we experimentally generated a series of positive control samples with insertions at different frequencies by mixing the genomic DNA (gDNA) extracted from the blood samples of two unrelated adults, ACC1 and ACC2 (see details in Materials and Methods). 172 ACC1 non-reference germline L1Hs insertions were identified by HAT-seq, 64 of which were confirmed to be ACC1-specific by 3’ junction PCR (3’ PCR) analysis of gDNA from ACC1 and ACC2 (Fig 2A; S2 Table; Appendix 2) and thus served as positive controls. Three HAT-seq libraries were generated from samples consisting of ACC2 gDNA spiked with 1%, 0.1%, or 0.01% of ACC1 gDNA. Considering that decreasing the number of cells pooled for sequencing increased the signal-to-noise ratio for detecting somatic insertions (Evrony *et al.*, 2016), each HAT-seq library was constructed from 20 ng input (about 3,000 cells).

**Fig 2.**
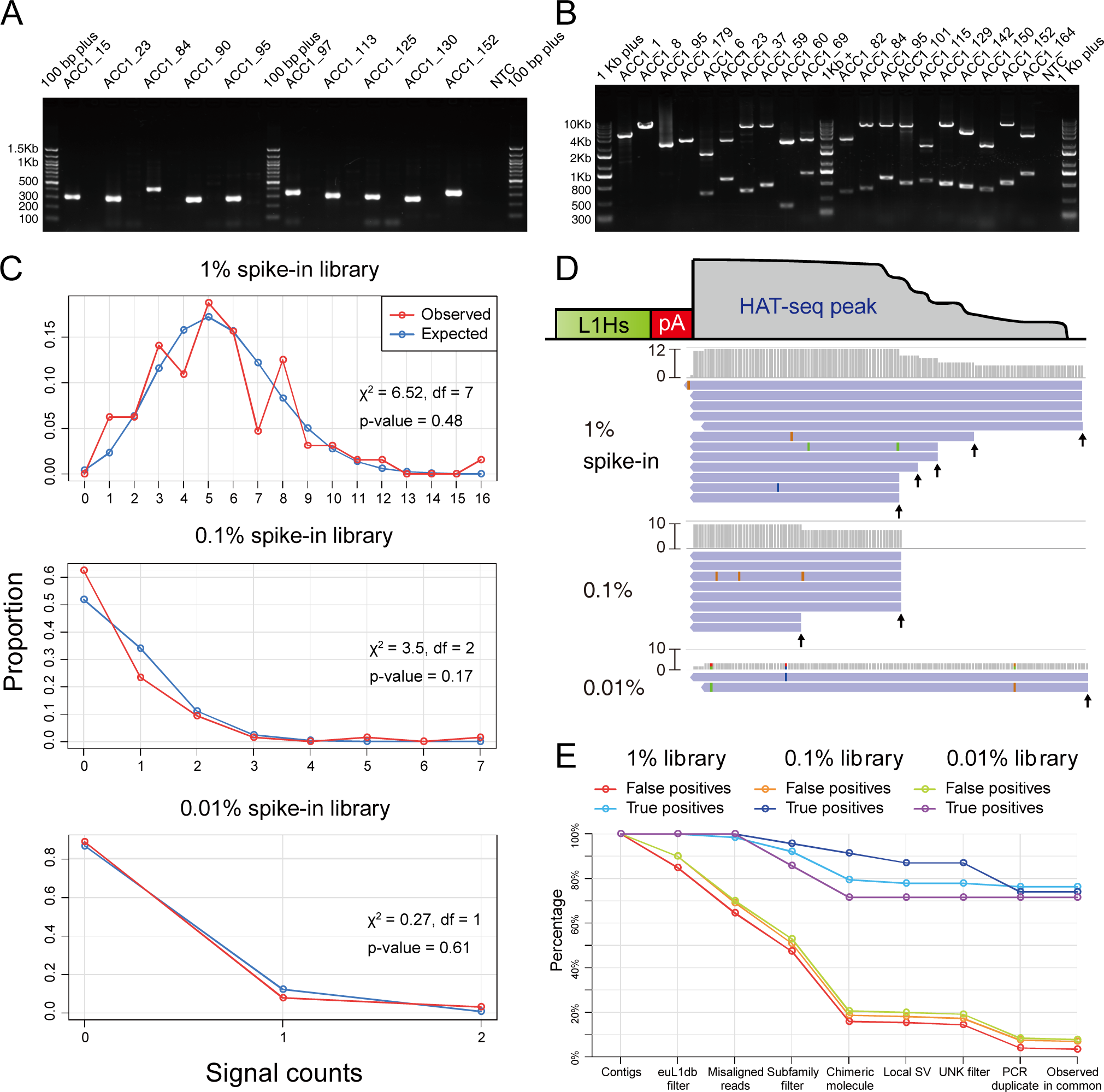
HAT-seq performance evaluation using a positive control. (A) Representative gel image used for the identification of ACC1-specific insertions based on 3’ PCR analysis. For each site, genomic DNA from ACC1 and ACC2 was amplified using the same protocol. NTC: negative control. (B) Representative gel image used for the zygosity analysis of ACC1-specific insertions based on full-length PCR. The four sites on the left were homozygous L1Hs insertions and the others were heterozygous L1Hs insertions. (C) The distributions of signal counts (reads with unique start positions) per ACC1-specific insertion closely followed Poisson distributions (chi-squared goodness-of-fit tests). (D) Representative ACC1-specific insertion (ACC1_132 at chr21:29069173) in 1%, 0.1%, and 0.01% spike-in libraries. Read coverage and supporting signal counts (unique start positions were indicated by black arrows) were positively correlated with the spike-in concentration. (E) The effectiveness of error filters. 64 ACC1-specific germline insertions in 1%, 0.1%, and 0.01% spike-in libraries were considered as “true positives”; all other signals were considered as “false positives”, which might include both background noise and some true somatic insertions present in the blood gDNA.

The zygosity of ACC1-specific L1Hs insertions was confirmed by full-length PCR: 49 of which were heterozygous, 9 of which were homozygous, and 6 of which were zygosity-undetermined (Fig 2B; S2 Table; Appendix 2). We detected all 64 ACC1-specific insertions in our positive control 1% ACC1 spike-in library, 49 (76.6%) of which passed all of error filters and subsequently were deemed “identified” by HAT-seq. In the 0.1% library, we detected 23 ACC1-specific insertions (16 heterozygous, 4 homozygous, and 3 zygosity-undetermined), 17 (73.9%) of which were identified. In the 0.01% library, we detected seven heterozygous ACC1-specific insertions, five (71.4%) of which were identified. The distributions of signal counts (reads with unique start positions) per ACC1-specific insertion followed the Poisson distribution (Fig 2C), indicating a similar probability for each of ACC1-specific insertions to be randomly sampled. In the 1%, 0.1%, and 0.01% libraries, each of ACC1-specific insertions was diluted to 30, 3, and 0.3 copies. Theoretically, by Poisson statistics, there would be 64, 60.81, and 16.59 ACC1-specific insertions being sampled and subsequently being used as the input of HAT-seq libraries (see details in Materials and Methods). Therefore, we estimated the sensitivity of HAT-seq for somatic L1Hs insertions in 1%, 0.1%, and 0.01% libraries as 76.6% (49/64), 28% (17/60.81), and 30.1% (5/16.59), respectively. Our data showed that, with about 3,000 cells as input, HAT-seq was able to detect somatic insertion events present in a single cell (Fig 2D and Appendix 3).

To further evaluate the efficacy of our L1Hs identification pipeline, we compared the proportions of true positives and false positives after applying all the error filters. For the most stringent evaluation, only those 64 ACC1-specific germline insertions in spike-in libraries were defined as “true positives”; all other signals were defined as “false positives”, which might include both background noise and some true somatic insertions present in the blood gDNA. As shown in Fig 2E, in three positive control experiments with 1%, 0.1%, and 0.01% ACC1 gDNA spike-in, 76.56%, 73.91%, and 71.43% of true positives remained after all filters, whereas only 3.40% (66), 6.90% (181), and 7.70% (183) of false positives remained after all filters (S3 Table). These results showed that HAT-seq performed in combination with our error filters could successfully remove most artifacts and identify very low-frequency somatic insertions in bulk DNA samples.

### Profiling of somatic L1Hs insertions in brain and non-brain human tissues

Next, we applied HAT-seq to 20 bulk samples obtained from postmortem neuronal (PFC neurons) and non-neuronal tissues (heart, eye, or fibroblast) from five Rett syndrome patients and five neurologically normal age-, gender-, and race-matched controls (Table 2 and S4–S7 Table). A total of 9,181 putative somatic L1Hs insertions were identified in these 20 HAT-seq libraries (S8 Table). A subset of 137 (1.49%) of these insertions were detected by reads with multiple start positions. Considering that the random fragmentation process in HAT-seq library preparation would result in only one start position shared by all reads generated from a single cell, these 137 insertions should be present in multiple cells in the bulk tissue, and thus classified as “clonal somatic insertions”. Based on the performance evaluation of HAT-seq, the lower bound of the precision of overall somatic L1Hs insertions was 60.14%. To demonstrate the validity of these identified somatic insertions *in silico*, we investigated whether they had the hallmark features of TPRT-mediated retrotransposition (see details in Materials and Methods). By exploiting the sequence information of L1 integration junctions, we found that such somatic insertions were significantly enriched in genomic regions containing L1 endonuclease cleavage motifs (L1 EN motifs) (p < 2.2×10^-16^, Wilcoxon rank–sum test; Fig 3A; S4 Fig; S9 Table). Moreover, our identified somatic insertions shared the 25-bp peak of poly-A tail length with the reference L1Hs insertions (Fig 3B and S9 Table), where some of the somatic insertions with shorter tails might be explained by non-TPRT mechanism (Morrish *et al.*, 2002). These features of somatic L1Hs insertions helped to elucidate the specificity of HAT-seq method.

**Fig 3.**
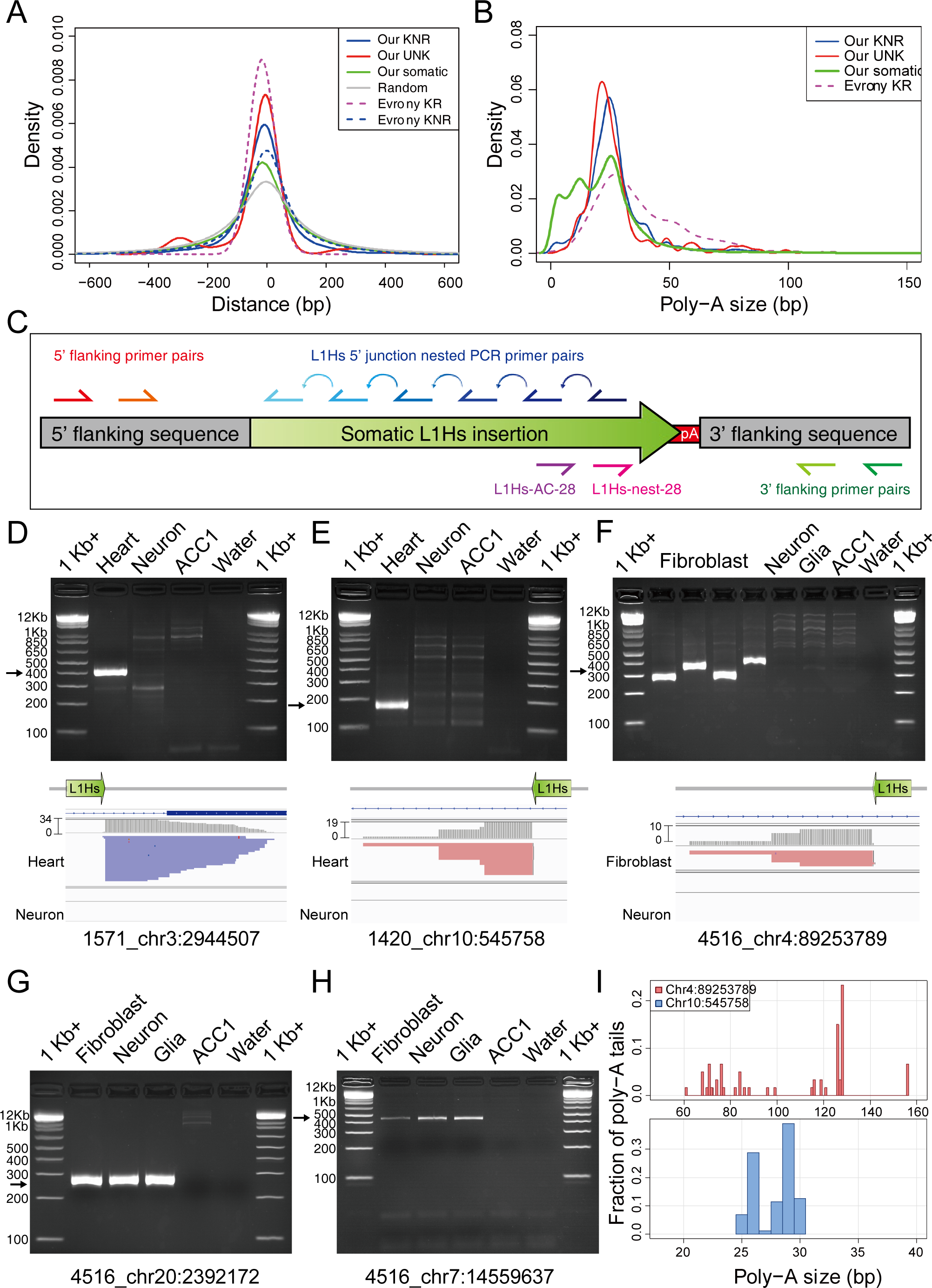
Profiling of somatic L1Hs insertions in multiple human tissues. The density distributions of L1 EN motifs around L1Hs integration sites. L1 EN motifs included seven specific motifs (TTAAAA, TTAAGA, TTAGAA, TTGAAA, TTAAAG, CTAAAA, TCAAAA). “Evrony KR” and “Evrony KNR” are germline L1Hs insertions identified in Evrony *et al*. 2012. (B) The density distributions of poly-A tail length for each category of L1Hs insertion. (C) The PCR validation scheme and locations of primers used. (D)–(H) Representative gel images of 3’ nested PCR validation for putative clonal somatic insertions. The Integrative Genomics Viewer screenshots for (D)–(F) showed the coverage track (gray) and the alignment track (blue for read strand [-]; red for read strand [+]) from HAT-seq data. Black arrows indicated bands with target size. 1Kb +: 1 Kb Plus DNA ladder. (I) Polymorphic poly-A tail sizes of clonal somatic insertions. Top: fibroblast-specific somatic L1Hs insertion at chr4:89253789 from Rett patient UMB#4516. Bottom: heart-specific somatic L1Hs insertion at chr10:545758 from Rett patient UMB#1420.

**Table 2.** Overview of postmortem human tissues.

| UMB ID<sup>a</sup> | Category | Mutation | Age | PMI | Brain tissue | Non-brain tissue | Matched ID | Number of somatic insertions |  | Rate of somatic insertions per cell |  | Known reference insertions (KRs) | Known non-reference insertions (KNRs) | Unknown insertions (UNKs) |
| --- | --- | --- | --- | --- | --- | --- | --- | --- | --- | --- | --- | --- | --- | --- |
|  |  |  |  |  |  |  |  | Cortical neuron | Non-brain | Cortical neuron | Non-brain |  |  |  |
| 4882 | Rett syndrome | c.763C>T (p.R255X) | 17 years<br>310 days | 18 hrs | PFC | Heart | 4591 | 855 | 364 | 1.47 | 1.01 | 819 | 194 | 10 |
| 1815 | Rett syndrome | IVS3-2 A>G | 18 years<br>130 days | 5 hrs | PFC | Eye | 1571 | 589 | 306 | 1.30 | 0.68 | 806 | 189 | 11 |
| 4852 | Rett syndrome | c.451G>T (p.D151Y) | 19 years<br>280 days | 13 hrs | PFC | Eye | 1347 | 380 | 257 | 0.89 | 0.54 | 824 | 171 | 8 |
| 4516 | Rett syndrome | c.763C>T (p.R255X) <sup>b</sup> | 20 years<br>356 days | 9 hrs | PFC | Fibroblast | 1846 | 759 | 411 | 1.82 | 0.69 | 814 | 195 | 7 |
| 1420 | Rett syndrome | No pathogenic mutations | 21 years<br>22 days | 18 hrs | PFC | Heart | 1455 | 708 | 216 | 1.31 | 0.37 | 824 | 182 | 8 |
| 4591 | Healthy control | NA | 16 years<br>223 days | 14 hrs | PFC | Heart | 4882 | 861 | 190 | 1.58 | 0.34 | 816 | 187 | 12 |
| 1571 | Healthy control | NA | 18 years<br>138 days | 8 hrs | PFC | Heart | 1815 | 384 | 221 | 0.63 | 0.57 | 813 | 175 | 11 |
| 1347 | Healthy control | NA | 19 years<br>76 days | 16 hrs | PFC | Heart | 4852 | 553 | 260 | 1.08 | 0.72 | 806 | 179 | 12 |
| 1846 | Healthy control | NA | 20 years<br>221 days | 9 hrs | PFC | Heart | 4516 | 744 | 296 | 1.66 | 0.68 | 817 | 175 | 8 |
| 1455 | Healthy control | NA | 25 years<br>149 days | 7 hrs | PFC | Heart | 1420 | 628 | 203 | 1.16 | 0.41 | 802 | 183 | 11 |

<sup>a</sup> The gender and race for all individuals were female and Caucasian. <sup>b</sup> Genetic variant identified by custom AmpliSeq capture panel (S4 Table).

Owing to the rarity of each somatic insertion in the cell population and to the sensitivity limits of various analytical methods, experimental validation of somatic insertions using unamplified bulk DNA, in particular when one of the primers is complementary to numerous homologous sequences in the human genome is very challenging (Appendix 4). In theory, if a somatic insertion was unique to a single cell, it would be impossible to detect it in any replicated gDNA extracted from the same tissue. To circumvent this, we performed single-copy cloning by adapting a modified version of digital nested 3’ PCR (Evrony *et al.*, 2015) that focused exclusively on clonal somatic insertions with three or more supporting signals, whose mosaicism (percentage of cells) were at least 0.1% based on our experimental design of HAT-seq library (Fig 3C). Five out of eight (62.5%) such clonal insertion sites were confirmed via 3’ nested PCR and Sanger sequencing of cloned amplification products (Fig 3D– H and S10 Table). Four of these clonal somatic insertions were located in introns of *TGM6*, *CNTN4*, *DIP2C*, and *DGKB*; three were sense-oriented to transcripts.

To our knowledge, no somatic insertions in non-brain tissues of healthy individuals has been reported (Faulkner and Billon, 2018). We identified and experimentally validated a heart-specific somatic L1Hs insertion from a healthy individual UMB#1571 (Fig 3D). Leveraging both the 3’ and 5’ junctions of somatic L1Hs insertions enable us to characterize the terminal site duplications (TSDs) and L1 endonuclease cleavage site of insertion. Because most of somatic L1Hs insertions were 5’ truncated with varied lengths, we screened and selected 22 high-quality step-wise primers covering the full-length L1Hs elements to capture their 5’ junction (Fig 3C; S11 Table; Appendix 4). Using 5’ junction nested PCR, we successfully re-captured and Sanger sequenced the 5’ junction of the heart-specific L1Hs insertion in the healthy individual (UMB#1571) (Fig 4A and S11 Table). We confirmed this insertion was a full-length somatic L1Hs insertion with 14 bp TSD and a cleavage site at 5’– TT/AAAG–3’, similar to the consensus L1 EN motif 5’–TT/AAAA–3’ (Fig 4B–D). Notably, we also validated this 5’ junction by combining full-length PCR with 5’ junction PCR (Fig 4E; see details in Materials and Methods).

**Fig 4.**
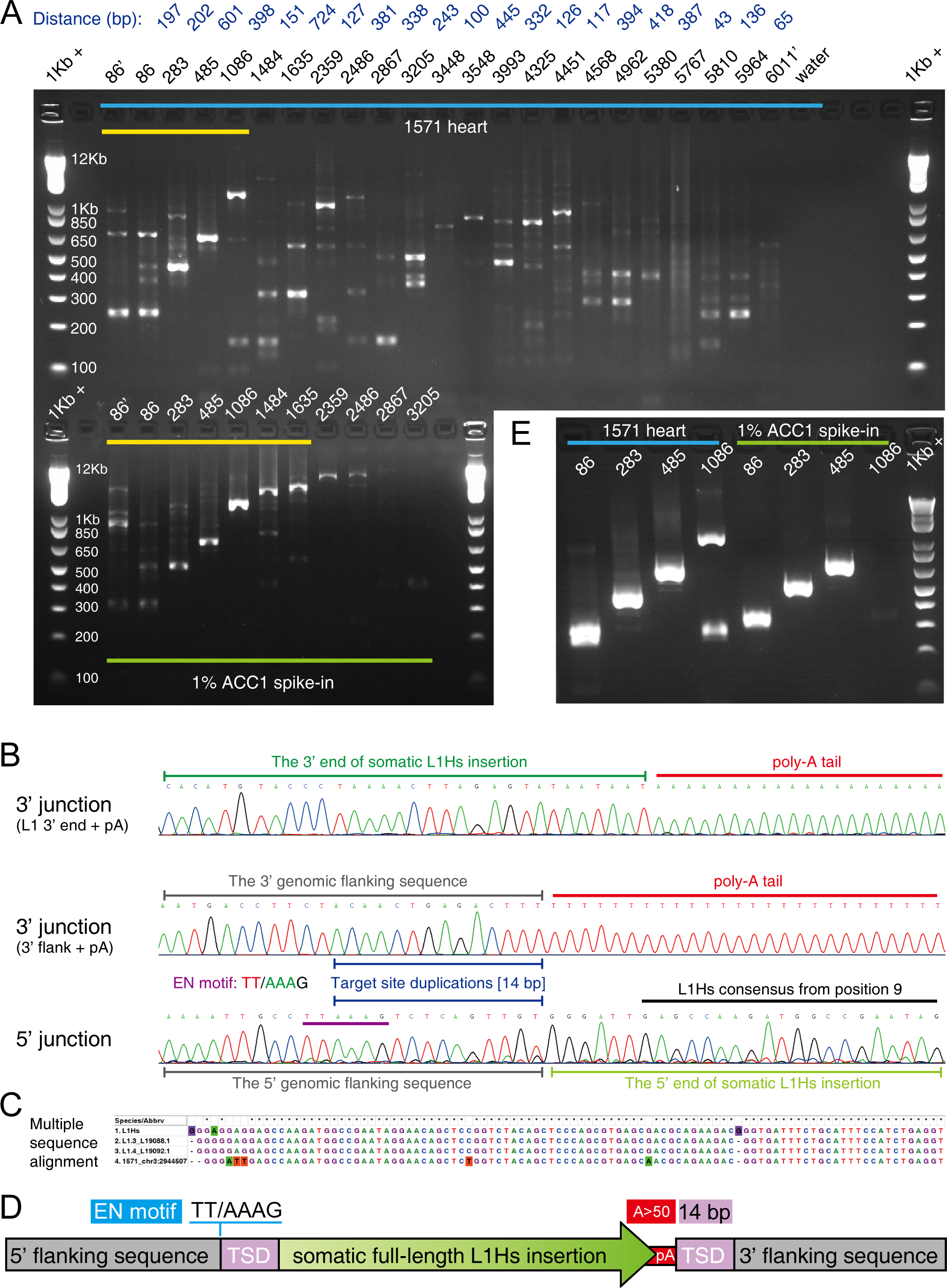
A full-length heart-specific L1Hs insertion (1571_chr3:2944507) in a healthy individual. (A) The agarose gel image of 5’ junction nested PCR validation for the heart-specific L1Hs insertion in the healthy individual (UMB#1571; upper panel). The locations of primers used in 5’ junction PCR assays were labeled on the top of each lane, where primers with the prime symbol denoted semi-nested PCR assays. The distances between each two adjacent 5’ step-wise primers were labeled on the top (dark blue). The lower panel represented a heterozygous, full-length L1Hs insertion (ACC1_16; S11 Table and Appendix 4) in 1% ACC1 spike-in gDNA as the positive control. The yellow line highlighted the expected stair-step bands in 5’ junction PCR. 1Kb +: 1 Kb Plus DNA ladder. (B) The Sanger sequencing chromatograms of the 3’ and 5’ junctions of the somatic insertion 1571_chr3:2944507. The L1 EN motif and TSD were indicated by purple and blue lines. (C) Multiple sequence alignment of the 5’ end between the identified somatic insertion and three L1Hs consensus sequences (L1Hs Repbase consensus and two hot L1s in human [L1.3 and L1.4]). (D) The schematic structure of 1571_chr3:2944507. (E) The agarose gel image of “full-length PCR + 5’ junction PCR” assays for 1571_chr3:2944507 and ACC1_16 positive control.

In addition, we verified one fibroblast-and another heart-specific L1Hs insertion in two patients with Rett syndrome (Fig 3E–F). The heart-specific L1Hs insertion in the Rett patient (UMB#1420) was further resolved to be a highly 5’ truncated L1Hs insertion (∼800 bp) with 9 bp TSD and a cleavage site at 5’–TT/TAAA–3’ (S5 Fig and S11 Table). The poly-A tails of these two clonal somatic insertions were experimentally measured to be polymorphic, indicating that they may involve multiple mutations after the original somatic retrotransposition events (Fig 3I and S10 Table). As previously reported (Evrony *et al.*, 2015; Grandi *et al.*, 2013), poly-A tail was shown to be a highly mutable sequence element and might undergo secondary mutations in descendant cells. Furthermore, we confirmed two additional somatic L1Hs insertions from Rett patient UMB#4516 were present in both PFC neurons, PFC glia, and fibroblasts (Fig 3G–H and S6 Fig), suggesting that they might retrotranspose during early embryonic development. Notably, the intronic somatic insertion (chr20:2392172) in *TGM6* was a full-length L1Hs insertion with 15 bp TSD and a cleavage site at 5’–AT/AAAA–3’ (S7 Fig and S11 Table). We further quantified the allele fractions of this insertion using custom droplet digital PCR (ddPCR) assay and found that 6.34% of fibroblasts and 2.87% of PFC neurons contained this L1Hs insertion (S8A–E Fig and S10 Table). Our observations demonstrated that endogenous L1Hs could retrotranspose in various types of non-brain tissues during human development.

### Abnormal L1Hs mobilization in patients with Rett syndrome

Our HAT-seq bulk sequencing data enabled us to perform statistical analysis of the exonic and intronic patterns of somatic L1Hs insertions in samples from Rett patients and matched healthy controls. We found 180 somatic insertions that were integrated into exonic regions: 9 of which were located in 5’ UTR, 102 of which were located in coding regions, and 69 of which were located in 3’ UTR (S12 Table). While no significant difference was observed in introns (OR = 0.97, p = 0.44, Fisher’s exact test), somatic insertions were significantly depleted in exons (OR = 0.59, p = 6.6×10-4, Fisher’s exact test) of Rett patients compared with matched healthy controls (Fig 5A and S13 Table). Previous studies have shown that dysregulation of long genes (> 100 kb) was linked to neurological disorders, including Rett syndrome (Gabel *et al.*, 2015) and autism spectrum disorder (King *et al.*, 2013). We used our HAT-seq data to investigate somatic insertional bias in both long (> 100 kb) and short genes (< 100 kb) of Rett patients. As a result, we found significant depletion of somatic insertions in exons of long genes (OR = 0.27, p = 5.2×10^-5^, Fisher’s exact test) but not short genes (OR = 0.76, p = 0.12, Fisher’s exact test; Fig 5B and S13 Table). Our speculation was that if an L1Hs inserted into the exonic regions, especially in important genes, of the *MECP2* mutated cell, the cell would have a higher risk of death and subsequently be cleared up; therefore, the observed exonic depletion of L1 insertions in Rett patients might be resulted from the negative selection acting on those “lethal” exonic insertions.

**Fig 5.**
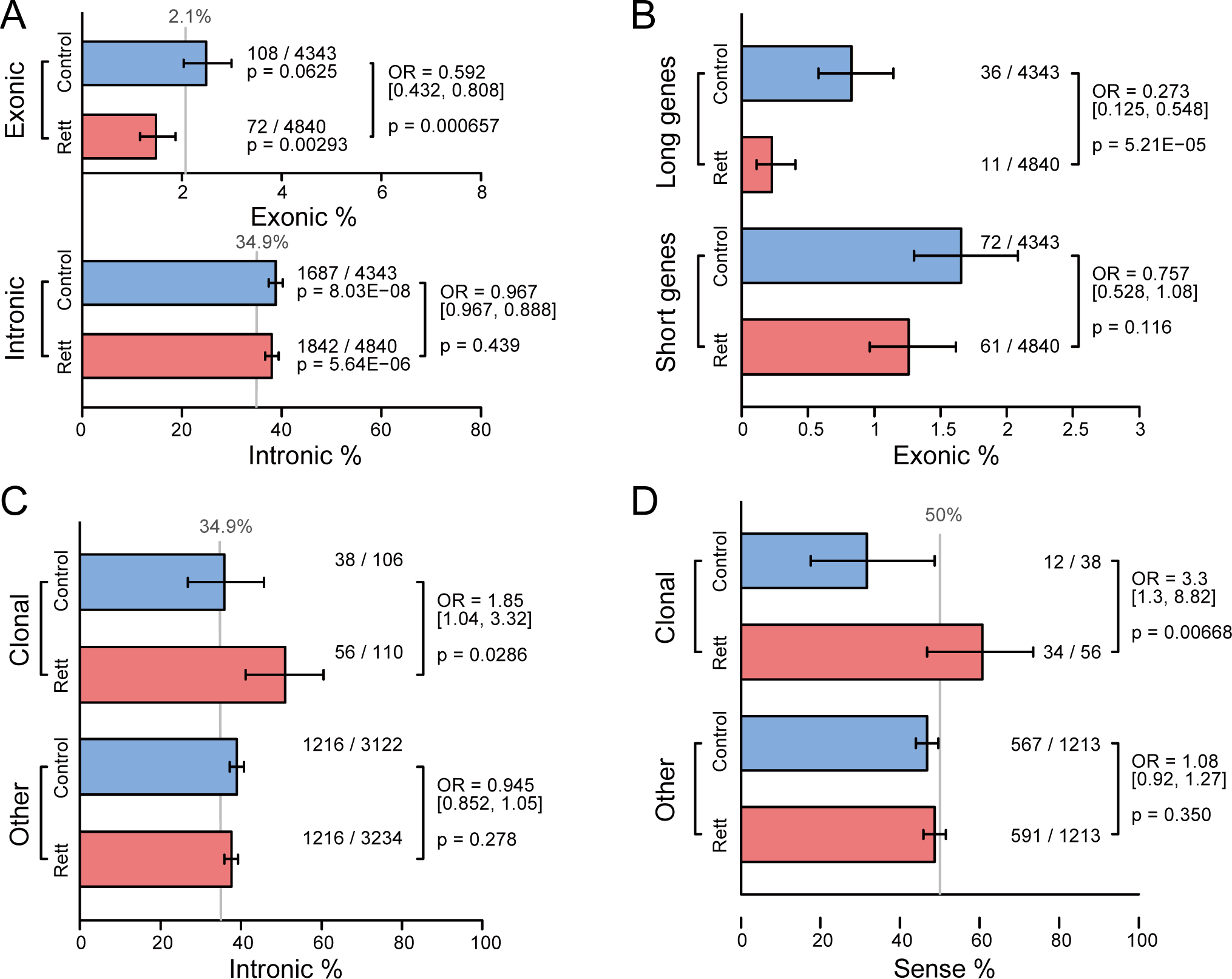
Abnormal L1Hs mobilization in patients with Rett syndrome. (A) Percentages of somatic L1Hs insertions in exons and introns. (B) Percentages of somatic L1Hs insertions in exons of long (> 100 kb) and short genes (< 100 kb). (C) Percentages of clonal somatic L1Hs insertions in introns. (D) Percentages of sense-oriented clonal somatic L1Hs insertions. The gray lines in (A) and (C) denoted the expected proportion determined by the exact base-pair count of that specific region relative to the human genome. The gray line in (D) represented the expected proportion if the insertions occurred randomly in both directions. Error bars in (A)–(D) indicated the 95% confidence intervals.

In contrast to germline insertions, the impact of somatic insertions depends not only on their genomic location, but also the number of cells carrying that insertion, highlighting the importance of clonal somatic insertions. We found that in cortical neurons of Rett patients, clonal somatic insertions were enriched in introns (OR = 1.85, p = 0.029, Fisher’s exact test; Fig 5C and S13 Table); these clonal intronic insertions were significantly enriched in the sense orientation to the transcripts (OR = 3.3, p = 0.0067, Fisher’s exact test; Fig 5D and S13 Table). The presence of L1 insertion in the sense orientation has been reported to interfere with transcriptional elongation of co-localized genes (Han *et al.*, 2004). Considering that clonal insertions are more likely to have occurred at an early stage of development and thus affect a relatively large proportion of cells, these distinct insertion pattern in cortical neurons of Rett patients might indicate potential transcriptional burden on the nervous system.

### Genomic patterns of somatic and germline L1Hs insertions

The design of HAT-seq method allowed for unbiased enrichment of both somatic and germline L1Hs insertions from each of bulk DNA samples. As germline insertion had constant genomic copy number in all tissues from the same donor, we used germline insertion as endogenous control to measure the relative copy number of genome-wide somatic insertions in the brain and non-brain tissues. We quantified the relative somatic L1Hs content by calculating the L1Hs-derived read count ratio of somatic to germline insertions using HAT-seq data of each sample (S14 Table; see details in Materials and Methods). Among all Rett patients and their matched controls, we observed a significant increase in the copy number of somatic L1Hs insertions in PFC neurons relative to matched non-brain tissues (heart, eye, or fibroblast) from the same donor (n = 10, p = 2.7×10^-4^, paired *t*-test; Fig 6A and S8F–G Fig). We also estimated the occurrence rate of somatic L1Hs insertions based on the germline insertion copy number of each individual (Fig 6B). This produced an average of 1.29 [95% CI: 1.03–1.55] somatic insertions per PFC neuron versus 0.60 [95% CI: 0.46–0.74] insertions per non-brain cell (S14 Table). Our observation of higher somatic L1Hs rate in PFC neurons from healthy individuals argued for the active retrotransposition of L1Hs in the human brain (Coufal *et al.*, 2009). One significant advantage of HAT-seq was the ability to distinguish signals of somatic insertions from the overwhelming copies of germline L1Hs insertions in the genome (see details in Materials and Methods). Inconsistent with the previous qPCR result (Muotri *et al.*, 2010), when comparing the group of Rett patients with matched healthy controls, we only observed a slight but not significant increase of somatic L1Hs insertion rate in the Rett group, with 1.36 [range: 0.89–1.82] versus 1.22 [range: 0.63–1.66] per PFC neuron and 0.66 [range: 0.37–1.01] versus 0.54 [range: 0.34–0.72] per non-brain cell (Fig 6C and S14 Table).

**Fig 6.**
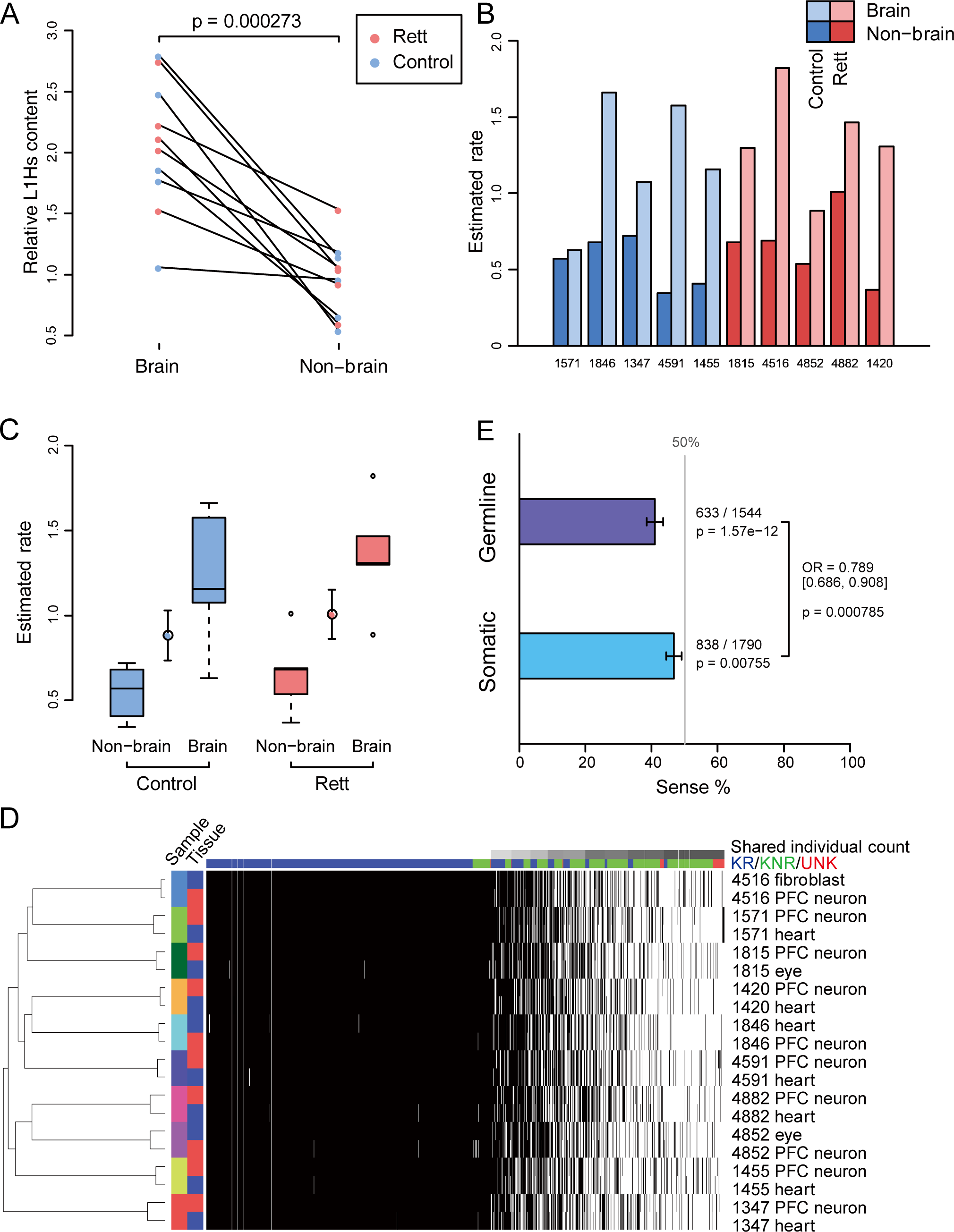
Genome-wide patterns of somatic and germline L1Hs insertions. (A) Relative somatic L1Hs content in PFC neurons and non-brain tissue from the same donor. The read count ratio of somatic insertions to germline KNR was calculated and then normalized relative to the average value of non-brain samples. The linked dots represented pairs of brain and non-brain samples obtained from the same individual. (B) Histogram of estimated rate of somatic L1Hs insertions in each of tissue samples from the same donor based on the germline KNR copy number of each individual. (C) Estimated rate of somatic L1Hs insertions for different tissue types and cohorts. Error bars denoted the standard error of the mean (S.E.M). (D) Hierarchical clustering of all samples sequenced in this study. Each row represented a sample, and each column represents an L1Hs germline insertion. Black and white squares indicated the presence or absence of insertion, respectively. Column annotations showed categories for known reference (KR; blue), known non-reference (KNR; green), and unknown (UNK; red) insertions. (E) Percentages of sense-oriented germline and somatic L1Hs insertions in transcripts. The gray line represented the expected proportion if the insertions occurred randomly in both directions. Error bars indicated the 95% confidence intervals.

We next characterized the genome-wide germline L1Hs insertions. HAT-seq yielded greater than 320-fold enrichment for KR, KNR, and UNK L1Hs insertions (S15 Table). On average, 814 KRs, 183 KNRs, and 10 UNKs were identified in each bulk sample (Table 2, S5–7 Table). Hierarchical clustering based on L1Hs profiles correctly paired all neuronal samples with the non-neuronal tissue samples of the same individual (Fig 6D). To experimentally validate the HAT-seq predicted germline insertions, we performed 3’ PCR validation on a random subset of polymorphic insertions from among the ten individuals, including 8 sites out of 160 polymorphic KRs, 20 sites out of 451 KNRs, and 2 sites out of 48 UNKs (S7 and S16 Table). As a result, all of the assayed sites were detected in 3’ PCR, with 98.4% (120/122) and 100% (168/168) sensitivity and specificity, respectively (S16 Table and Appendix 5). These results support that HAT-seq can reliably detect germline L1Hs insertions with high sensitivity and specificity.

Previous studies have shown that intronic germline L1Hs insertions are sense-depleted (Ewing and Kazazian, 2010; Smit, 1999; Upton *et al.*, 2015). As expected, the germline insertions identified in this study were significantly sense-depleted to the transcripts (633/1,544 [41%], p = 1.6×10^-12^, binomial test; Fig 6E and S13 Table). It is important to ask the question: whether such orientation bias for germline insertions is resulted from natural selection or insertional preference? To address this, we chose somatic L1Hs insertions as internal reference to control confounding factors. We compared the orientation bias between germline and somatic L1Hs insertions in transcripts and found that germline insertions were significantly sense-depleted than somatic insertions (odds ratio [OR] = 0.79, p = 7.9×10^-4^, Fisher’s exact test; Fig 6E and S13 Table). Because somatic L1Hs insertions only affected a small proportion of cells and thus they should undergo weaker selective pressure than germline insertions, our results suggested that natural selection may play a major role in shaping the sense-depleted distribution of germline L1Hs insertions.

## Discussion

Here, we present HAT-seq, a bulk DNA sequencing method to profile genome-wide L1Hs insertions from physiologically normal and pathological human tissues. We demonstrated that, in addition to neuronal cells (Erwin *et al.*, 2016; Evrony *et al.*, 2012; Evrony *et al.*, 2015; Macia *et al.*, 2017; Upton *et al.*, 2015), L1Hs also retrotransposed in a variety of non-brain tissues and cell types during normal development and contributed to the inter-cellular diversity of the human genome. Using high-throughput sequencing-based quantitative analysis, we found that somatic insertions occurred at a higher rate in brain than in non-brain tissues, consistent with previous studies (Coufal *et al.*, 2009).

Previous qPCR and single-cell genomic studies have resulted in conflicting estimates of the frequency of somatic insertions in neurons: ∼80 L1 insertions per neuron (Coufal *et al.*, 2009), < 0.04–0.6 L1 insertions per neuron (Evrony *et al.*, 2012), 13.7 L1 insertions per neuron (Upton *et al.*, 2015), or ∼0.58–1 somatic L1-associated variants per neuron (Erwin *et al.*, 2016). Differential estimates might result from differences in WGA and signal enrichment methods. Using a bulk DNA sequencing approach, we estimated the rate of somatic insertions to be 0.63–1.66 L1Hs insertions per PFC neuron in healthy individuals (Fig 6B and S14 Table).

Clonally distributed insertions are prevalent in normal brain (Evrony *et al.*, 2015). Increasing evidence suggests that neuronal L1s retrotransposition contributes to the susceptibility to and pathophysiology of neurological disorders, including Rett syndrome (Muotri *et al.*, 2010), schizophrenia (Bundo *et al.*, 2014) and Alzheimer’s disease (Guo *et al.*, 2018). We observed that, in PFC neurons of Rett patients, clonal somatic insertions were enriched in introns, and these clonal intronic insertions were significantly enriched in the sense orientation (Fig 5C–D). In particular, in Rett patient UMB#4516, we found a full-length, sense-orientated, intronic somatic insertion (chr20:2392172) in *TGM6* (S7 Fig and S11 Table), a gene associated with central nervous system development and motor function (Thomas *et al.*, 2013), which could potentially dysregulate gene expression (Han *et al.*, 2004). We found that 6.34% of fibroblasts and 2.87% of PFC neurons contained this insertion (S8A–E Fig and S10 Table), suggesting that it might occur in the 16-cell or 32-cell stages during morula stage. Mutations in *TGM6* are associated with spinocerebellar ataxia type 35, one of a group of genetic disorders characterized by poor coordination of hands, gait, speech, and eye movements as well as frequent atrophy of the cerebellum (Guo *et al.*, 2014; Li *et al.*, 2013; Wang *et al.*, 2010). According to the clinical records, UMB#4516 had slight cerebral atrophy and cerebellar degeneration, could not hold things in her hands, and her speech development ceased at 16 months of age; these phenotypes were absent in the other four patients with Rett syndrome. Taken together, our data indicated that this clonal L1Hs insertion of *TGM6* might be correlated with the distinct clinical phenotype of UMB#4516.

Previous studies have provided evidence for significant selection against older L1 elements that are non-polymorphic (Boissinot *et al.*, 2001; Ewing and Kazazian, 2010). To characterize the insertion pattern of L1 with minimal influence from selective pressure, experimental methods were developed for recovery of novel L1 insertions in cultured cells (Gilbert *et al.*, 2005; Symer *et al.*, 2002). Using HAT-seq method, we were able to distinguish somatic L1Hs insertions from germline L1Hs insertions within the same individual. To determine whether the sense-depleted germline insertion was resulted from natural selection or insertional preference, we used somatic insertion as internal reference to control confounding factors such as intrinsic insertion preference and compared germline with somatic insertions. Our results suggested that natural selection shaped a sense-depleted distribution of germline L1Hs insertions in the human genome.

Several PCR-based bulk sequencing methods, such as ATLAS (Badge *et al.*, 2003), L1-seq (Ewing and Kazazian, 2010), TIP-seq (Tang *et al.*, 2017), bulk SLAV-seq (Erwin *et al.*, 2016), and ATLAS-seq (Philippe and Cristofari, 2016), have been developed to identify germline L1Hs insertions. Furthermore, L1-seq and TIP-seq have been successfully used in the identification of somatic insertions in tumors (Achanta *et al.*, 2016; Doucet-O’Hare *et al.*, 2015; Ewing *et al.*, 2015; Solyom *et al.*, 2012; Tang *et al.*, 2017). Due to clonal expansion during tumorigenesis, such insertions could affect numerous cells in tumors. To our knowledge, HAT-seq is the first PCR-based bulk sequencing method to identify rare somatic insertions in a subset of cells—even unique cells—in non-tumor tissues. HAT-seq provides not only the genomic positions of somatic insertions but also the allele fraction of each insertion, which is informative for inferring the timing when the insertion has occurred. The sensitivities of HAT-seq for low-frequency somatic L1Hs insertions were relatively low (∼30% for insertions present in < 1% fraction of cells). One possible explanation was that some signals of insertion were lost during library construction and NGS sequencing, e.g. sonic fragmentation, clean-ups, size selection, and loading library to sequencer. Single-cell whole genome and targeted sequencing approaches have been used to identify both TPRT-mediated and endonuclease-independent insertions (Erwin *et al.*, 2016; Evrony *et al.*, 2012; Evrony *et al.*, 2015; Upton *et al.*, 2015), where the signal of somatic insertions can be as high as germline heterozygous insertions in single-cell level. However, such single-cell approaches cannot achieve increased sensitivity without cost (Evrony *et al.*, 2016). For example, to detect a given insertion with 0.1% mosaicism, more than 1,000 single cells may need to be amplified and sequenced. Therefore, compared with single-cell approaches, HAT-seq was eligible to identify a large number of somatic L1Hs insertions in a more cost-effective way.

Based on our experimental design, assembling overlapped read pairs into contigs can provide sequence information fully spanning the L1Hs integration sites, enabling downstream false-positive filtering based on both sequence features and read-count. However, a portion of read pairs were unable to be merged into contigs because the inaccurate size-selection during library construction. Applying the same filtering strategy, we re-analyzed these unassembled read pairs and revealed 11 clonal insertion candidates. Further PCR experiments only validate one of these candidates (9%, S10 Table). Because the key filter “chimera within poly-A tail” was not applicable for unassembled read pairs, our sequence analysis suggested that chimeric molecules bridging within the poly-A tail was the major source of false-positives for unassembled data (see details in Materials and Methods). As shown in the statistics of positive control libraries (S2 Table) and experimental validation, the unassembled data could provide additional signals of somatic L1Hs insertions but require careful analysis and rigorous validation to address technical artifacts. Further gains in statistical power will be benefited from increased sample size and improved efficiency of HAT-seq.

Several unresolved technical challenges might constrain the total number of detectable L1Hs insertions by the current version of HAT-seq, including the identification of insertions in repetitive regions with low mappability (such as pre-existing L1 germline insertions) and 3’ truncated insertions. With rapid innovations in sequencing technology, higher throughput and longer read length will markedly improve the performance of HAT-seq. Future studies that profile all active retrotransposons (i.e., L1Hs, Alu, and SVA) in a variety of cell types, tissues, and developmental stages will shed new light on the dynamics of somatic retrotransposition under host regulation and help to uncover their roles in human disease.

## Supporting information

Appendix 1. Cell type-specific sorting for postmortem human brain samples

Appendix 2. Identification of ACC1-specific insertions and their zygosity

Appendix 3. Detection of somatic insertions in positive control experiments

Appendix 4. Benchmarking PCR validation assays for low-frequency somatic insertions

Appendix 5. Experimental validation of polymorphic germline L1Hs insertions

## Acknowledgements

We acknowledge the UMB Brain and Tissue Bank (University of Maryland, Baltimore, MD) and the Lieber Institute for Brain Development (Baltimore, MD) for providing postmortem human tissues. We are grateful to Drs Eunjung Alice Lee, Daniel R. Weinberger, Li-Lin Du, Meng-Qiu Dong, Yu Zhang, Ge Gao, Louis Tao, Cheng Li, Jian Lu, and Manyuan Long for their insightful comments and suggestions. We thank Drs Kazuya Iwamoto and Miki Bundo for providing detailed protocols of nuclei isolation. We thank Dr. Timothy W. Yu for providing experimental resources in the revision. We thank the reviewers for constructive feedback on the manuscript. This study was supported by the National Natural Science Foundation of China (31530092) and the Ministry of Science and Technology 863 Grant (2015AA020108). Q-R.L. was supported in part by the Intramural Research Program at the National Institute on Aging.

## Author contributions

Adam Yongxin Ye

Formal analysis, Methodology, Visualization, Writing-original draft

August Yue Huang

Conceptualization, Resources, Supervision, Writing—review and editing

Boxun Zhao

Conceptualization, Data curation, Formal analysis, Investigation, Methodology, Project administration, Validation, Visualization, Writing—original draft, Writing—review and editing

Jing Guo

Data curation, Methodology, Validation

Liping Wei

Conceptualization, Funding acquisition, Resources, Supervision, Writing—review and editing

Linlin Yan Formal analysis

Thomas M. Hyde Resources

Qing-Rong Liu Methodology

Qixi Wu

Conceptualization, Investigation, Methodology, Writing—review and editing

Xianing Zheng Methodology

Xiaoxu Yang

Validation, Visualization

## Supporting Information

### Supplementary Figures

**S1 Fig.**
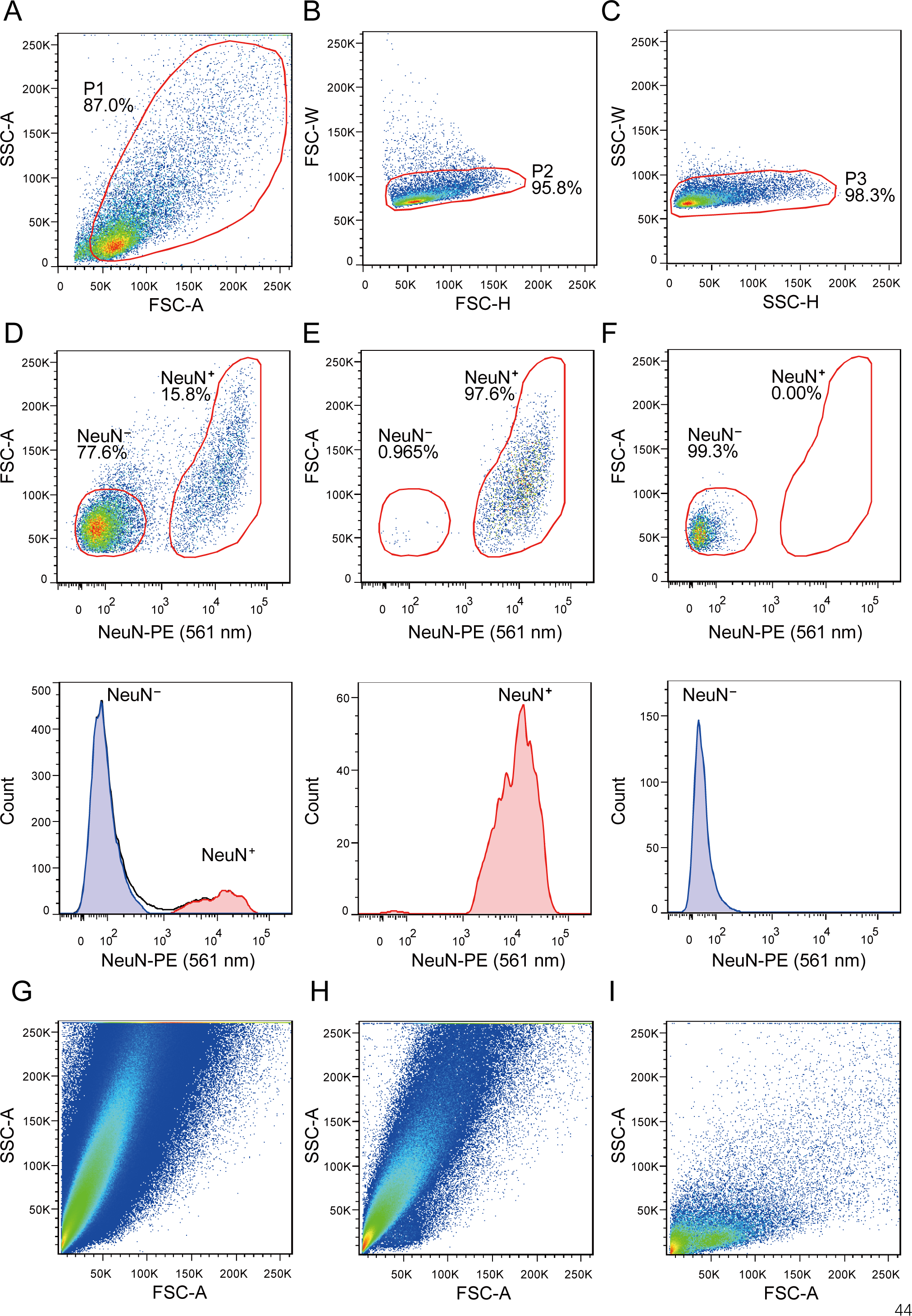
Nuclei isolation and NeuN^+^ fluorescence-activated cell sorting (FACS). (A)–(D) Purify neuronal nuclei from human PFC. (A) The first gate (P1) was set as an FSC-A vs. SSC-A plot to discriminate the population containing small-size debris. (B)–(C) The second (P2) and third (P3) gates were set as FSC-H vs. FSC-W and SSC-H vs. SSC-D plots, respectively, to remove doublets and clumps. (D) Top: NeuN^−^ and NeuN^+^ gates were set in the NeuN-PE (561 nm) vs. FSC-A plot. Bottom: a count plot of NeuN-stained nuclei. (E) Purity analysis of sorted neurons. Top: sorted NeuN^+^ nuclei were re-analyzed by FACS to confirm the sort purity. Bottom: a count plot of re-analyzed NeuN^+^ nuclei. (F) Purity analysis of sorted glia. Top: sorted NeuN^−^ nuclei were re-analyzed by FACS to confirm the sort purity. Bottom: a count plot of re-analyzed NeuN^−^ nuclei. (G) FSC vs. SSC plot of brain homogenate. Brain homogenate contained a huge amount of cell debris and myelin debris. (H) FSC vs. SSC plot of debris-detached single-nuclei homogenate. Minced brain tissue was soaked overnight before homogenization and then incubated with nonionic detergent, Nonidet P-40, to remove cell debris from nuclear membrane. (I) FSC vs. SSC plot of debris removed nuclei fraction. Cell debris and myelin were separated from nuclei using Percoll density gradient centrifugation.

**S2 Fig.**
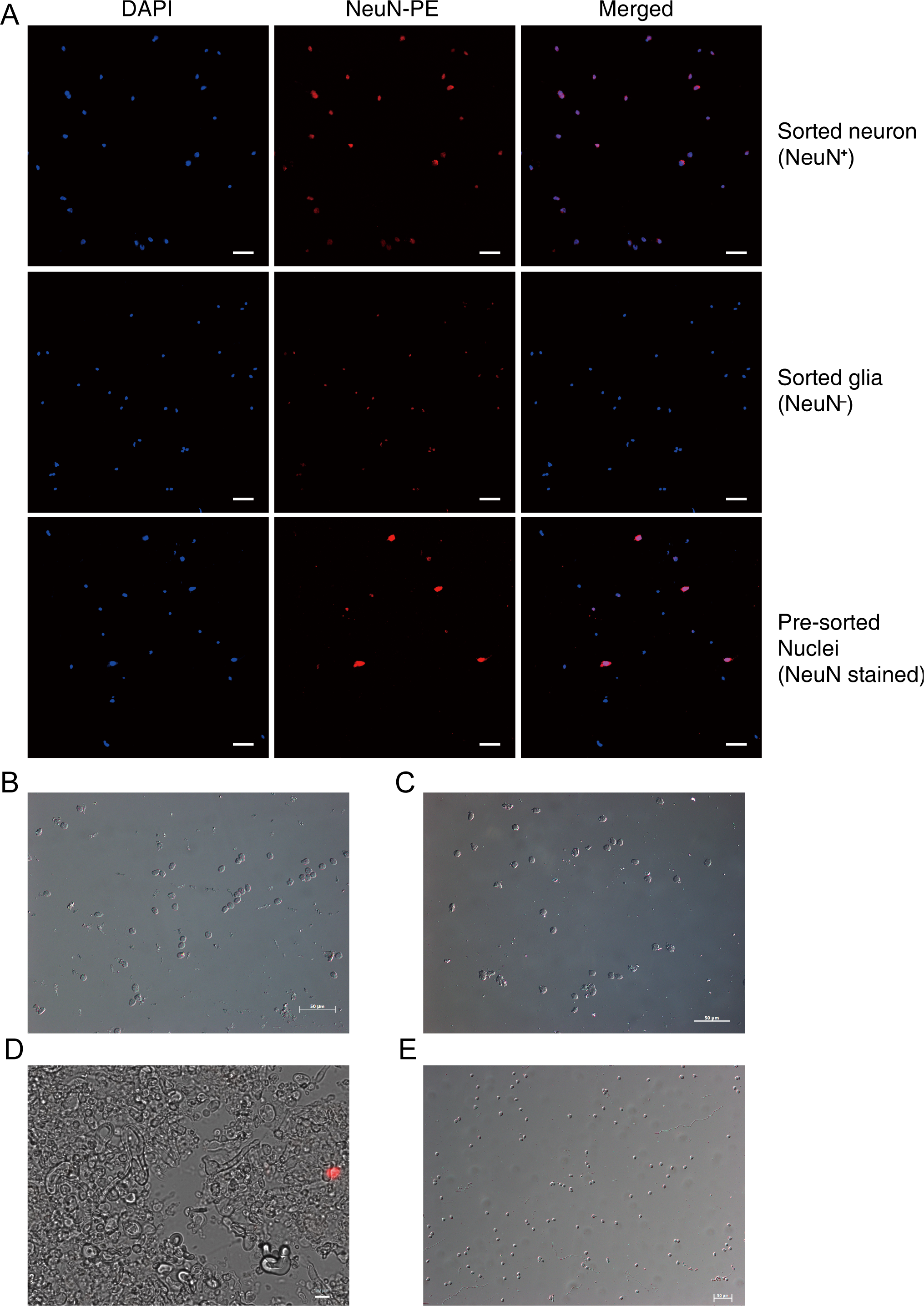
Confirmation of NeuN^+^ FACS purity and integrity. (A) Example of fluorescence microscopy confirmation of isolated nuclei. The purity of each fraction was > 95% for NeuN^+^ and NeuN^−^ nuclei. Bar = 50 μm. (B)–(C) Examples of integrity confirmation using differential interference contrast (DIC) of sorted neurons (B) and glia (C). Bar = 50 μm. (D) Example of the myelin, lipid, and cell debris layers (12% Percoll) after Percoll density gradient centrifugation. Nuclei were stained with a red fluorescent nuclear counterstain, propidium iodide (PI). Bar = 20 μm. (E) Example of the nuclei fraction layer (35% Percoll) after Percoll density gradient centrifugation. Bar = 50 μm. DAPI, 4’,6-diamidino-2-phenylindole; NeuN-PE, PE-conjugated anti-NeuN antibody.

**S3 Fig.**
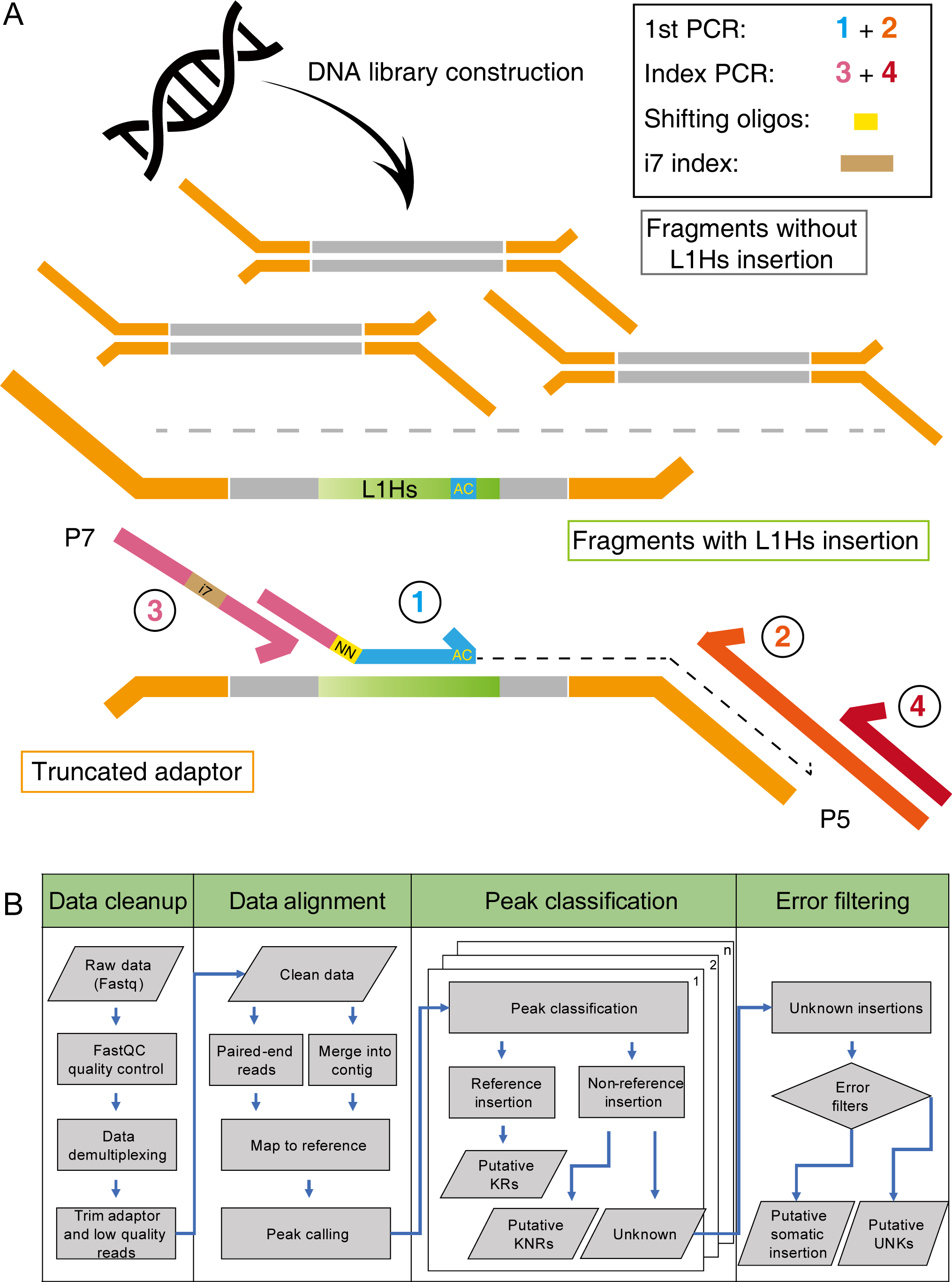
Schematic diagrams of HAT-seq library construction and computational analysis pipeline. (A) Schematic of the HAT-seq library construction. The fragmented genomic DNA was ligated with P7 truncated adaptors, and then used as template for L1Hs amplification PCR. Primers 1 (P7_Ns_L1Hs) was specific to L1Hs diagnostic ‘‘AC’’ motif. See S1 Table for primer sequences. (B) Schematic of the HAT-seq data analysis pipeline; full details are provided in the Materials and Methods.

**S4 Fig.**
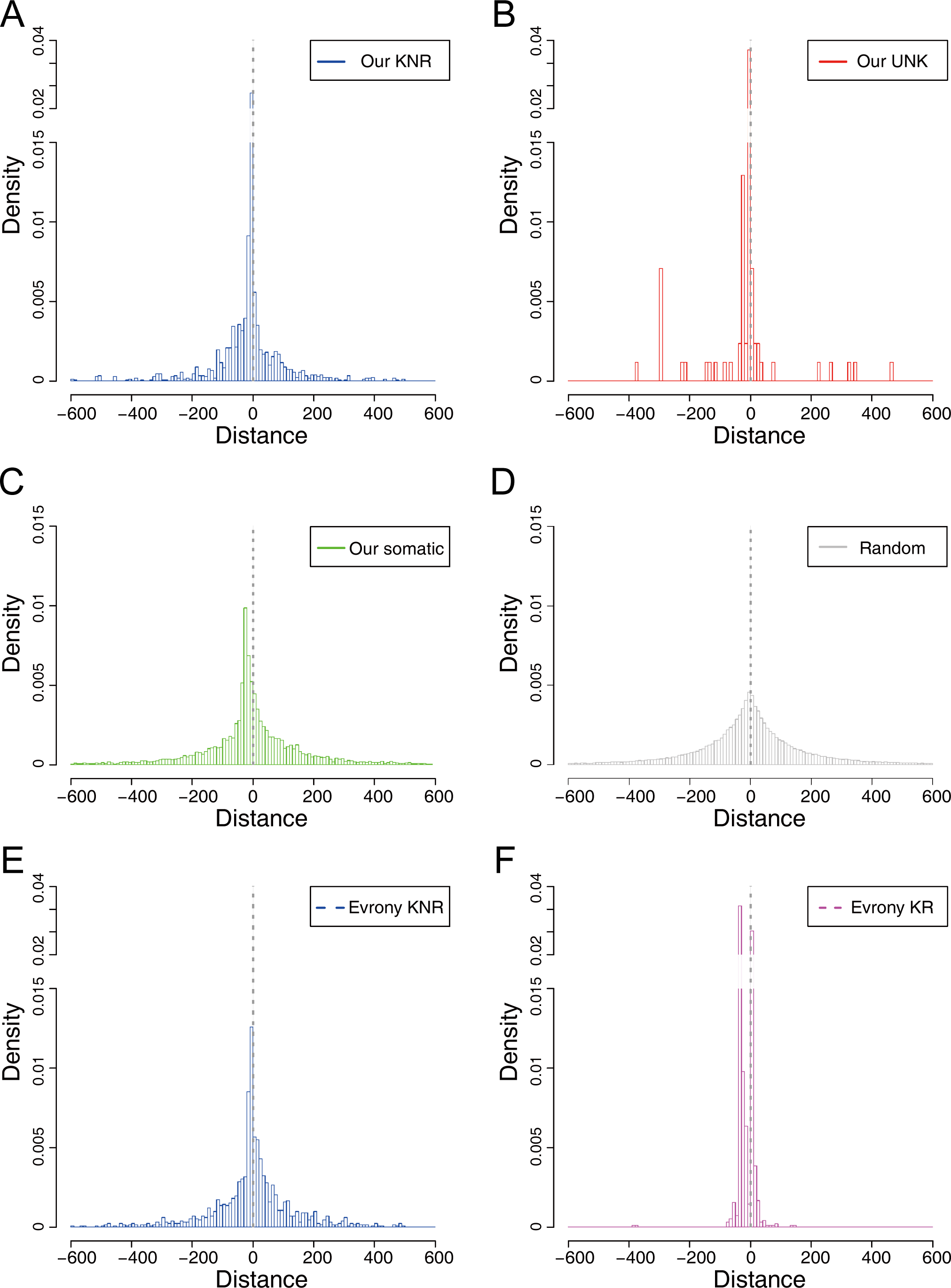
EN motif enrichment analysis across all categories of L1Hs insertions. The density distributions of L1 EN motifs around germline KNR (A), UNK (B), somatic insertions (C), randomly sampled positions (D), “Evrony KR” (E), and “Evrony KNR” (F). The lists of “Evrony KR” and “Evrony KNR” were extracted from Evrony *et al*. 2012. The bin size of histogram was 10 bp. L1 EN motifs included seven specific motifs (TTAAAA, TTAAGA, TTAGAA, TTGAAA, TTAAAG, CTAAAA, TCAAAA).

**S5 Fig.**
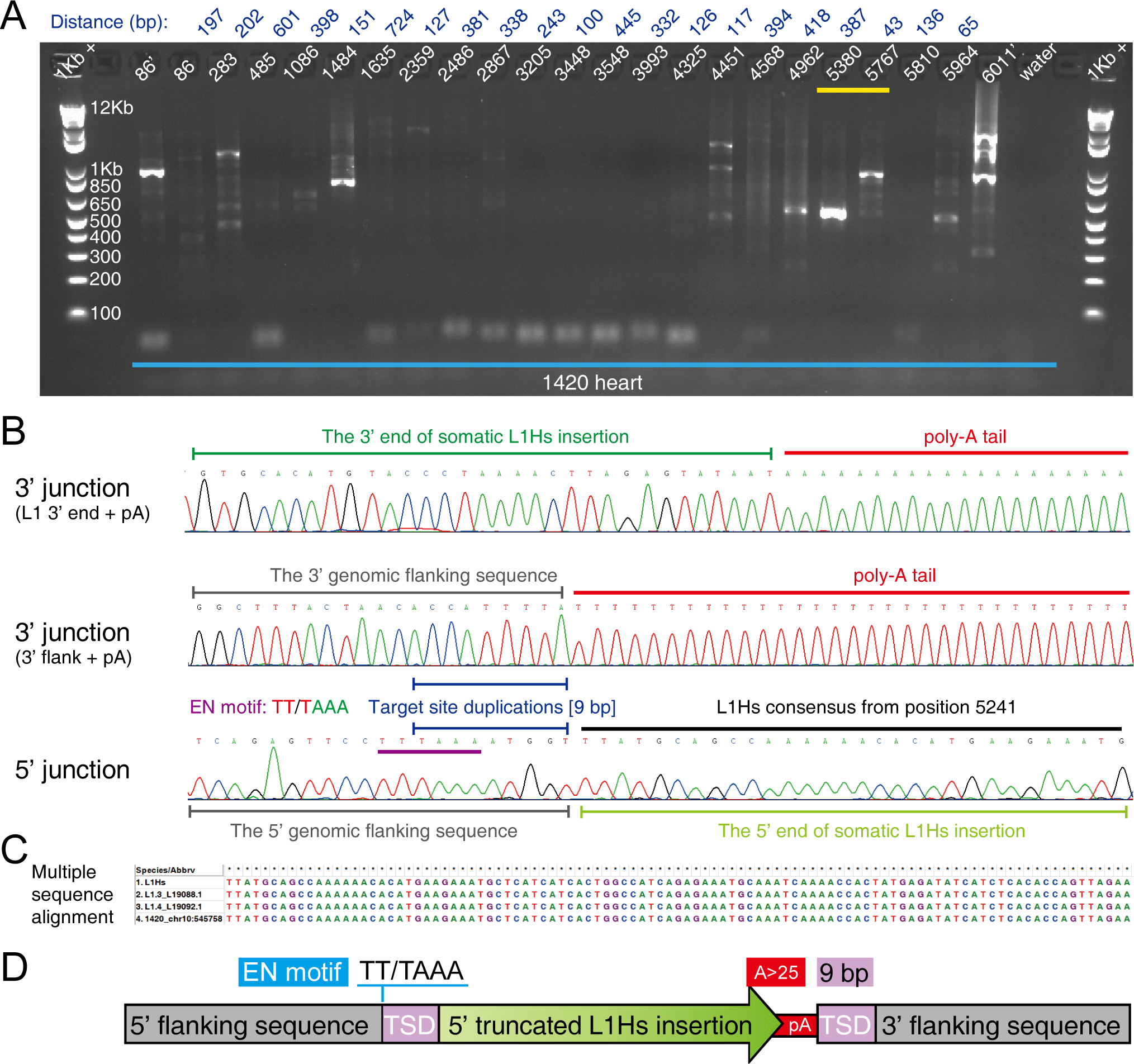
A 5’ truncated heart-specific L1Hs insertion (1420_chr10:545758) in a Rett patient. (A) The agarose gel image of 5’ junction nested PCR validation for the heart-specific L1Hs insertion in the Rett patient (UMB#1420). The locations of primers used in 5’ junction PCR assays were labeled on the top of each lane, where primers with the prime symbol denoted semi-nested PCR assays. The distances between each two adjacent 5’ step-wise primers were labeled on the top (dark blue). The yellow line highlighted the expected stair-step bands in 5’ junction PCR. 1Kb +: 1 Kb Plus DNA ladder. (A) The Sanger sequencing chromatograms of the 3’ and 5’ junctions of the somatic insertion (1420_chr10:545758). The L1 EN motif and TSD were indicated by purple and blue lines. (C) Multiple sequence alignment of the 5’ end between the identified somatic insertion and three L1Hs consensus sequences (L1Hs Repbase consensus and two hot L1s in human [L1.3 and L1.4]). (D) The schematic structure of the highly 5’ truncated (∼800 bp) L1Hs insertion 1420_chr10:545758.

**S6 Fig.**
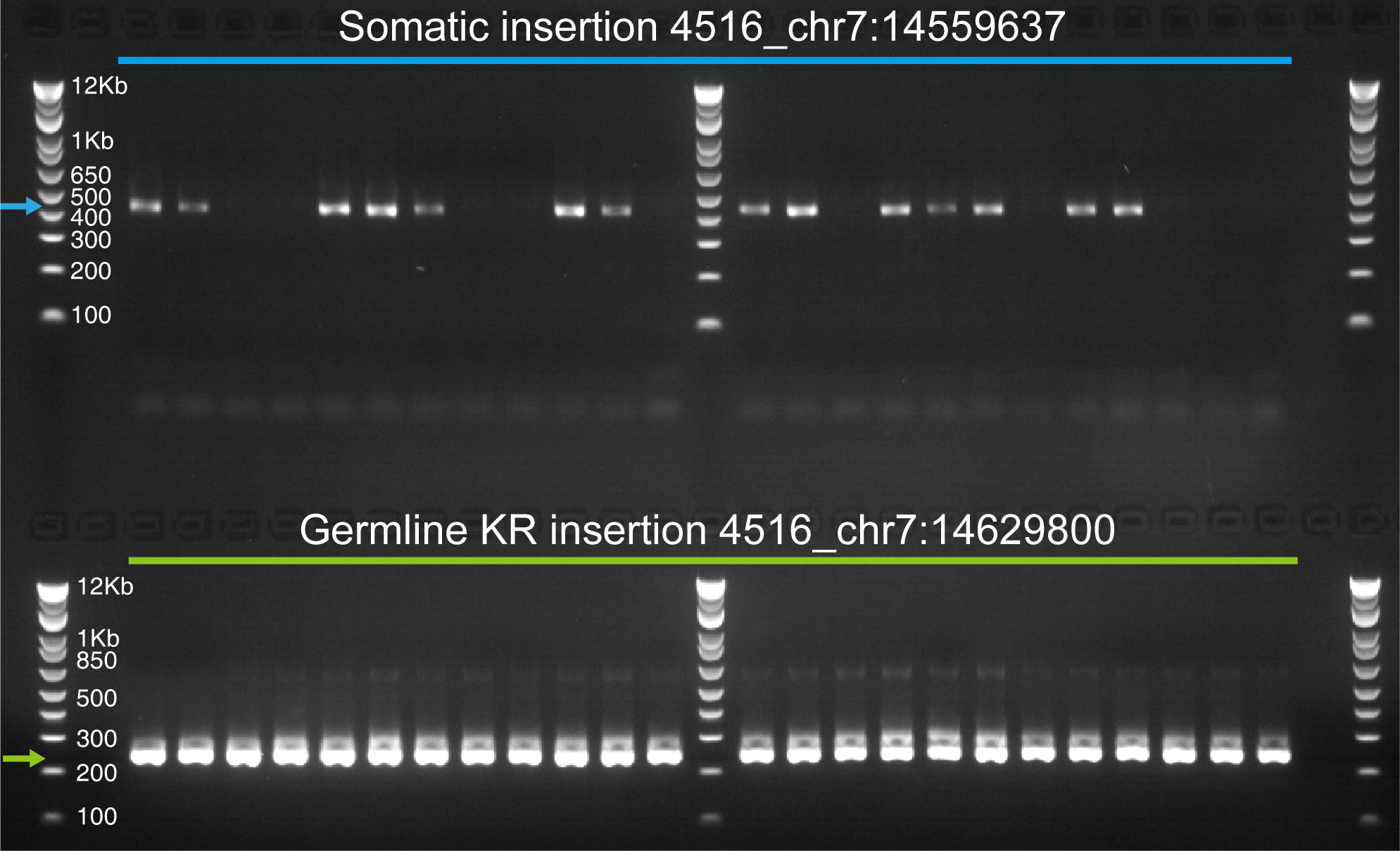
The somatic status of an L1Hs insertions (4516_chr7:14559637) in a Rett patient. The somatic insertion 4516_chr7:14559637 was present in 14 out of 24 nested 3’ PCR wells, compared to 24 out of 24 wells for a germline KR insertion (chr7:14629800) from the same donor. DNA sample was diluted to ∼300 cells per well. Blue and green arrows indicated bands with target size.

**S7 Fig.**
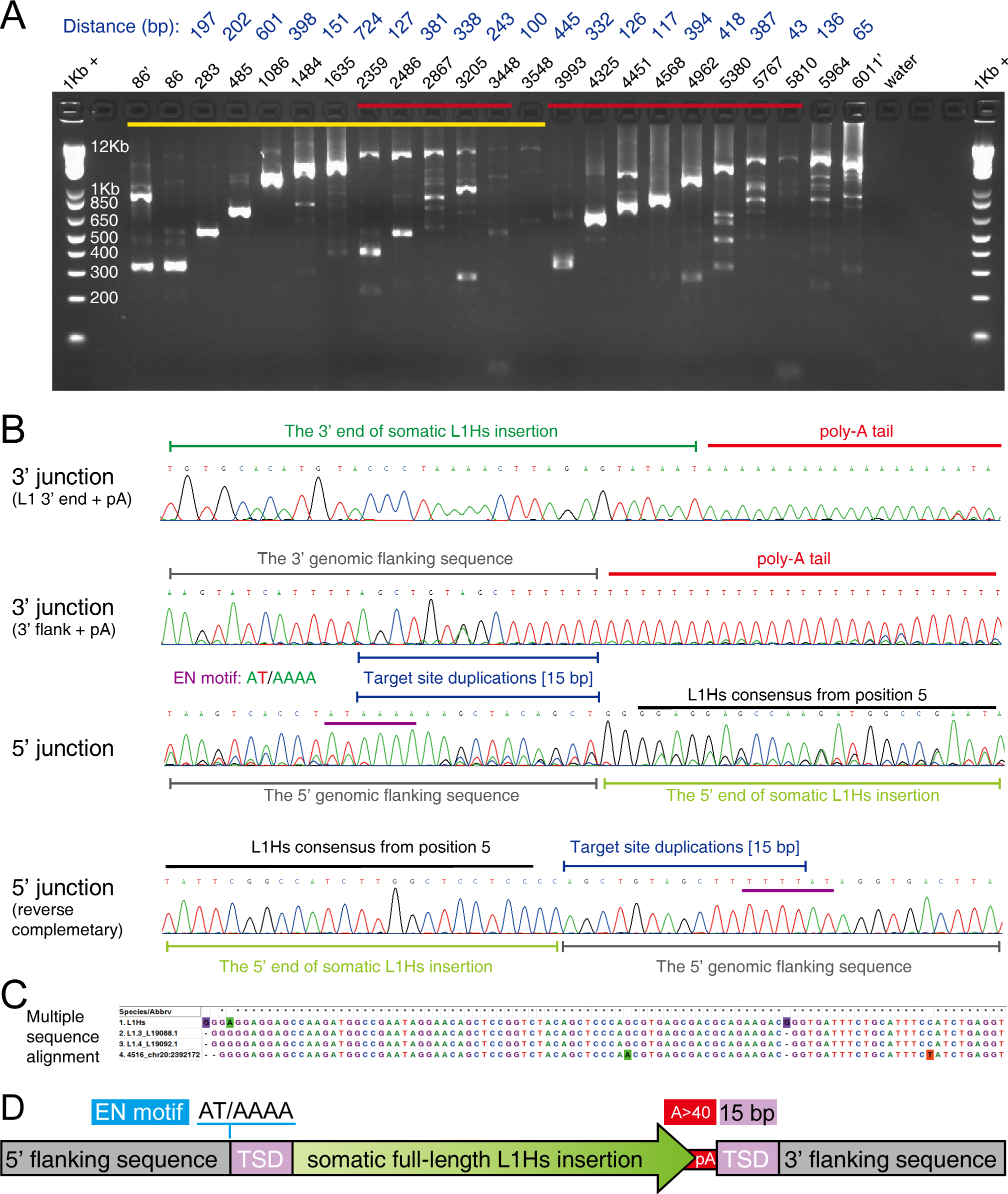
A full-length embryonic somatic L1Hs insertion (4516_chr20:2392172) in a Rett patient. (A) The agarose gel image of 5’ junction nested PCR validation for the embryonic somatic L1Hs insertion (4516_chr20:2392172) in the Rett patient (UMB#4516). The locations of primers used in 5’ junction PCR assays were labeled on the top of each lane. Step-wise primers with the prime symbol were used twice in semi-nested PCR assays. The distances between each primer pairs were labeled on the top (dark blue). The yellow line highlighted the expected stair-step bands in 5’ junction PCR, while the red lines indicated false positives resulted from non-specific amplification of L1PA subfamilies. 1Kb +: 1 Kb Plus DNA ladder. (B) The Sanger sequencing chromatograms of the 3’ and 5’ junctions of somatic insertion (4516_chr20:2392172). The nucleotides shifted chromatogram in 5’ junction might result from the DNA polymerase slippage at homopolymers in the upstream region (L1MB3 element), and its sequence was confirmed from the reverse direction. The L1 EN motif and TSD were indicated by purple and blue lines. (C) Multiple sequence alignment of the 5’ end between the identified somatic insertion and three L1Hs consensus sequences (L1Hs Repbase consensus and two hot L1s in human [L1.3 and L1.4]). (D) The schematic structure of 4516_chr20:2392172.

**S8 Fig.**
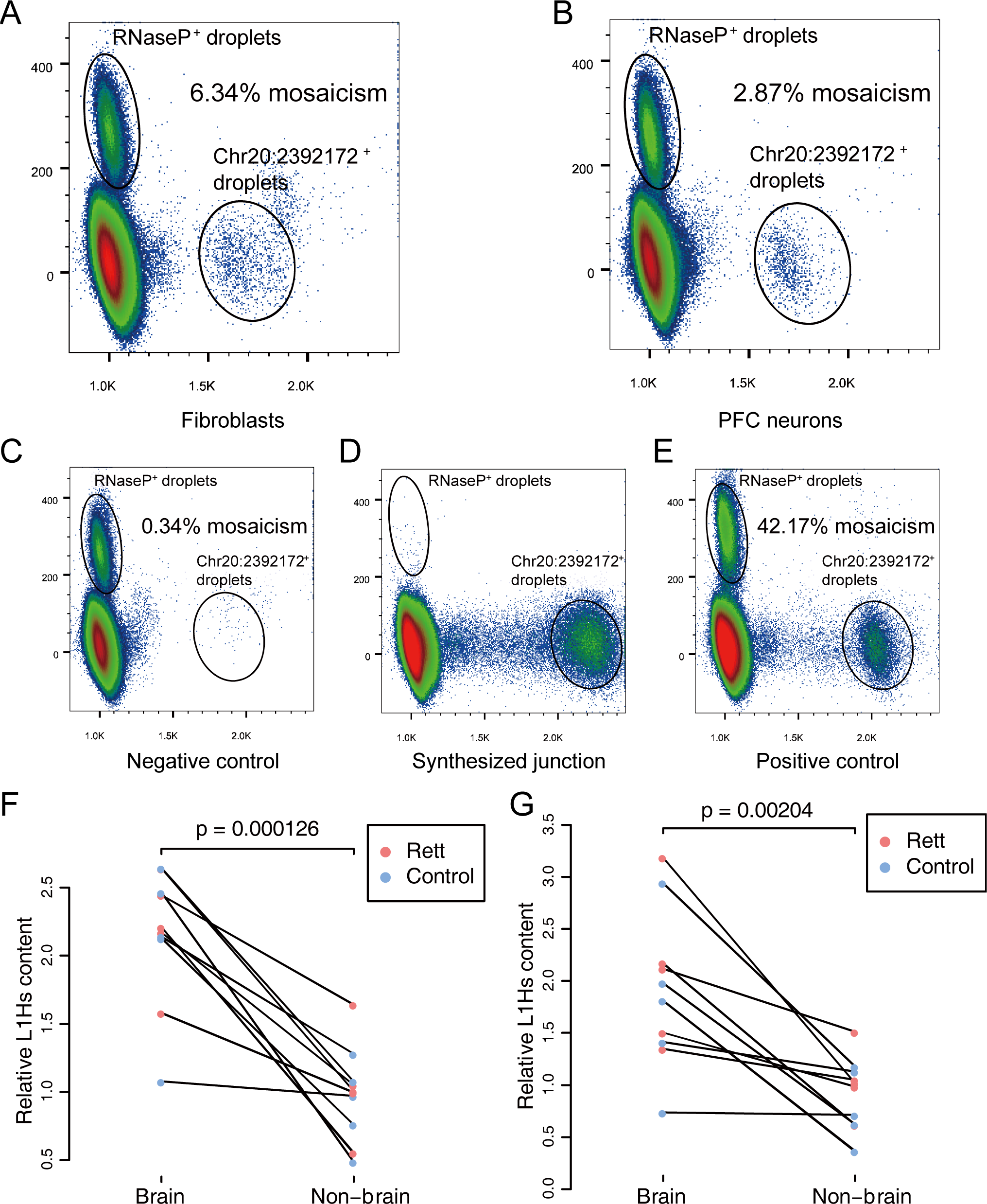
Qualitative and quantitative analysis of L1Hs insertions. (A)–(E) Droplet digital PCR (ddPCR) assays to quantify mosaicism (percentage of cells) of somatic L1Hs insertions at chr20:2392172 in fibroblasts (A) and PFC neurons (B) from Rett patient UMB#4516. Fragmented ACC1 blood gDNA was used as template for negative control assay (C). A mixed template containing fragmented ACC1 blood gDNA and diluted synthesized L1Hs genome junction oligos (D) was used for positive control assay (E). RNaseP served as a genomic copy number reference (copy number = 2). L1Hs and RNaseP assays were labeled with FAM and VIC, respectively. (F)–(G) Relative somatic L1Hs content in PFC neurons and non-brain tissue from the same donor, normalized by the read count of KRs (F) or UNKs (G) from the same tissue sample.

### Supplementary Tables

S1 Table. Primer sequences of HAT-seq library

S2 Table. ACC1-specific insertions in positive control experiments

S3 Table. Statistics of error filters in positive control experiments

S4 Table. Clinical characterization of patients with Rett syndrome

S5 Table. Statistics of HAT-seq libraries

S6 Table. Statistics of known reference insertions among all samples

S7 Table. Statistics of polymorphic insertions among all samples

S8 Table. Statistics of somatic insertions among all samples

S9 Table. TPRT hallmark annotation for all somatic insertions

S10 Table. 3’ junction nested PCR and digital droplet PCR validation

S11 Table. 5’ junction nested PCR validation

S12 Table. Annotation for somatic exonic insertions

S13 Table. Raw data for statistical analyses

S14 Table. Quantification statistics among all samples

S15 Table. L1Hs enrichment analysis on HAT-seq

S16 Table. 3’ junction PCR validation for germline insertions

### Supplementary Files

Appendix 1. Cell type-specific sorting for postmortem human brain samples

Appendix 2. Identification of ACC1-specific insertions and their zygosity

Appendix 3. Detection of somatic insertions in positive control experiments

Appendix 4. Benchmarking PCR validation assays for low-frequency somatic insertions

Appendix 5. Experimental validation of polymorphic germline L1Hs insertions

