## Appendix 1. Cell type-specific sorting for postmortem human brain samples for "Somatic LINE-1 retrotransposition in cortical neurons and non-brain tissues of Rett patients and healthy individuals"

**Appendix 1. Cell type-specific sorting by BD Aria II using postmortem human brain sample.**

An overview of cell type-specific sorting using postmortem samples of prefrontal cortex.

| UMB_ID | Category | Date | Weight (mg) | NeuN$^+$ | NeuN$^-$ | NeuN$^+$ gDNA (ng) | NeuN$^-$ gDNA (ng) | Purity (%) |
|---|---|---|---|---|---|---|---|---|
| 1815 | Rett | 20141112 | 32 | 206,157 | 359,601 | 486 | 382 | 99.7 |
| 1571 | control | 20141113 | 97 | 607,553 | 1,454,375 | 1,378 | 2,587 | 99.6 |
| 4516 | Rett | 20141120 | 125 | 1,447,909 | 8,068,812 | 3,666 | 22,100 | 99 |
| 1846 | control | 20141119 | 77 | 1,073,946 | 1,226,869 | 3,042 | 2,015 | 99.2 |
| 4852 | Rett | 20141029 | 104 | 32,136 | 5,297,495 | 101 | 2,756 | NA |
| 1347 | control | 20141030 | 45 | 404,093 | 1,135,614 | 1,144 | 1,128 | 99.6 |
| 4882 | Rett | 20141104 | 67 | 1,100,590 | 4,426,782 | 928 | 1,404 | 94.8 |
| 4591 | control | 20141204 | 54 | NA | NA | 1,755 | 1,173 | 99.1 |
| 1420 | Rett | 20141127 | NA | 691,671 | 2,046,284 | 1,677 | 4,394 | 99.7 |
| 1455 | control | 20141203 | 132 | 1,492,461 | 3,809,266 | 2,782 | 3,952 | 98.6 |
| Average | | | 81 | 784,057 | 3,091,678 | 1,696 | 4,189 | 98.8 |

Default parameters of BD Aril II:

FSC=260; SSC=330; DAPI=250; PE=457

Description of the FACS gating:

The first gate (P1) was set as an FSC-A vs. SSC-A plot to discriminate the population containing small size debris. The second (P2) and third (P3) gates were set as FSC-H vs. FSC-W and SSC-H vs. SSC-D plots, respectively, to remove doublets and clumps. The fourth (P4) NeuN$^+$ and fifth (P5) NeuN$^-$ gates were set in the NeuN-PE (561 nm) vs. FSC-A plot. The sixth (P6) NeuN$^-$ and seventh (P7) NeuN$^+$ gates were set in a count plot of NeuN-stained nuclei.

Description of the FACS sorting log files:

Before staining nuclei, we kept 20 µl of the nuclei fraction from the human brain tissue sample in another tube and add 180 µl of 1% BSA in PBS. This served as an unstained control sample (TUBE: PBS CONTROL) for flow cytometry. The rest of nuclei fraction was stained with NeuN-PE. Before sorting samples, the cytometer settings should be optimized for scatter and fluorescence parameters by adjusting forward scatter (FSC) and side scatter (SSC) voltages. The FSC threshold should be optimized by using an unstained control sample (TUBE: PBS CONTROL) to display the population of nuclei appropriately on the FSC vs. SSC plot. Fluorescence photomultiplier tube (PMT) voltage for the signal from blue laser should be optimized using a small volume of the stained sample (TUBE: PE ONLY CONTROL). Adjusting P4 (NeuN$^+$) and P5 (NeuN$^-$) sorting gates using a small volume of the stained sample with DAPI (TUBE: PE + DAPI). After the adjustment, we sorted the nuclei fraction by P4 (NeuN$^+$) and P5 (NeuN$^-$) gates using NeuN-PE stained sample (TUBE: PE SAMPLE). Collect the sorted nuclei in the collection tubes. After sorting, some of the sorted fractions should be reanalyzed to verify the purity (TUBE: GLIA BACK and TUBE: NEURON BACK).

### The BD Arial II FACS sorting reports of UMB#1815

Sample: prefrontal cortex

Category: Rett patient

### FACSDiva Version 6.1.3

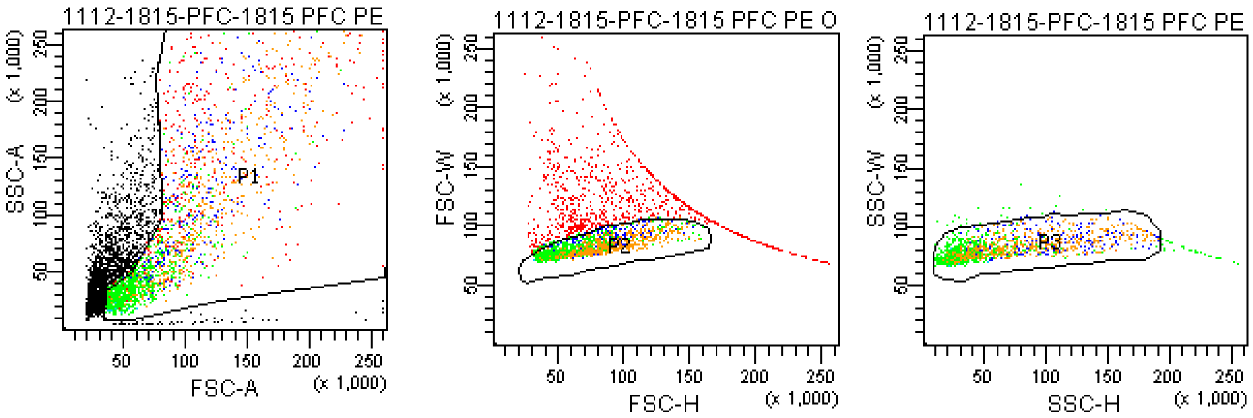


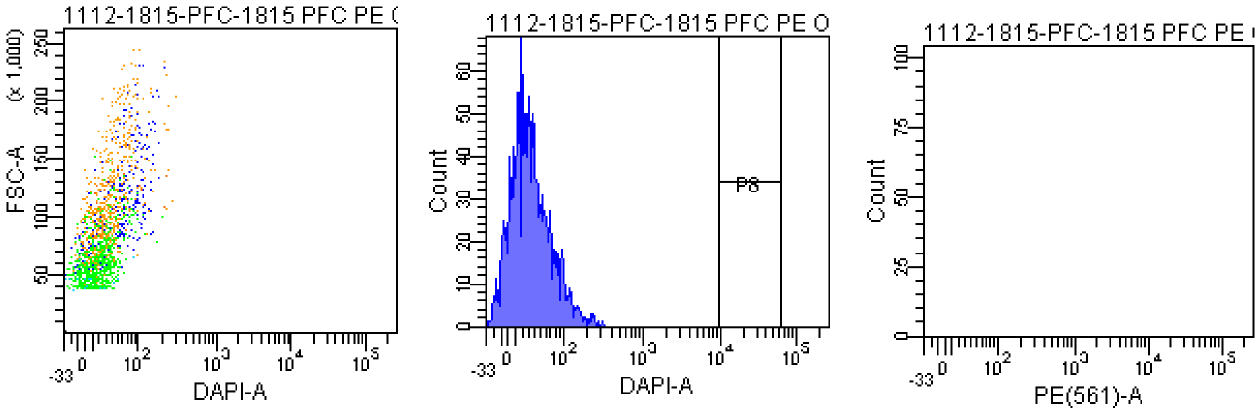


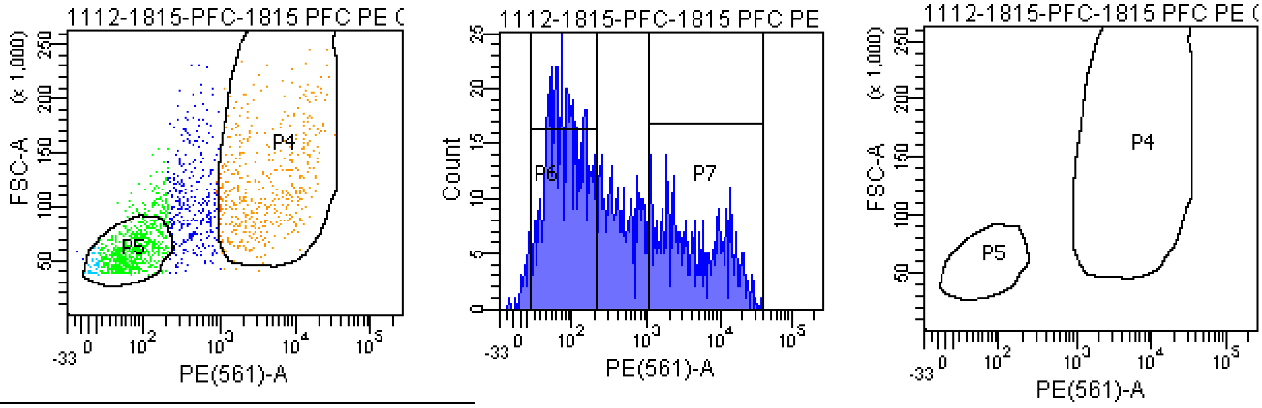


Tube: 1815 PFC PE ONLY 400V

| Population | #Events | %Parent |
|---|---|---|
| All Events | 4,566 | #### |
| P1 | 2,627 | 57.5 |
| P2 | 1,668 | 63.5 |
| P3 | 1,588 | 95.2 |
| P4 | 473 | 29.8 |
| P5 | 683 | 43.0 |
| P6 | 738 | 46.5 |
| P7 | 450 | 28.3 |
| P8 | 0 | 0.0 |

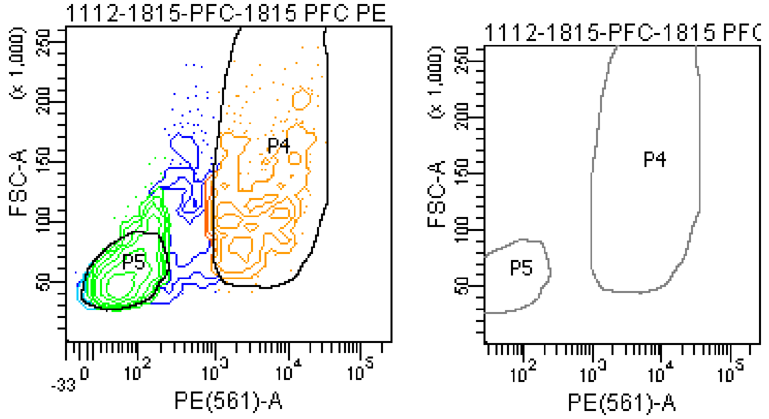

### FACSDiva Version 6.1.3

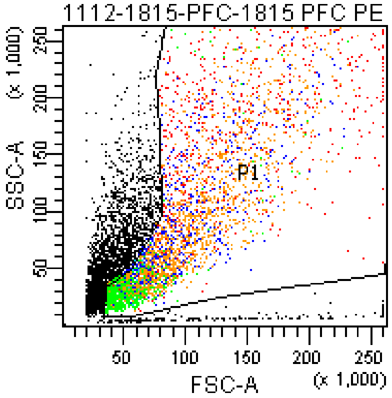


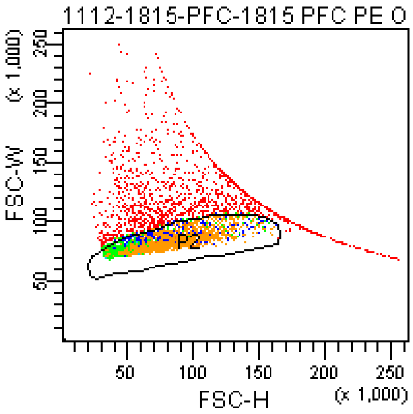


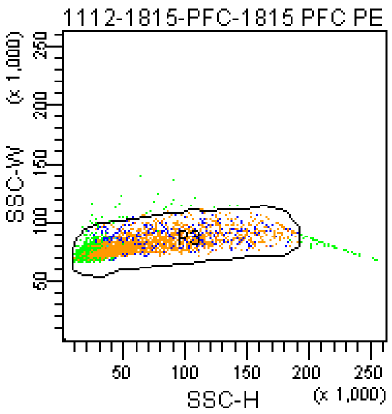


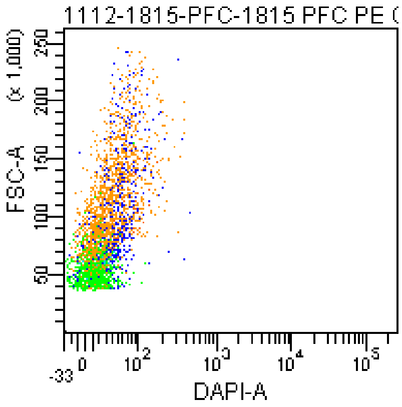


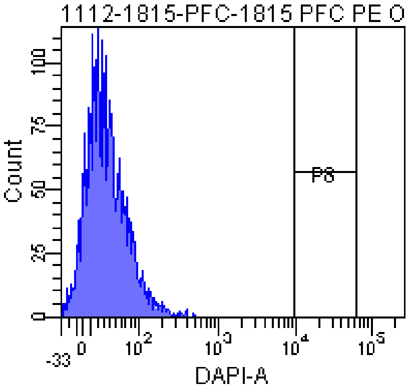


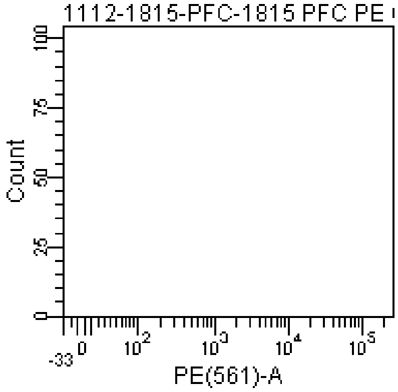


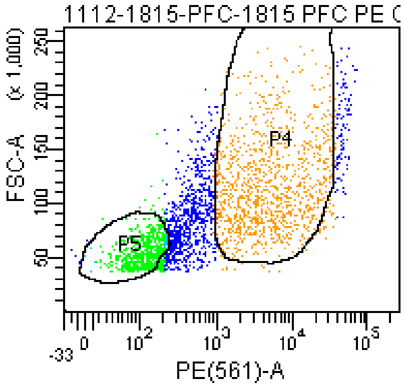


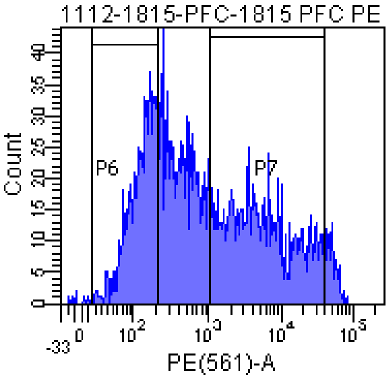


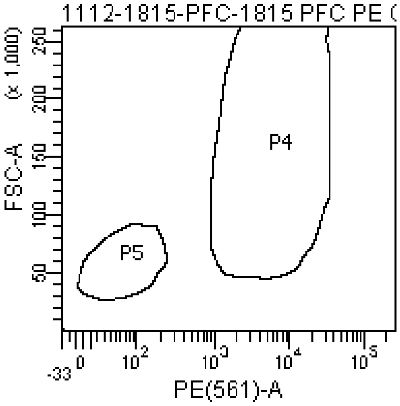


Tube: 1815 PFC PE ONLY 457V

| Population | #Events | %Parent |
|---|---|---|
| All Events | 7,474 | #### |
| P1 | 4,508 | 60.3 |
| P2 | 3,008 | 66.7 |
| P3 | 2,906 | 96.6 |
| P4 | 1,094 | 37.6 |
| P5 | 703 | 24.2 |
| P6 | 776 | 26.7 |
| P7 | 1,120 | 38.5 |
| P8 | 0 | 0.0 |

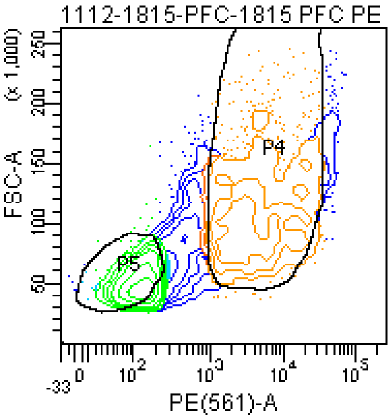


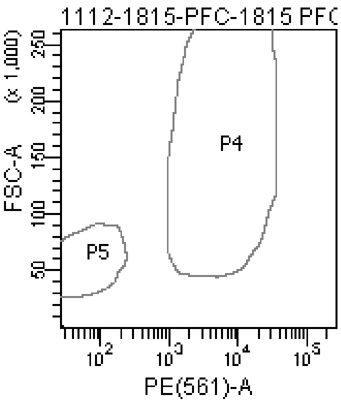

### FACSDiva Version 6.1.3

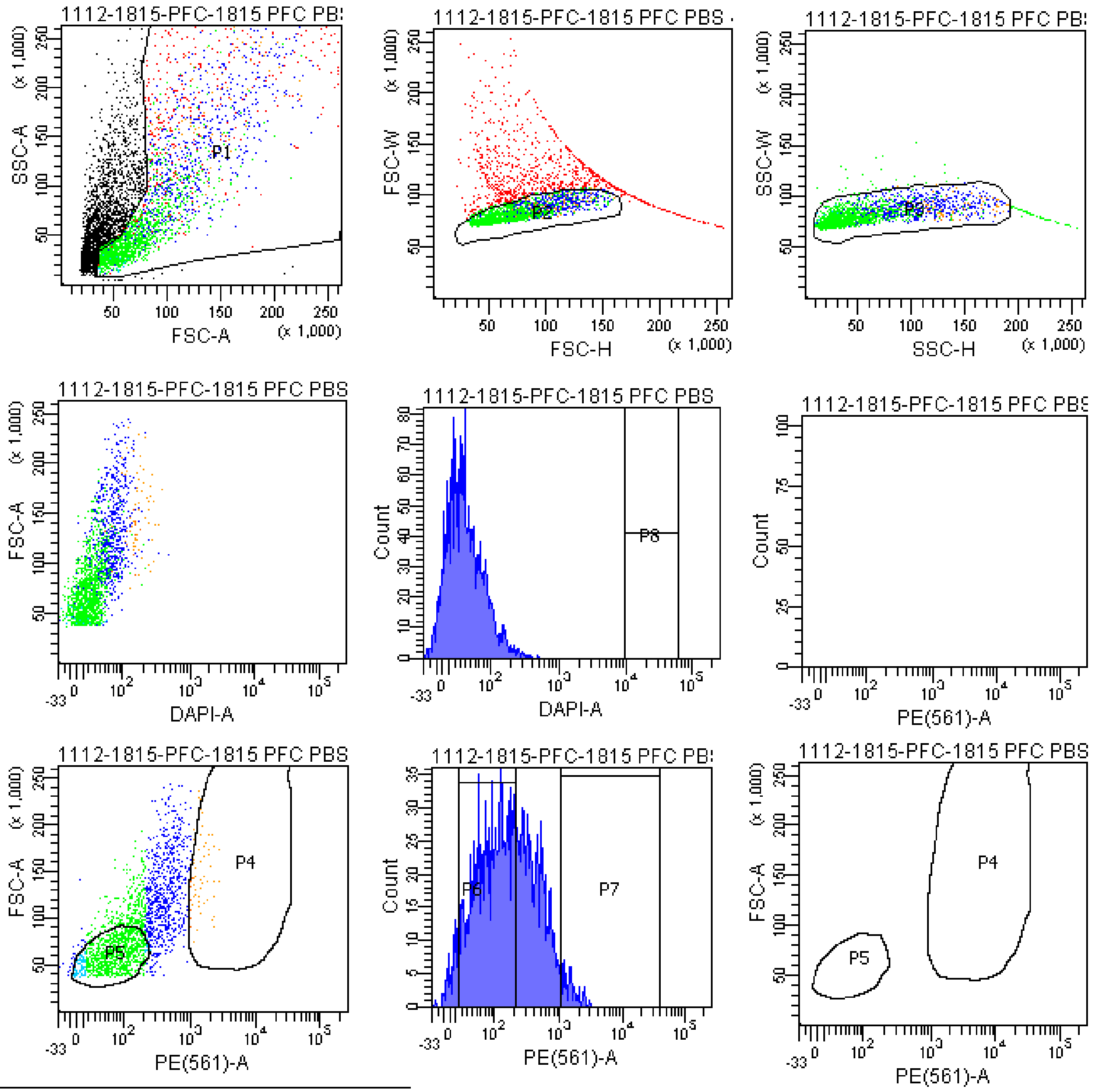


| Tube: 1815 PFC PBS 457V | | |
|---|---|---|
| Population | #Events | %Parent |
| All Events | 5,294 | #### |
| P1 | 3,116 | 58.9 |
| P2 | 2,178 | 69.9 |
| P3 | 2,071 | 95.1 |
| P4 | 65 | 3.1 |
| P5 | 989 | 47.8 |
| P6 | 1,190 | 57.5 |
| P7 | 57 | 2.8 |
| P8 | 0 | 0.0 |

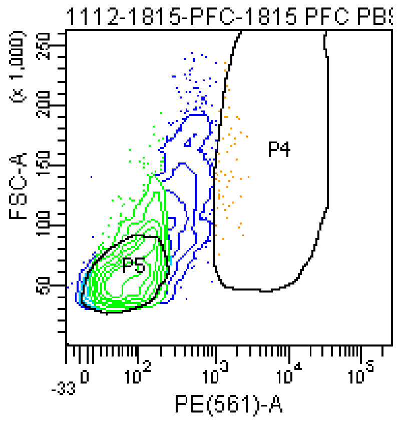


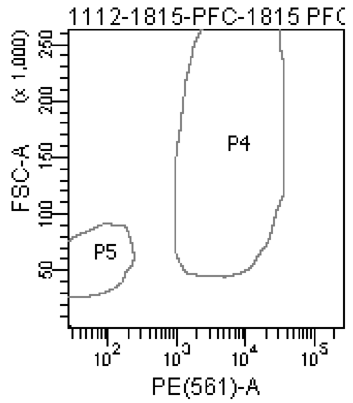

### FACSDiva Version 6.1.3

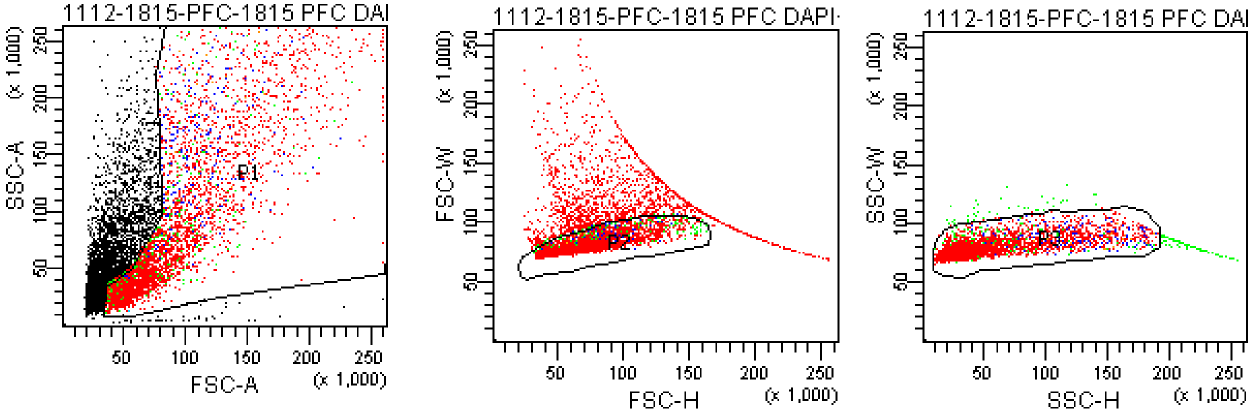


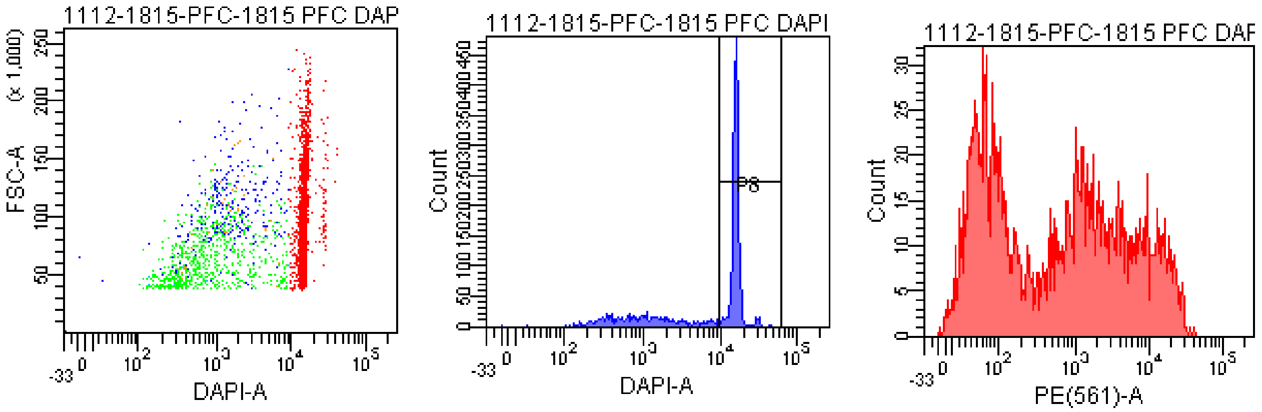


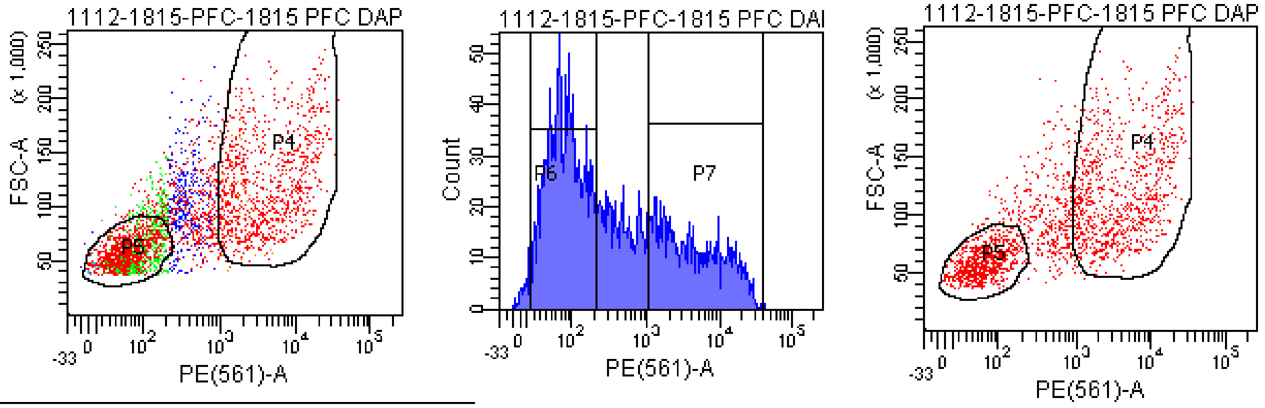


Tube: 1815 PFC DAPI+PE 400V

| Population | #Events | %Parent |
|---|---|---|
| All Events | 8,814 | #### |
| P1 | 5,049 | 57.3 |
| P2 | 3,363 | 66.6 |
| P3 | 3,159 | 93.9 |
| P4 | 979 | 31.0 |
| P5 | 1,381 | 43.7 |
| P6 | 1,489 | 47.1 |
| P7 | 941 | 29.8 |
| P8 | 2,121 | 67.1 |

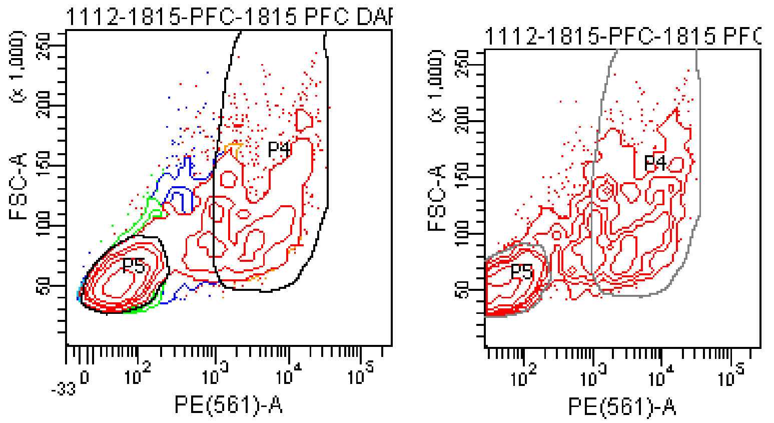

### FACSDiva Version 6.1.3

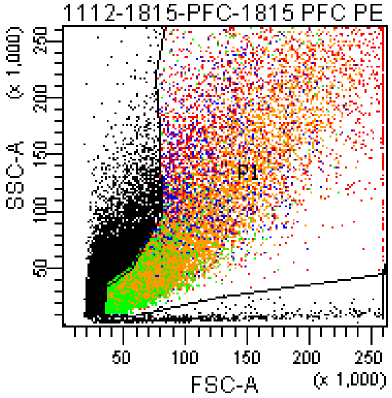


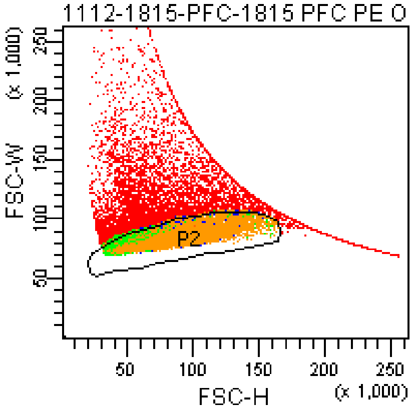


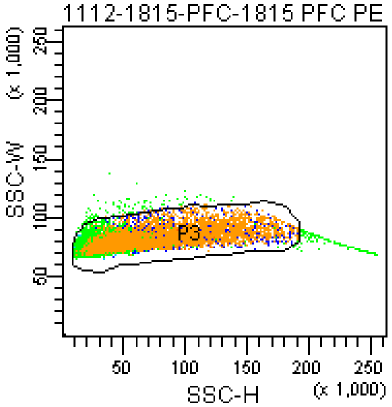


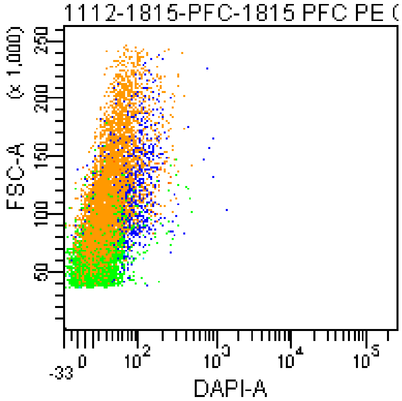


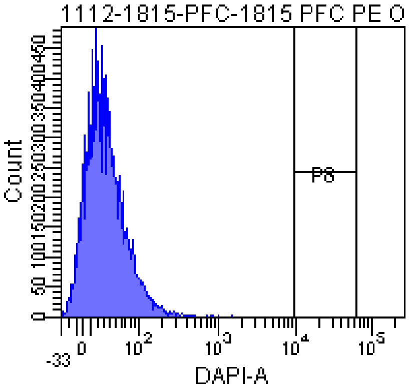


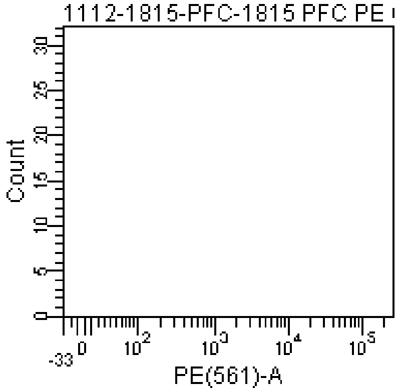


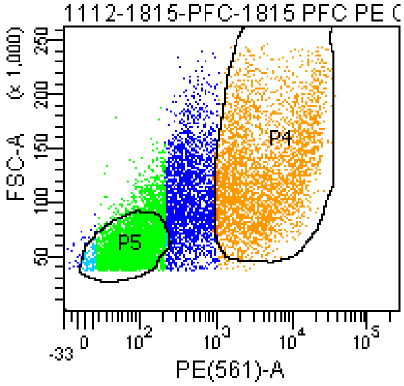


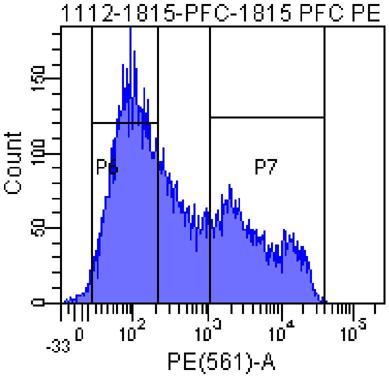


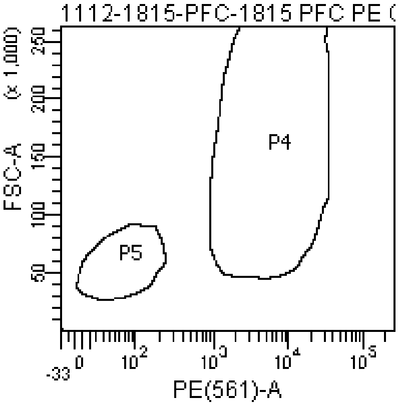


Tube: 1815 PFC PE ONLY SAMPLE

| Population | #Events | %Parent |
|---|---|---|
| All Events | 31,883 | #### |
| P1 | 20,000 | 62.7 |
| P2 | 12,215 | 61.1 |
| P3 | 11,896 | 97.4 |
| P4 | 3,670 | 30.9 |
| P5 | 5,067 | 42.6 |
| P6 | 5,500 | 46.2 |
| P7 | 3,578 | 30.1 |
| P8 | 0 | 0.0 |

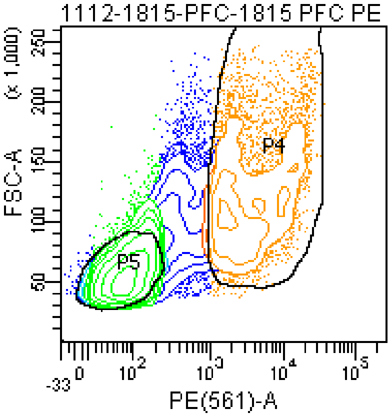


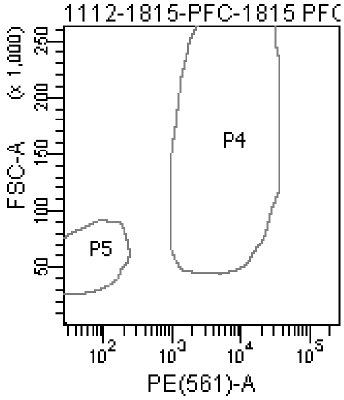

### FACSDiva Version 6.1.3

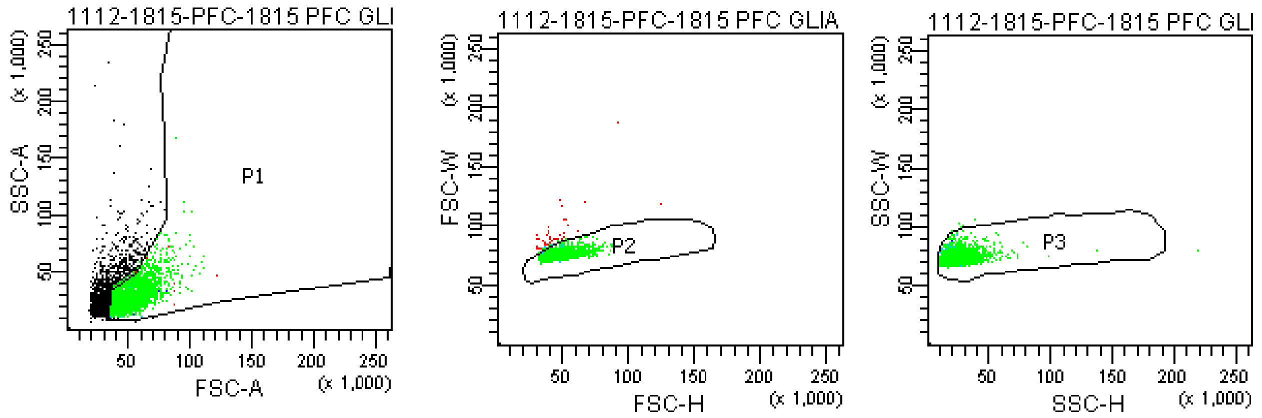


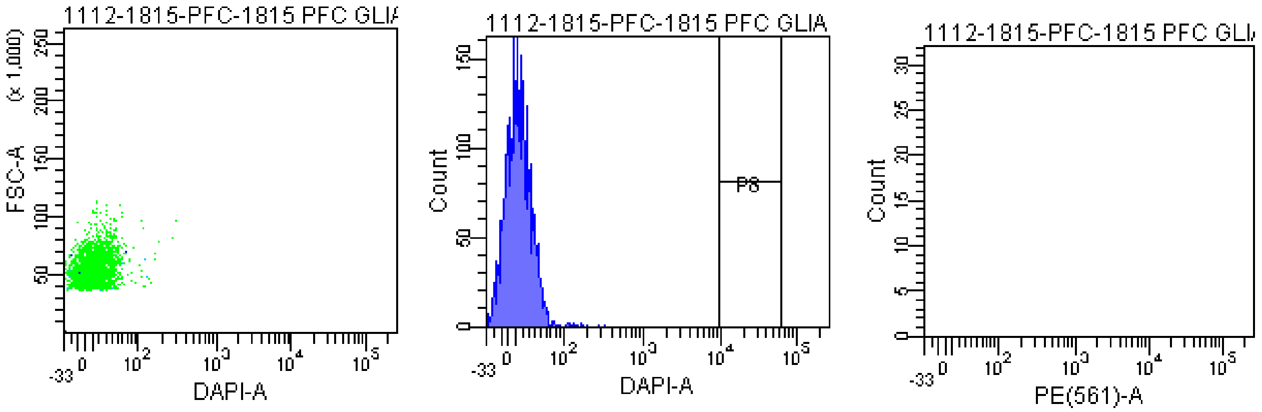


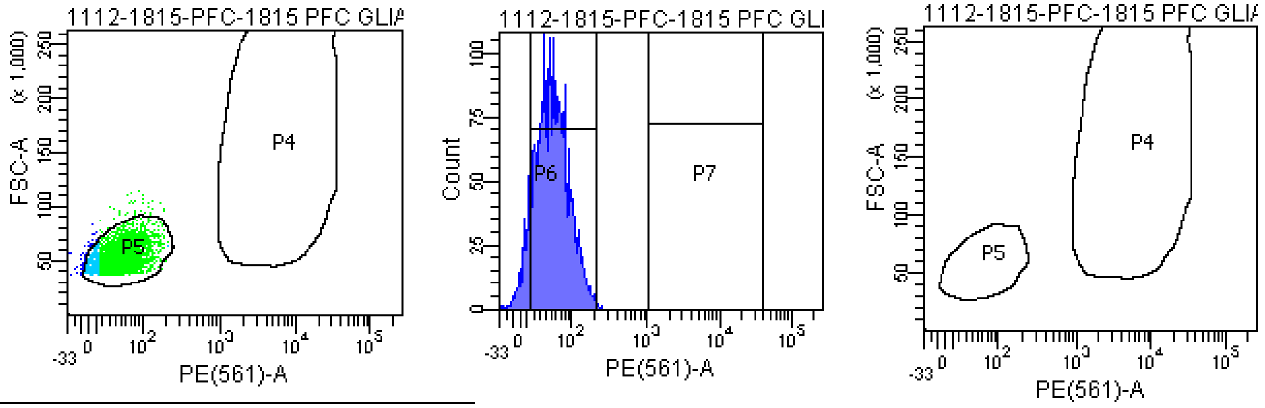


Tube: 1815 PFC GLIA BACK

| Population | #Events | %Parent |
|---|---|---|
| All Events | 4,338 | #### |
| P1 | 2,956 | 68.1 |
| P2 | 2,913 | 98.5 |
| P3 | 2,905 | 99.7 |
| P4 | 0 | 0.0 |
| P5 | 2,816 | 96.9 |
| P6 | 2,603 | 89.6 |
| P7 | 0 | 0.0 |
| P8 | 0 | 0.0 |

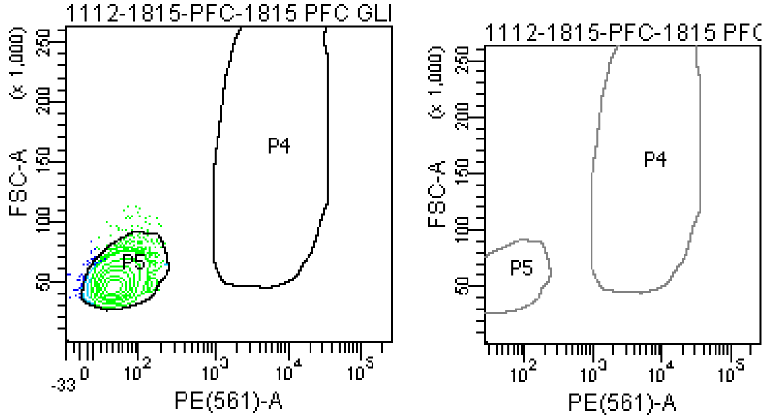

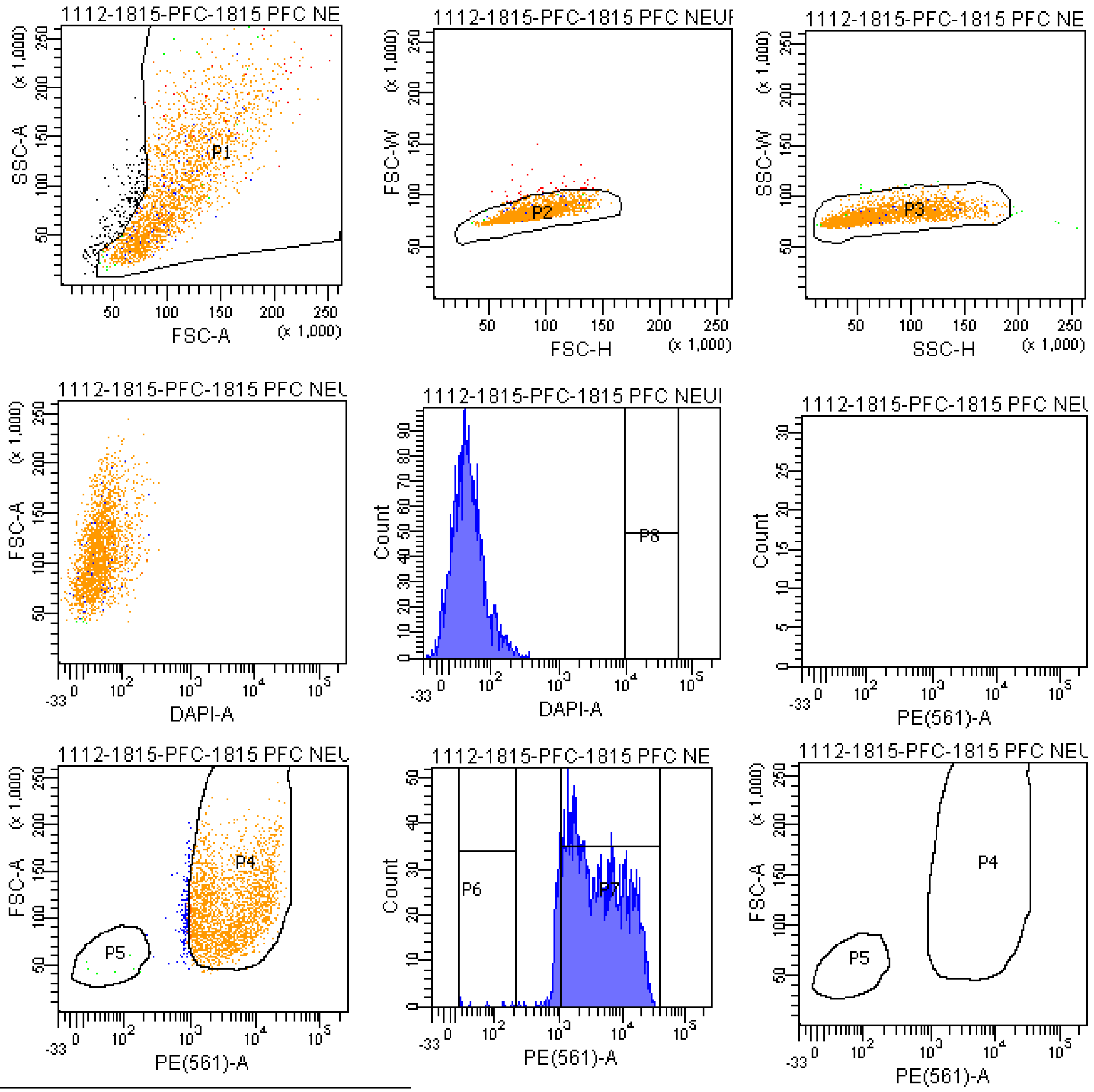


Tube: 1815 PFC NEURON BACK

| Population | #Events | %Parent |
|---|---|---|
| All Events | 2,716 | #### |
| P1 | 2,519 | 92.7 |
| P2 | 2,476 | 98.3 |
| P3 | 2,461 | 99.4 |
| P4 | 2,286 | 92.9 |
| P5 | 6 | 0.2 |
| P6 | 7 | 0.3 |
| P7 | 2,218 | 90.1 |
| P8 | 0 | 0.0 |

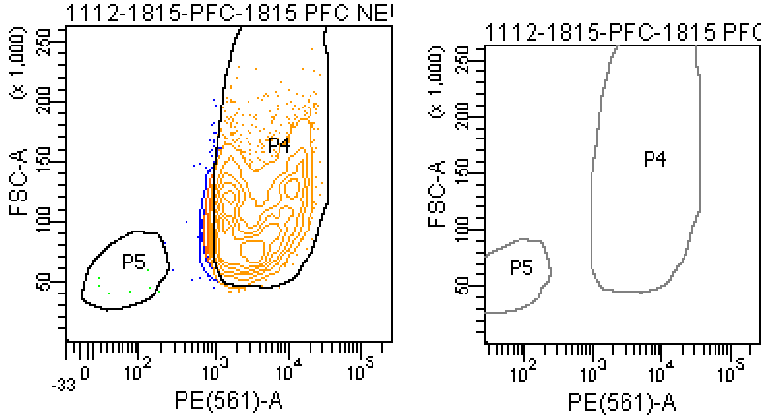

**The BD Arial II FACS sorting reports of UMB#1571**

Sample: prefrontal cortex

Category: healthy control

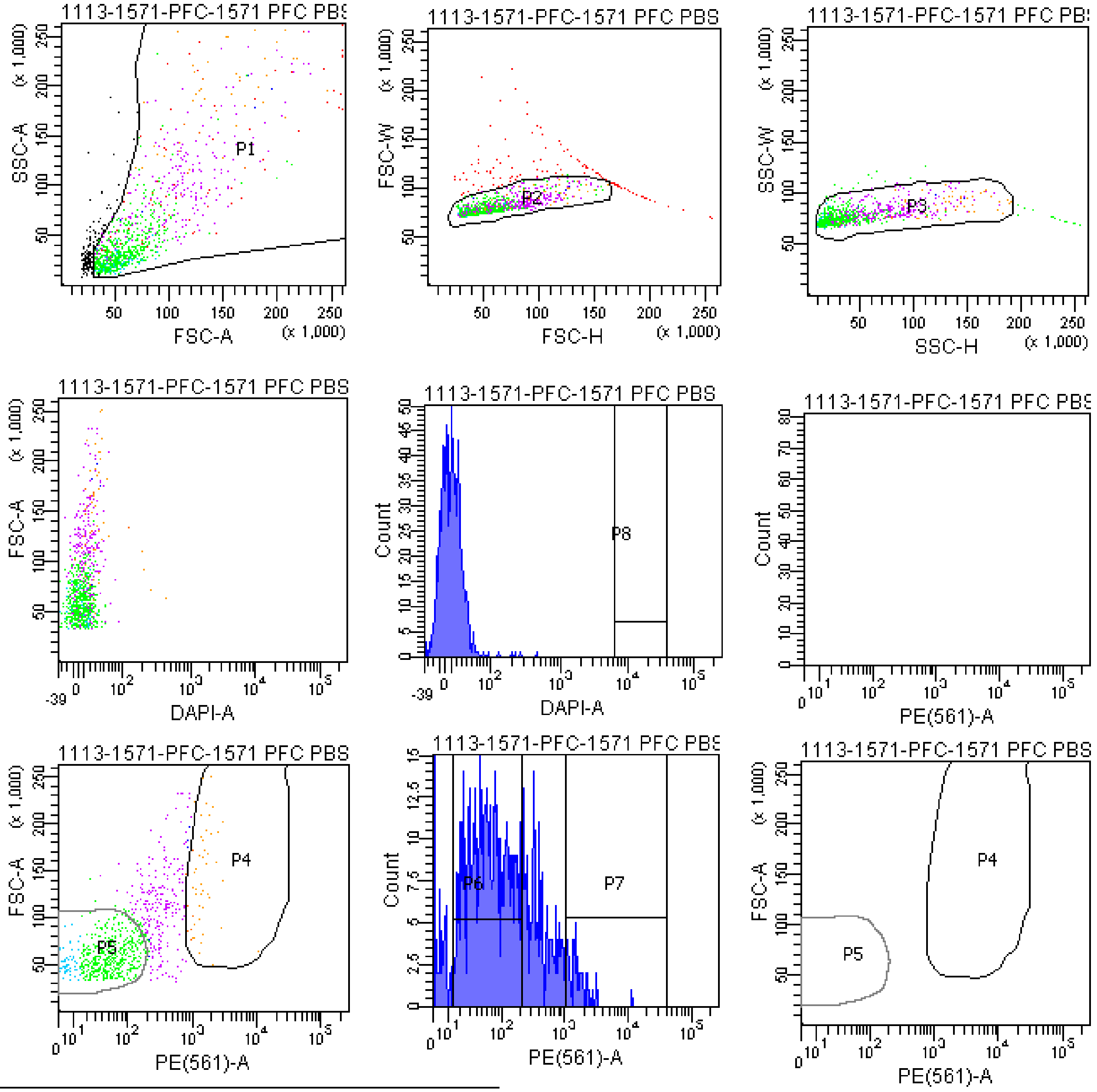


Tube: 1571 PFC PBS 500V

| Population | #Events | %Parent | %Total |
|---|---|---|---|
| All Events | 1,195 | #### | 100.0 |
| P1 | 1,022 | 85.5 | 85.5 |
| P2 | 901 | 88.2 | 75.4 |
| P3 | 867 | 96.2 | 72.6 |
| P4 | 54 | 6.2 | 4.5 |
| P5 | 568 | 65.5 | 47.5 |
| P6 | 532 | 61.4 | 44.5 |
| P7 | 44 | 5.1 | 3.7 |
| P8 | 0 | 0.0 | 0.0 |
| P9 | 215 | 24.8 | 18.0 |

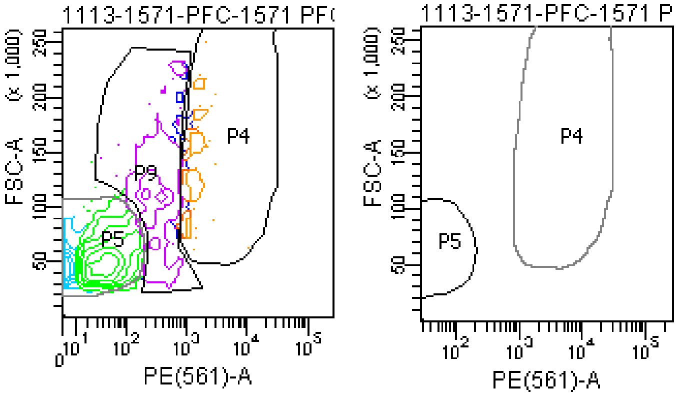

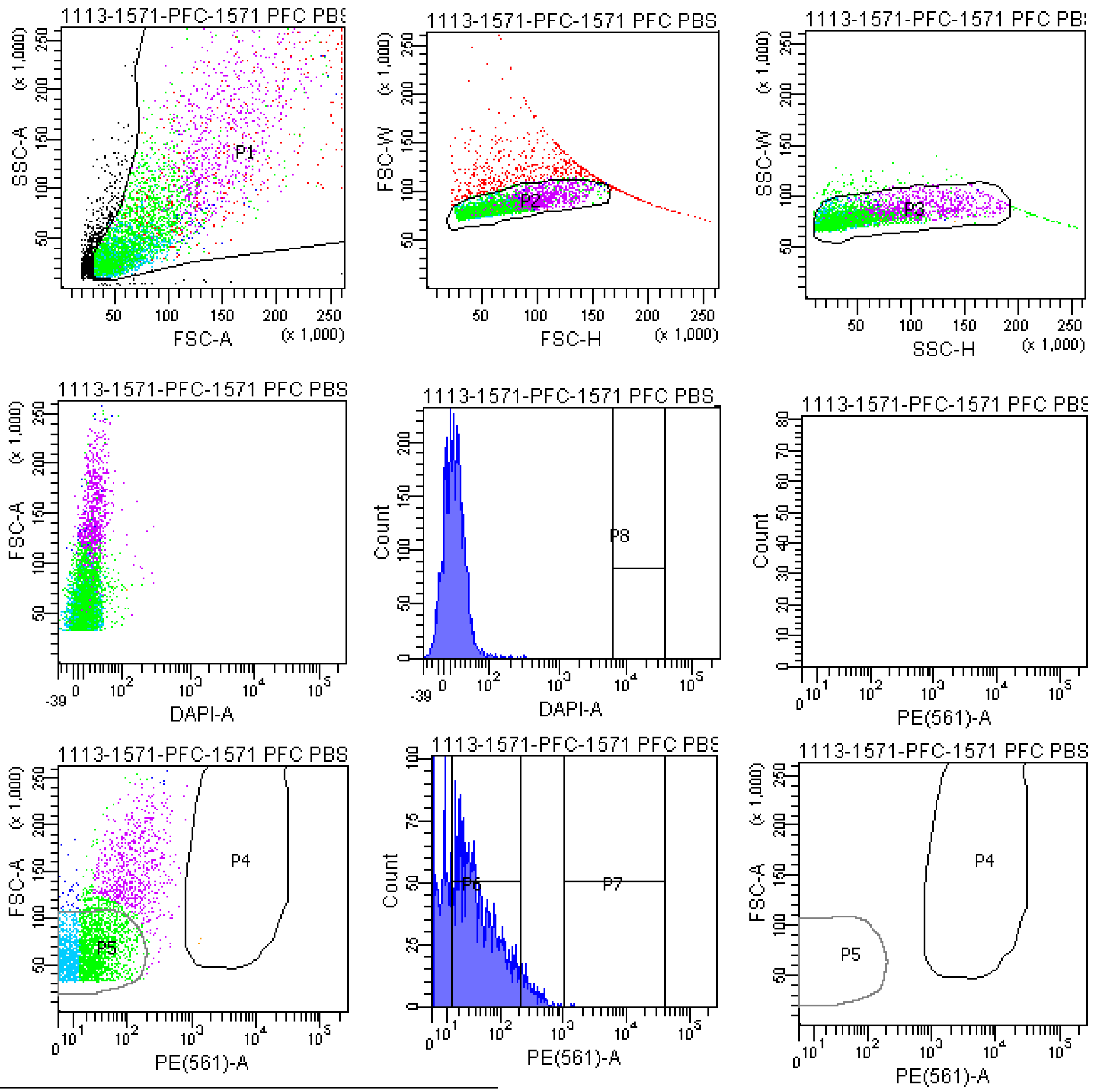


Tube: 1571 PFC PBS_400V

| Population | #Events | %Parent | %Total |
|---|---|---|---|
| All Events | 6,083 | #### | 100.0 |
| P1 | 5,232 | 86.0 | 86.0 |
| P2 | 4,495 | 85.9 | 73.9 |
| P3 | 4,345 | 96.7 | 71.4 |
| P4 | 2 | 0.0 | 0.0 |
| P5 | 3,413 | 78.6 | 56.1 |
| P6 | 2,350 | 54.1 | 38.6 |
| P7 | 2 | 0.0 | 0.0 |
| P8 | 0 | 0.0 | 0.0 |
| P9 | 706 | 16.2 | 11.6 |

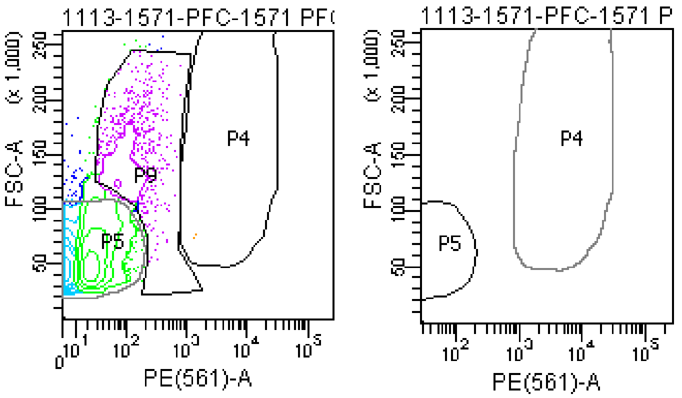

#### FACSDiva Version 6.1.3

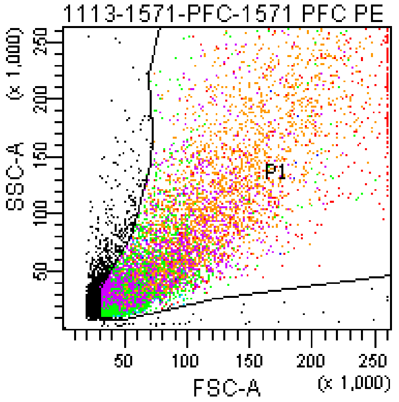


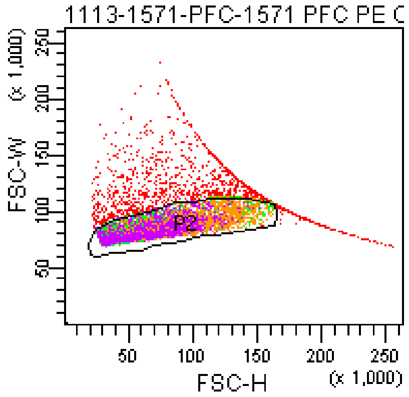


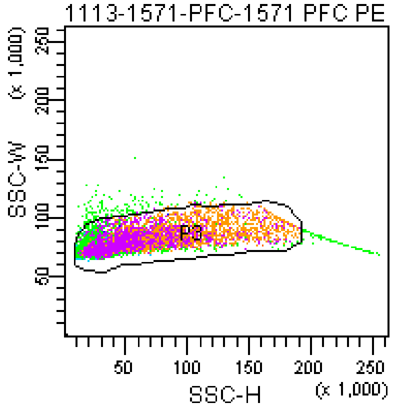


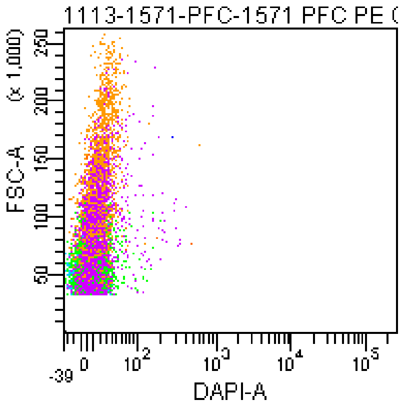


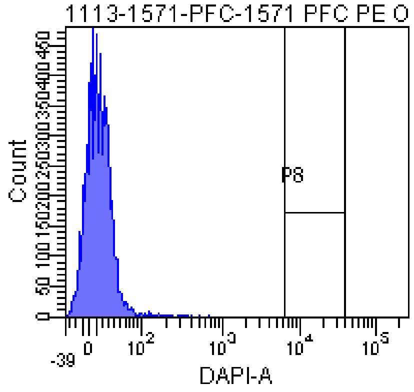


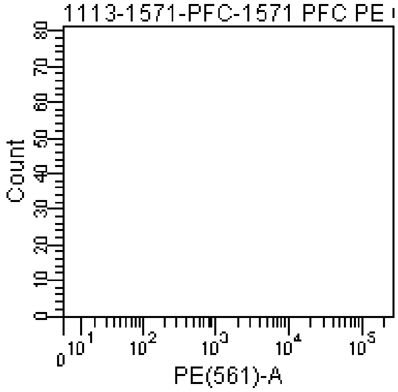


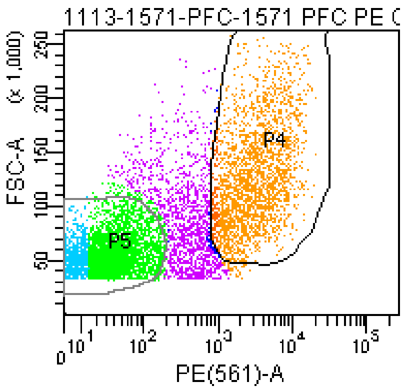


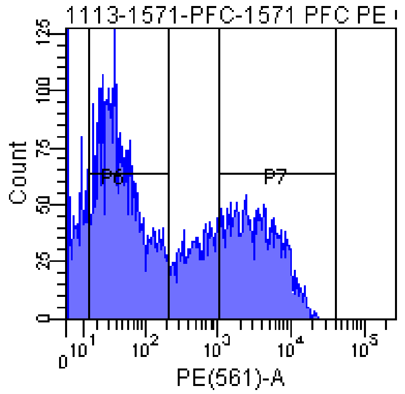


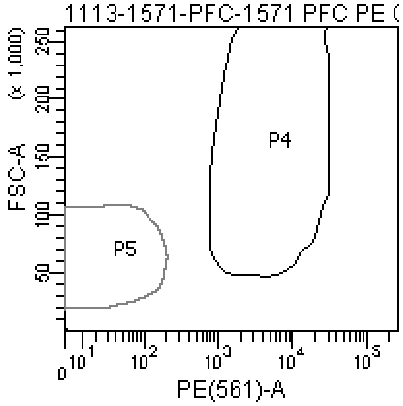


Tube: 1571 PFC PE ONLY

| Population | #Events | %Parent | %Total |
|---|---|---|---|
| All Events | 11,886 | #### | 100.0 |
| P1 | 10,301 | 86.7 | 86.7 |
| P2 | 8,790 | 85.3 | 74.0 |
| P3 | 8,473 | 96.4 | 71.3 |
| P4 | 2,462 | 29.1 | 20.7 |
| P5 | 4,548 | 53.7 | 38.3 |
| P6 | 3,550 | 41.9 | 29.9 |
| P7 | 2,372 | 28.0 | 20.0 |
| P8 | 0 | 0.0 | 0.0 |
| P9 | 1,235 | 14.6 | 10.4 |

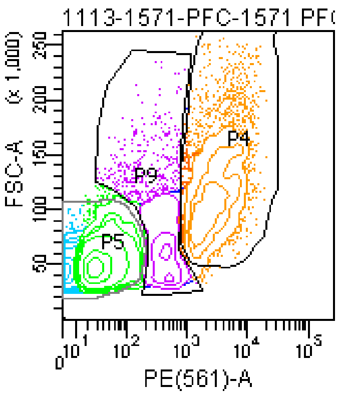


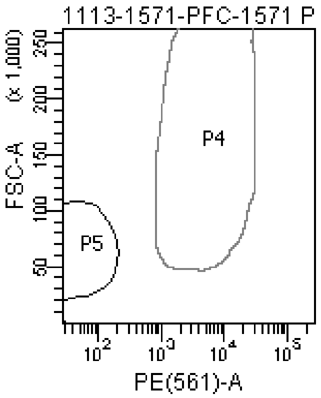

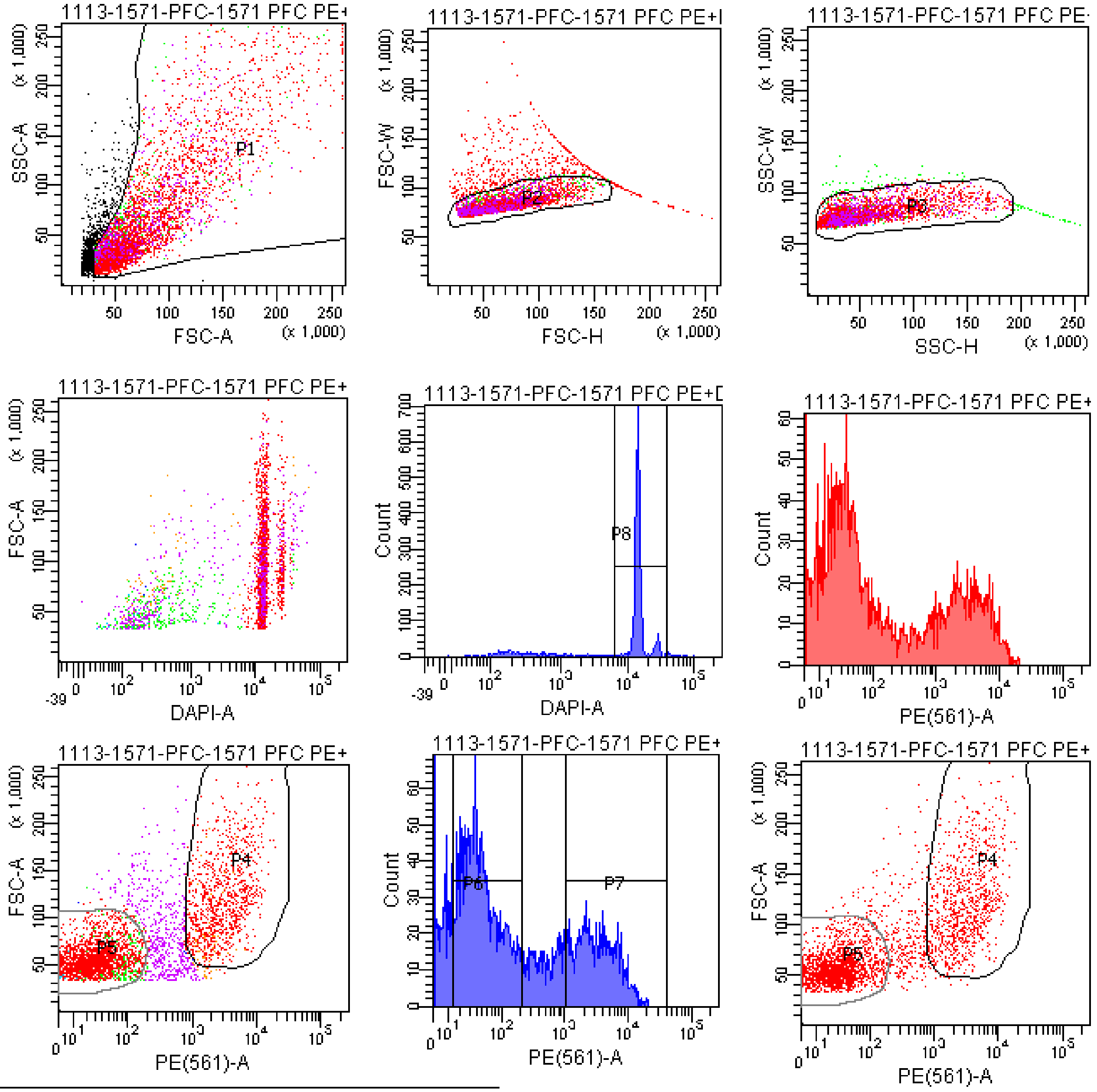


Tube: 1571 PFC PE+DAPI

| Population | #Events | %Parent | %Total |
|---|---|---|---|
| All Events | 5,509 | #### | 100.0 |
| P1 | 4,699 | 85.3 | 85.3 |
| P2 | 4,116 | 87.6 | 74.7 |
| P3 | 4,002 | 97.2 | 72.6 |
| P4 | 993 | 24.8 | 18.0 |
| P5 | 2,388 | 59.7 | 43.3 |
| P6 | 1,844 | 46.1 | 33.5 |
| P7 | 970 | 24.2 | 17.6 |
| P8 | 3,366 | 84.1 | 61.1 |
| P9 | 529 | 13.2 | 9.6 |

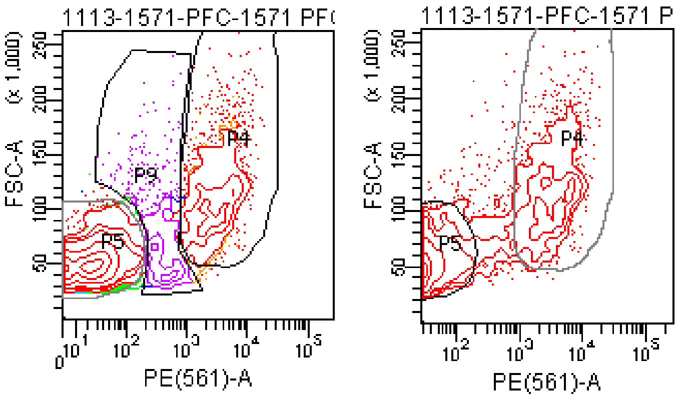

### FACSDiva Version 6.1.3

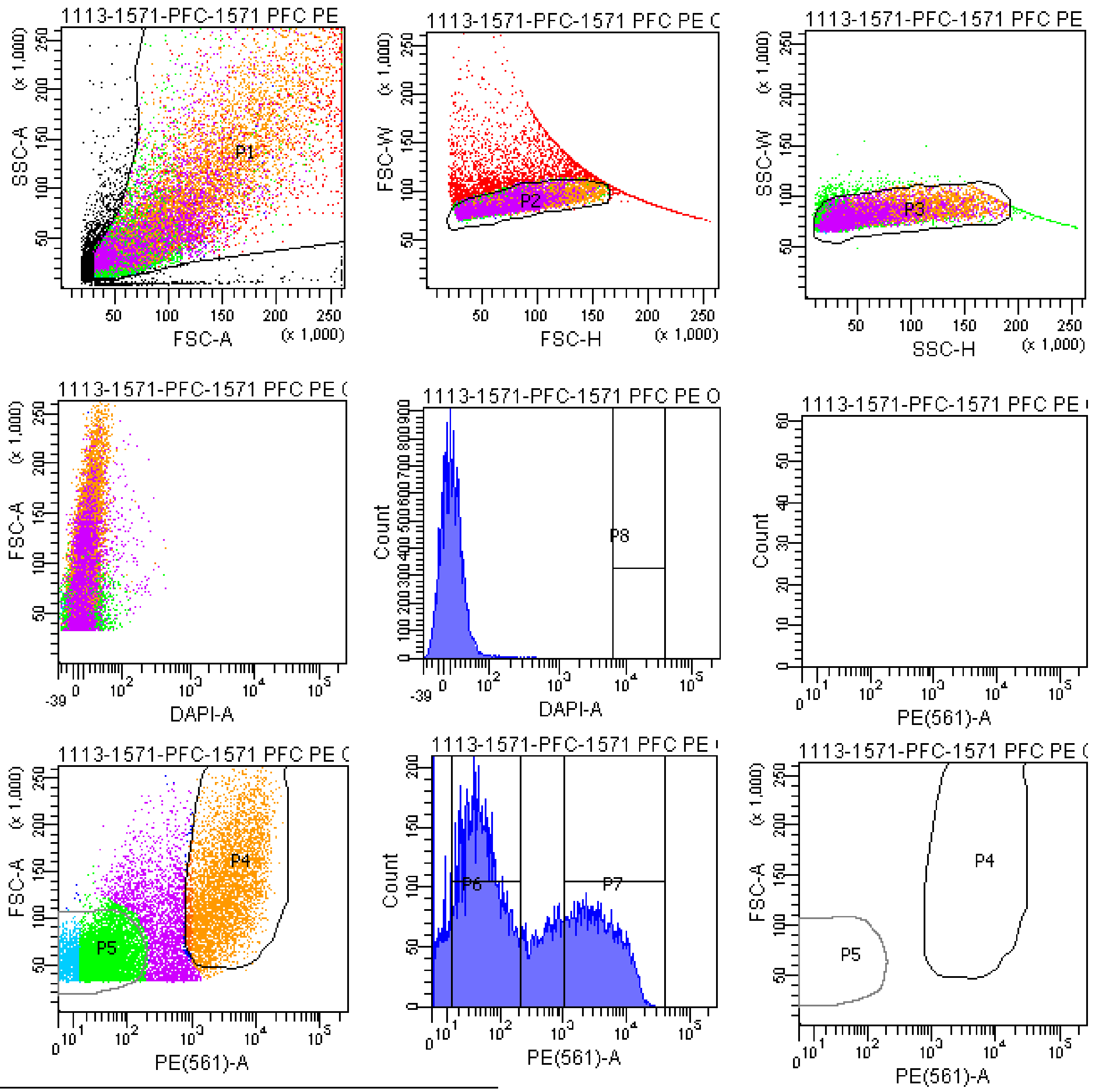


Tube: 1571 PFC PE ONLY SAMPLE

| Population | #Events | %Parent | %Total |
|---|---|---|---|
| All Events | 23,315 | #### | 100.0 |
| P1 | 20,000 | 85.8 | 85.8 |
| P2 | 16,430 | 82.2 | 70.5 |
| P3 | 15,992 | 97.3 | 68.6 |
| P4 | 4,610 | 28.8 | 19.8 |
| P5 | 8,262 | 51.7 | 35.4 |
| P6 | 7,234 | 45.2 | 31.0 |
| P7 | 4,424 | 27.7 | 19.0 |
| P8 | 0 | 0.0 | 0.0 |
| P9 | 2,632 | 16.5 | 11.3 |

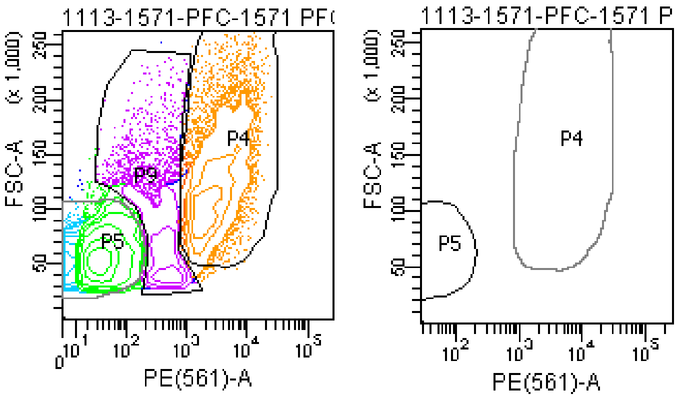

### FACSDiva Version 6.1.3

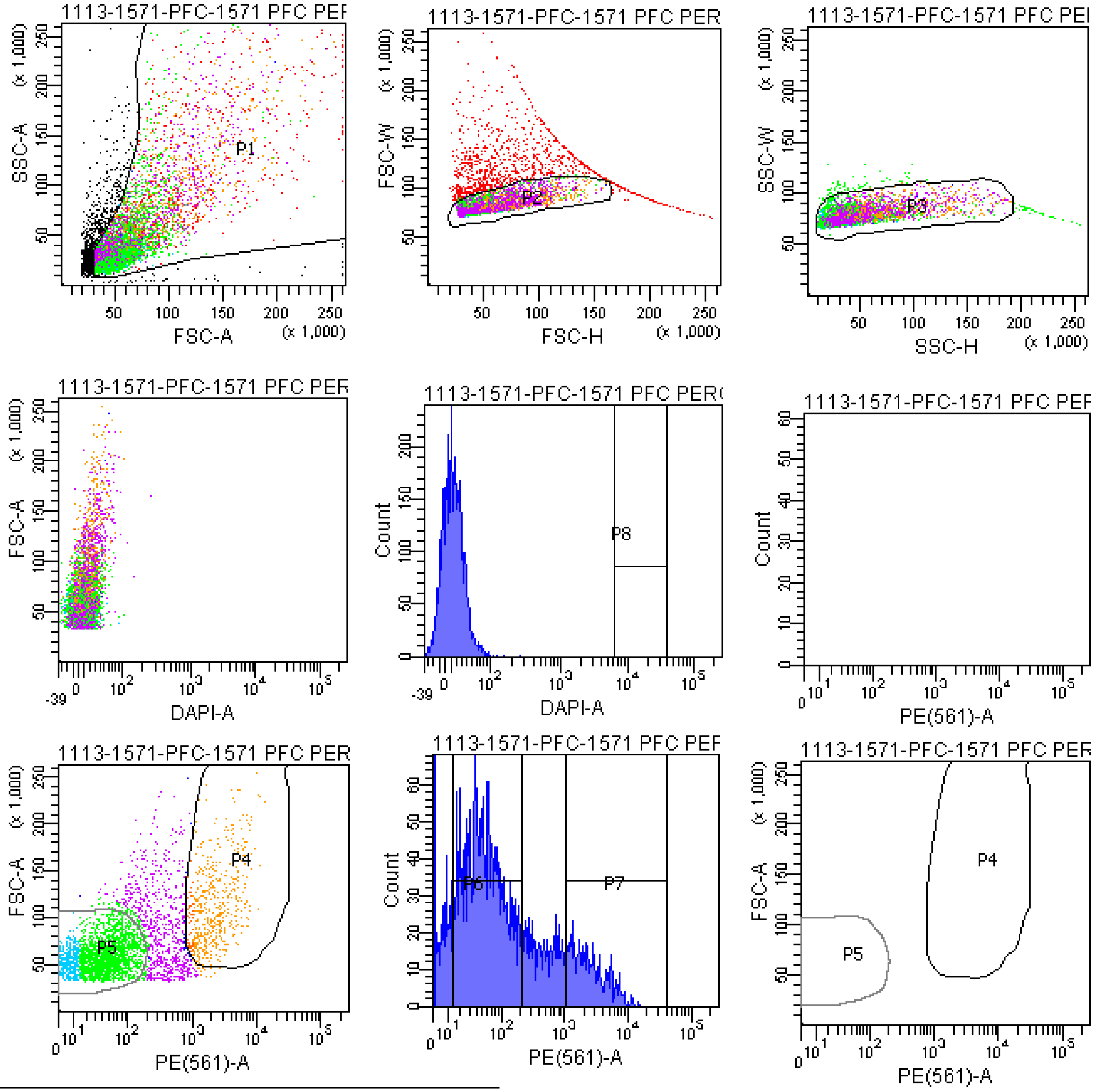


Tube: 1571 PFC PERCOLL 2 PE

| Population | #Events | %Parent | %Total |
|---|---|---|---|
| All Events | 6,172 | #### | 100.0 |
| P1 | 5,176 | 83.9 | 83.9 |
| P2 | 4,112 | 79.4 | 66.6 |
| P3 | 3,989 | 97.0 | 64.6 |
| P4 | 558 | 14.0 | 9.0 |
| P5 | 2,607 | 65.4 | 42.2 |
| P6 | 2,212 | 55.5 | 35.8 |
| P7 | 526 | 13.2 | 8.5 |
| P8 | 0 | 0.0 | 0.0 |
| P9 | 690 | 17.3 | 11.2 |

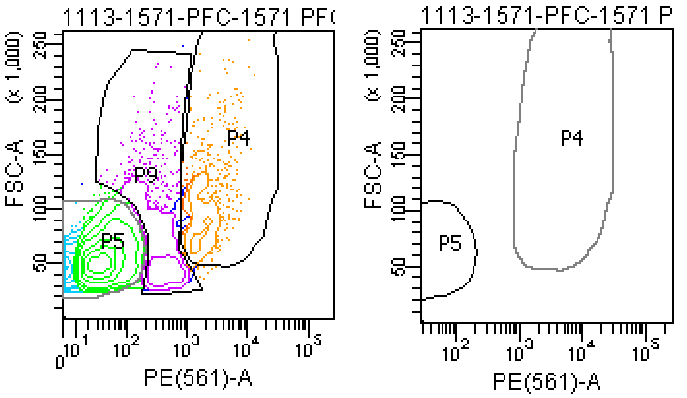

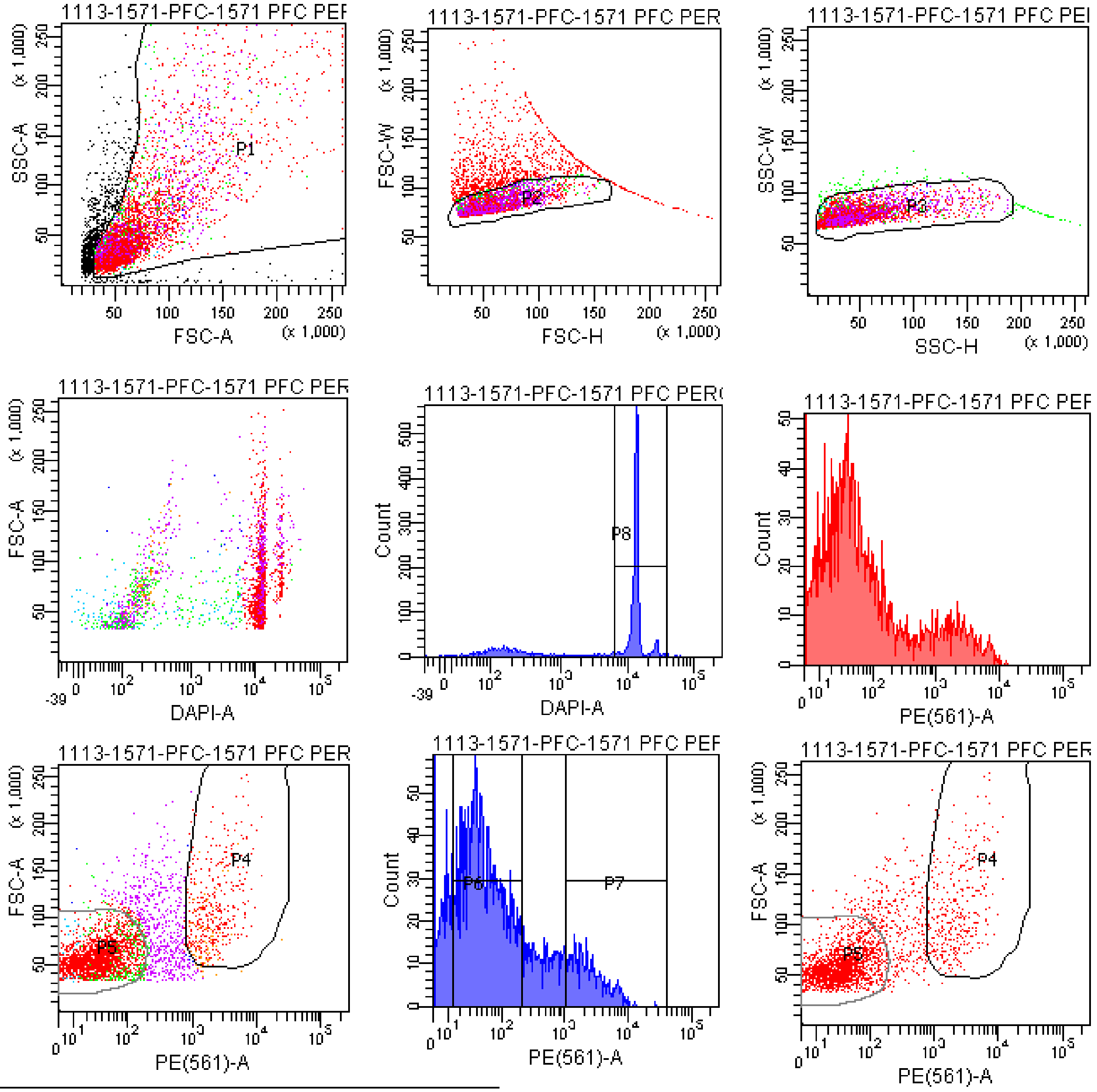


| Tube: | 1571 PFC PERCOLL 2 DAPI+PE | | |
|---|---|---|---|
| Population | #Events | %Parent | %Total |
| All Events | 5,493 | #### | 100.0 |
| P1 | 4,498 | 81.9 | 81.9 |
| P2 | 3,526 | 78.4 | 64.2 |
| P3 | 3,411 | 96.7 | 62.1 |
| P4 | 413 | 12.1 | 7.5 |
| P5 | 2,343 | 68.7 | 42.7 |
| P6 | 1,910 | 56.0 | 34.8 |
| P7 | 386 | 11.3 | 7.0 |
| P8 | 2,562 | 75.1 | 46.6 |
| P9 | 526 | 15.4 | 9.6 |

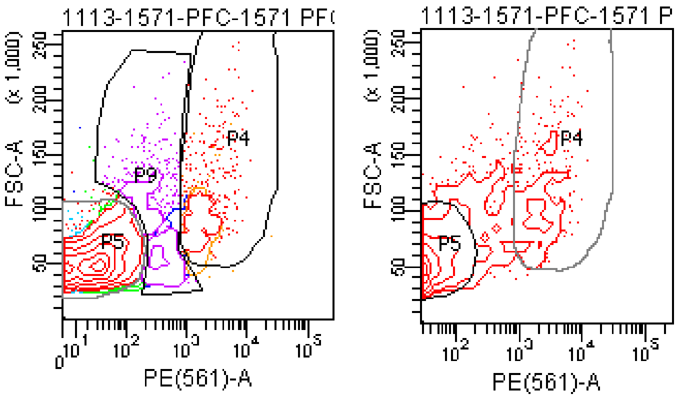

### FACSDiva Version 6.1.3

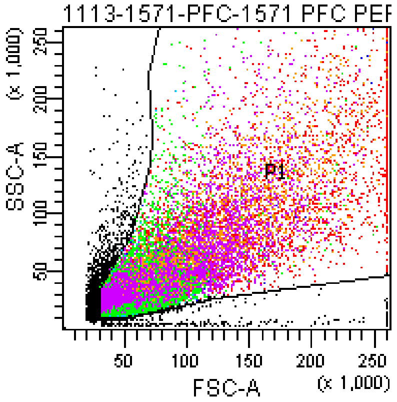


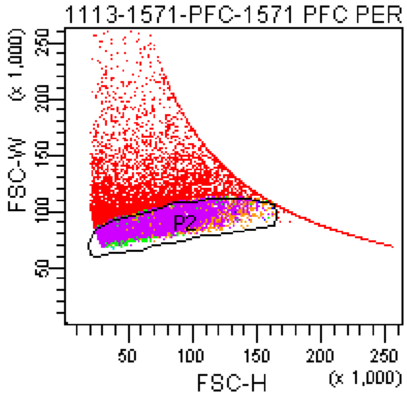


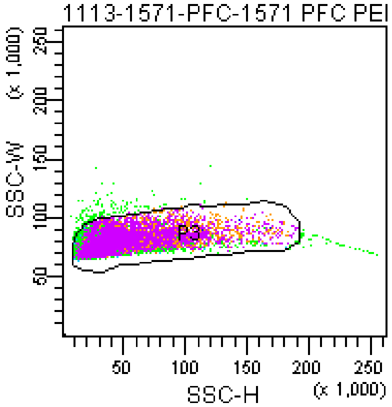


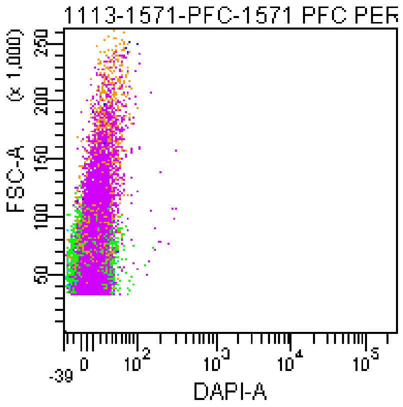


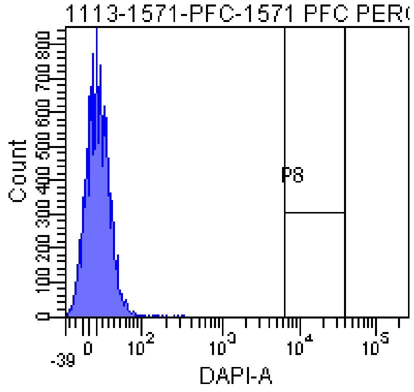


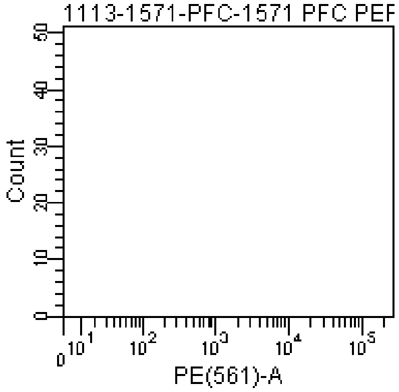


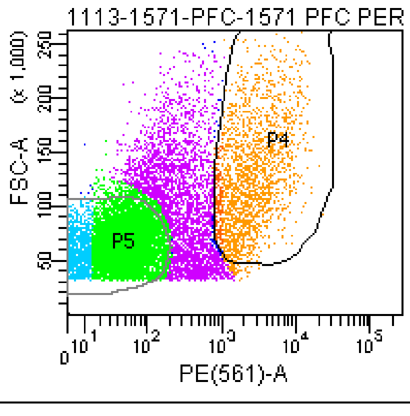


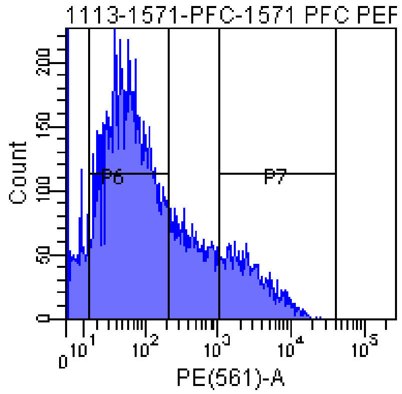


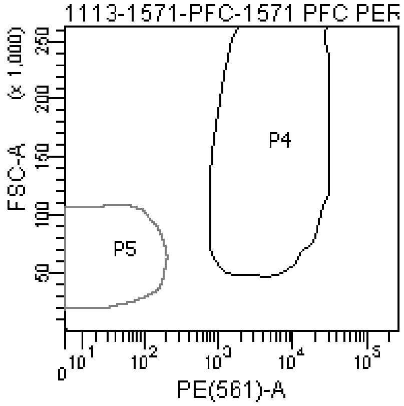


Tube: 1571 PFC PERCOLL 2 SAMPLE

| Population | #Events | %Parent | %Total |
|---|---|---|---|
| All Events | 22,960 | #### | 100.0 |
| P1 | 20,000 | 87.1 | 87.1 |
| P2 | 14,489 | 72.4 | 63.1 |
| P3 | 14,265 | 98.5 | 62.1 |
| P4 | 2,051 | 14.4 | 8.9 |
| P5 | 8,499 | 59.6 | 37.0 |
| P6 | 8,291 | 58.1 | 36.1 |
| P7 | 1,975 | 13.8 | 8.6 |
| P8 | 0 | 0.0 | 0.0 |
| P9 | 3,146 | 22.1 | 13.7 |

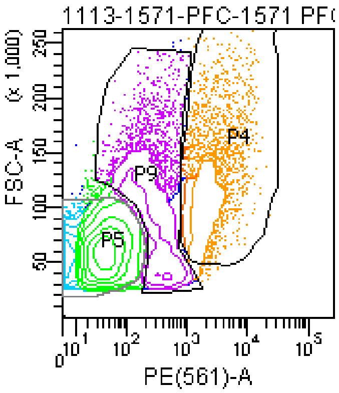


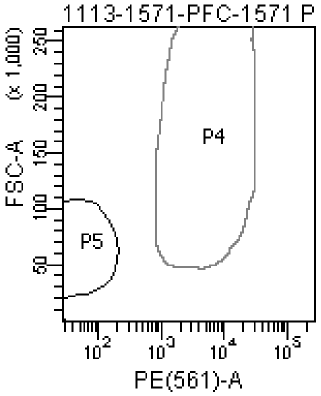

### FACSDiva Version 6.1.3

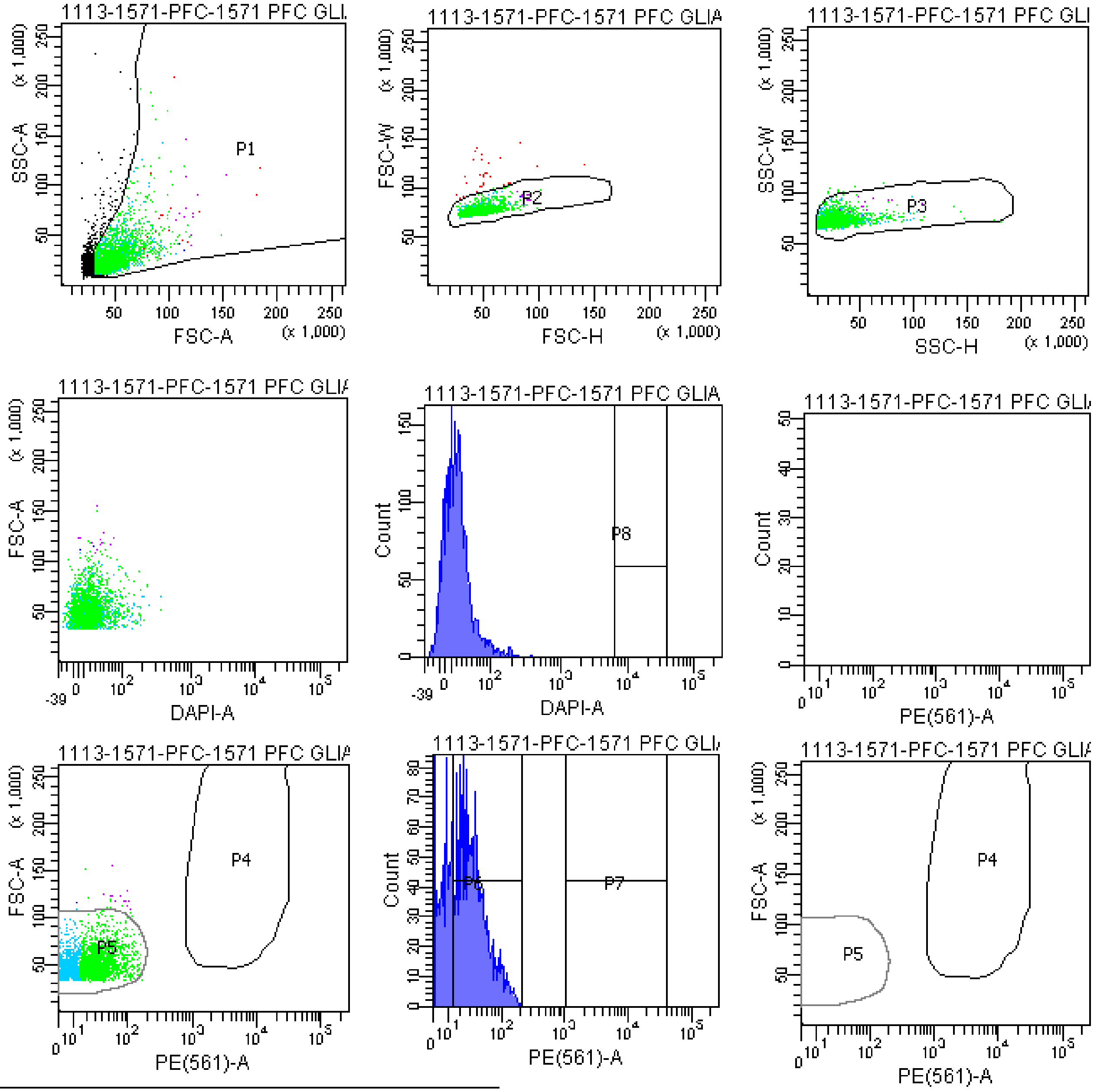


Tube: 1571 PFC GLIA BACK

| Population | #Events | %Parent | %Total |
|---|---|---|---|
| All Events | 3,688 | #### | 100.0 |
| P1 | 3,142 | 85.2 | 85.2 |
| P2 | 3,113 | 99.1 | 84.4 |
| P3 | 3,099 | 99.6 | 84.0 |
| P4 | 0 | 0.0 | 0.0 |
| P5 | 3,070 | 99.1 | 83.2 |
| P6 | 1,697 | 54.8 | 46.0 |
| P7 | 0 | 0.0 | 0.0 |
| P8 | 0 | 0.0 | 0.0 |
| P9 | 12 | 0.4 | 0.3 |

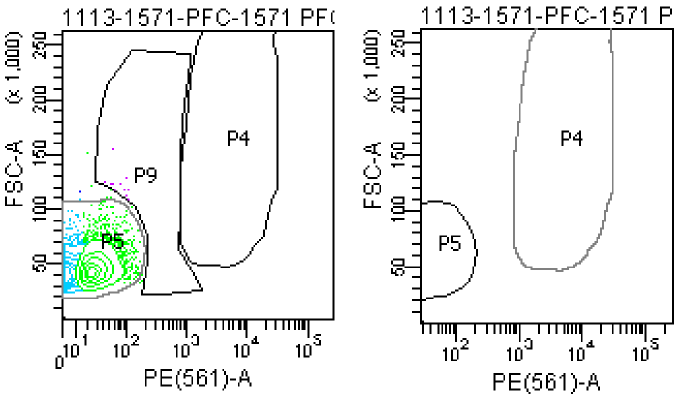

### FACSDiva Version 6.1.3

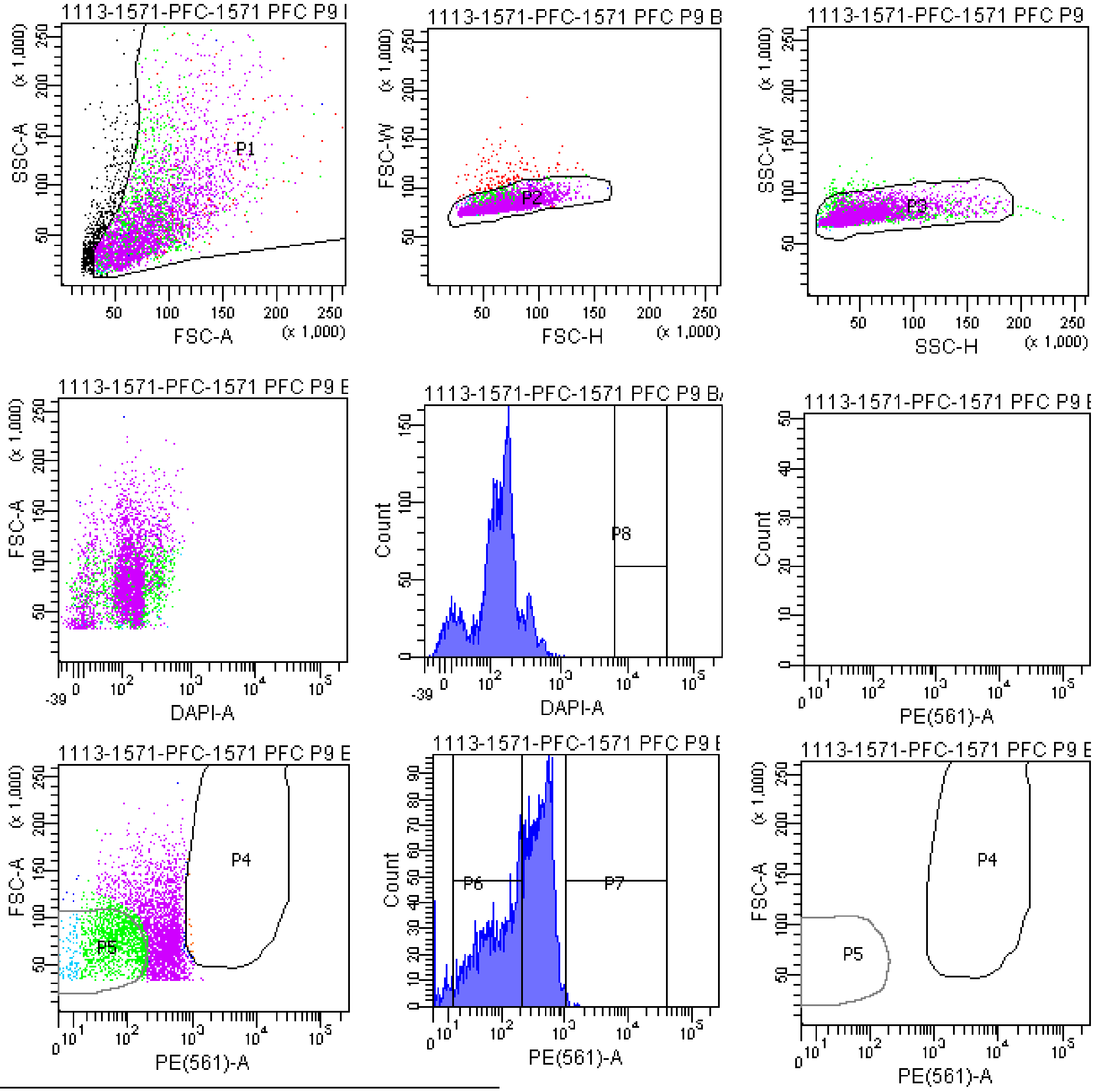


Tube: 1571 PFC P9 BACK

| Population | #Events | %Parent | %Total |
|---|---|---|---|
| All Events | 4,921 | #### | 100.0 |
| P1 | 4,306 | 87.5 | 87.5 |
| P2 | 4,161 | 96.6 | 84.6 |
| P3 | 4,090 | 98.3 | 83.1 |
| P4 | 13 | 0.3 | 0.3 |
| P5 | 1,241 | 30.3 | 25.2 |
| P6 | 1,727 | 42.2 | 35.1 |
| P7 | 6 | 0.1 | 0.1 |
| P8 | 0 | 0.0 | 0.0 |
| P9 | 2,576 | 63.0 | 52.3 |

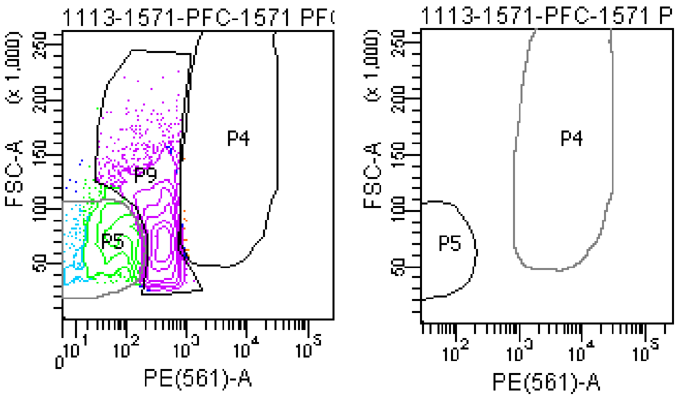

### FACSDiva Version 6.1.3

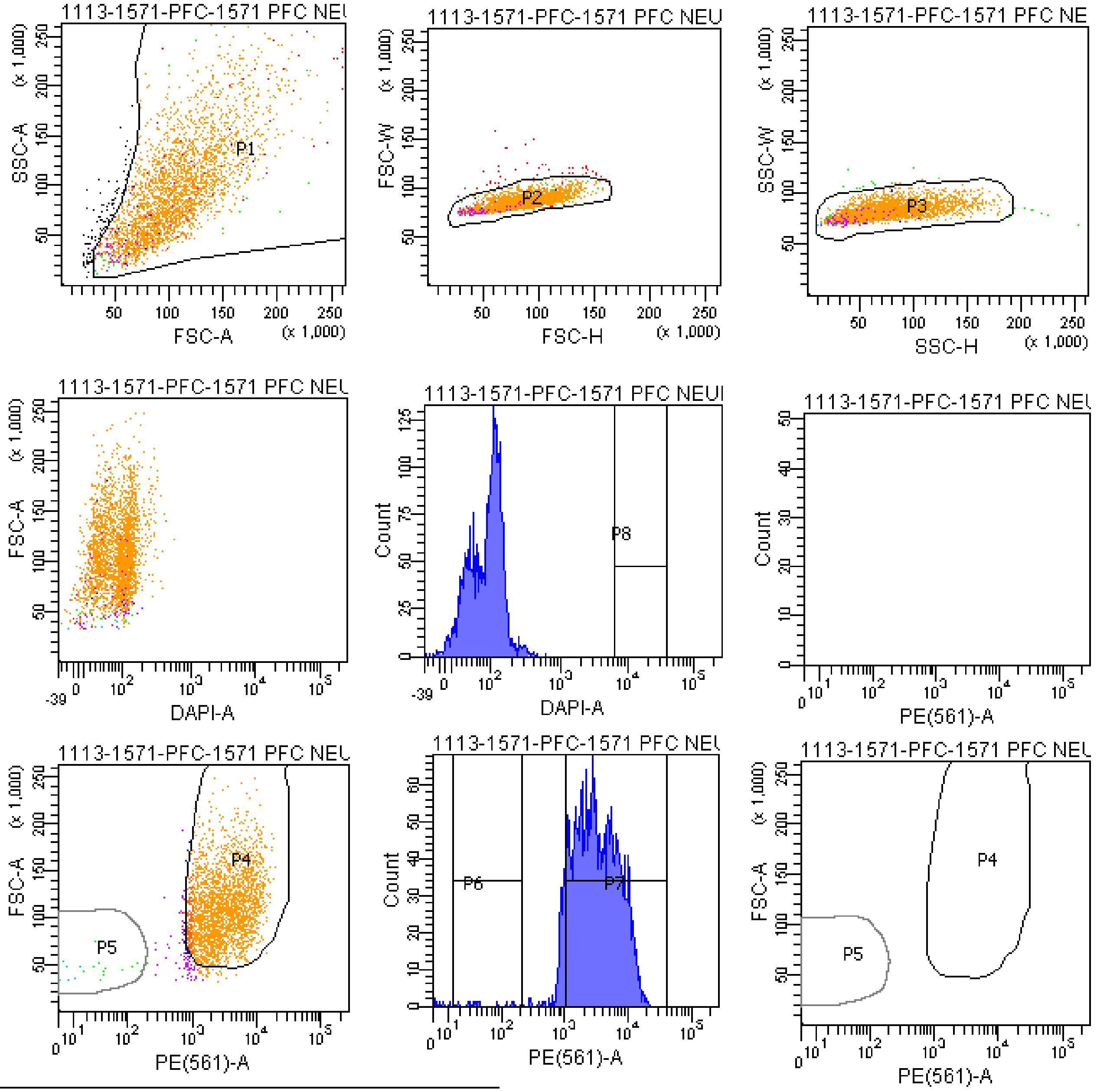


Tube: 1571 PFC NEURON BACK

| Population | #Events | %Parent | %Total |
|---|---|---|---|
| All Events | 3,149 | #### | 100.0 |
| P1 | 3,051 | 96.9 | 96.9 |
| P2 | 3,002 | 98.4 | 95.3 |
| P3 | 2,982 | 99.3 | 94.7 |
| P4 | 2,811 | 94.3 | 89.3 |
| P5 | 20 | 0.7 | 0.6 |
| P6 | 13 | 0.4 | 0.4 |
| P7 | 2,709 | 90.8 | 86.0 |
| P8 | 0 | 0.0 | 0.0 |
| P9 | 67 | 2.2 | 2.1 |

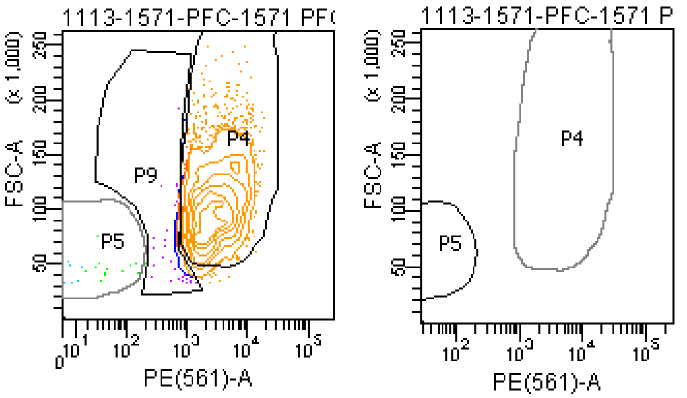

**The BD Arial II FACS sorting reports of UMB#4516**

Sample: prefrontal cortex

Category: Rett patient

### FACSDiva Version 6.1.3

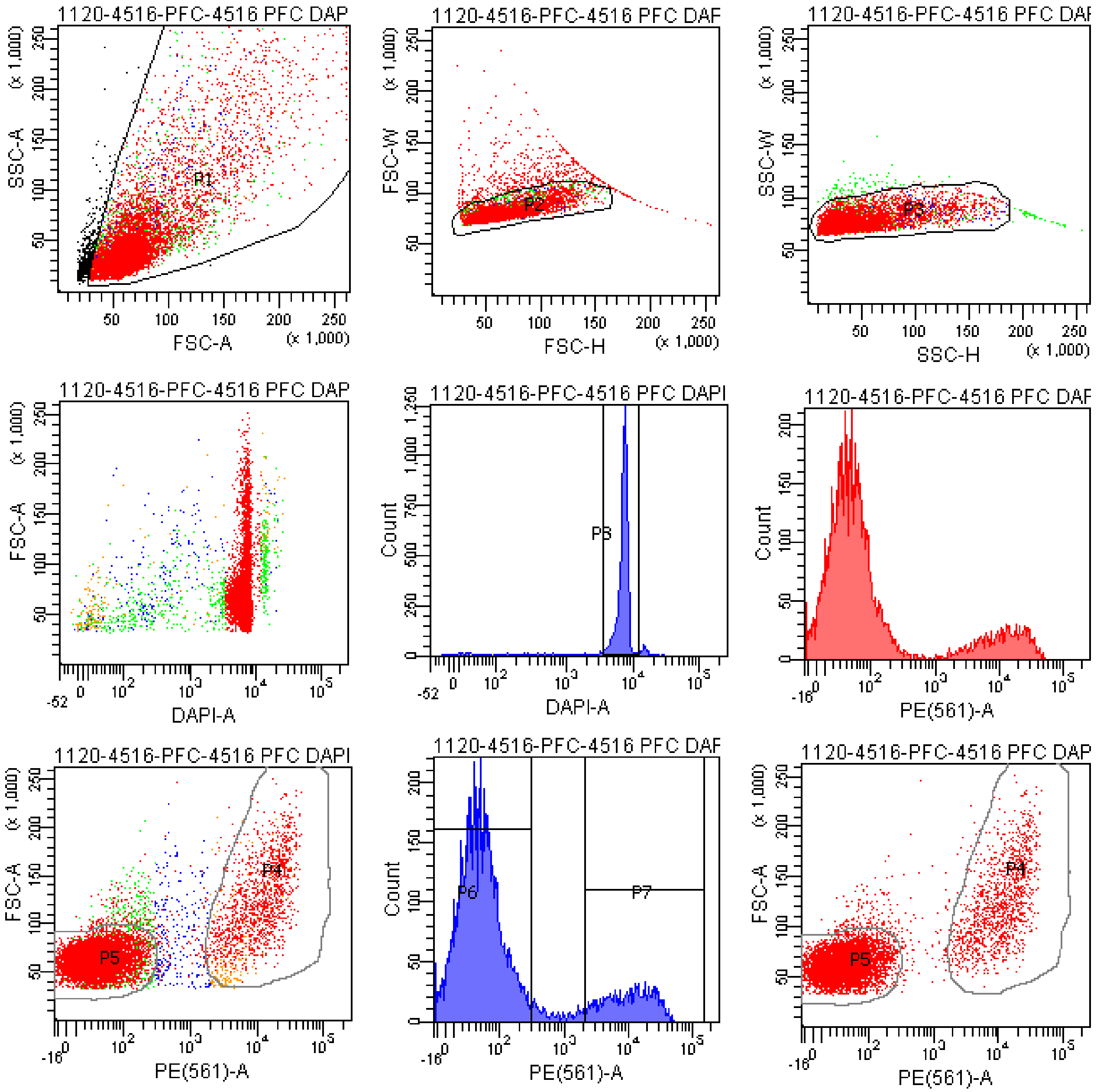


Tube: 4516 PFC DAPI+PE

| Population | #Events | %Parent | %Total |
|---|---|---|---|
| All Events | 10,582 | #### | 100.0 |
| P1 | 10,025 | 94.7 | 94.7 |
| P2 | 9,486 | 94.6 | 89.6 |
| P3 | 9,343 | 98.5 | 88.3 |
| P4 | 1,411 | 15.1 | 13.3 |
| P5 | 7,324 | 78.4 | 69.2 |
| P6 | 7,590 | 81.2 | 71.7 |
| P7 | 1,429 | 15.3 | 13.5 |
| P8 | 8,346 | 89.3 | 78.9 |
| P9 | 481 | 5.1 | 4.5 |

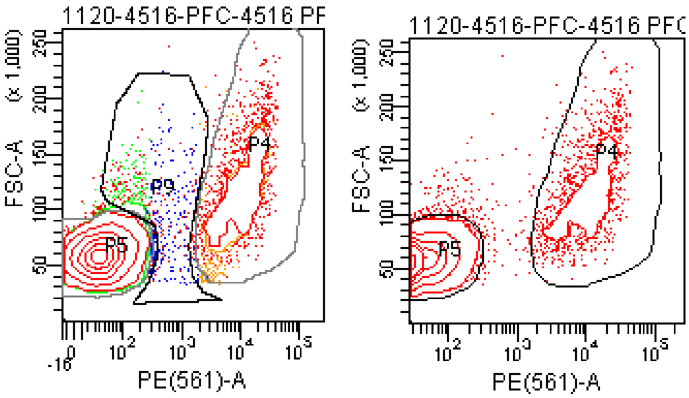

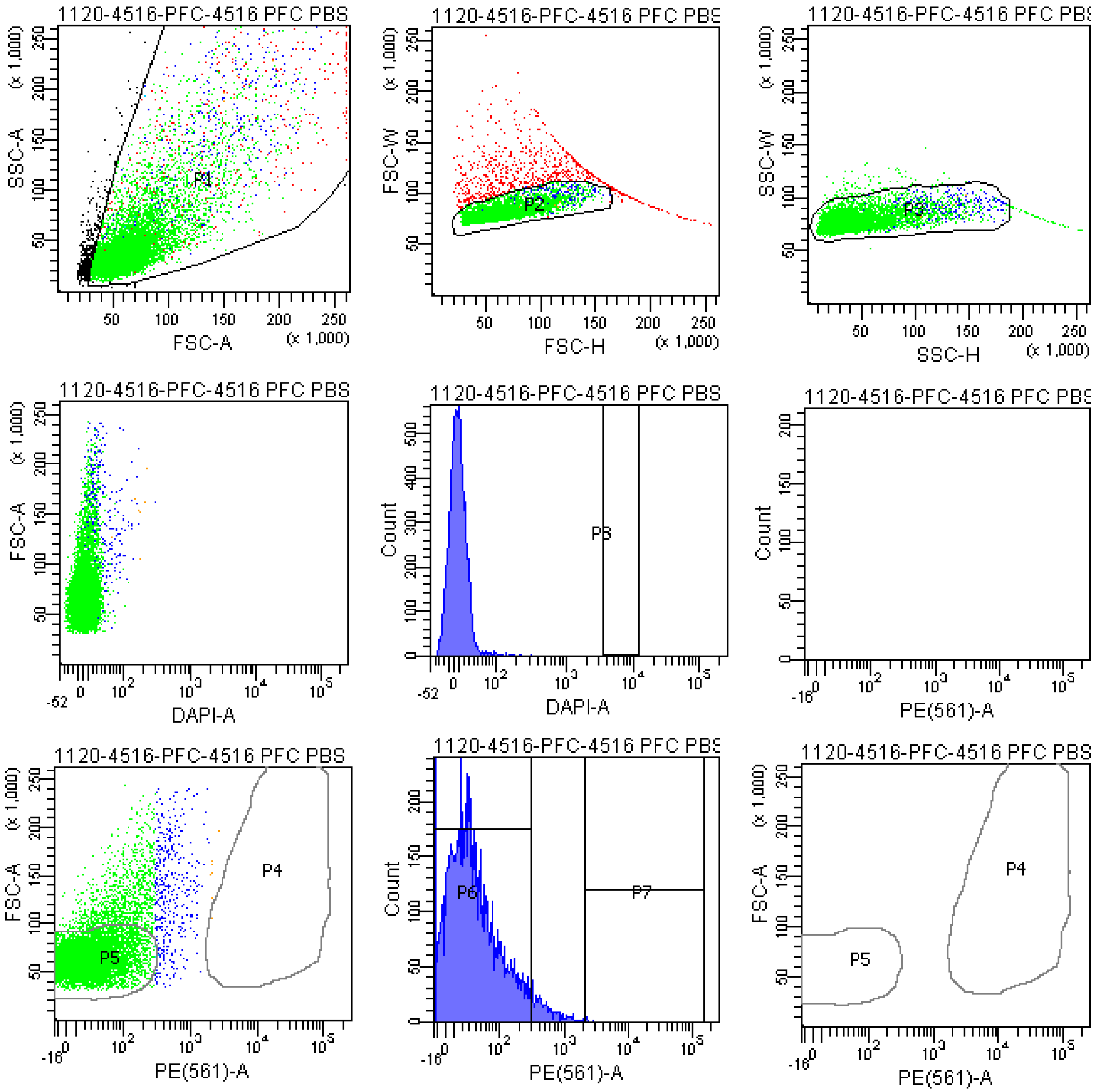


Tube: 4516 PFC PBS CONTROL

| Population | #Events | %Parent | %Total |
|---|---|---|---|
| All Events | 10,656 | #### | 100.0 |
| P1 | 10,026 | 94.1 | 94.1 |
| P2 | 9,237 | 92.1 | 86.7 |
| P3 | 9,094 | 98.5 | 85.3 |
| P4 | 1 | 0.0 | 0.0 |
| P5 | 7,662 | 84.3 | 71.9 |
| P6 | 8,658 | 95.2 | 81.2 |
| P7 | 7 | 0.1 | 0.1 |
| P8 | 0 | 0.0 | 0.0 |
| P9 | 1,107 | 12.2 | 10.4 |

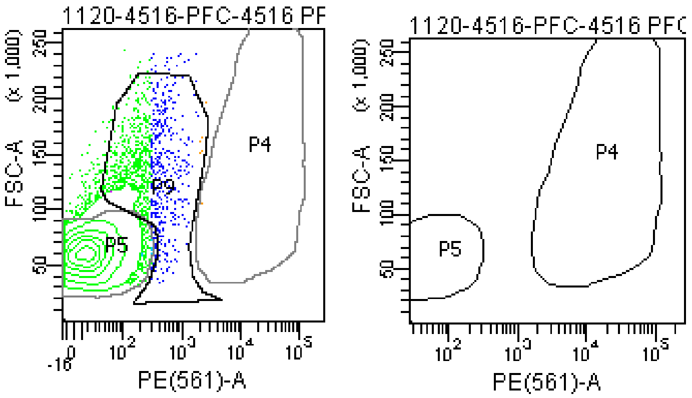

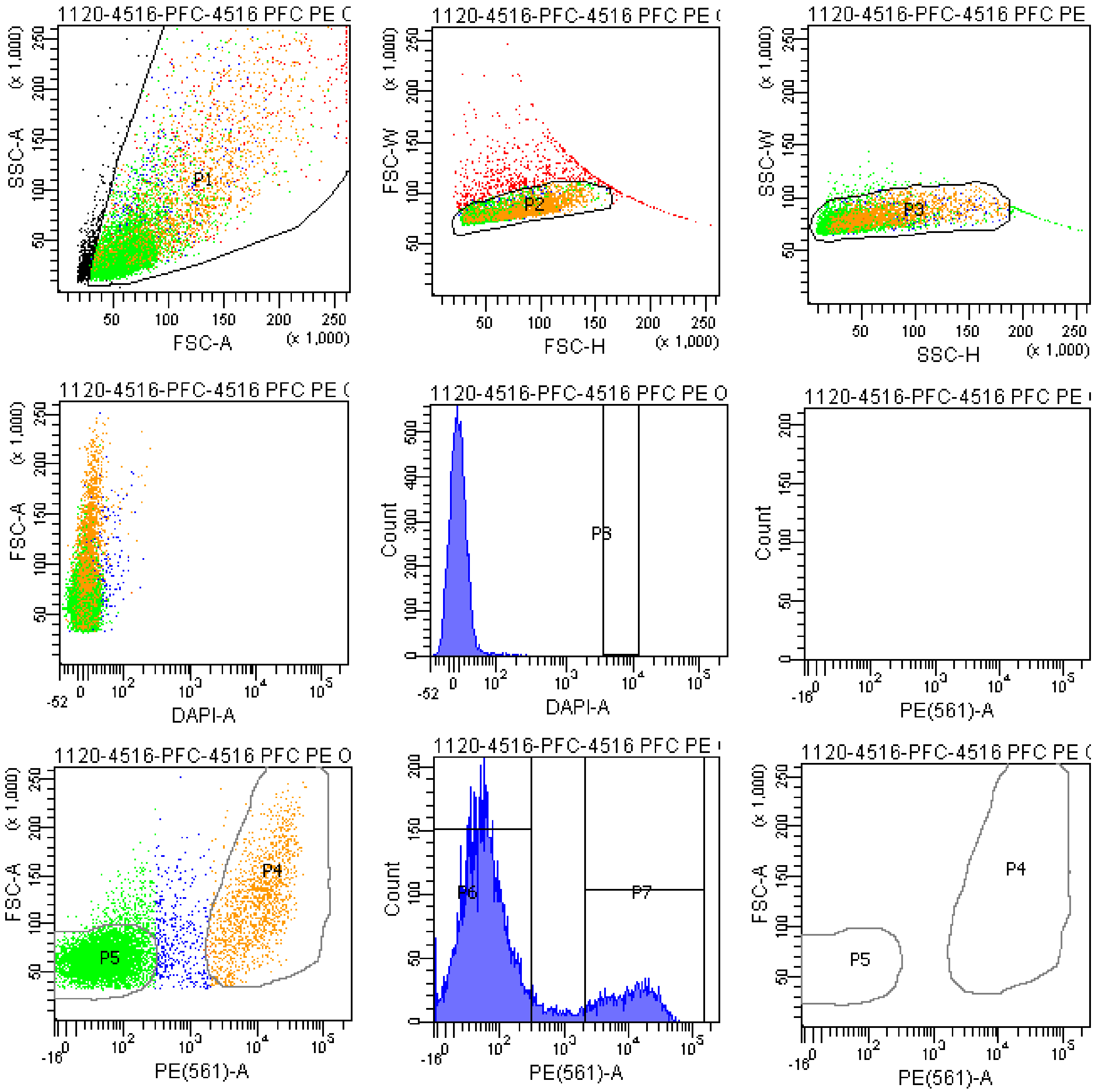


Tube: 4516 PFC PE ONLY CONTROL

| Population | #Events | %Parent | %Total |
|---|---|---|---|
| All Events | 10,485 | #### | 100.0 |
| P1 | 9,866 | 94.1 | 94.1 |
| P2 | 9,196 | 93.2 | 87.7 |
| P3 | 9,048 | 98.4 | 86.3 |
| P4 | 1,399 | 15.5 | 13.3 |
| P5 | 6,903 | 76.3 | 65.8 |
| P6 | 7,202 | 79.6 | 68.7 |
| P7 | 1,413 | 15.6 | 13.5 |
| P8 | 0 | 0.0 | 0.0 |
| P9 | 603 | 6.7 | 5.8 |

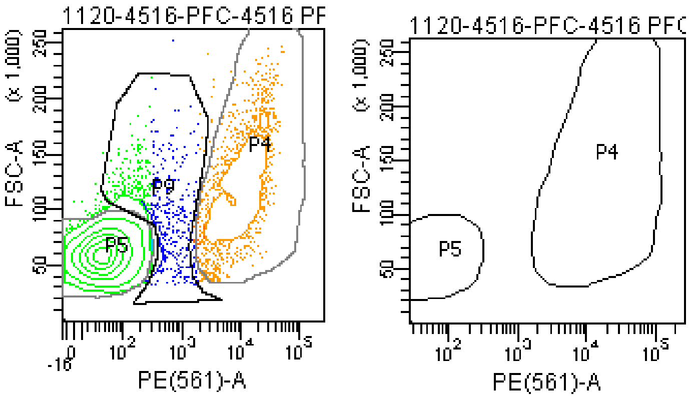

### FACSDiva Version 6.1.3

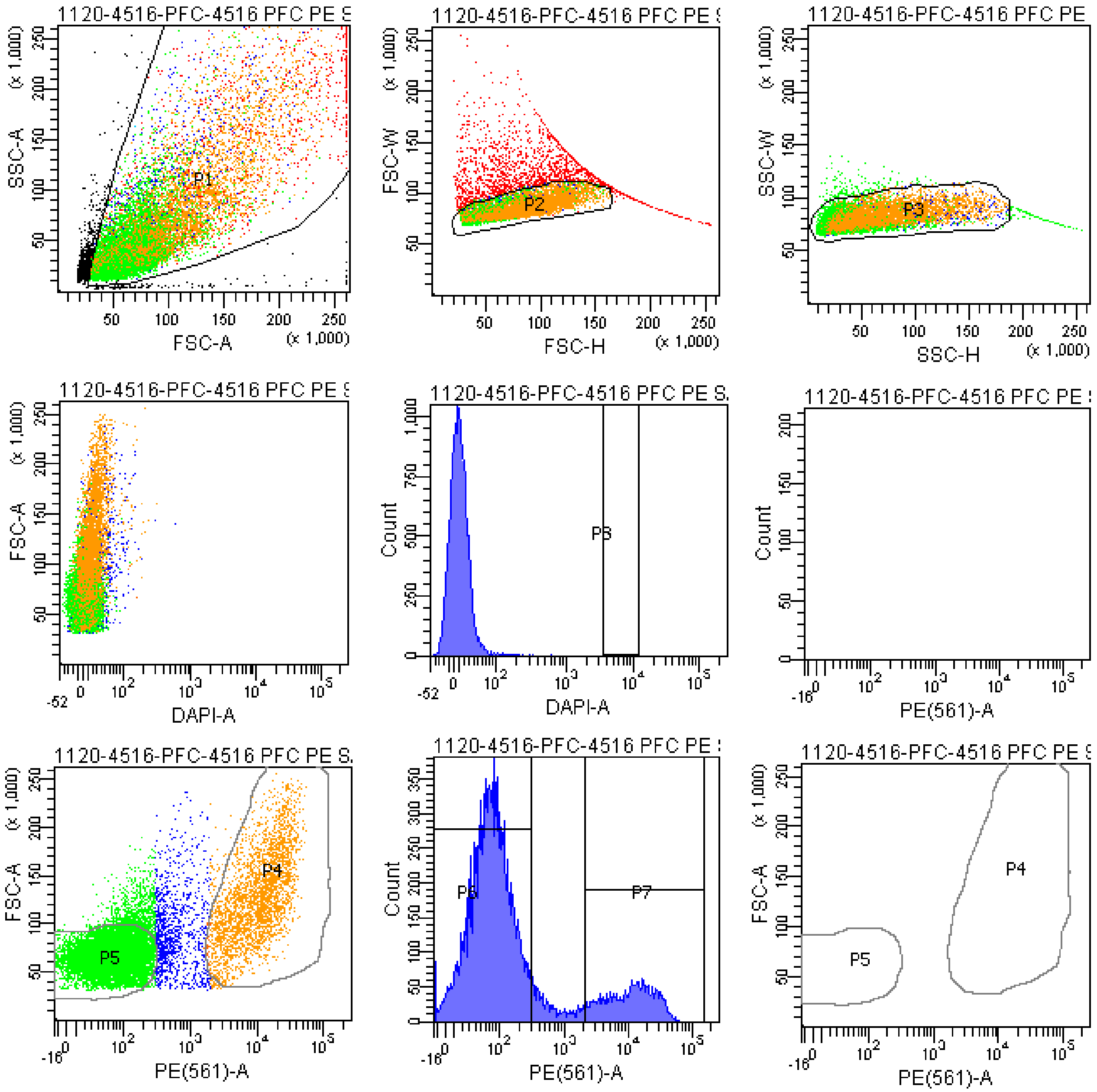


Tube: 4516 PFC PE SAMPLE

| Population | #Events | %Parent | %Total |
|---|---|---|---|
| All Events | 21,013 | #### | 100.0 |
| P1 | 20,000 | 95.2 | 95.2 |
| P2 | 18,205 | 91.0 | 86.6 |
| P3 | 17,966 | 98.7 | 85.5 |
| P4 | 2,808 | 15.6 | 13.4 |
| P5 | 13,366 | 74.4 | 63.6 |
| P6 | 14,065 | 78.3 | 66.9 |
| P7 | 2,843 | 15.8 | 13.5 |
| P8 | 0 | 0.0 | 0.0 |
| P9 | 1,537 | 8.6 | 7.3 |

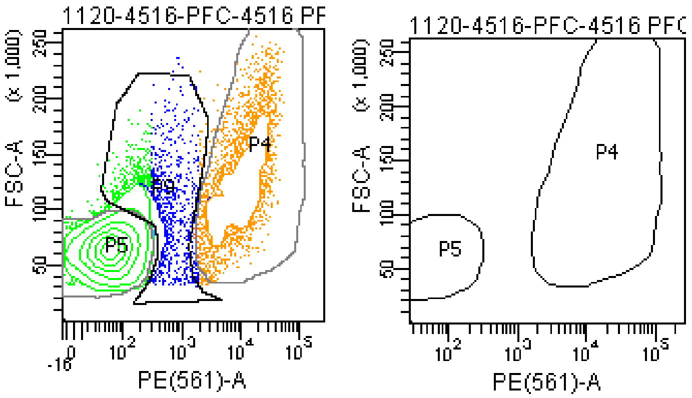

### FACSDiva Version 6.1.3

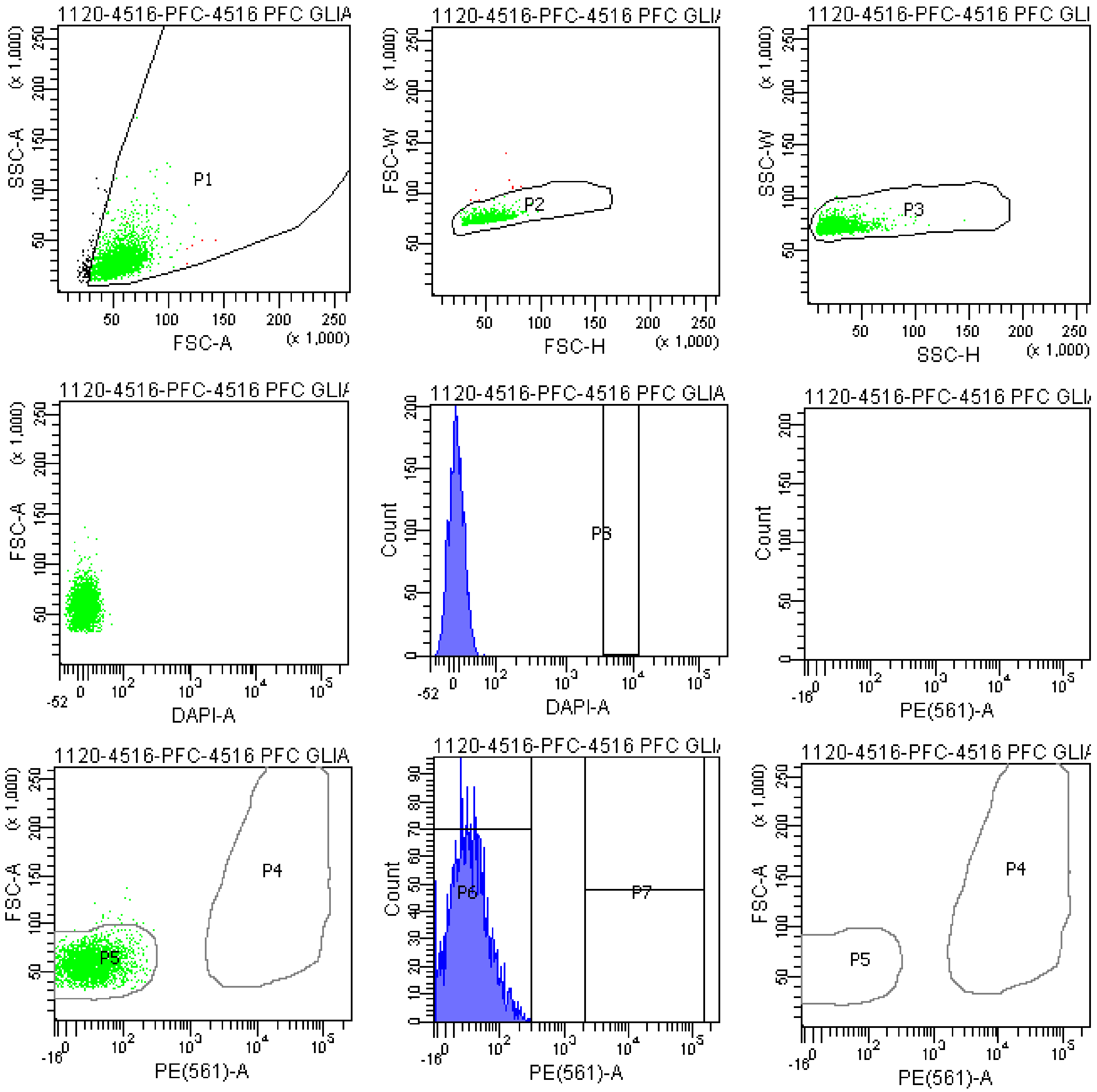


Tube: 4516 PFC GLIA BACK

| Population | #Events | %Parent | %Total |
|---|---|---|---|
| All Events | 2,964 | #### | 100.0 |
| P1 | 2,864 | 96.6 | 96.6 |
| P2 | 2,856 | 99.7 | 96.4 |
| P3 | 2,854 | 99.9 | 96.3 |
| P4 | 0 | 0.0 | 0.0 |
| P5 | 2,835 | 99.3 | 95.6 |
| P6 | 2,854 | 100.0 | 96.3 |
| P7 | 0 | 0.0 | 0.0 |
| P8 | 0 | 0.0 | 0.0 |
| P9 | 11 | 0.4 | 0.4 |

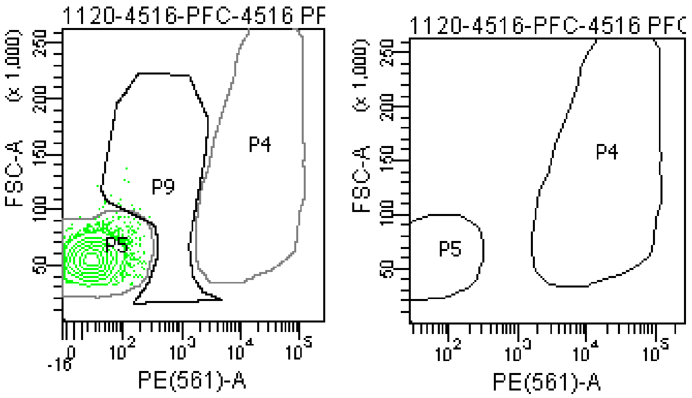

### FACSDiva Version 6.1.3

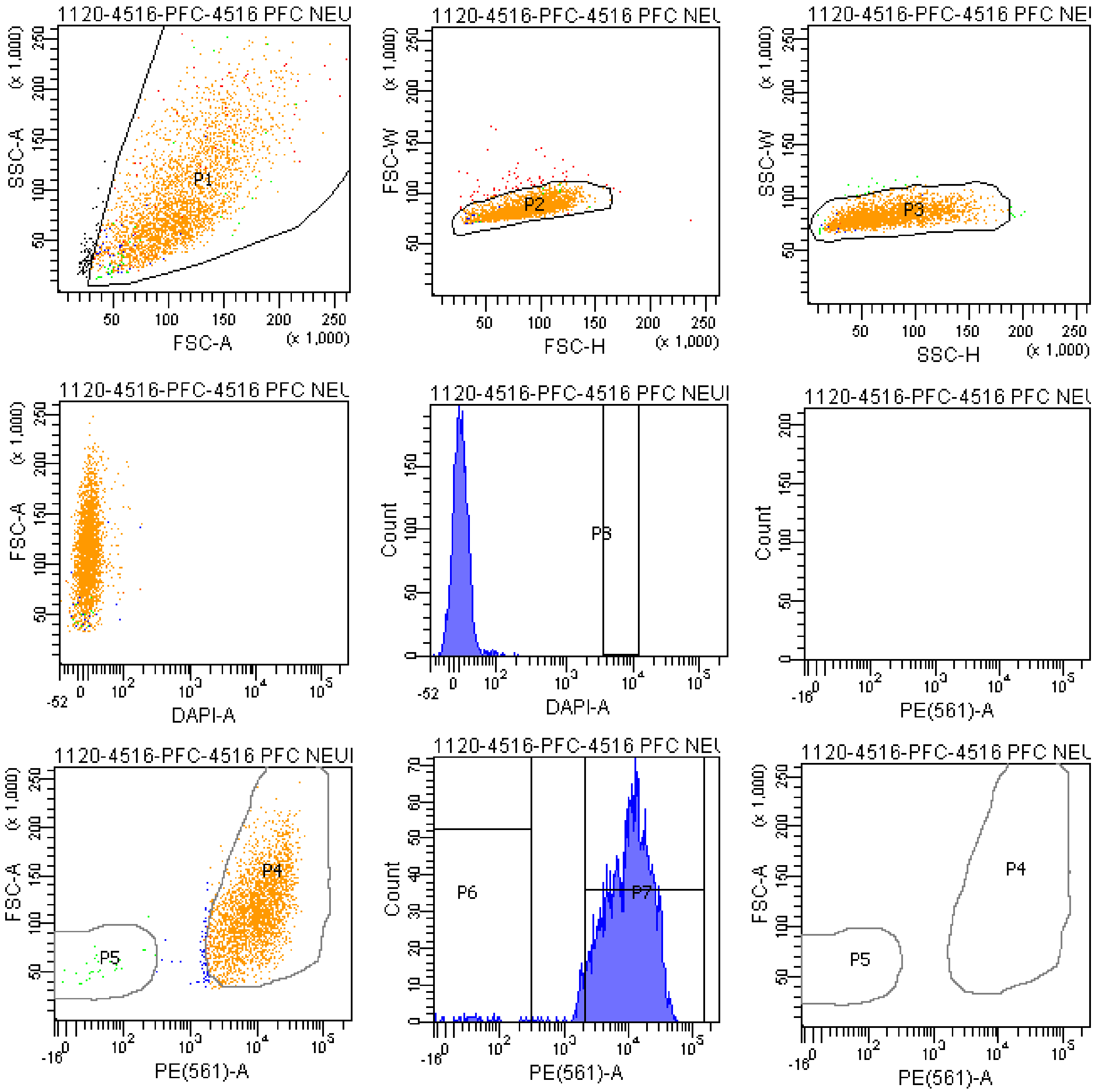


Tube: 4516 PFC NEURON BACK

| Population | #Events | %Parent | %Total |
|---|---|---|---|
| All Events | 3,072 | #### | 100.0 |
| P1 | 2,984 | 97.1 | 97.1 |
| P2 | 2,918 | 97.8 | 95.0 |
| P3 | 2,899 | 99.3 | 94.4 |
| P4 | 2,787 | 96.1 | 90.7 |
| P5 | 29 | 1.0 | 0.9 |
| P6 | 30 | 1.0 | 1.0 |
| P7 | 2,778 | 95.8 | 90.4 |
| P8 | 0 | 0.0 | 0.0 |
| P9 | 20 | 0.7 | 0.7 |

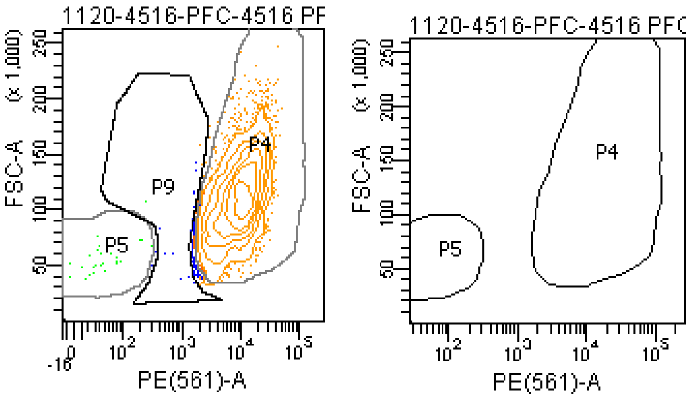

**The BD Arial II FACS sorting reports of UMB#1846**

Sample: prefrontal cortex

Category: healthy control

### FACSDiva Version 6.1.3

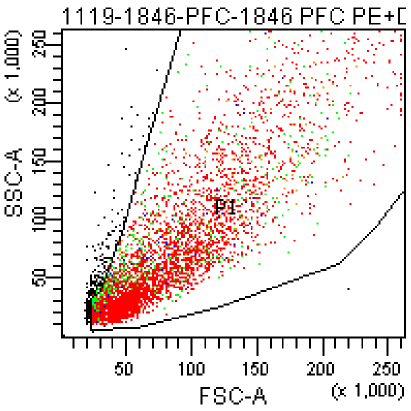


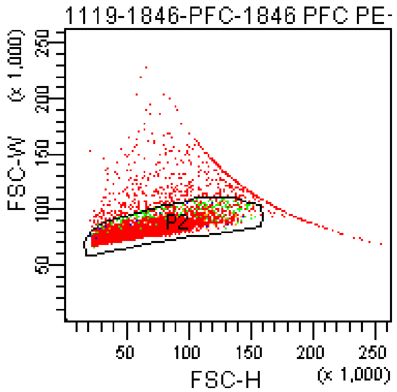


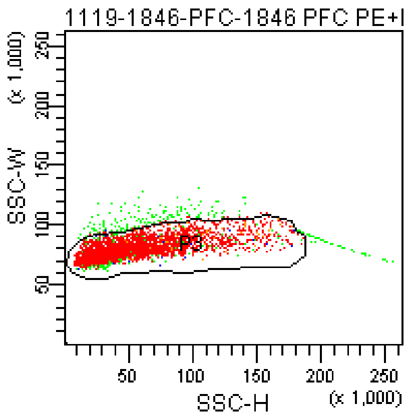


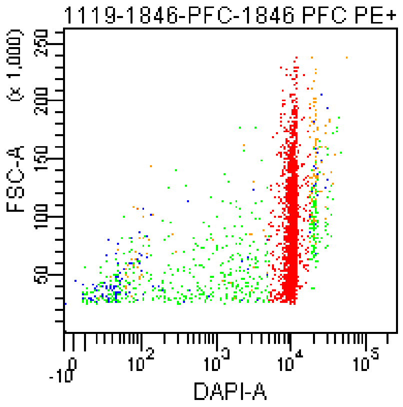


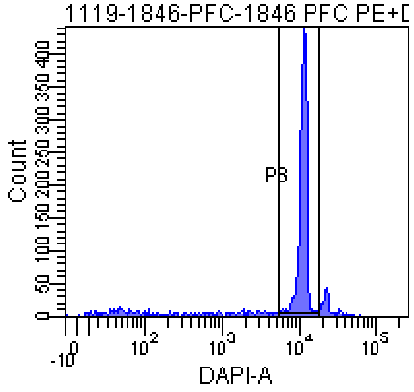


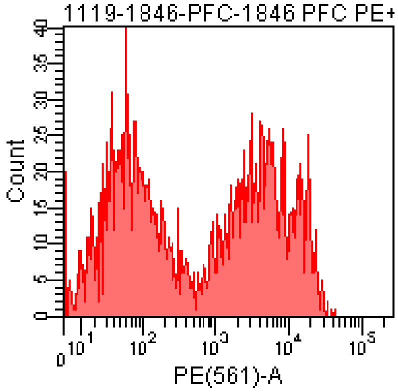


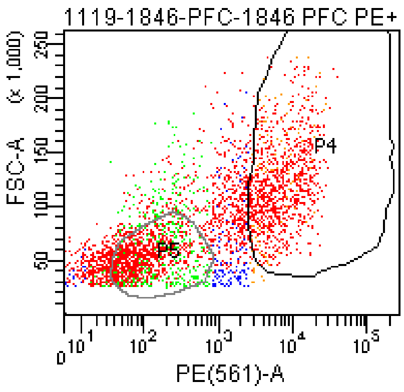


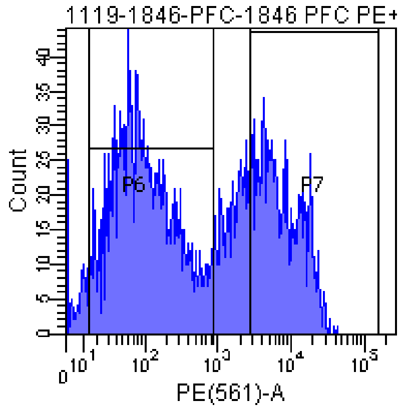


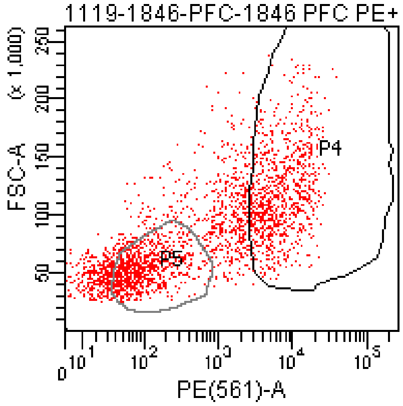


Tube: 1846 PFC PE+DAPI 400V

| Population | #Events | %Parent | %Total |
|---|---|---|---|
| All Events | 4,559 | #### | 100.0 |
| P1 | 4,289 | 94.1 | 94.1 |
| P2 | 3,664 | 85.4 | 80.4 |
| P3 | 3,467 | 94.6 | 76.0 |
| P4 | 1,056 | 30.5 | 23.2 |
| P5 | 1,189 | 34.3 | 26.1 |
| P6 | 1,778 | 51.3 | 39.0 |
| P7 | 1,059 | 30.5 | 23.2 |
| P8 | 2,671 | 77.0 | 58.6 |
| P9 | 653 | 18.8 | 14.3 |

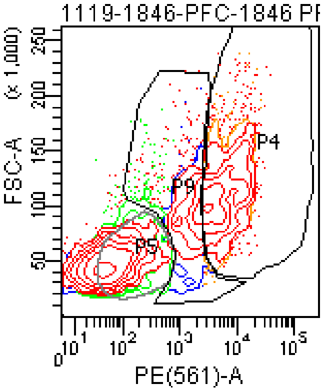


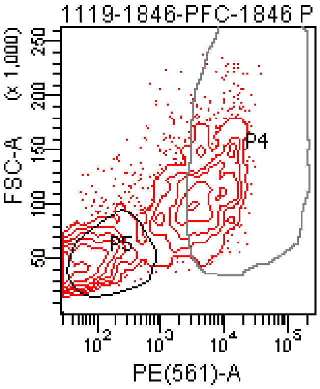

### FACSDiva Version 6.1.3

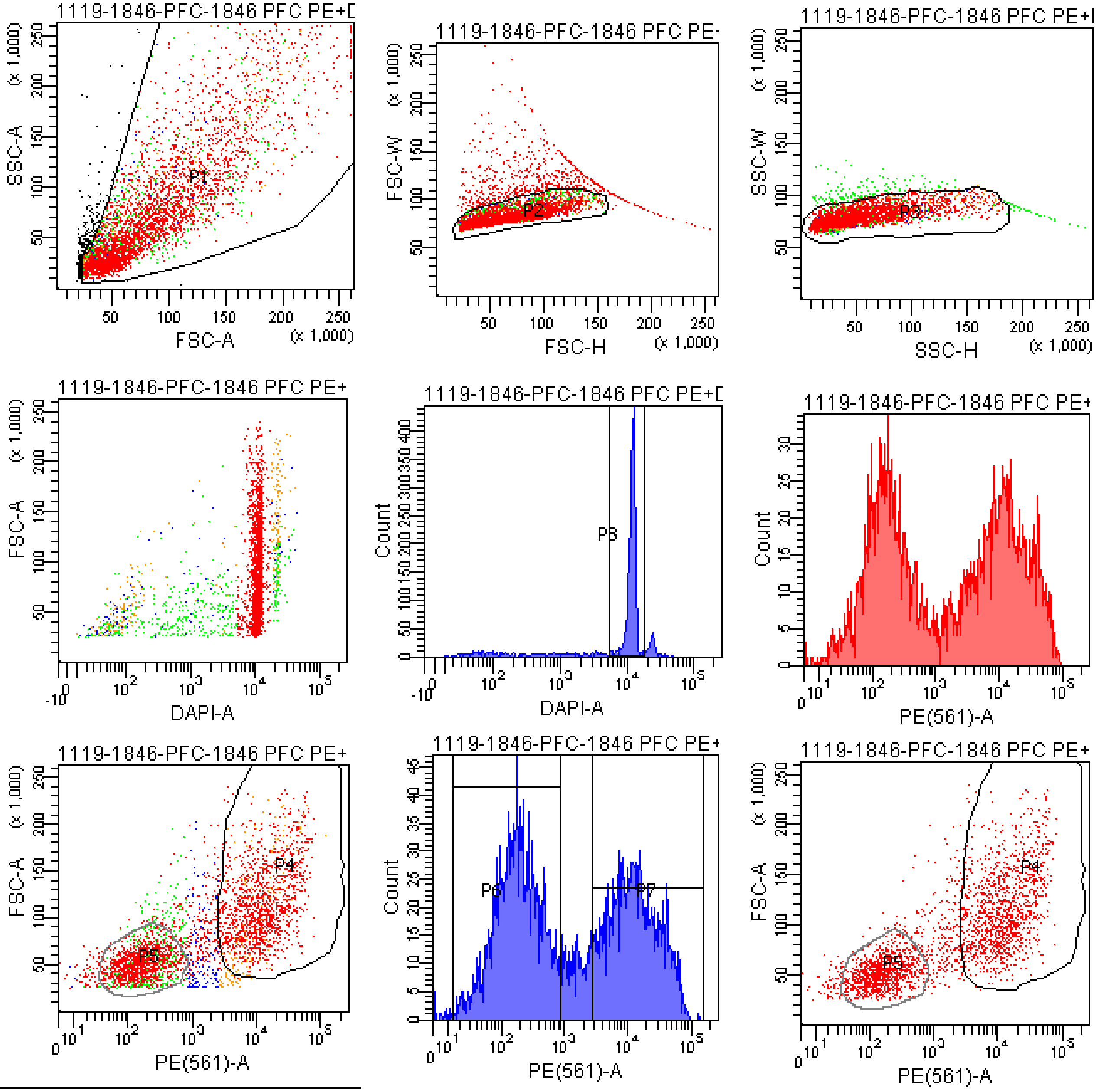


Tube: 1846 PFC PE+DAPI 457V

| Population | #Events | %Parent | %Total |
|---|---|---|---|
| All Events | 4,480 | #### | 100.0 |
| P1 | 4,218 | 94.2 | 94.2 |
| P2 | 3,577 | 84.8 | 79.8 |
| P3 | 3,398 | 95.0 | 75.8 |
| P4 | 1,416 | 41.7 | 31.6 |
| P5 | 1,414 | 41.6 | 31.6 |
| P6 | 1,623 | 47.8 | 36.2 |
| P7 | 1,460 | 43.0 | 32.6 |
| P8 | 2,640 | 77.7 | 58.9 |
| P9 | 438 | 12.9 | 9.8 |

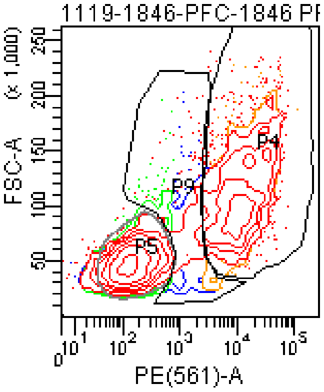


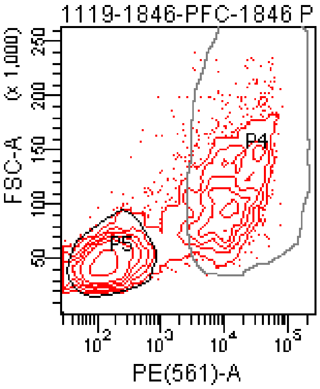

### FACSDiva Version 6.1.3

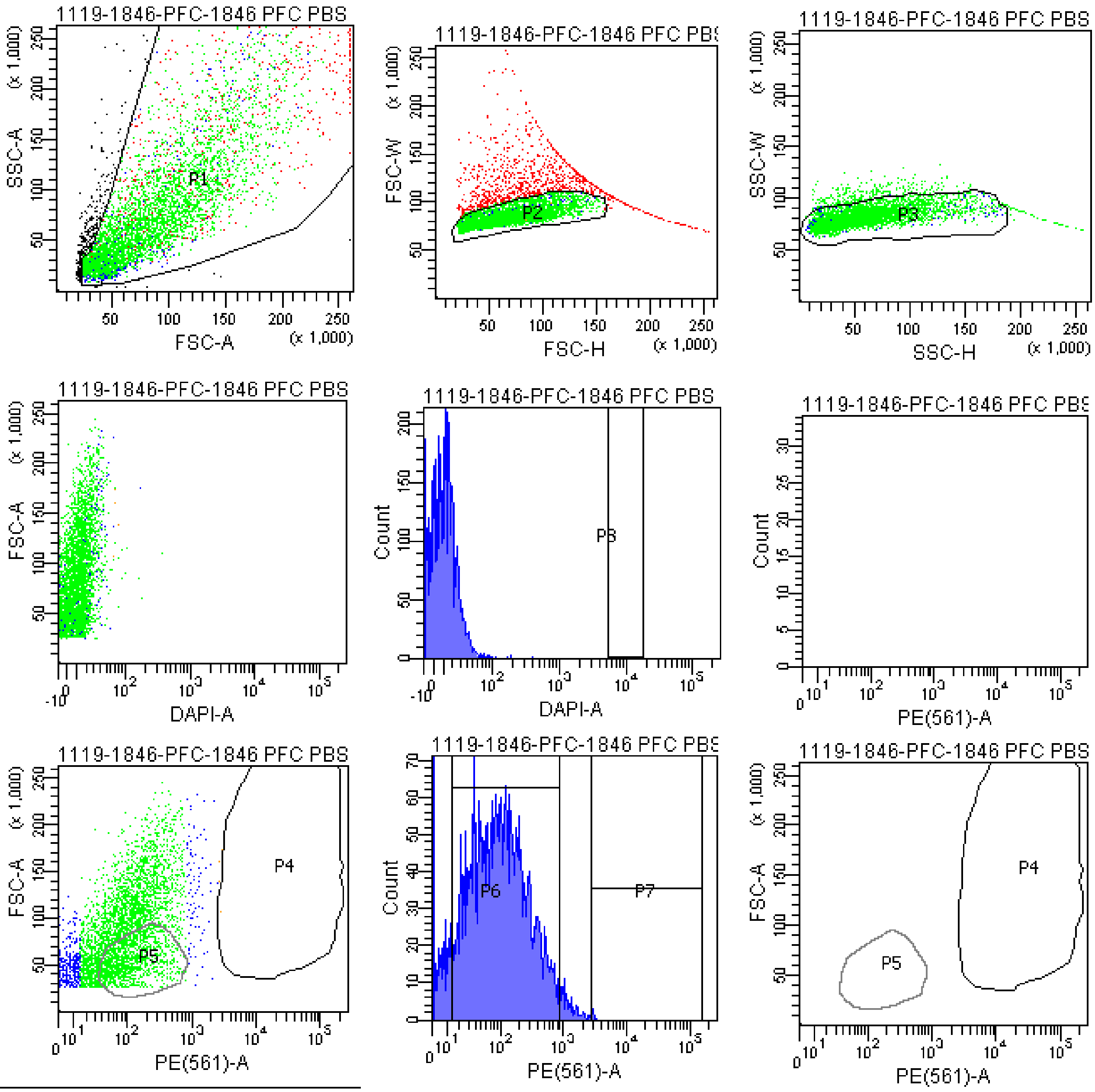


| Tube: 1846 PFC PBS CONTROL | | | |
|---|---|---|---|
| **Population** | **#Events** | **%Parent** | **%Total** |
| All Events | 5,760 | #### | 100.0 |
| P1 | 5,418 | 94.1 | 94.1 |
| P2 | 4,329 | 79.9 | 75.2 |
| P3 | 4,057 | 93.7 | 70.4 |
| P4 | 2 | 0.0 | 0.0 |
| P5 | 1,214 | 29.9 | 21.1 |
| P6 | 3,466 | 85.4 | 60.2 |
| P7 | 4 | 0.1 | 0.1 |
| P8 | 0 | 0.0 | 0.0 |
| P9 | 867 | 21.4 | 15.1 |

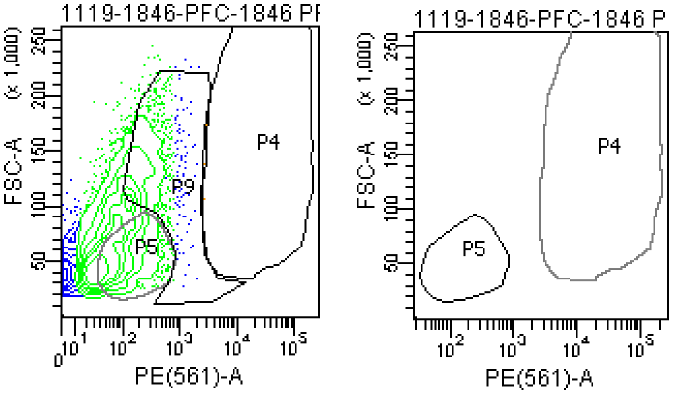

### FACSDiva Version 6.1.3

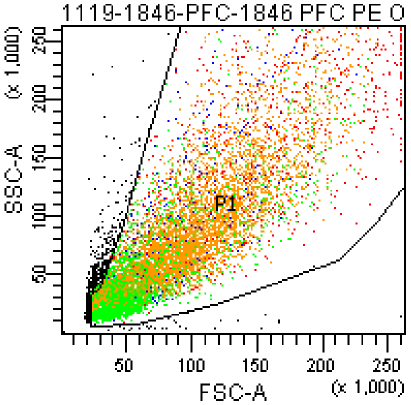


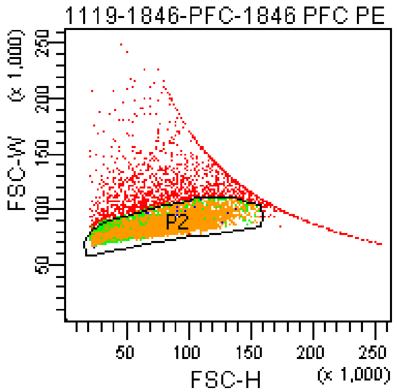


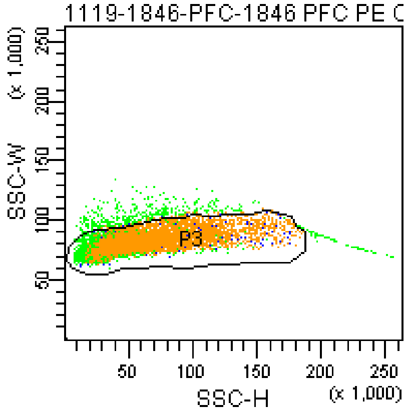


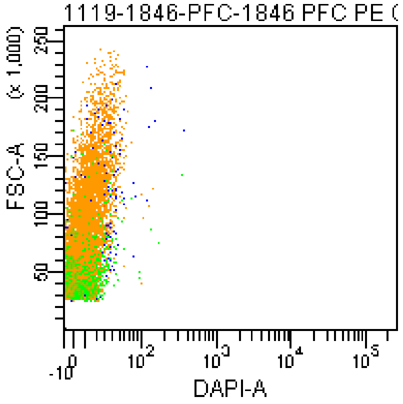


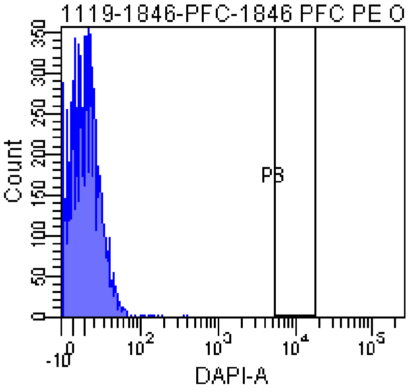


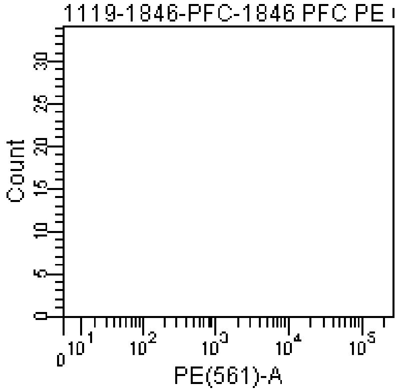


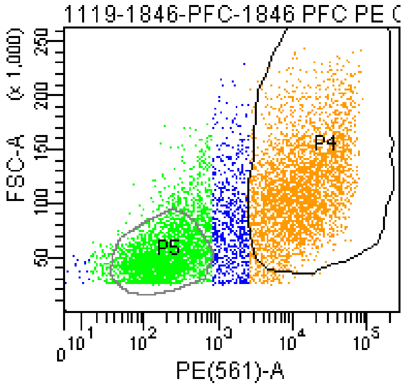


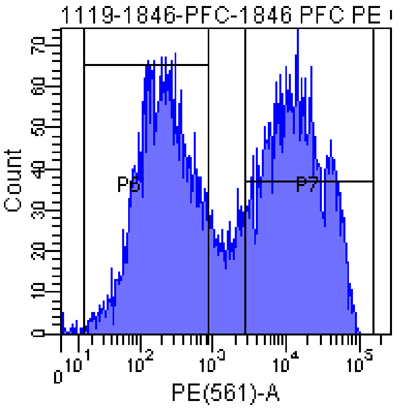


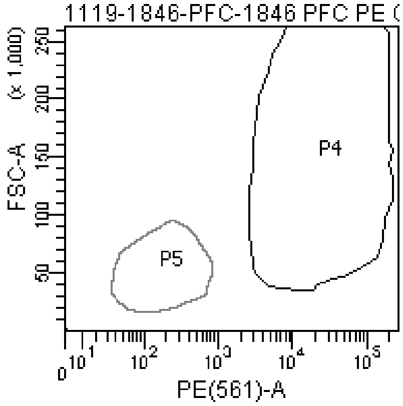


Tube: 1846 PFC PE ONLY CONTROL

| Population | #Events | %Parent | %Total |
|---|---|---|---|
| All Events | 10,146 | #### | 100.0 |
| P1 | 9,589 | 94.5 | 94.5 |
| P2 | 7,909 | 82.5 | 78.0 |
| P3 | 7,464 | 94.4 | 73.6 |
| P4 | 3,408 | 45.7 | 33.6 |
| P5 | 2,737 | 36.7 | 27.0 |
| P6 | 3,231 | 43.3 | 31.8 |
| P7 | 3,530 | 47.3 | 34.8 |
| P8 | 0 | 0.0 | 0.0 |
| P9 | 1,116 | 15.0 | 11.0 |

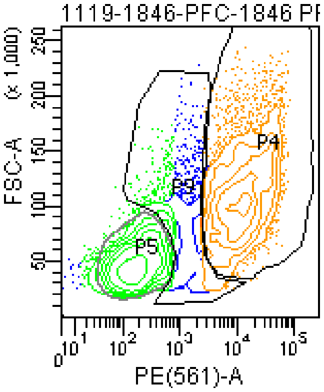


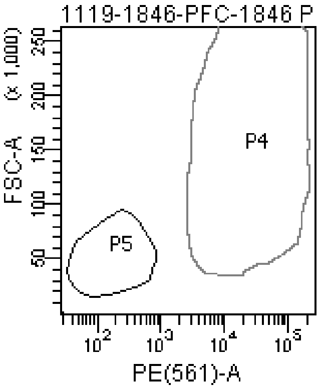

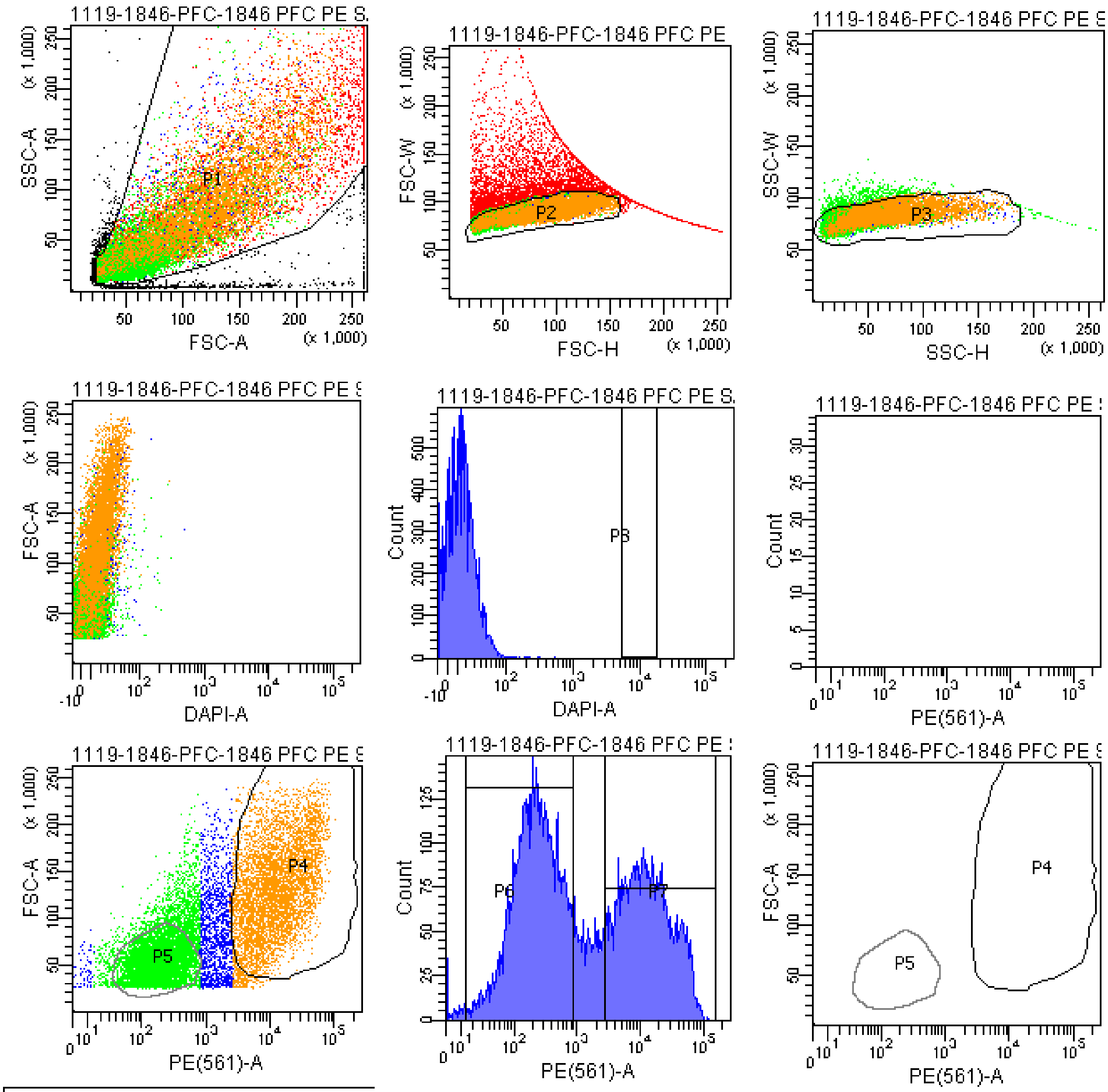


Tube: 1846 PFC PE SAMPLE

| Population | #Events | %Parent | %Total |
|---|---|---|---|
| All Events | 21,428 | #### | 100.0 |
| P1 | 20,000 | 93.3 | 93.3 |
| P2 | 13,835 | 69.2 | 64.6 |
| P3 | 13,341 | 96.4 | 62.3 |
| P4 | 5,175 | 38.8 | 24.2 |
| P5 | 5,177 | 38.8 | 24.2 |
| P6 | 6,553 | 49.1 | 30.6 |
| P7 | 5,380 | 40.3 | 25.1 |
| P8 | 0 | 0.0 | 0.0 |
| P9 | 2,416 | 18.1 | 11.3 |

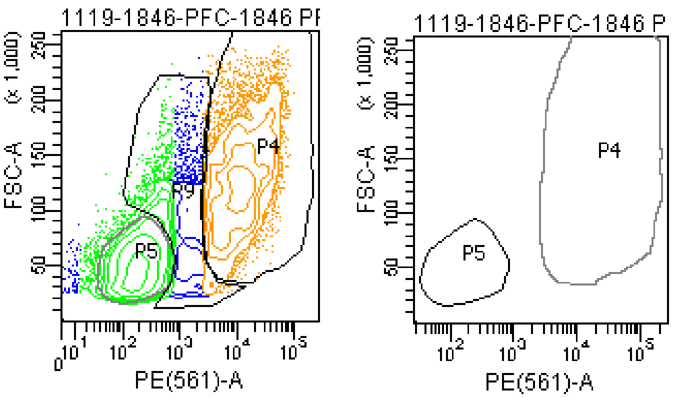

### FACSDiva Version 6.1.3

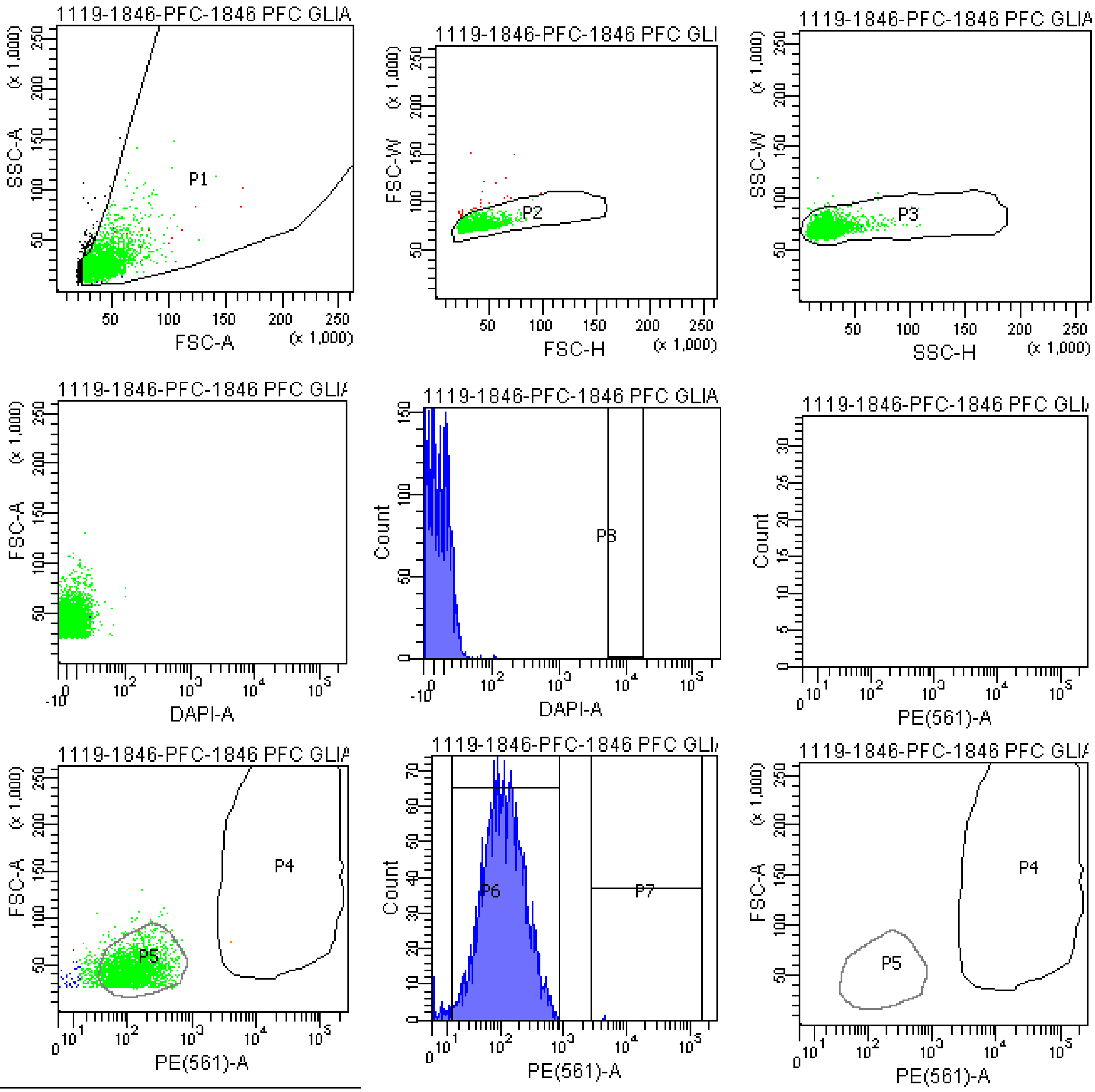


Tube: 1846 PFC GLIA BACK

| Population | #Events | %Parent | %Total |
|---|---|---|---|
| All Events | 3,330 | #### | 100.0 |
| P1 | 3,025 | 90.8 | 90.8 |
| P2 | 2,998 | 99.1 | 90.0 |
| P3 | 2,977 | 99.3 | 89.4 |
| P4 | 1 | 0.0 | 0.0 |
| P5 | 2,682 | 90.1 | 80.5 |
| P6 | 2,928 | 98.4 | 87.9 |
| P7 | 1 | 0.0 | 0.0 |
| P8 | 0 | 0.0 | 0.0 |
| P9 | 16 | 0.5 | 0.5 |

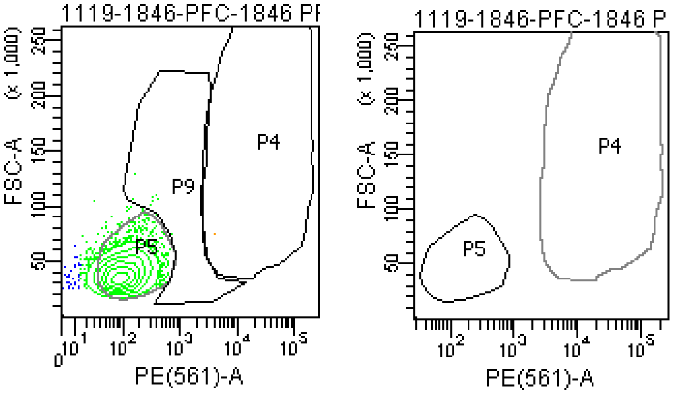

### FACSDiva Version 6.1.3

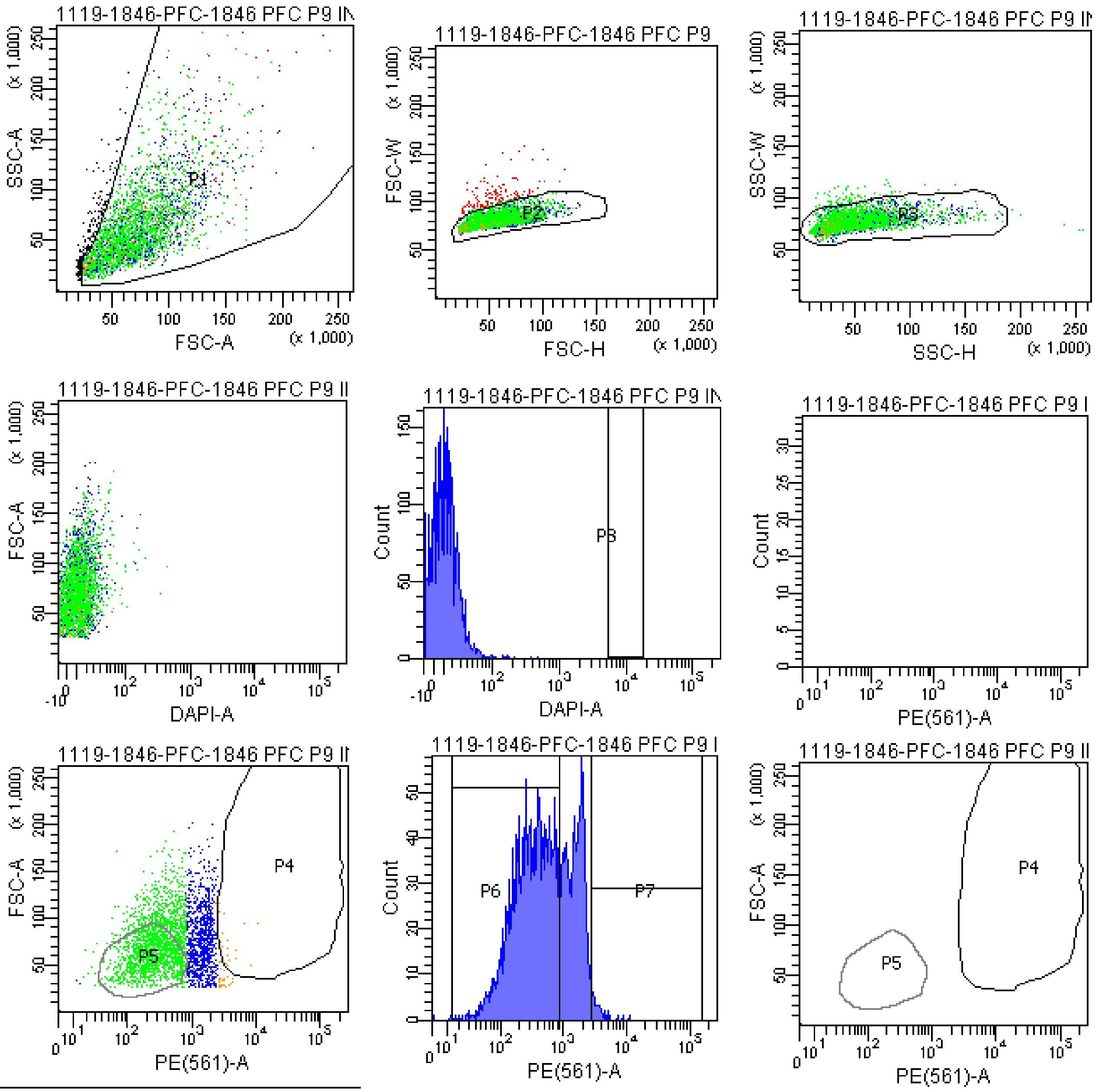


| Tube: 1846 PFC P9 INTER BACK | | | |
|---|---|---|---|
| Population | #Events | %Parent | %Total |
| All Events | 3,379 | #### | 100.0 |
| P1 | 3,195 | 94.6 | 94.6 |
| P2 | 3,054 | 95.6 | 90.4 |
| P3 | 2,985 | 97.7 | 88.3 |
| P4 | 22 | 0.7 | 0.7 |
| P5 | 1,354 | 45.4 | 40.1 |
| P6 | 1,974 | 66.1 | 58.4 |
| P7 | 43 | 1.4 | 1.3 |
| P8 | 0 | 0.0 | 0.0 |
| P9 | 1,483 | 49.7 | 43.9 |

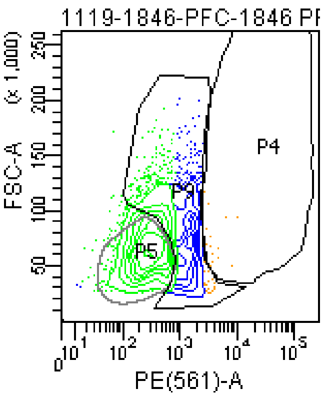


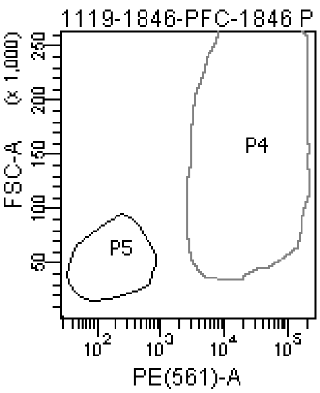

### FACSDiva Version 6.1.3

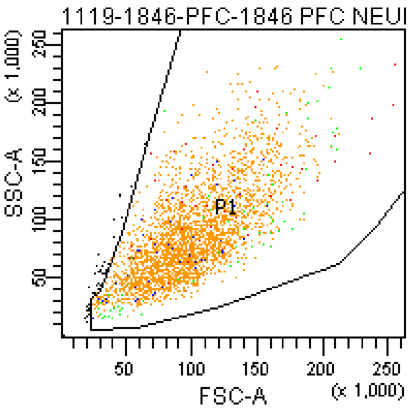


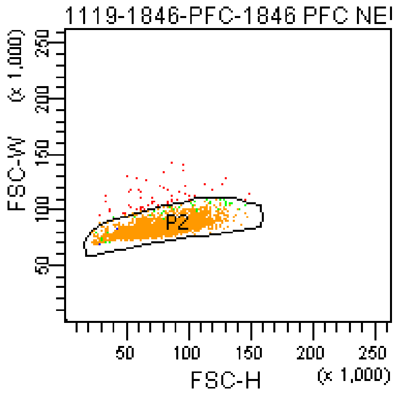


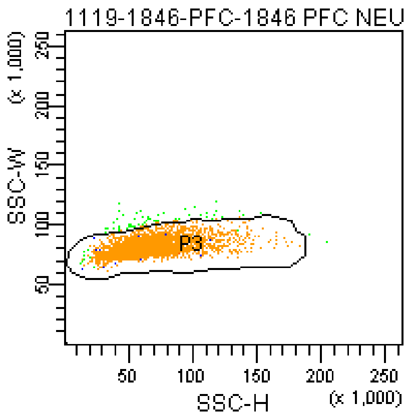


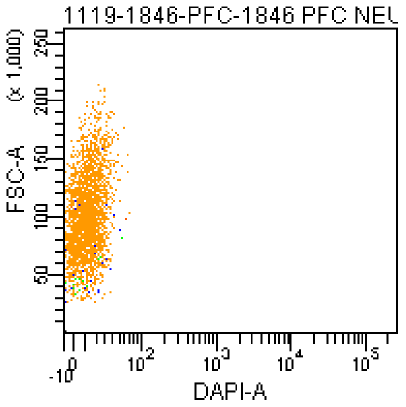


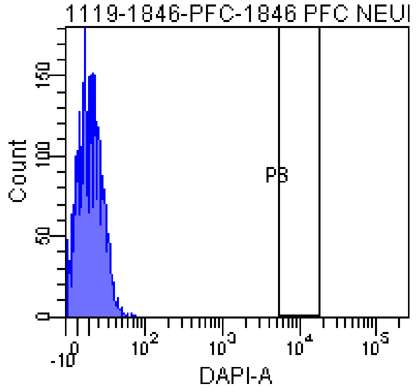


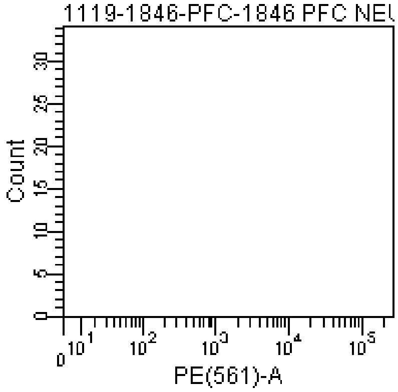


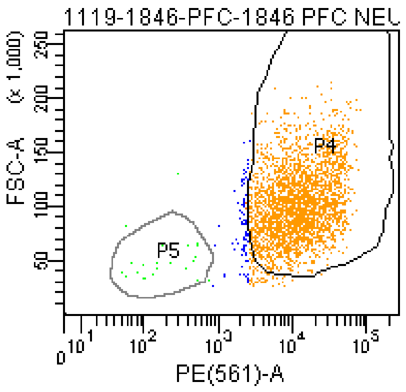


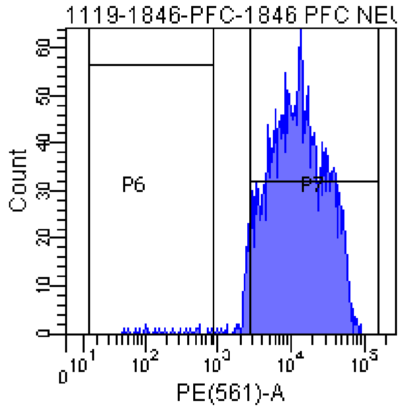


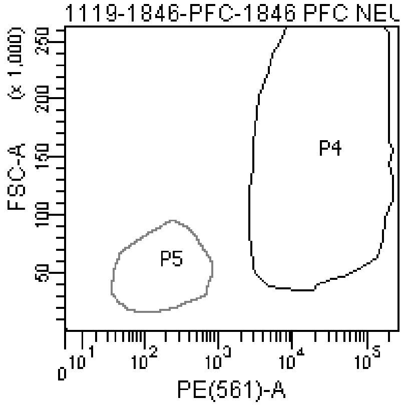


Tube: 1846 PFC NEURON BACK

| Population | #Events | %Parent | %Total |
|---|---|---|---|
| All Events | 3,095 | #### | 100.0 |
| P1 | 3,052 | 98.6 | 98.6 |
| P2 | 2,997 | 98.2 | 96.8 |
| P3 | 2,938 | 98.0 | 94.9 |
| P4 | 2,758 | 93.9 | 89.1 |
| P5 | 19 | 0.6 | 0.6 |
| P6 | 24 | 0.8 | 0.8 |
| P7 | 2,817 | 95.9 | 91.0 |
| P8 | 0 | 0.0 | 0.0 |
| P9 | 137 | 4.7 | 4.4 |

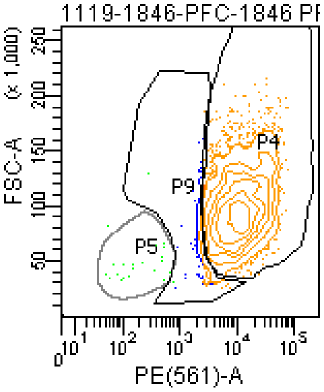


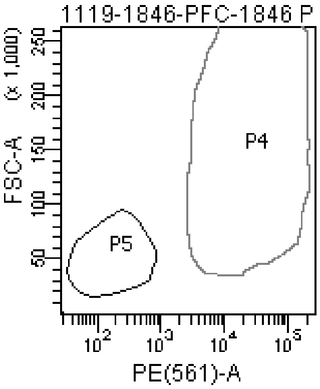

**The BD Arial II FACS sorting reports of UMB#4852**

Sample: prefrontal cortex

Category: Rett patient

### FACSDiva Version 6.1.3

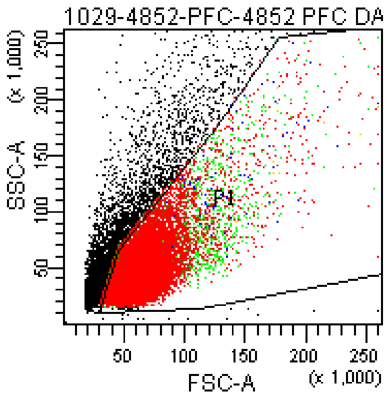


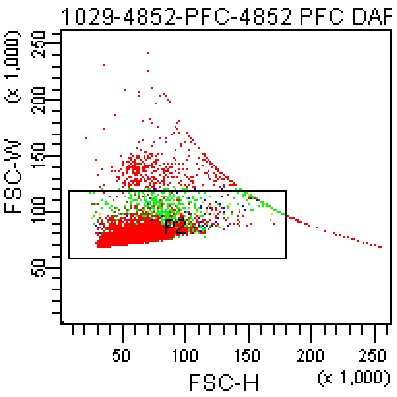


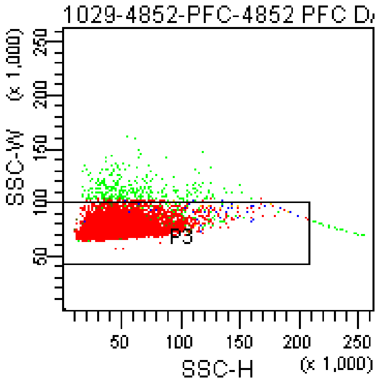


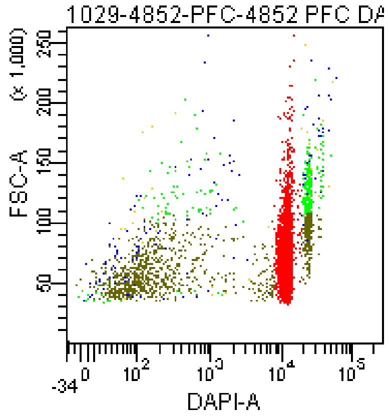


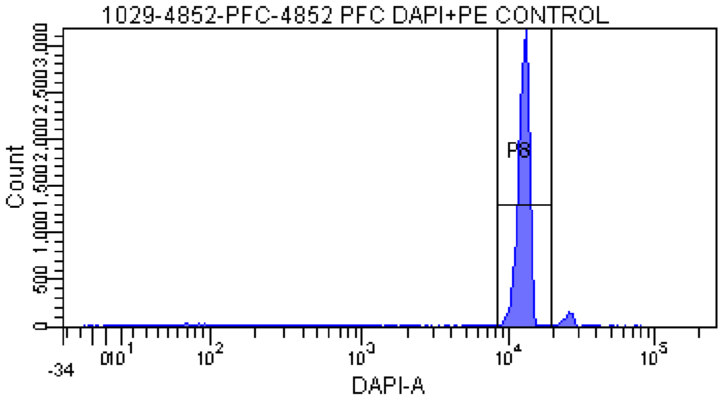


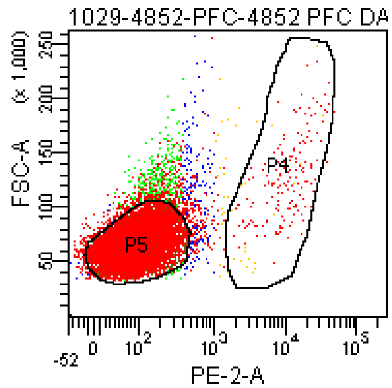


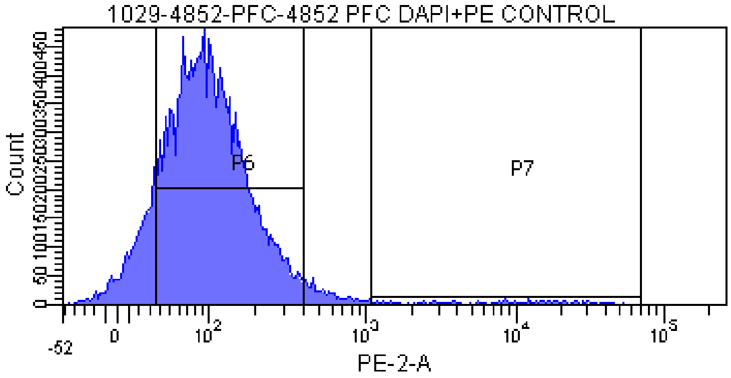


Tube: 4852 PFC DAPI+PE CONTROL

| Population | #Events | %Parent | %Total |
|---|---|---|---|
| All Events | 23,010 | #### | 100.0 |
| P1 | 18,266 | 79.4 | 79.4 |
| P2 | 17,818 | 97.5 | 77.4 |
| P3 | 17,432 | 97.8 | 75.8 |
| P8 | 15,762 | 90.4 | 68.5 |
| P5 | 16,014 | 91.9 | 69.6 |
| P6 | 14,551 | 83.5 | 63.2 |
| P7 | 224 | 1.3 | 1.0 |
| P4 | 184 | 1.1 | 0.8 |

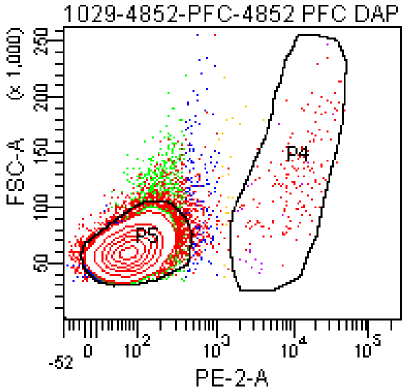

### FACSDiva Version 6.1.3

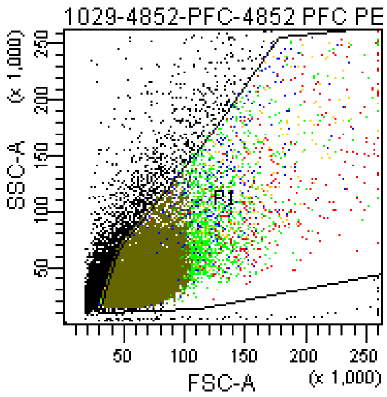


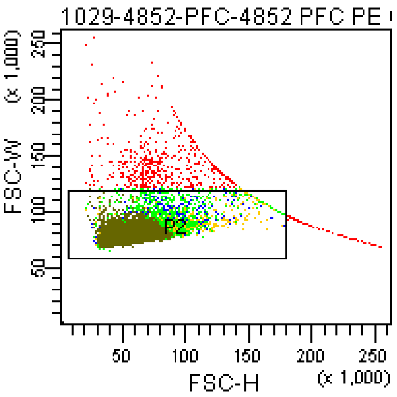


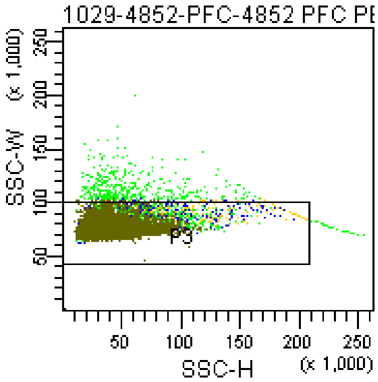


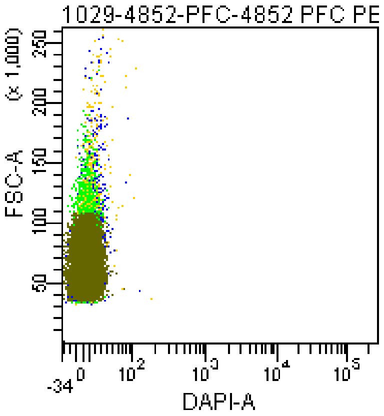


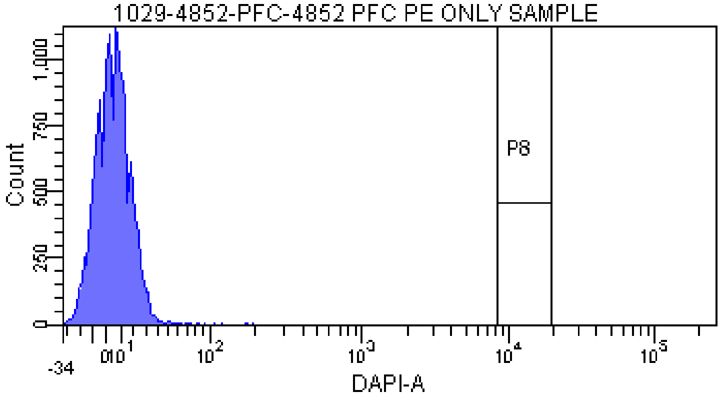


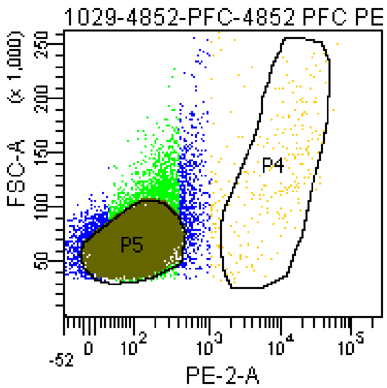


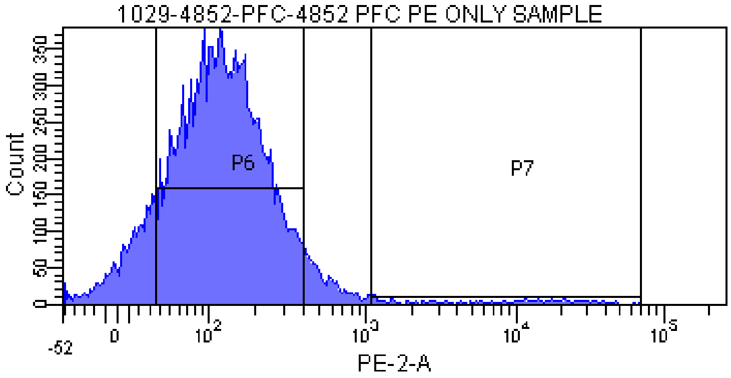


Tube: 4852 PFC PE ONLY SAMPLE

| Population | #Events | %Parent | %Total |
|---|---|---|---|
| All Events | 21,831 | #### | 100.0 |
| P1 | 17,690 | 81.0 | 81.0 |
| P2 | 17,071 | 96.5 | 78.2 |
| P3 | 16,604 | 97.3 | 76.1 |
| P8 | 0 | 0.0 | 0.0 |
| P5 | 14,100 | 84.9 | 64.6 |
| P6 | 13,509 | 81.4 | 61.9 |
| P7 | 287 | 1.7 | 1.3 |
| P4 | 202 | 1.2 | 0.9 |

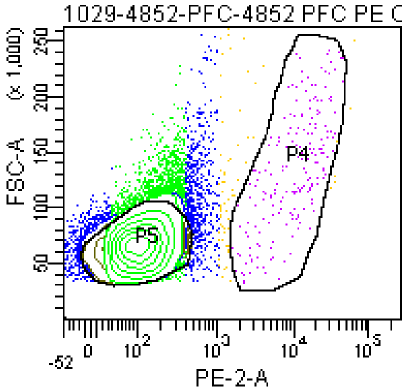

### FACSDiva Version 6.1.3

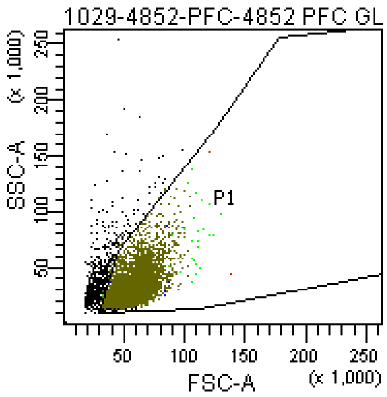


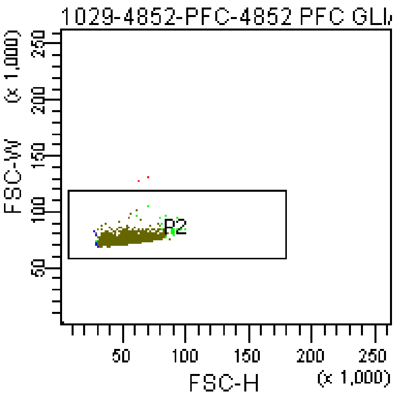


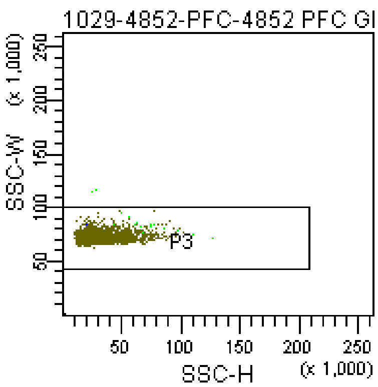


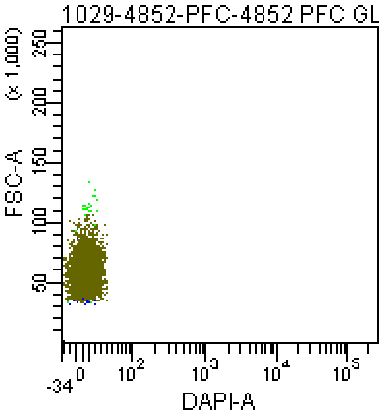


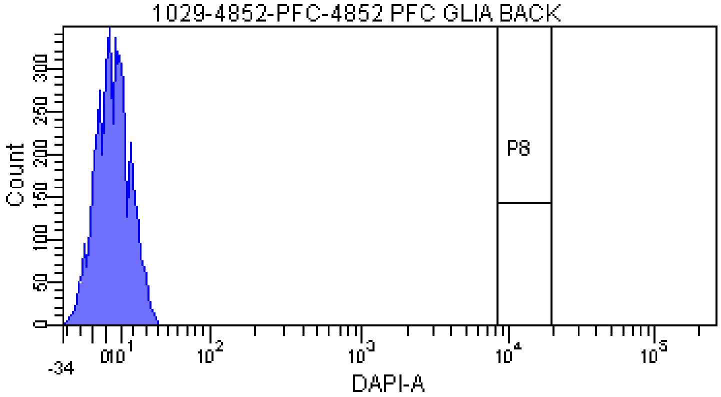


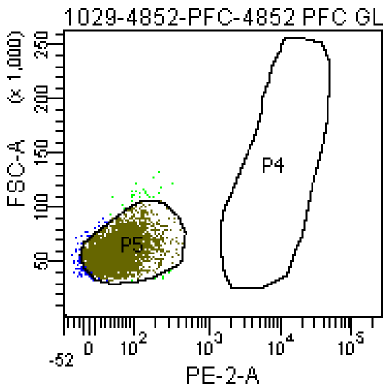


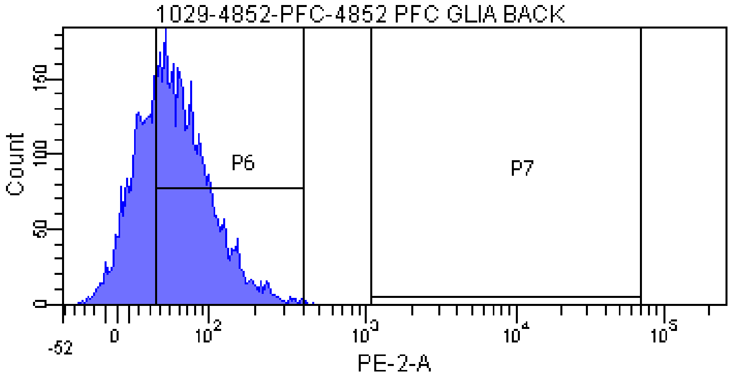


Tube: 4852 PFC GLIA BACK

| Population | #Events | %Parent | %Total |
|---|---|---|---|
| All Events | 5,855 | #### | 100.0 |
| P1 | 5,230 | 89.3 | 89.3 |
| P2 | 5,228 | 100.0 | 89.3 |
| P3 | 5,226 | 100.0 | 89.3 |
| P8 | 0 | 0.0 | 0.0 |
| P5 | 5,075 | 97.1 | 86.7 |
| P6 | 3,349 | 64.1 | 57.2 |
| P7 | 0 | 0.0 | 0.0 |
| P4 | 0 | 0.0 | 0.0 |

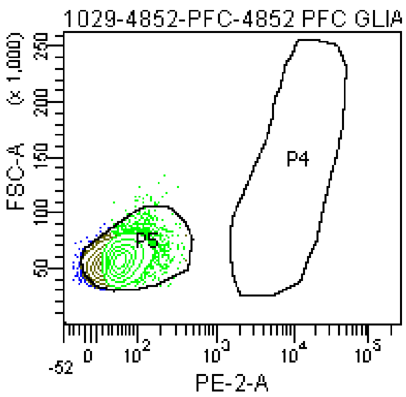

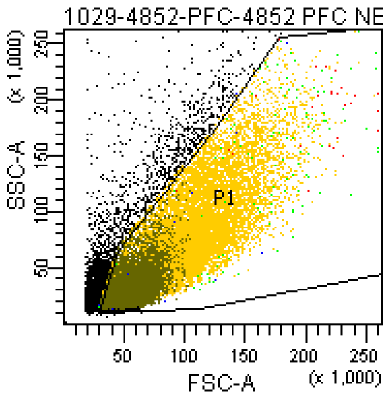


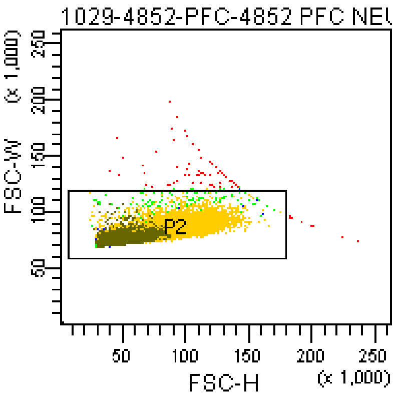


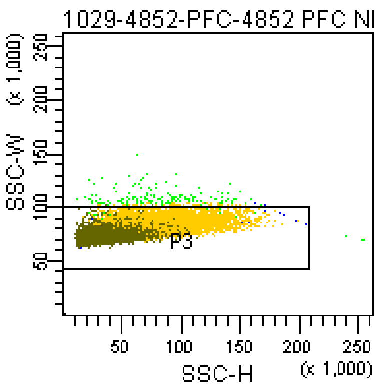


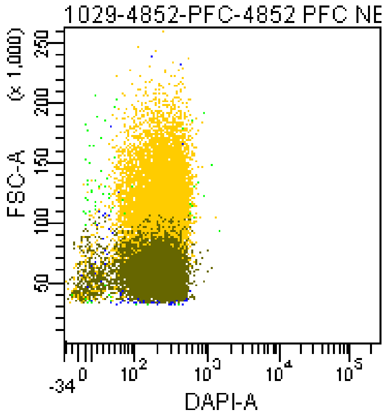


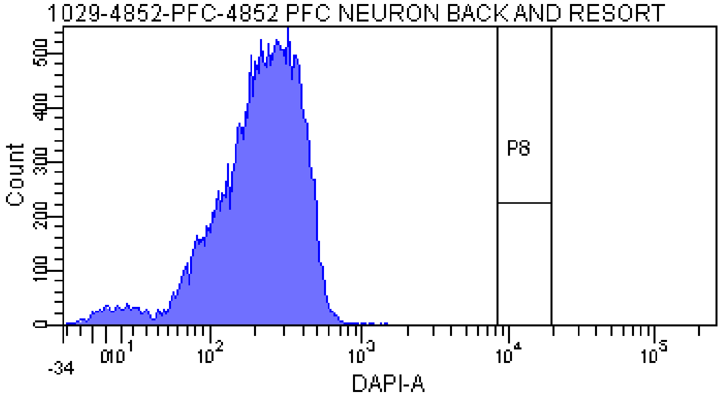


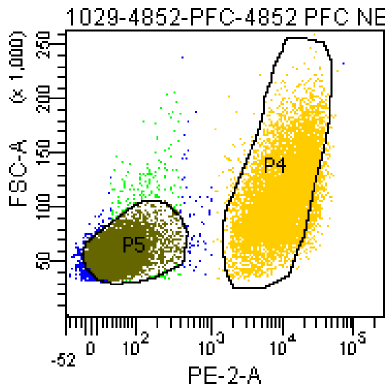


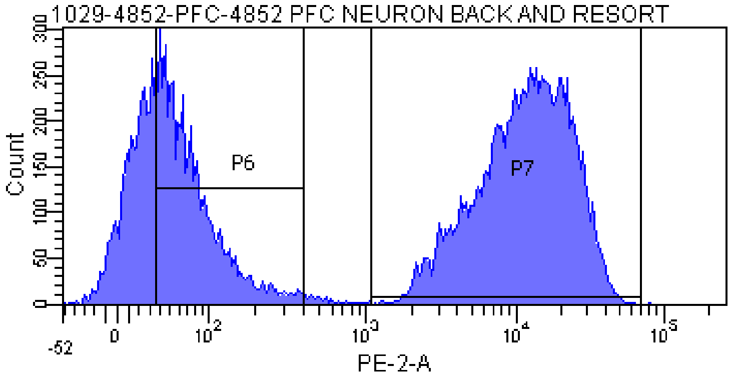


Tube: 4852 PFC NEURON BACK AND RESORT

| Population | #Events | %Parent | %Total |
|---|---|---|---|
| All Events | 22,746 | #### | 100.0 |
| P1 | 19,674 | 86.5 | 86.5 |
| P2 | 19,600 | 99.6 | 86.2 |
| P3 | 19,437 | 99.2 | 85.5 |
| P8 | 0 | 0.0 | 0.0 |
| P5 | 7,655 | 39.4 | 33.7 |
| P6 | 4,773 | 24.6 | 21.0 |
| P7 | 11,160 | 57.4 | 49.1 |
| P4 | 10,484 | 53.9 | 46.1 |

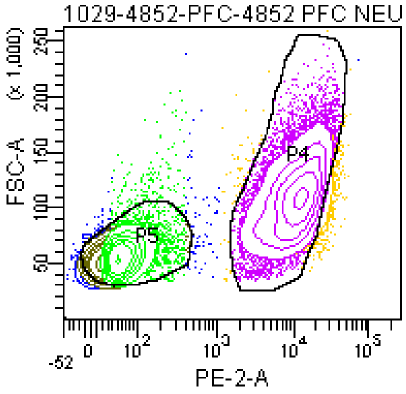

**The BD Arial II FACS sorting reports of UMB#1347**

Sample: prefrontal cortex

Category: healthy control

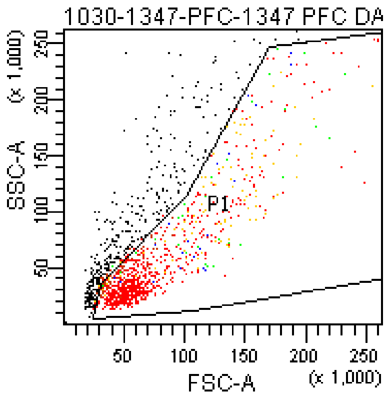


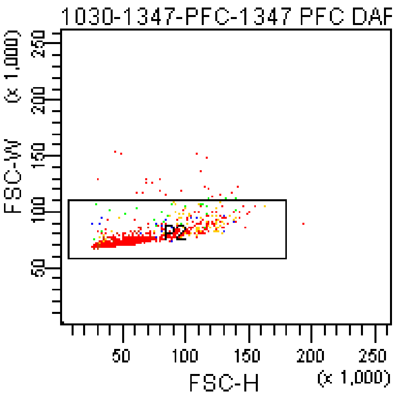


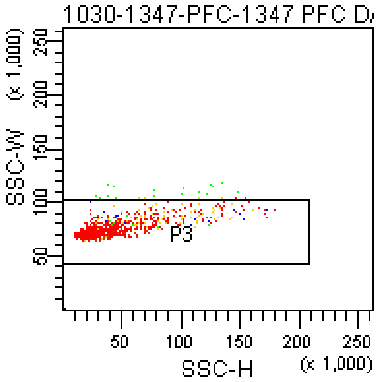


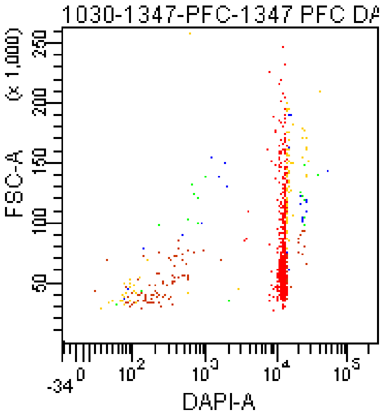


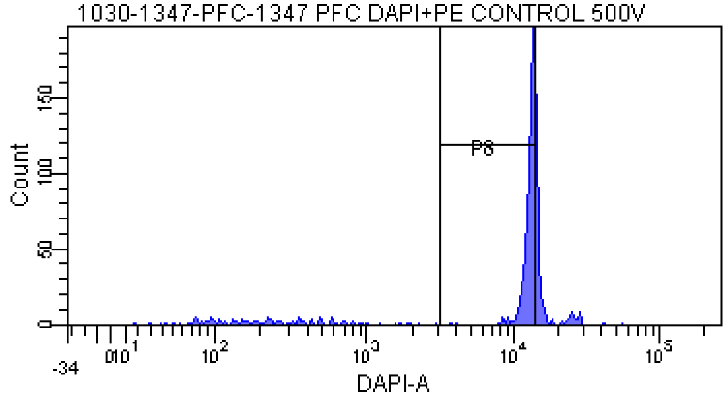


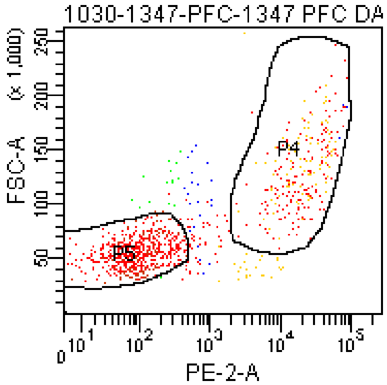


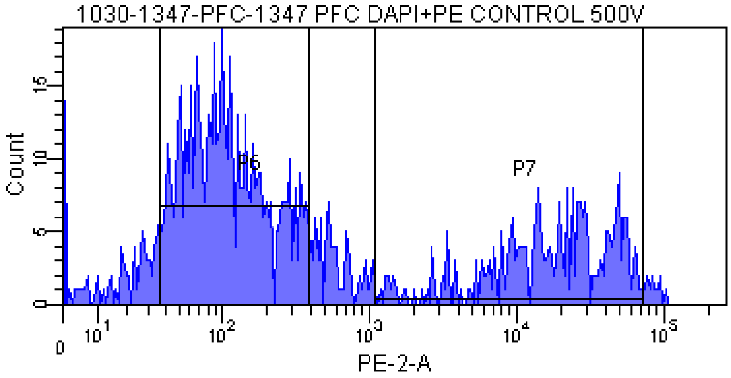


Tube: 1347 PFC DAPI+PE CONTROL 500V

| Population | #Events | %Parent | %Total |
|---|---|---|---|
| All Events | 1,436 | #### | 100.0 |
| P1 | 994 | 69.2 | 69.2 |
| P2 | 972 | 97.8 | 67.7 |
| P3 | 958 | 98.6 | 66.7 |
| P8 | 705 | 73.6 | 49.1 |
| P5 | 593 | 61.9 | 41.3 |
| P6 | 528 | 55.1 | 36.8 |
| P7 | 270 | 28.2 | 18.8 |
| P4 | 237 | 24.7 | 16.5 |

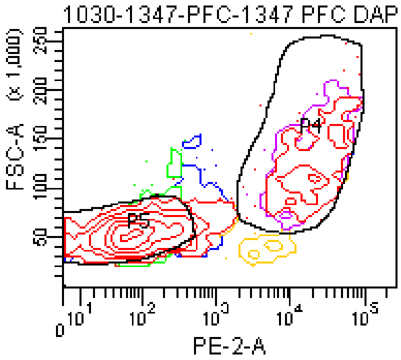

### FACSDiva Version 6.1.3

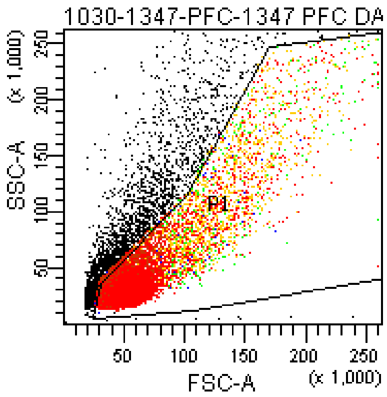


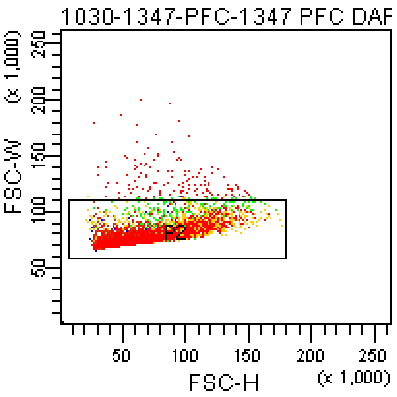


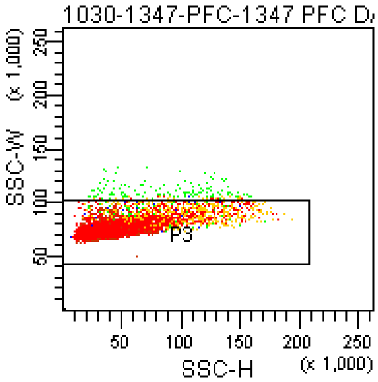


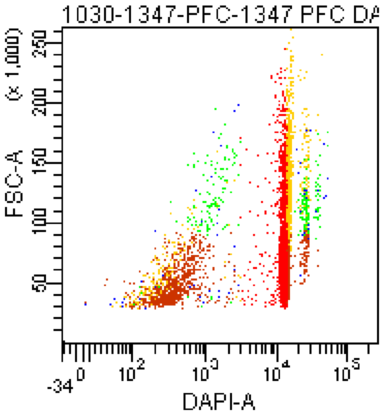


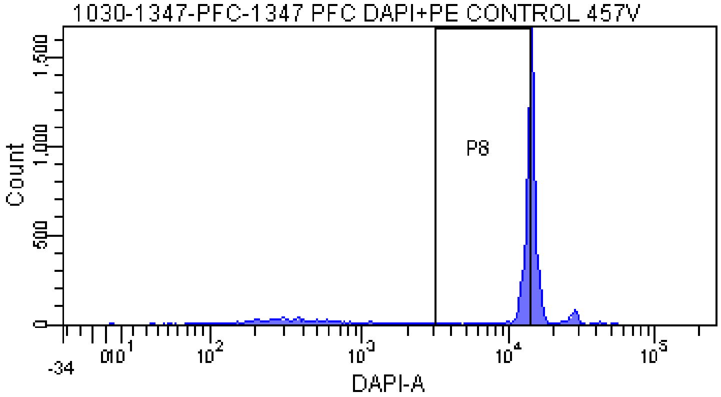


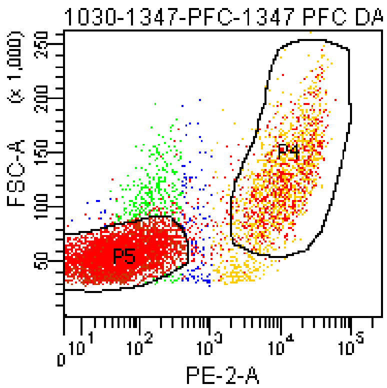


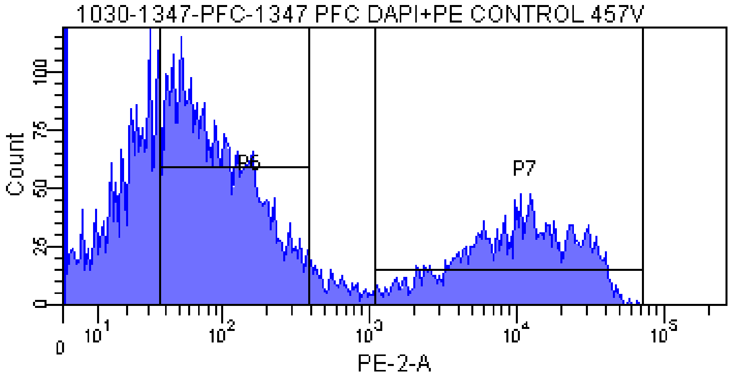


Tube: 1347 PFC DAPI+PE CONTROL 457V

| Population | #Events | %Parent | %Total |
|---|---|---|---|
| All Events | 11,230 | #### | 100.0 |
| P1 | 7,871 | 70.1 | 70.1 |
| P2 | 7,737 | 98.3 | 68.9 |
| P3 | 7,614 | 98.4 | 67.8 |
| P8 | 3,736 | 49.1 | 33.3 |
| P5 | 5,047 | 66.3 | 44.9 |
| P6 | 3,375 | 44.3 | 30.1 |
| P7 | 2,021 | 26.5 | 18.0 |
| P4 | 1,766 | 23.2 | 15.7 |

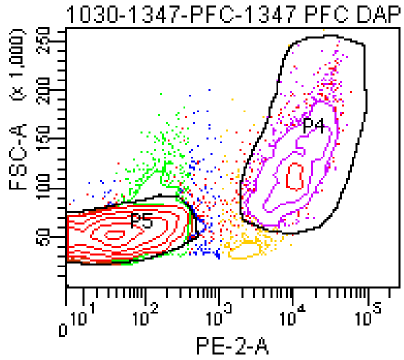

### FACSDiva Version 6.1.3

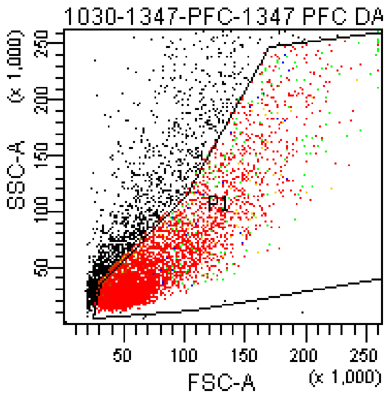


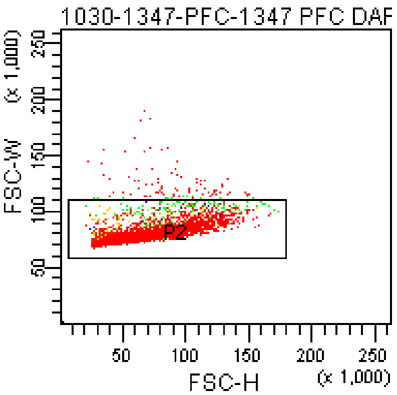


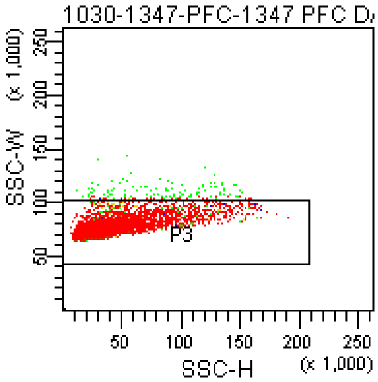


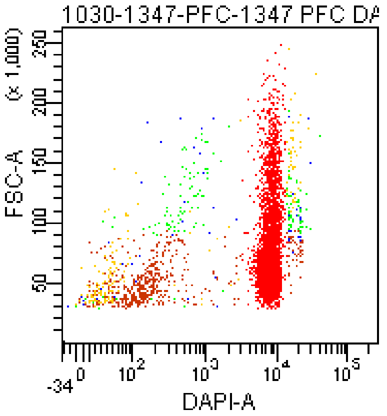


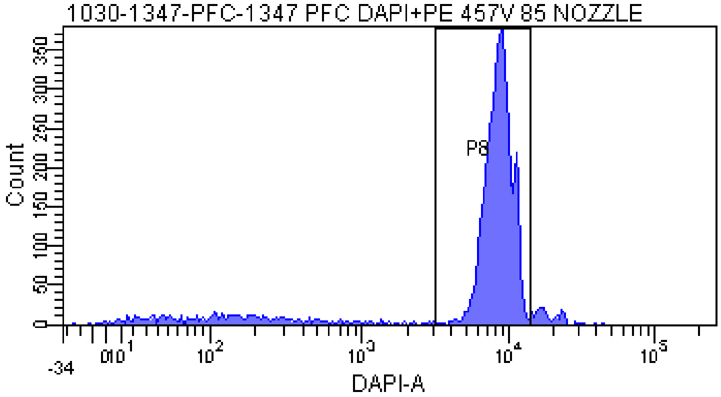


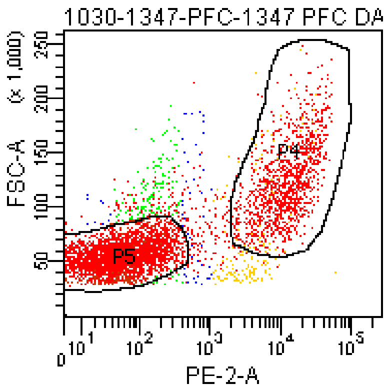


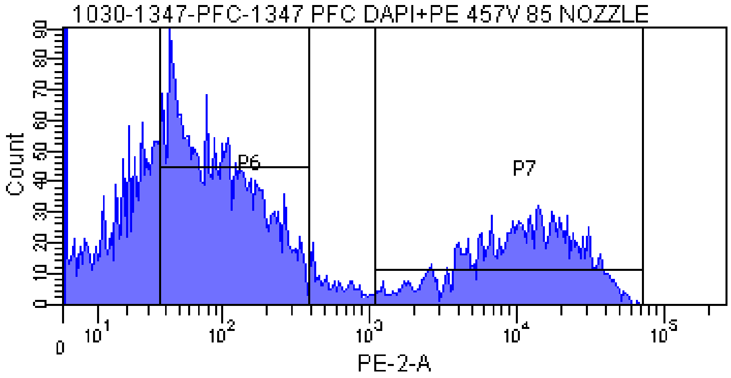


Tube: 1347 PFC DAPI+PE 457V 85 NOZZLE

| Population | #Events | %Parent | %Total |
|---|---|---|---|
| All Events | 7,432 | #### | 100.0 |
| P1 | 5,430 | 73.1 | 73.1 |
| P2 | 5,355 | 98.6 | 72.1 |
| P3 | 5,258 | 98.2 | 70.7 |
| P8 | 4,397 | 83.6 | 59.2 |
| P5 | 3,454 | 65.7 | 46.5 |
| P6 | 2,326 | 44.2 | 31.3 |
| P7 | 1,429 | 27.2 | 19.2 |
| P4 | 1,266 | 24.1 | 17.0 |

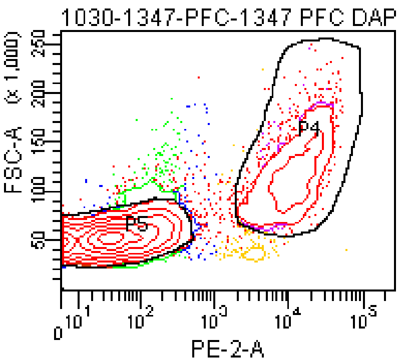

### FACSDiva Version 6.1.3

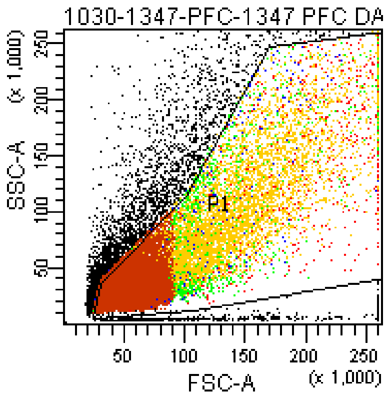


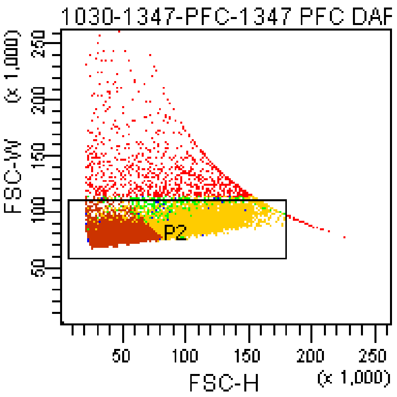


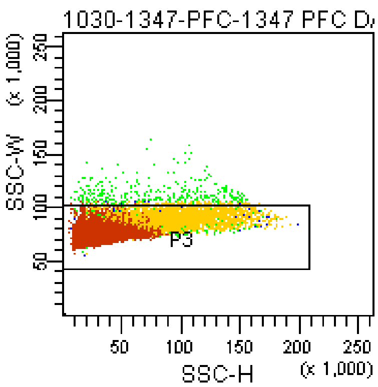


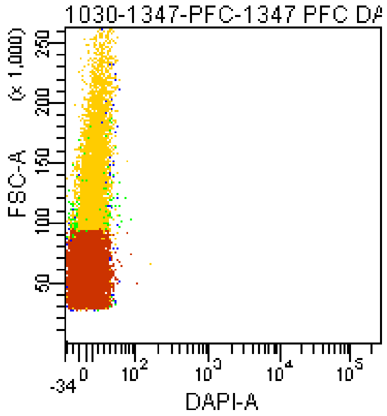


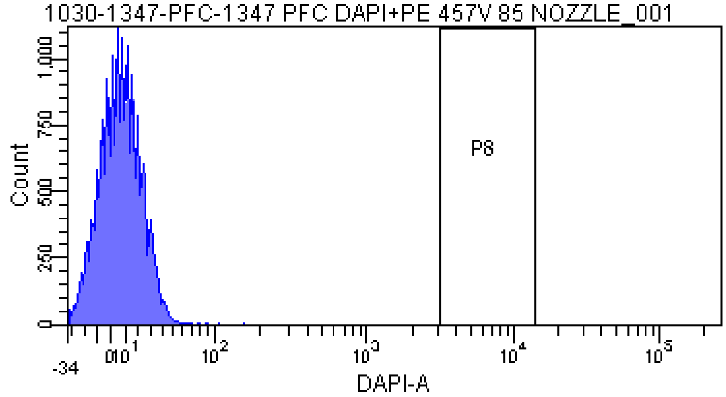


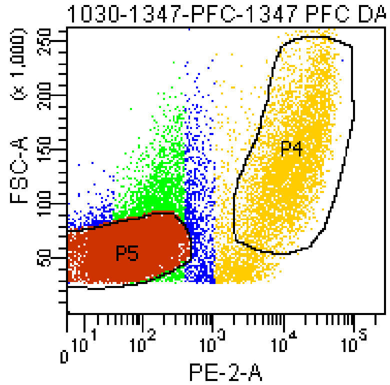


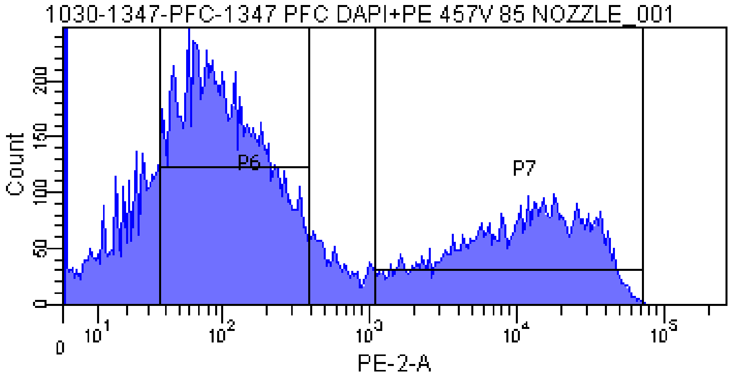


Tube: 1347 PFC DAPI+PE 457V 85 NOZZLE_001

| Population | #Events | %Parent | %Total |
|---|---|---|---|
| All Events | 26,621 | #### | 100.0 |
| P1 | 20,000 | 75.1 | 75.1 |
| P2 | 18,950 | 94.8 | 71.2 |
| P3 | 18,622 | 98.3 | 70.0 |
| P8 | 0 | 0.0 | 0.0 |
| P5 | 10,486 | 56.3 | 39.4 |
| P6 | 8,958 | 48.1 | 33.7 |
| P7 | 5,507 | 29.6 | 20.7 |
| P4 | 4,360 | 23.4 | 16.4 |

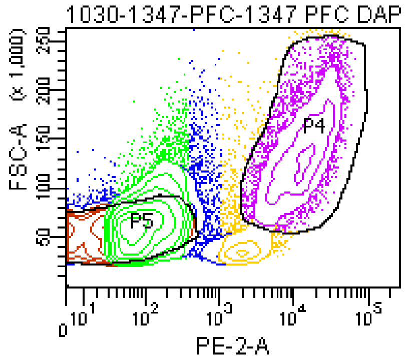

### FACSDiva Version 6.1.3

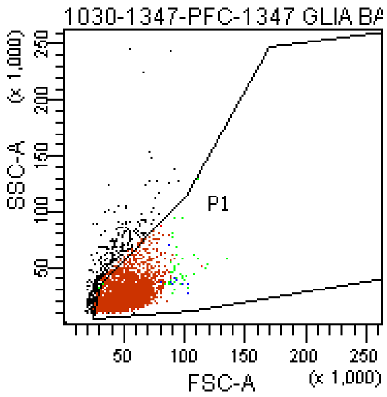


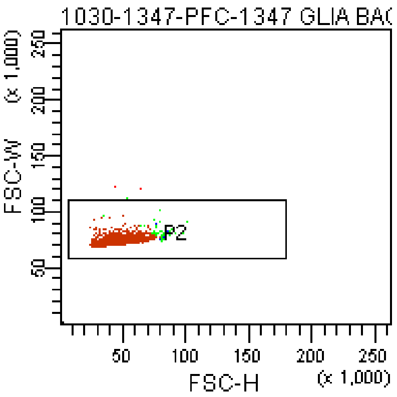


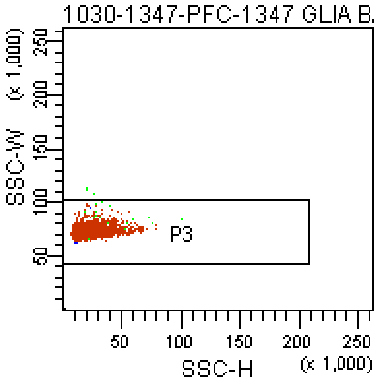


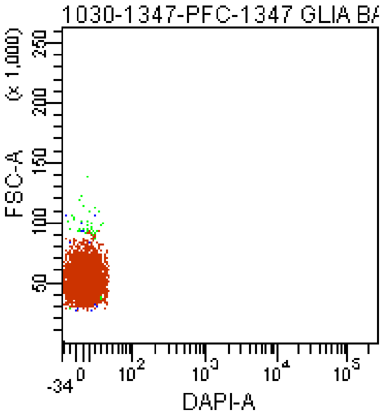


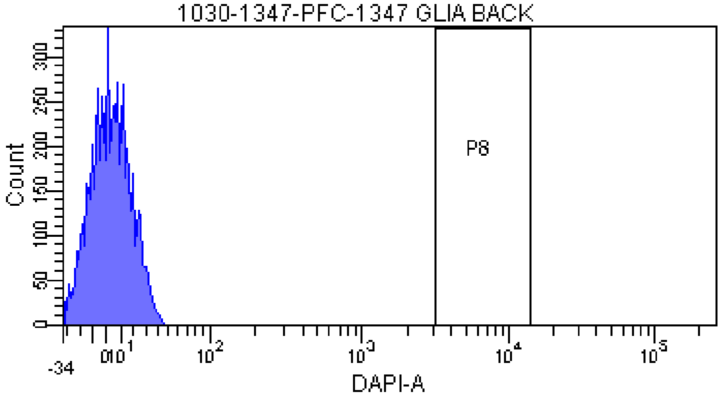


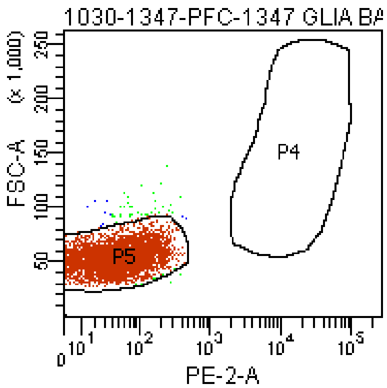


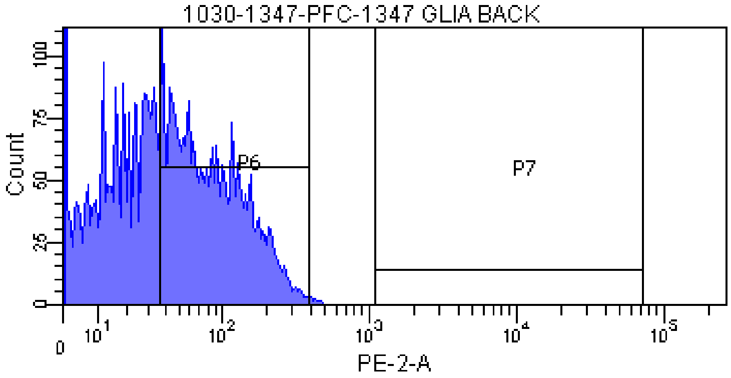


Tube: 1347 GLIA BACK

| Population | #Events | %Parent | %Total |
|---|---|---|---|
| All Events | 5,393 | #### | 100.0 |
| P1 | 5,043 | 93.5 | 93.5 |
| P2 | 5,041 | 100.0 | 93.5 |
| P3 | 5,038 | 99.9 | 93.4 |
| P8 | 0 | 0.0 | 0.0 |
| P5 | 4,962 | 98.5 | 92.0 |
| P6 | 2,408 | 47.8 | 44.7 |
| P7 | 0 | 0.0 | 0.0 |
| P4 | 0 | 0.0 | 0.0 |

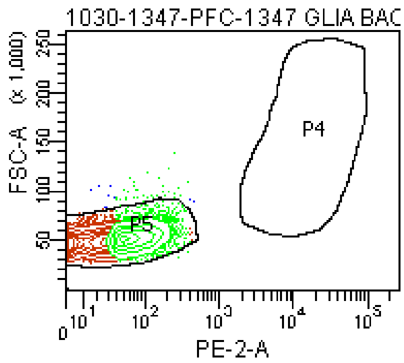

### FACSDiva Version 6.1.3

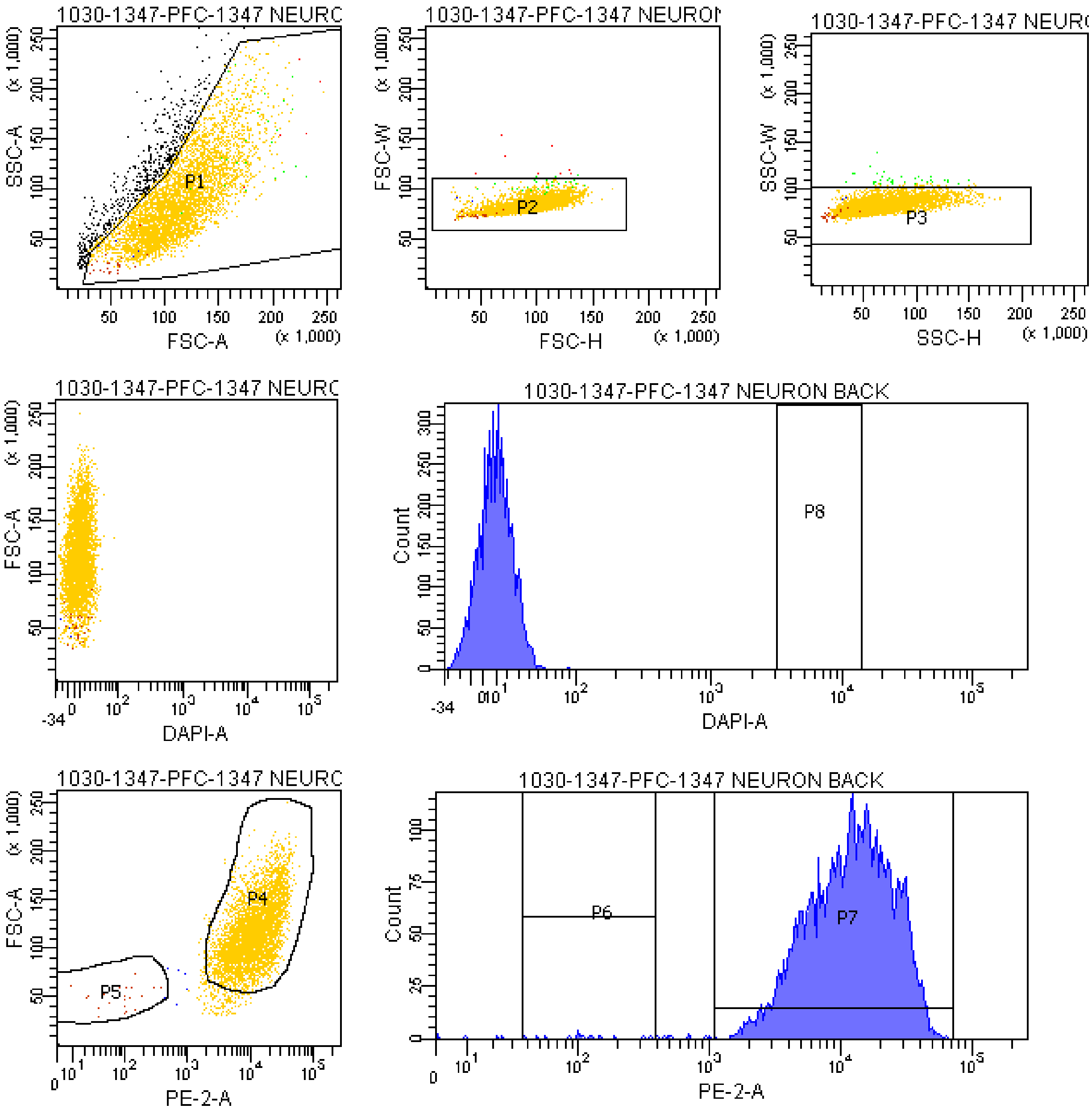


Tube: 1347 NEURON BACK

| Population | #Events | %Parent | %Total |
|---|---|---|---|
| All Events | 5,318 | #### | 100.0 |
| P1 | 4,752 | 89.4 | 89.4 |
| P2 | 4,744 | 99.8 | 89.2 |
| P3 | 4,707 | 99.2 | 88.5 |
| P8 | 0 | 0.0 | 0.0 |
| P5 | 25 | 0.5 | 0.5 |
| P6 | 18 | 0.4 | 0.3 |
| P7 | 4,676 | 99.3 | 87.9 |
| P4 | 4,499 | 95.6 | 84.6 |

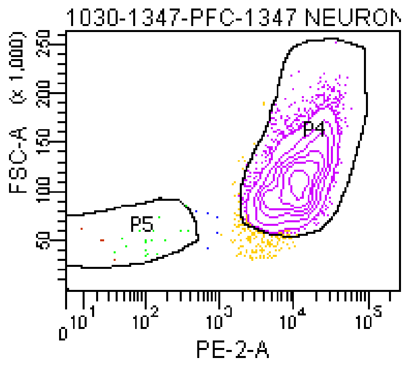

**The BD Arial II FACS sorting reports of UMB#4882**

Sample: prefrontal cortex

Category: Rett patient

### FACSDiva Version 6.1.3

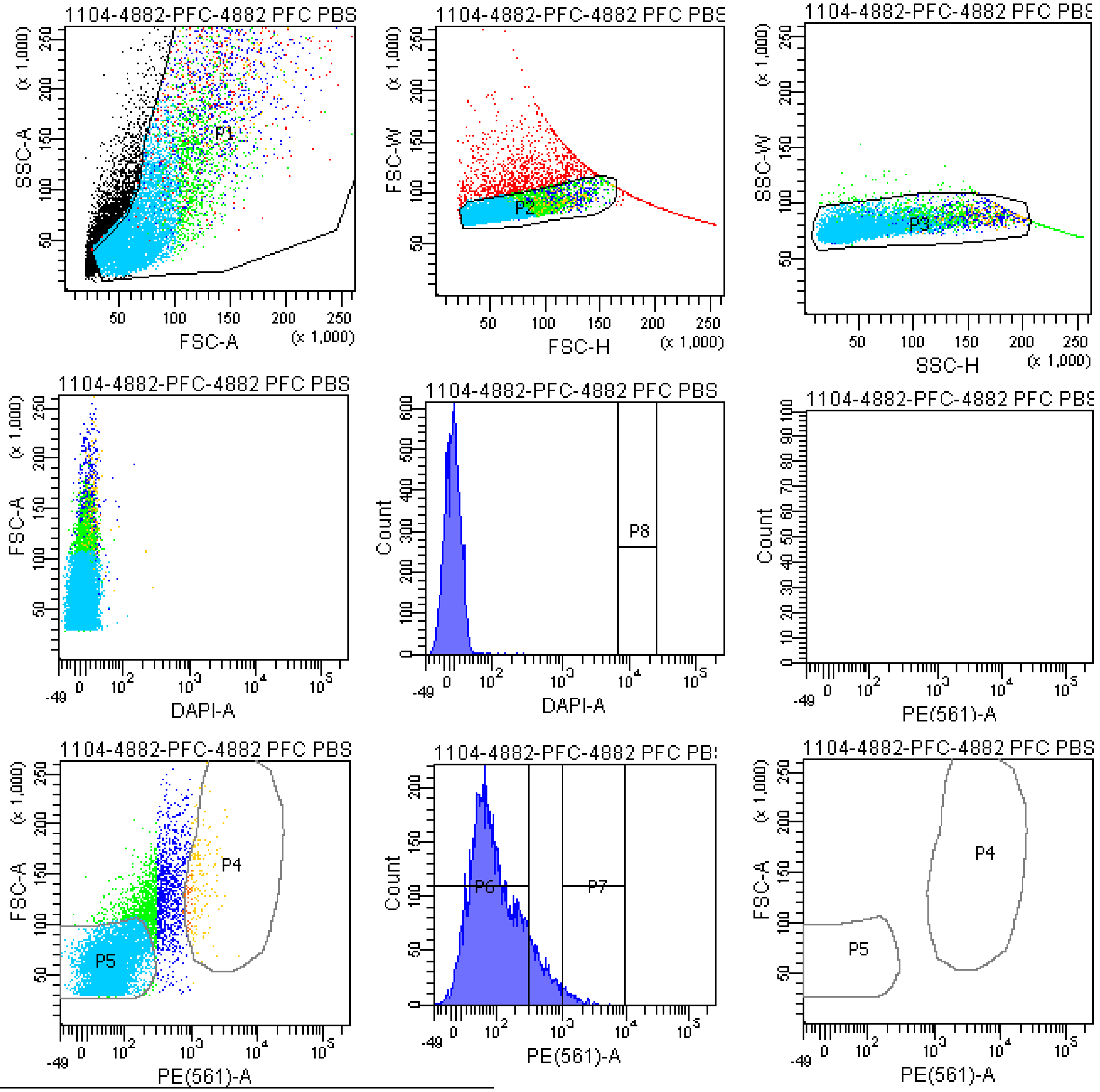


Tube: 4882 PFC PBS CONTROL

| Population | #Events | %Parent | %Total |
|---|---|---|---|
| All Events | 15,116 | #### | 100.0 |
| P1 | 11,238 | 74.3 | 74.3 |
| P2 | 9,605 | 85.5 | 63.5 |
| P3 | 9,193 | 95.7 | 60.8 |
| P4 | 207 | 2.3 | 1.4 |
| P5 | 6,914 | 75.2 | 45.7 |
| P6 | 7,986 | 86.9 | 52.8 |
| P7 | 162 | 1.8 | 1.1 |
| P8 | 0 | 0.0 | 0.0 |
| P9 | 1,519 | 16.5 | 10.0 |

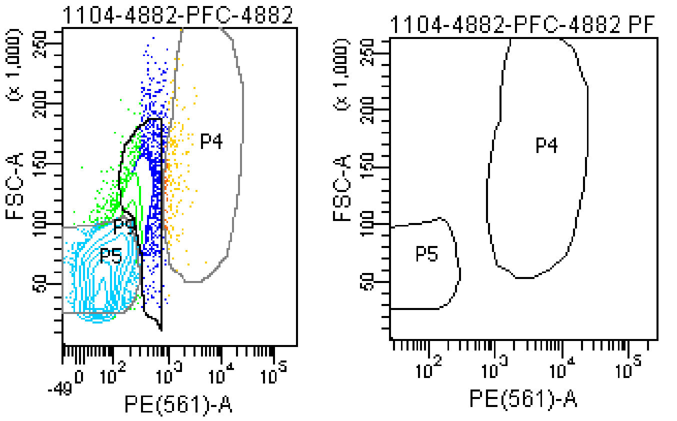

### FACSDiva Version 6.1.3

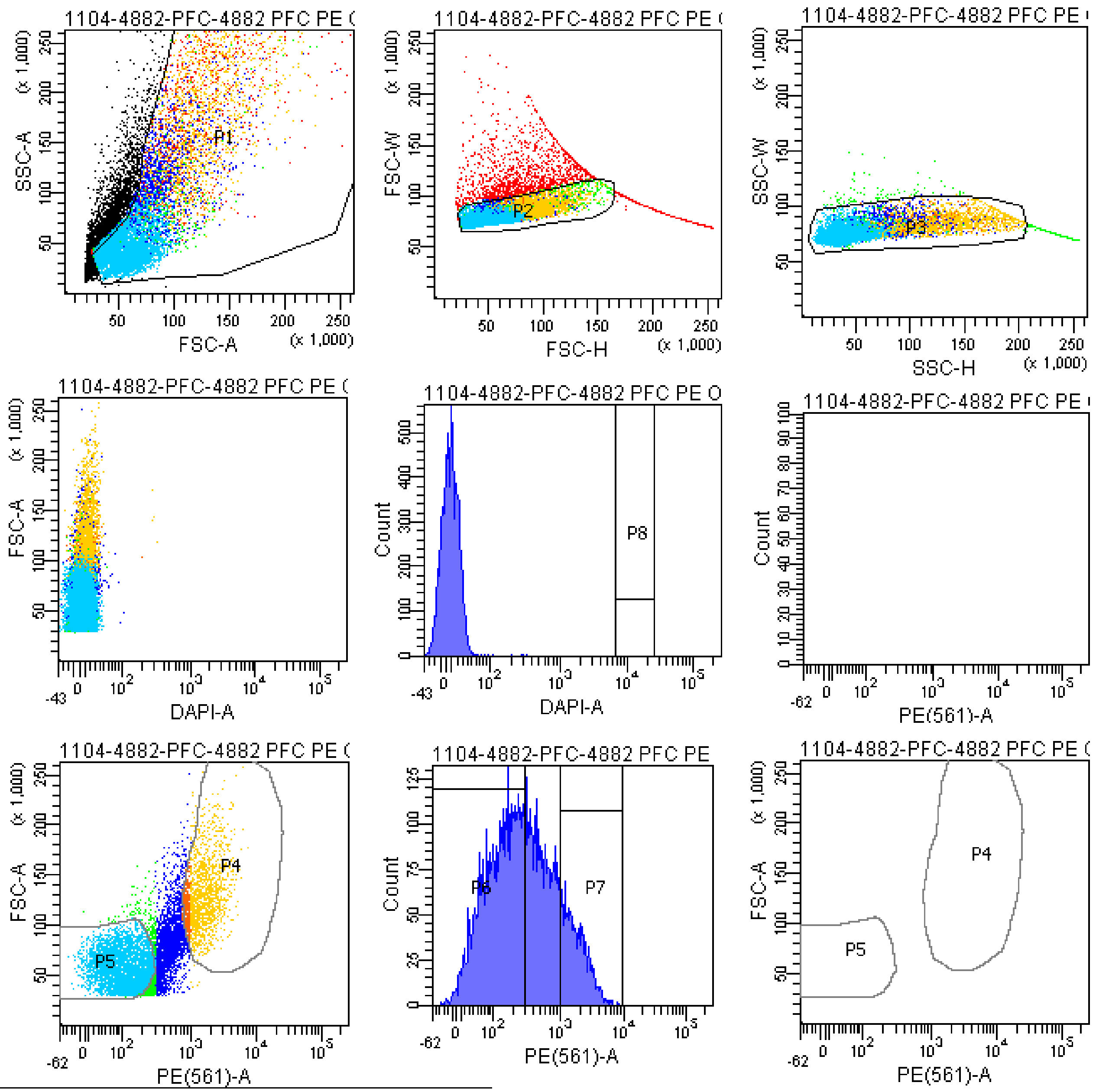


| | Tube: 4882 PFC PE ONLY | | |
|---|---|---|---|
| Population | #Events | %Parent | %Total |
| All Events | 15,420 | #### | 100.0 |
| P1 | 11,140 | 72.2 | 72.2 |
| P2 | 9,110 | 81.8 | 59.1 |
| P3 | 8,549 | 93.8 | 55.4 |
| P4 | 1,599 | 18.7 | 10.4 |
| P5 | 4,048 | 47.4 | 26.3 |
| P6 | 4,744 | 55.5 | 30.8 |
| P7 | 1,279 | 15.0 | 8.3 |
| P8 | 0 | 0.0 | 0.0 |
| P9 | 1,876 | 21.9 | 12.2 |

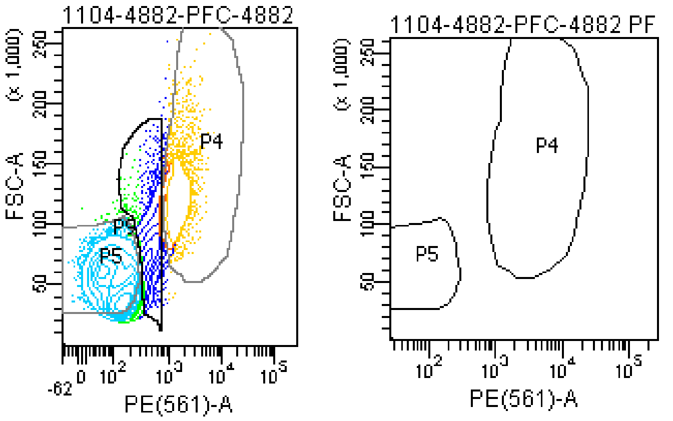

### FACSDiva Version 6.1.3

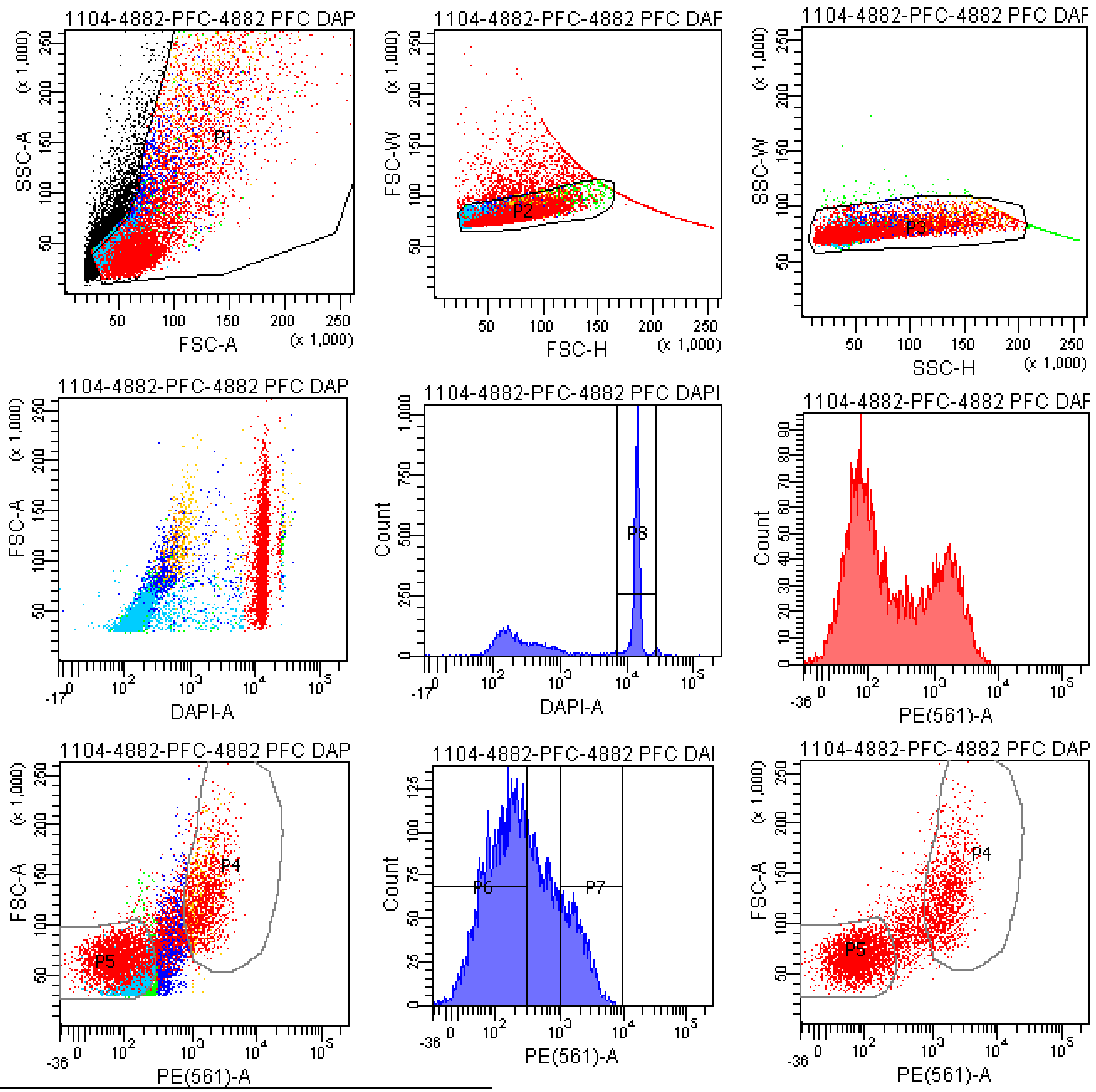


Tube: 4882 PFC DAPI+PE

| Population | #Events | %Parent | %Total |
|---|---|---|---|
| All Events | 15,537 | #### | 100.0 |
| P1 | 11,189 | 72.0 | 72.0 |
| P2 | 9,518 | 85.1 | 61.3 |
| P3 | 9,011 | 94.7 | 58.0 |
| P4 | 1,724 | 19.1 | 11.1 |
| P5 | 4,533 | 50.3 | 29.2 |
| P6 | 5,213 | 57.9 | 33.6 |
| P7 | 1,464 | 16.2 | 9.4 |
| P8 | 4,714 | 52.3 | 30.3 |
| P9 | 1,758 | 19.5 | 11.3 |

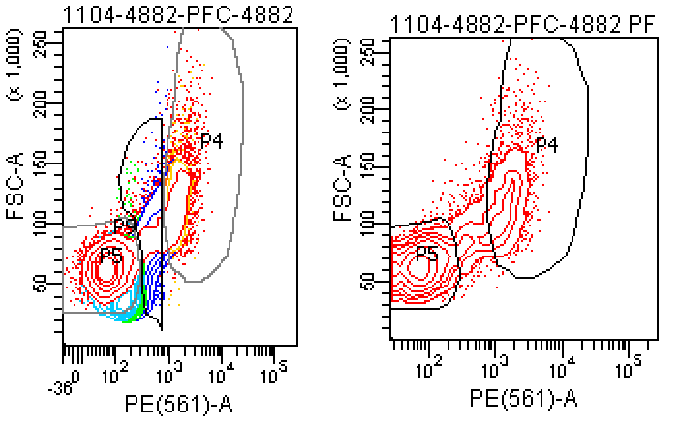

### FACSDiva Version 6.1.3

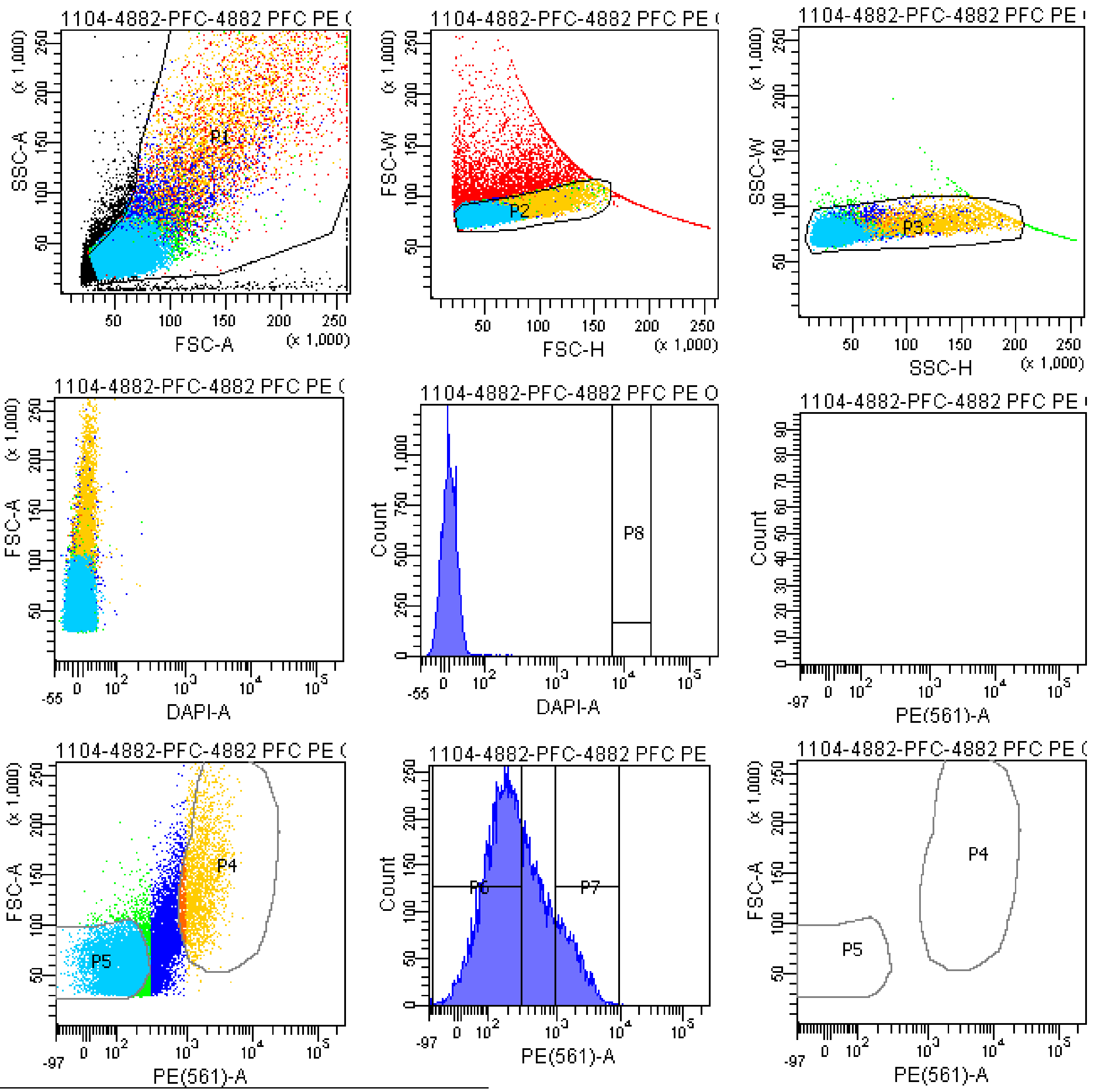


Tube: 4882 PFC PE ONLY SAMPLE

| Population | #Events | %Parent | %Total |
|---|---|---|---|
| All Events | 26,520 | #### | 100.0 |
| P1 | 21,160 | 79.8 | 79.8 |
| P2 | 15,967 | 75.5 | 60.2 |
| P3 | 15,537 | 97.3 | 58.6 |
| P4 | 2,428 | 15.6 | 9.2 |
| P5 | 8,047 | 51.8 | 30.3 |
| P6 | 9,170 | 59.0 | 34.6 |
| P7 | 2,111 | 13.6 | 8.0 |
| P8 | 0 | 0.0 | 0.0 |
| P9 | 3,369 | 21.7 | 12.7 |

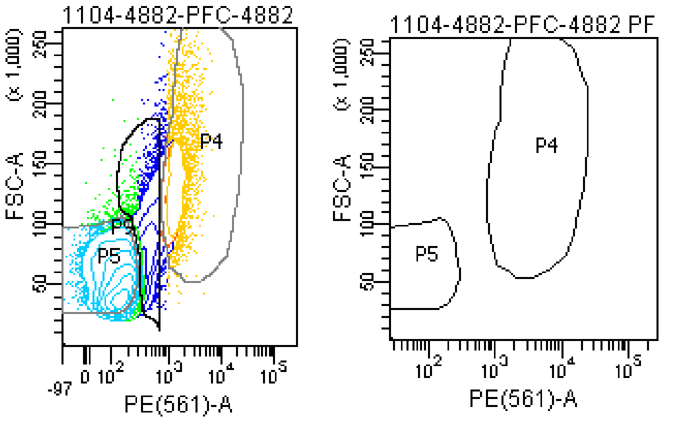

### FACSDiva Version 6.1.3

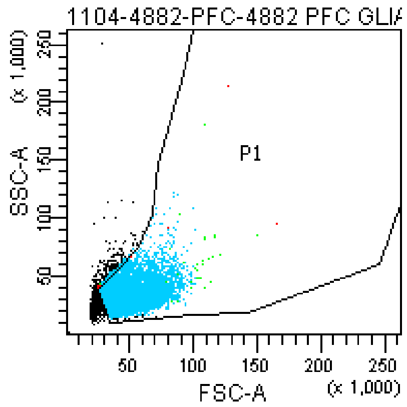


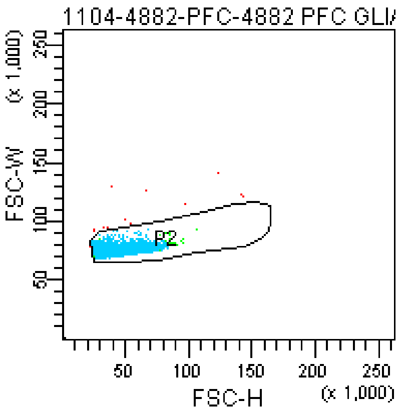


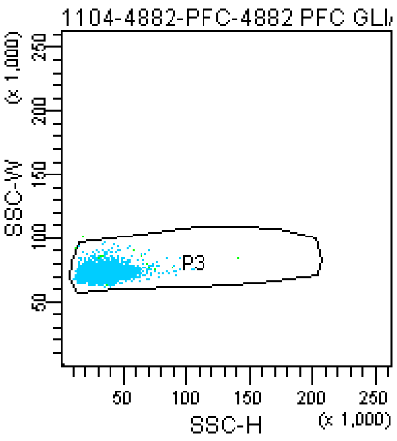


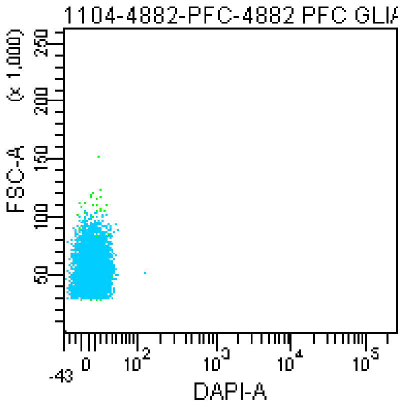


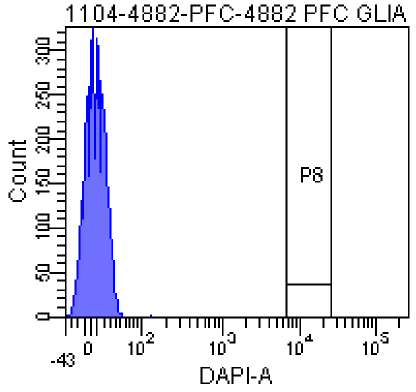


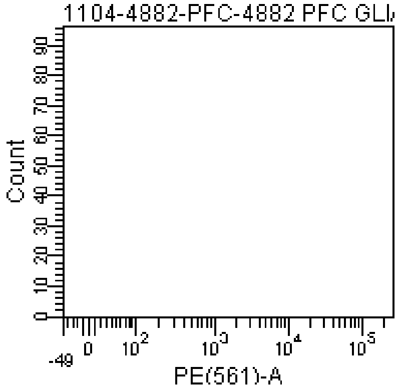


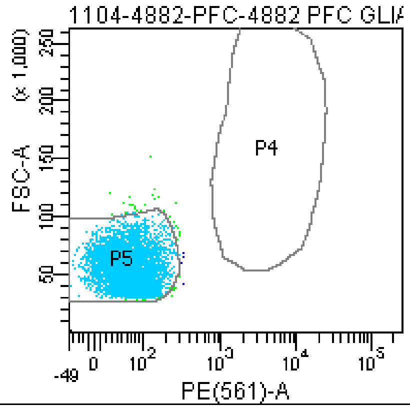


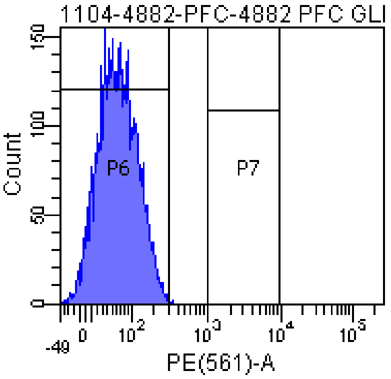


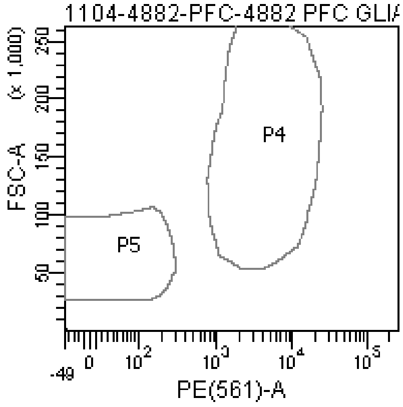


Tube: 4882 PFC GLIA BACK

| Population | #Events | %Parent | %Total |
|---|---|---|---|
| All Events | 6,855 | #### | 100.0 |
| P1 | 5,429 | 79.2 | 79.2 |
| P2 | 5,391 | 99.3 | 78.6 |
| P3 | 5,389 | 100.0 | 78.6 |
| P4 | 0 | 0.0 | 0.0 |
| P5 | 5,325 | 98.8 | 77.7 |
| P6 | 5,384 | 99.9 | 78.5 |
| P7 | 0 | 0.0 | 0.0 |
| P8 | 0 | 0.0 | 0.0 |
| P9 | 10 | 0.2 | 0.1 |

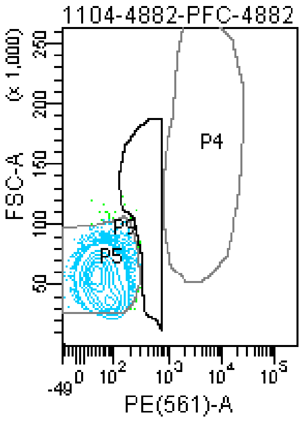


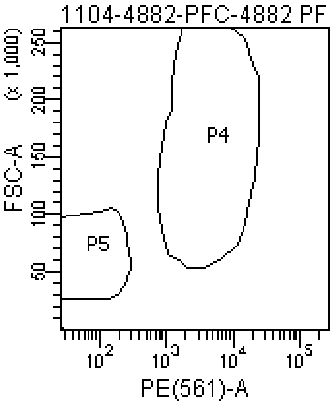

### FACSDiva Version 6.1.3

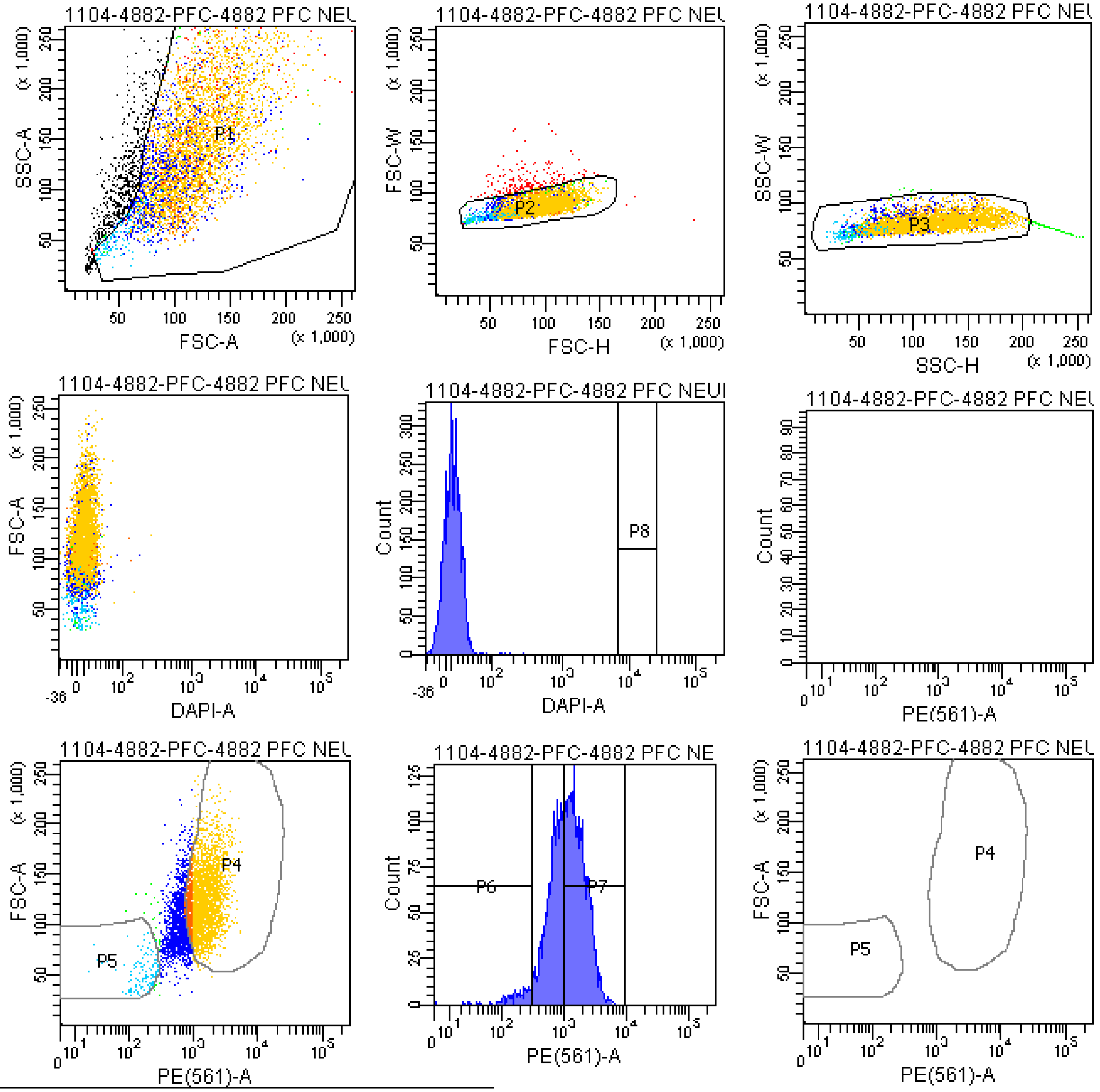


Tube: 4882 PFC NEURON BACK

| Population | #Events | %Parent | %Total |
|---|---|---|---|
| All Events | 5,432 | #### | 100.0 |
| P1 | 4,845 | 89.2 | 89.2 |
| P2 | 4,652 | 96.0 | 85.6 |
| P3 | 4,551 | 97.8 | 83.8 |
| P4 | 3,016 | 66.3 | 55.5 |
| P5 | 124 | 2.7 | 2.3 |
| P6 | 158 | 3.5 | 2.9 |
| P7 | 2,586 | 56.8 | 47.6 |
| P8 | 0 | 0.0 | 0.0 |
| P9 | 965 | 21.2 | 17.8 |

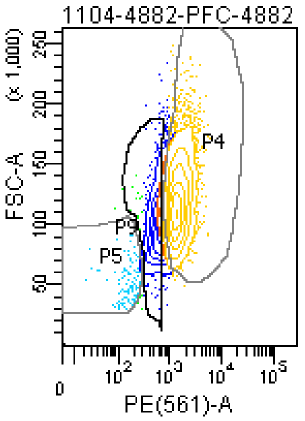


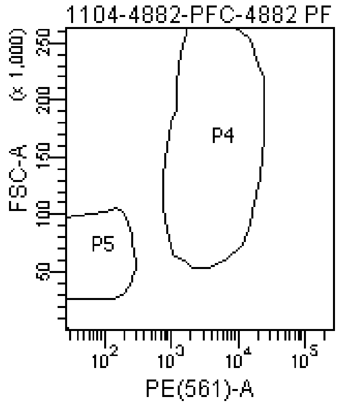

### FACSDiva Version 6.1.3

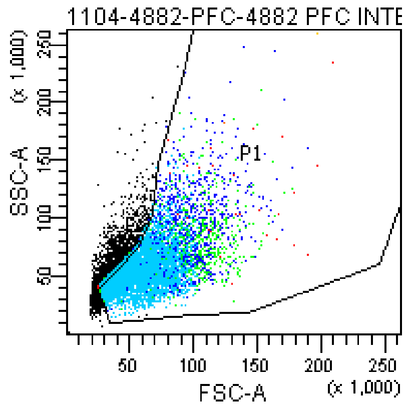


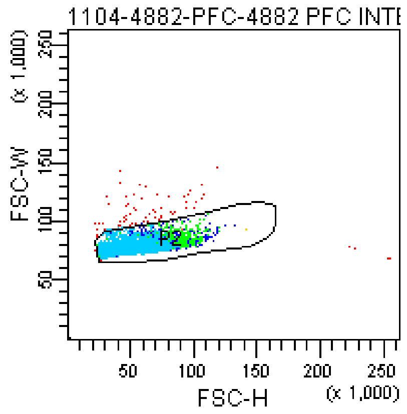


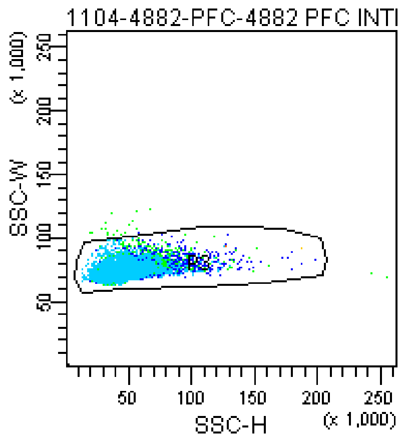


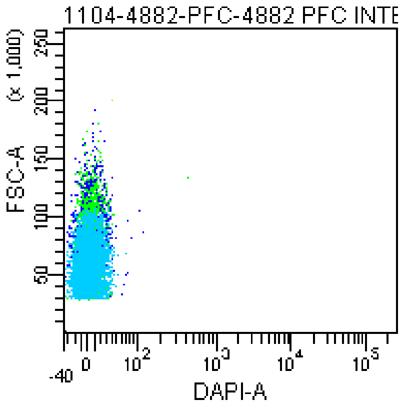


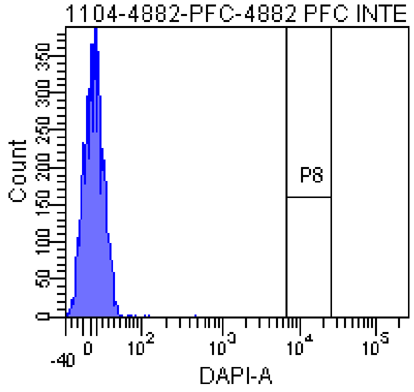


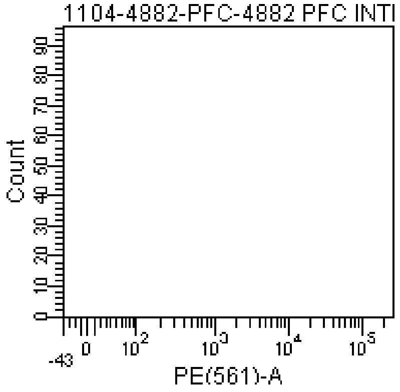


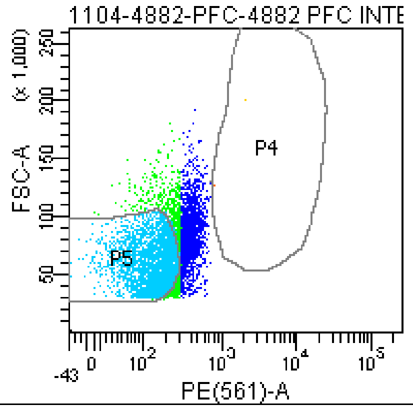


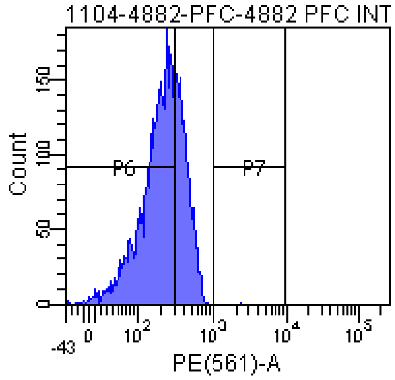


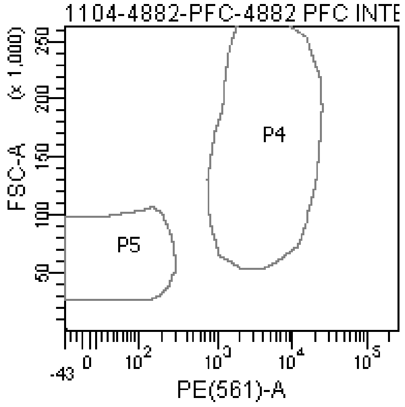


Tube: 4882 PFC INTER BACK

| Population | #Events | %Parent | %Total |
|---|---|---|---|
| All Events | 7,851 | #### | 100.0 |
| P1 | 6,105 | 77.8 | 77.8 |
| P2 | 6,009 | 98.4 | 76.5 |
| P3 | 5,990 | 99.7 | 76.3 |
| P4 | 2 | 0.0 | 0.0 |
| P5 | 3,438 | 57.4 | 43.8 |
| P6 | 4,214 | 70.4 | 53.7 |
| P7 | 1 | 0.0 | 0.0 |
| P8 | 0 | 0.0 | 0.0 |
| P9 | 1,939 | 32.4 | 24.7 |

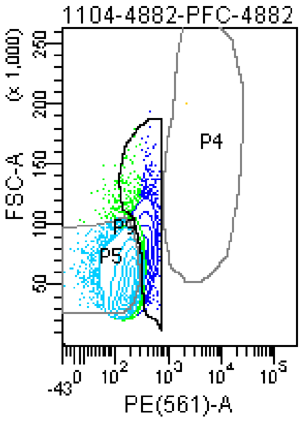


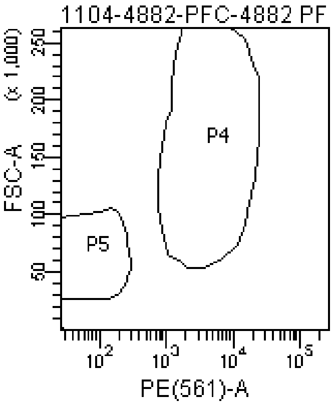

### FACSDiva Version 6.1.3

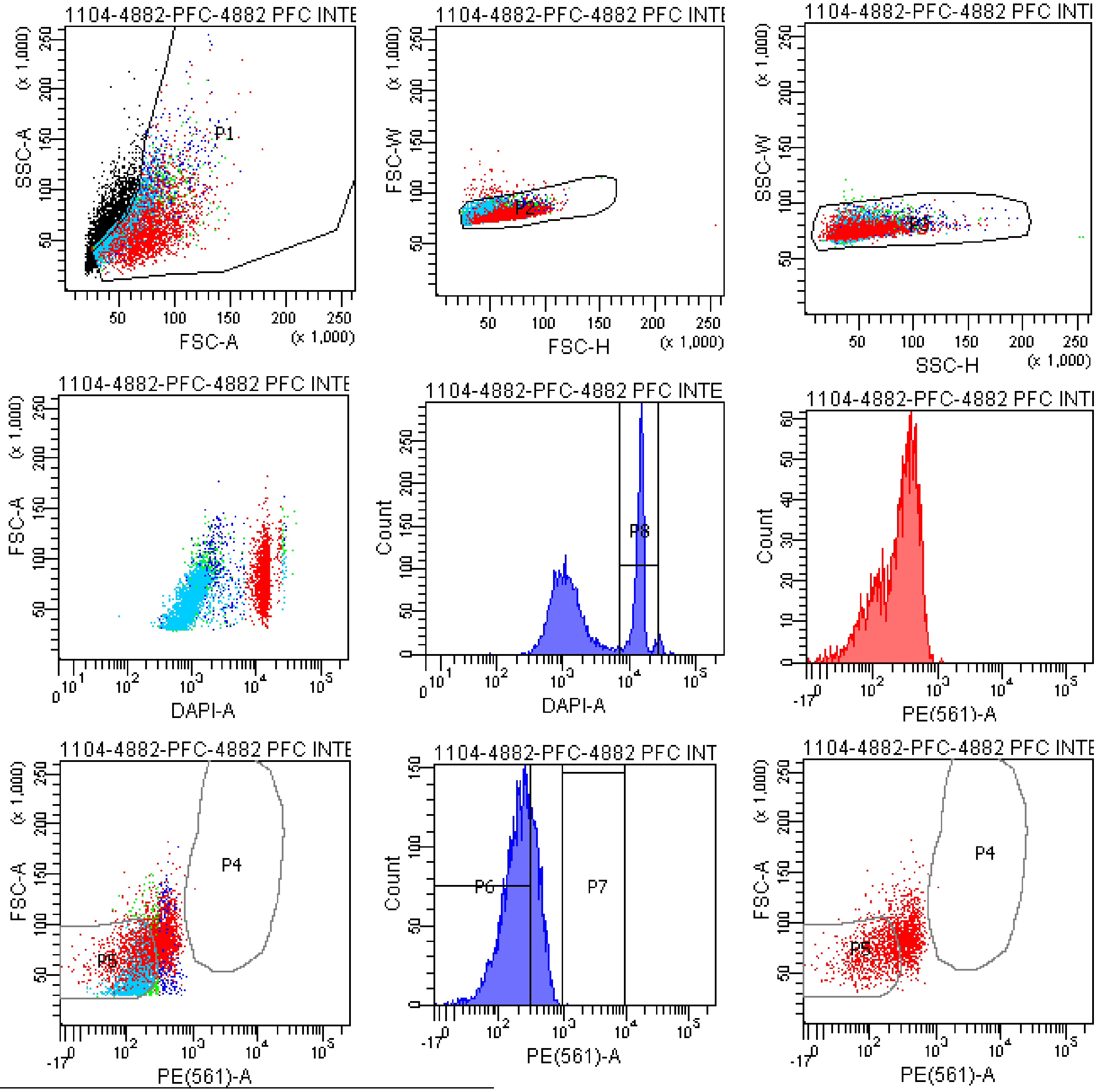


Tube: 4882 PFC INTER BACK+DAPI

| Population | #Events | %Parent | %Total |
|---|---|---|---|
| All Events | 7,269 | #### | 100.0 |
| P1 | 5,122 | 70.5 | 70.5 |
| P2 | 5,017 | 98.0 | 69.0 |
| P3 | 5,002 | 99.7 | 68.8 |
| P4 | 2 | 0.0 | 0.0 |
| P5 | 3,056 | 61.1 | 42.0 |
| P6 | 3,625 | 72.5 | 49.9 |
| P7 | 1 | 0.0 | 0.0 |
| P8 | 1,882 | 37.6 | 25.9 |
| P9 | 1,553 | 31.0 | 21.4 |

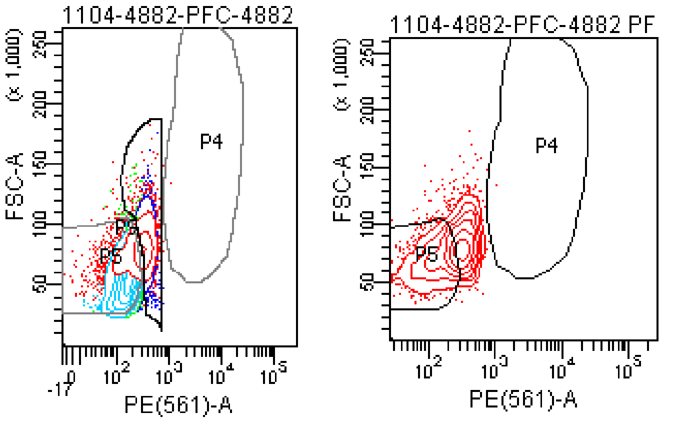

### FACSDiva Version 6.1.3

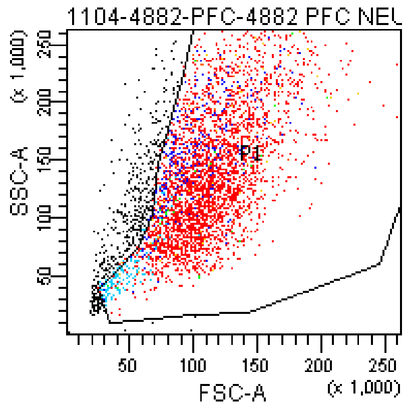


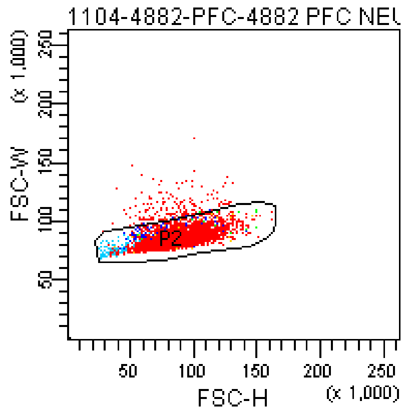


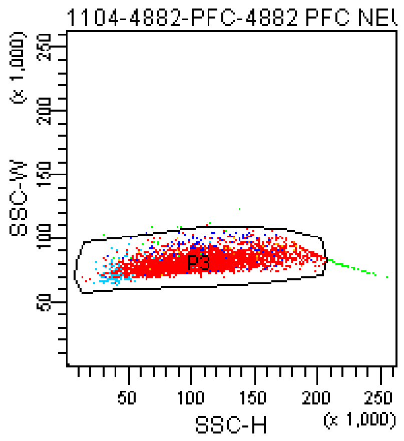


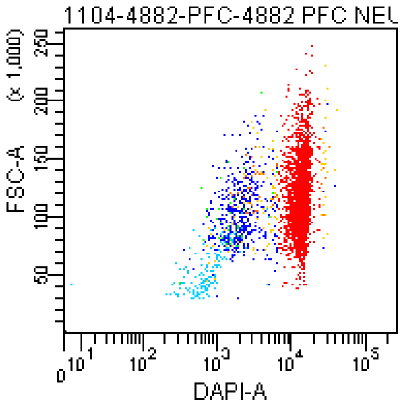


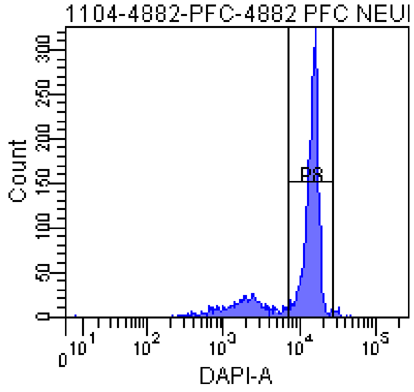


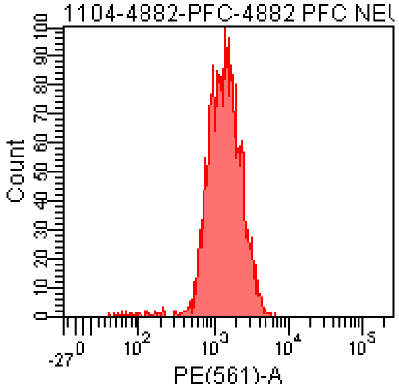


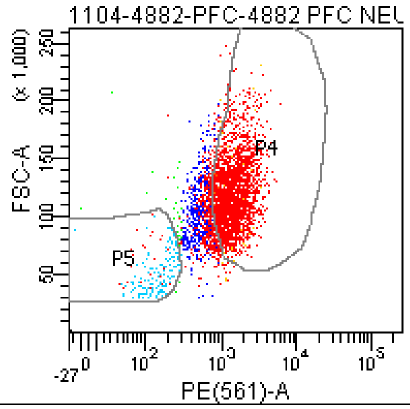


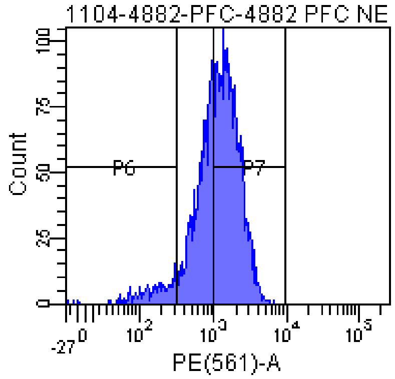


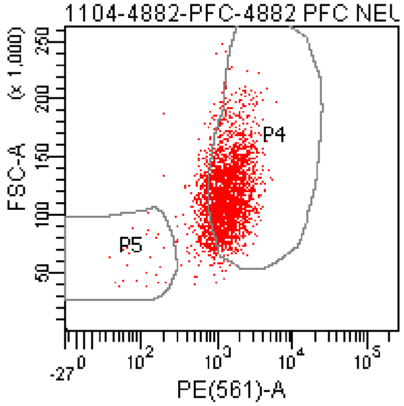


Tube: 4882 PFC NEURON BACK+DAPI

| Population | #Events | %Parent | %Total |
|---|---|---|---|
| All Events | 4,252 | #### | 100.0 |
| P1 | 3,789 | 89.1 | 89.1 |
| P2 | 3,627 | 95.7 | 85.3 |
| P3 | 3,564 | 98.3 | 83.8 |
| P4 | 2,385 | 66.9 | 56.1 |
| P5 | 183 | 5.1 | 4.3 |
| P6 | 223 | 6.3 | 5.2 |
| P7 | 2,041 | 57.3 | 48.0 |
| P8 | 2,752 | 77.2 | 64.7 |
| P9 | 670 | 18.8 | 15.8 |

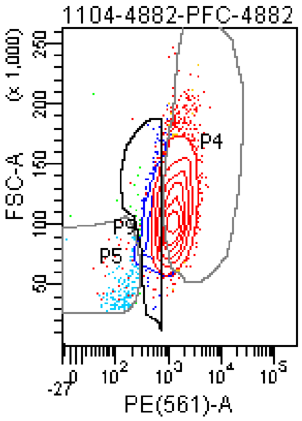


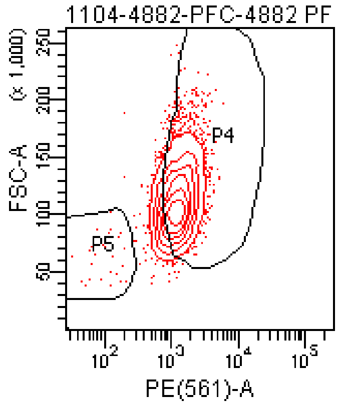

**The BD Arial II FACS sorting reports of UMB#4591**

Sample: prefrontal cortex

Category: healthy control

### FACSDiva Version 6.1.3

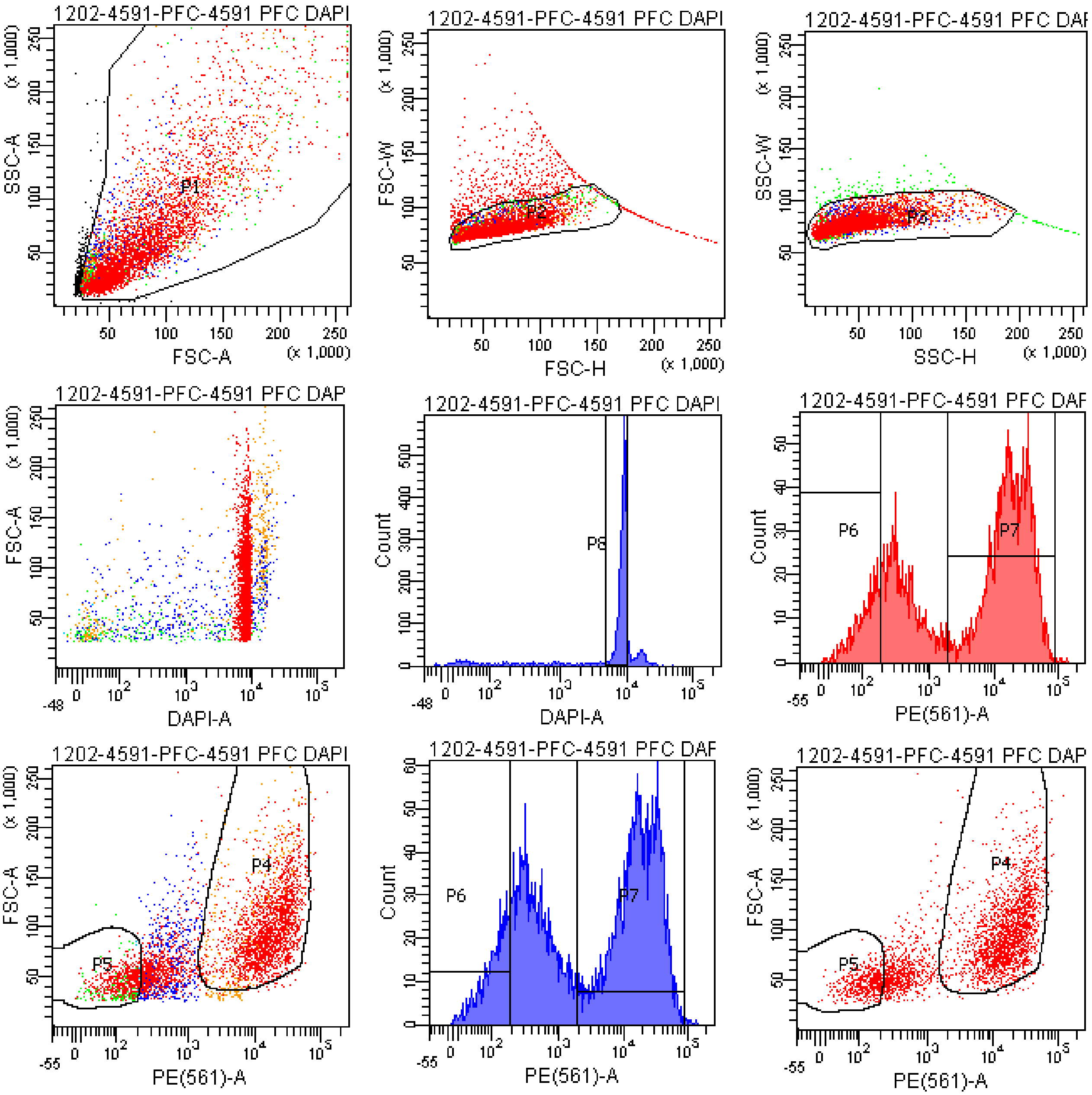


Tube: 4591 PFC DAPI+PE 457V

| Population | #Events | %Parent | %Total |
|---|---|---|---|
| All Events | 5,491 | #### | 100.0 |
| P1 | 5,243 | 95.5 | 95.5 |
| P2 | 4,525 | 86.3 | 82.4 |
| P3 | 4,348 | 96.1 | 79.2 |
| P4 | 2,280 | 52.4 | 41.5 |
| P5 | 708 | 16.3 | 12.9 |
| P6 | 592 | 13.6 | 10.8 |
| P7 | 2,383 | 54.8 | 43.4 |
| P8 | 3,262 | 75.0 | 59.4 |
| P9 | 1,245 | 28.6 | 22.7 |

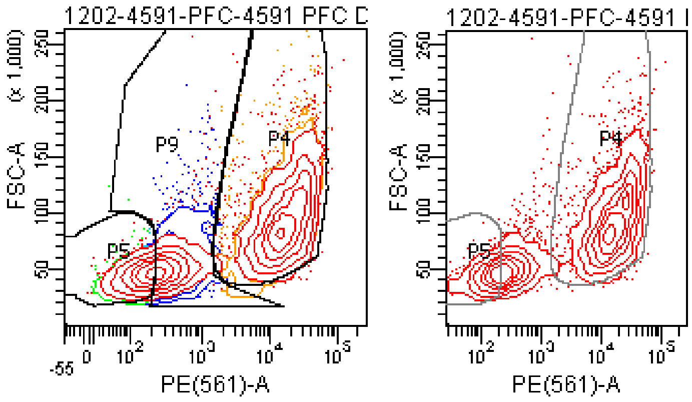

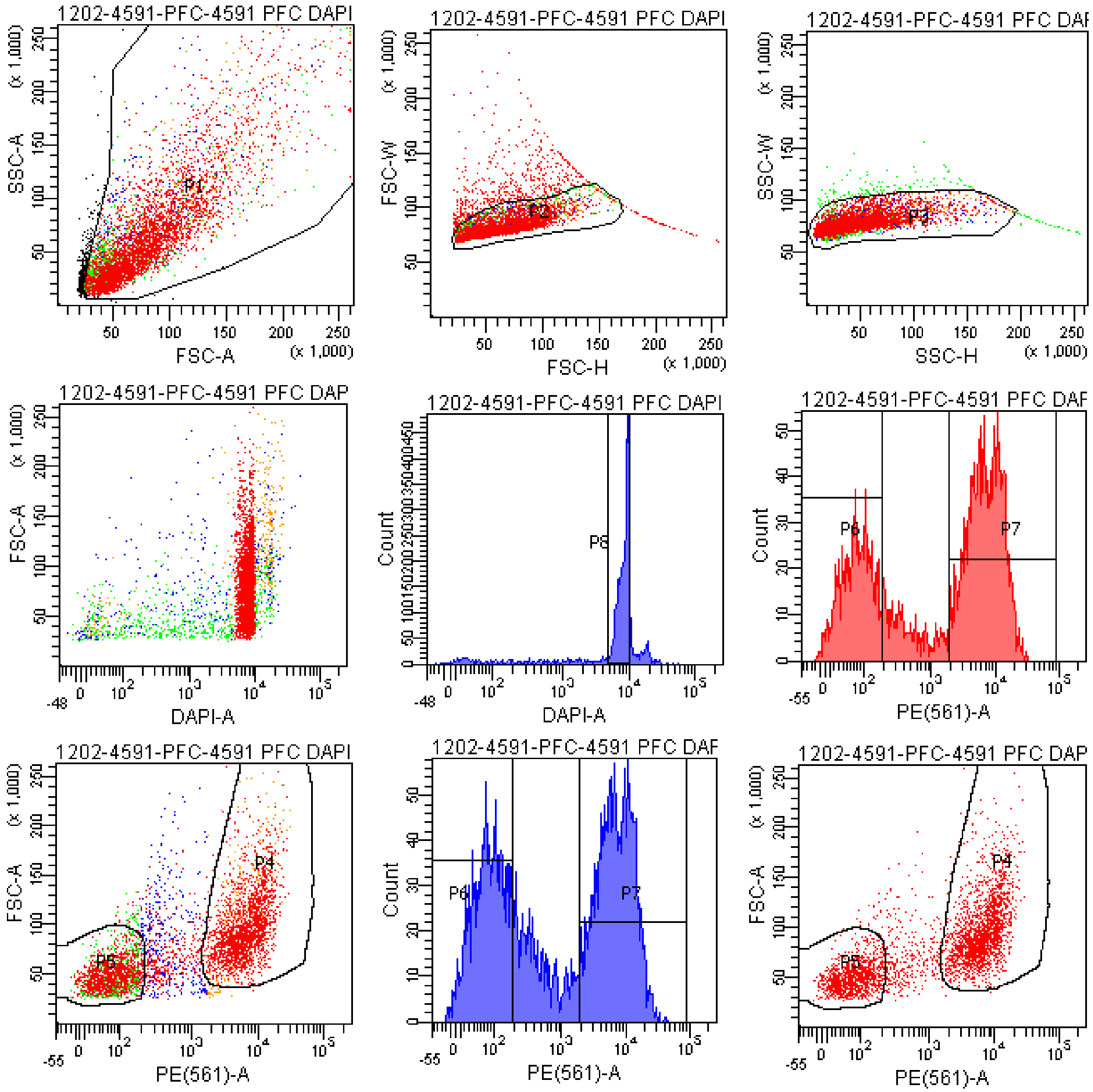


Tube: 4591 PFC DAPI+PE 400V

| Population | #Events | %Parent | %Total |
|---|---|---|---|
| All Events | 5,537 | #### | 100.0 |
| P1 | 5,246 | 94.7 | 94.7 |
| P2 | 4,640 | 88.4 | 83.8 |
| P3 | 4,459 | 96.1 | 80.5 |
| P4 | 2,142 | 48.0 | 38.7 |
| P5 | 1,594 | 35.7 | 28.8 |
| P6 | 1,542 | 34.6 | 27.8 |
| P7 | 2,160 | 48.4 | 39.0 |
| P8 | 3,273 | 73.4 | 59.1 |
| P9 | 700 | 15.7 | 12.6 |

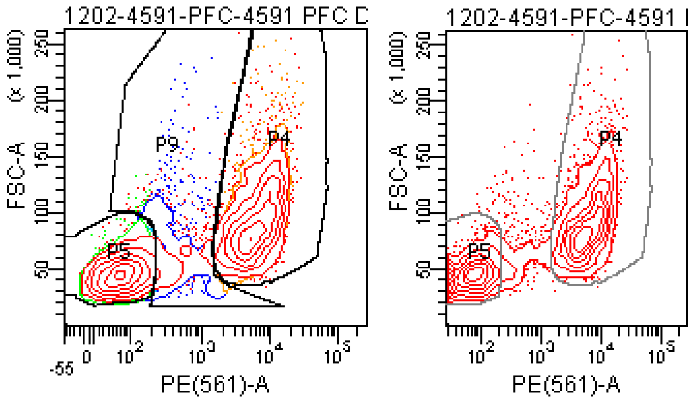

### FACSDiva Version 6.1.3

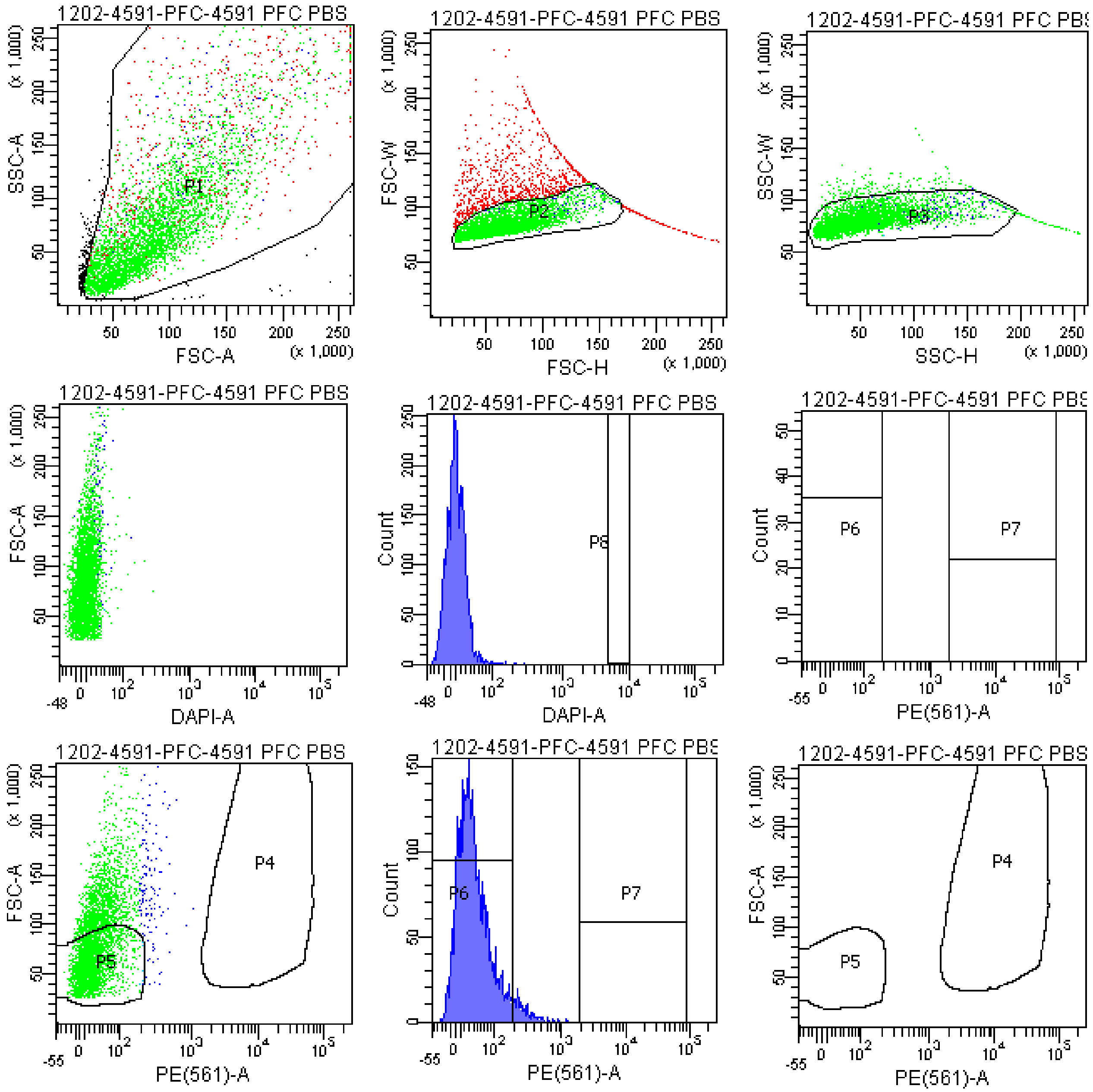


Tube: 4591 PFC PBS CONTROL

| Population | #Events | %Parent | %Total |
|---|---|---|---|
| All Events | 5,512 | #### | 100.0 |
| P1 | 5,258 | 95.4 | 95.4 |
| P2 | 4,238 | 80.6 | 76.9 |
| P3 | 4,001 | 94.4 | 72.6 |
| P4 | 0 | 0.0 | 0.0 |
| P5 | 2,456 | 61.4 | 44.6 |
| P6 | 3,843 | 96.1 | 69.7 |
| P7 | 0 | 0.0 | 0.0 |
| P8 | 0 | 0.0 | 0.0 |
| P9 | 704 | 17.6 | 12.8 |

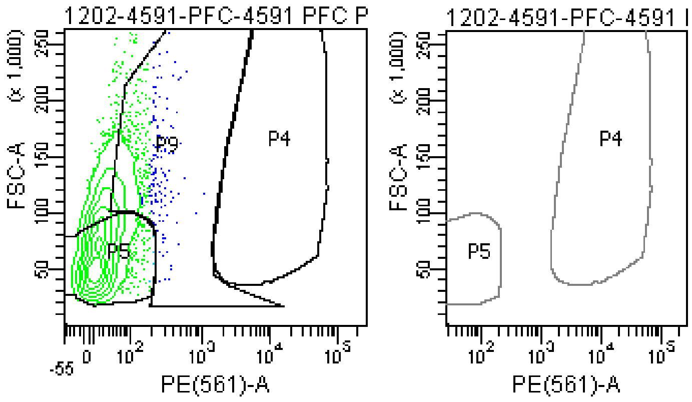

### FACSDiva Version 6.1.3

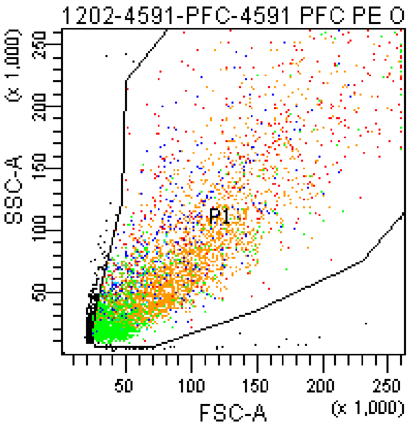


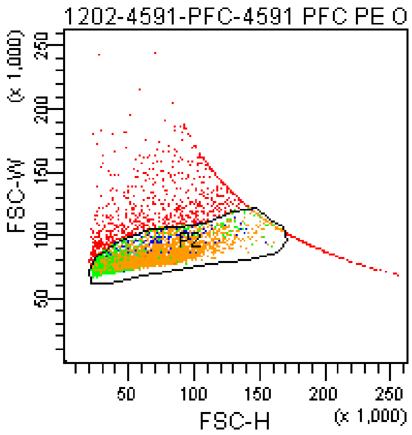


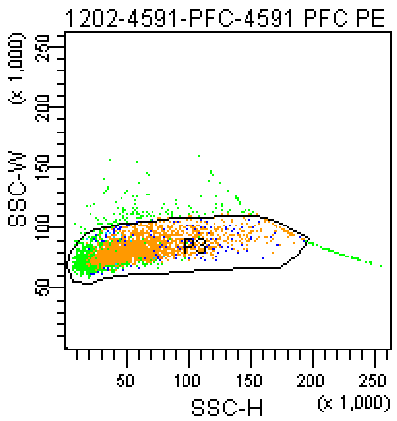


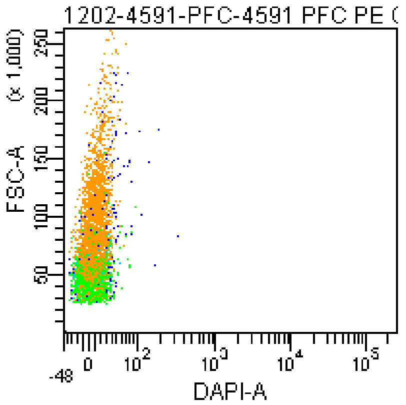


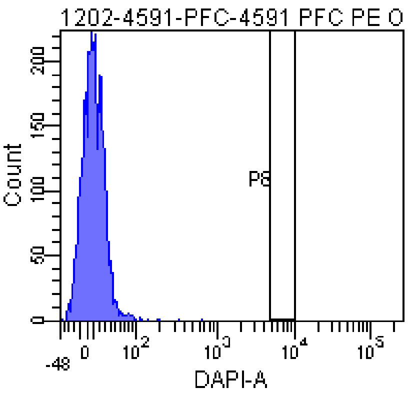


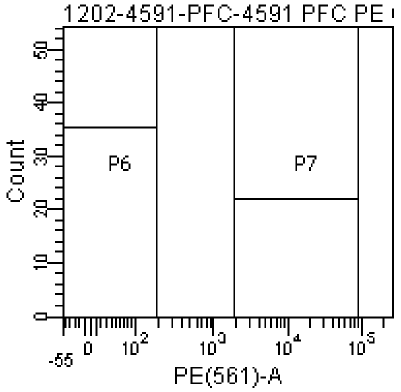


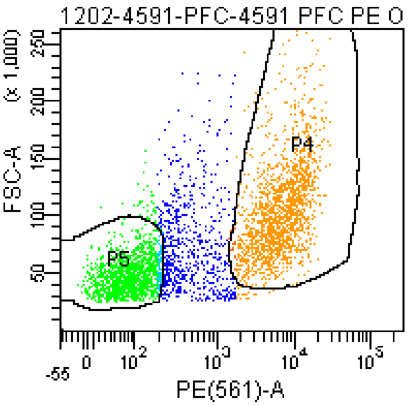


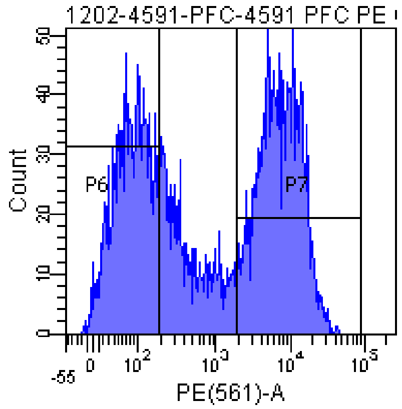


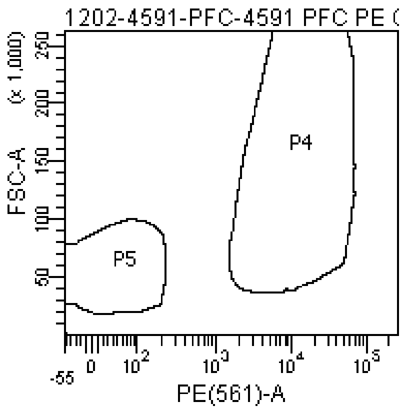


Tube: 4591 PFC PE ONLY CONTROL

| Population | #Events | %Parent | %Total |
|---|---|---|---|
| All Events | 5,570 | #### | 100.0 |
| P1 | 5,188 | 93.1 | 93.1 |
| P2 | 4,300 | 82.9 | 77.2 |
| P3 | 4,070 | 94.7 | 73.1 |
| P4 | 1,824 | 44.8 | 32.7 |
| P5 | 1,501 | 36.9 | 26.9 |
| P6 | 1,417 | 34.8 | 25.4 |
| P7 | 1,839 | 45.2 | 33.0 |
| P8 | 0 | 0.0 | 0.0 |
| P9 | 719 | 17.7 | 12.9 |

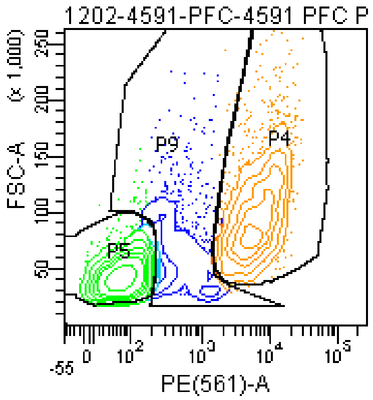


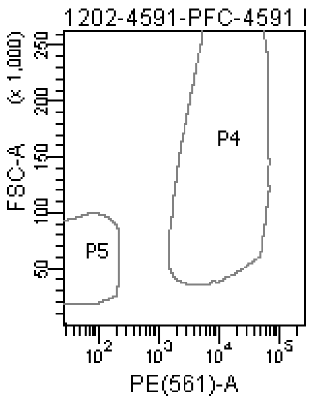

### FACSDiva Version 6.1.3

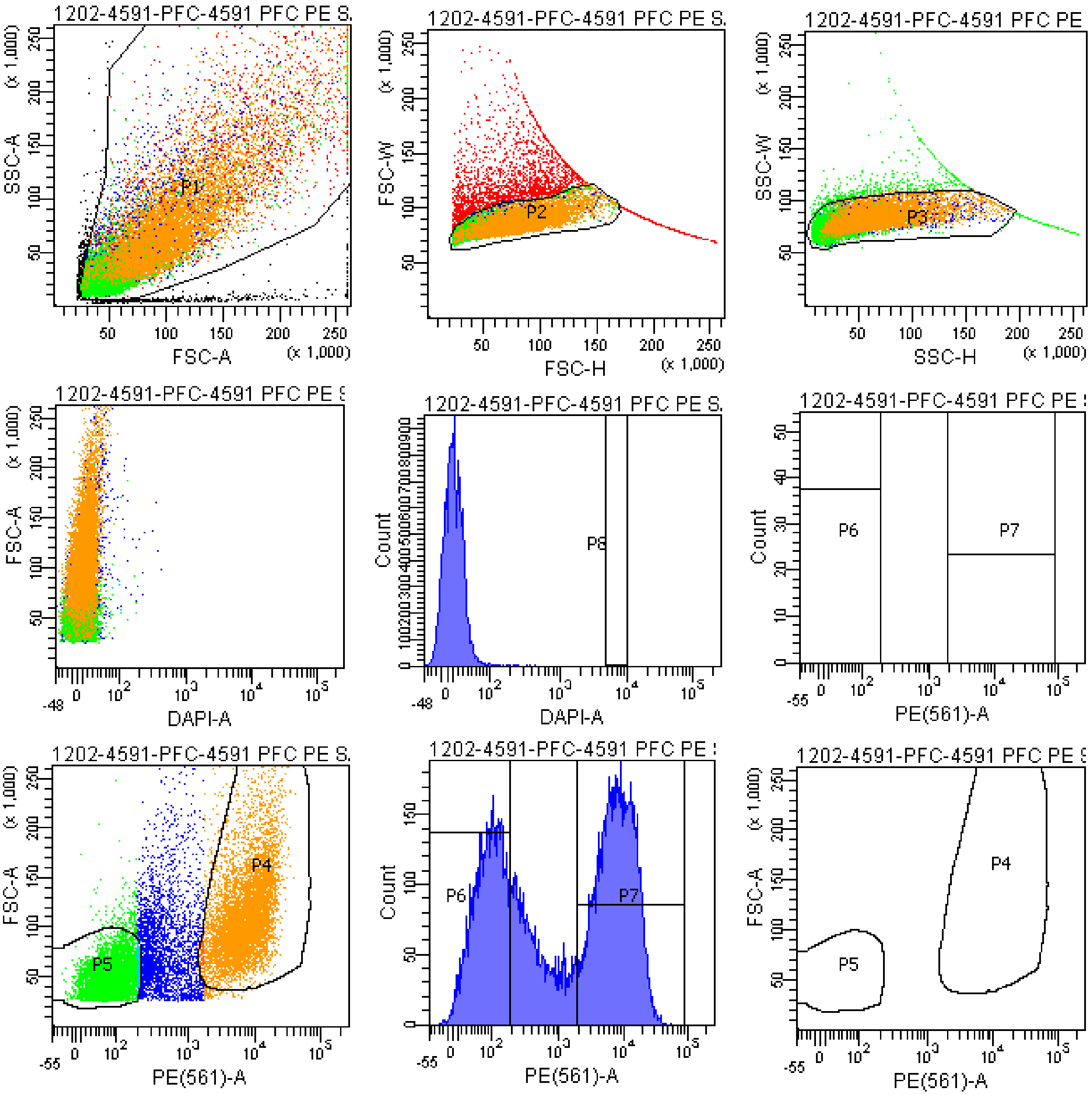


Tube: 4591 PFC PE SAMPLE

| Population | #Events | %Parent | %Total |
|---|---|---|---|
| All Events | 20,982 | #### | 100.0 |
| P1 | 20,000 | 95.3 | 95.3 |
| P2 | 16,723 | 83.6 | 79.7 |
| P3 | 16,035 | 95.9 | 76.4 |
| P4 | 7,586 | 47.3 | 36.2 |
| P5 | 5,352 | 33.4 | 25.5 |
| P6 | 5,137 | 32.0 | 24.5 |
| P7 | 7,646 | 47.7 | 36.4 |
| P8 | 0 | 0.0 | 0.0 |
| P9 | 3,021 | 18.8 | 14.4 |

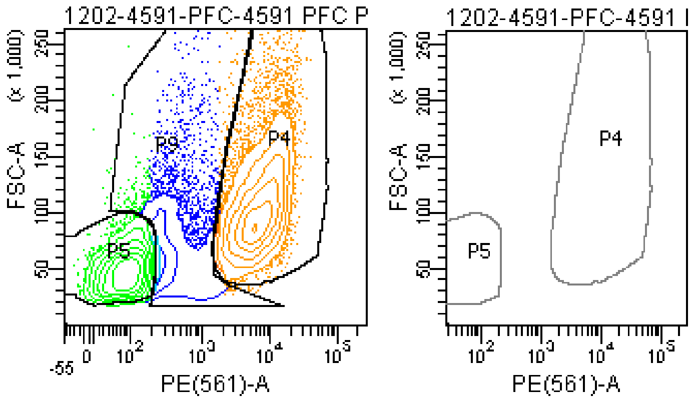

### FACSDiva Version 6.1.3

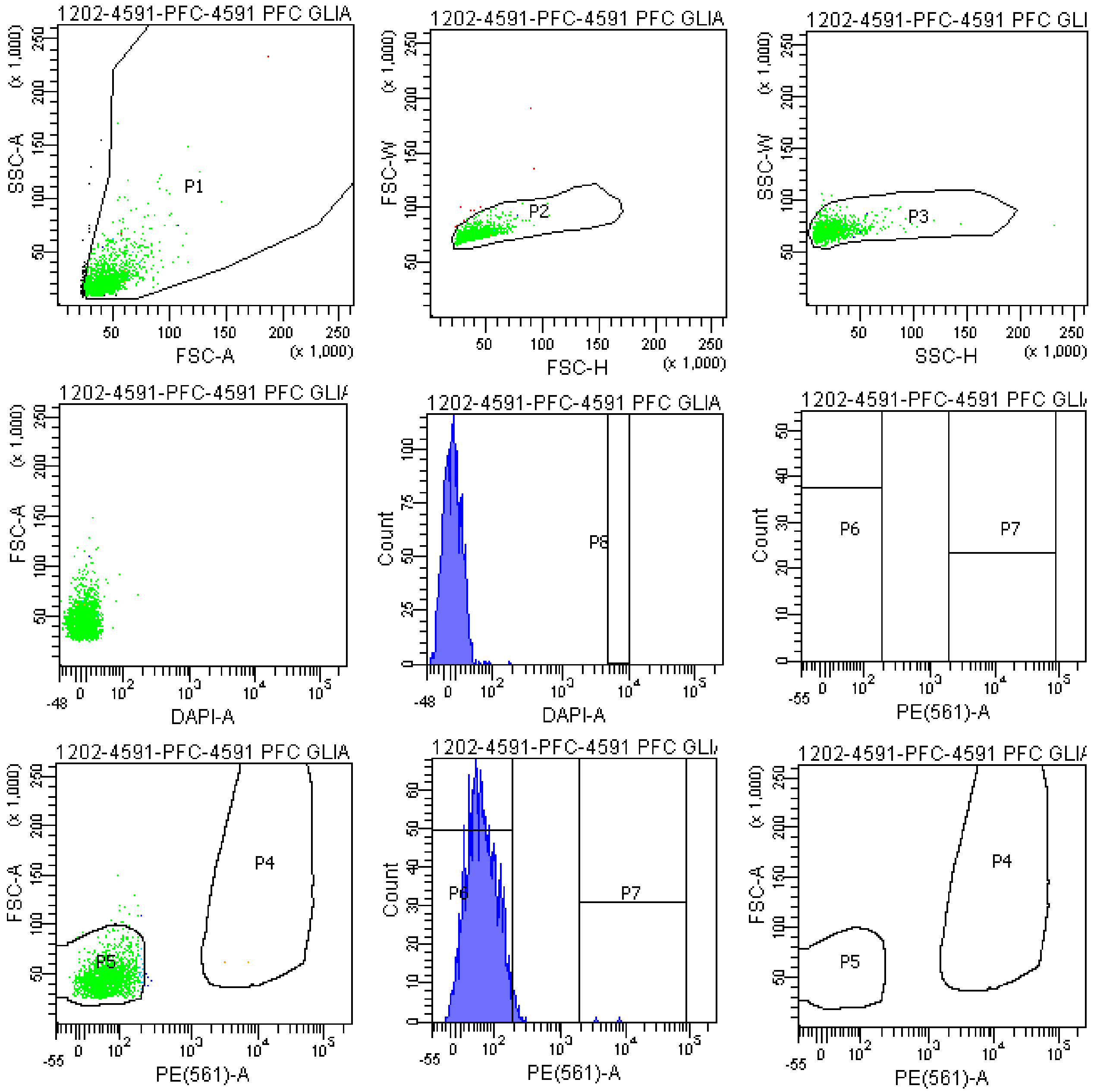


Tube: 4591 PFC GLIA BACK

| Population | #Events | %Parent | %Total |
|---|---|---|---|
| All Events | 2,145 | #### | 100.0 |
| P1 | 2,052 | 95.7 | 95.7 |
| P2 | 2,043 | 99.6 | 95.2 |
| P3 | 2,036 | 99.7 | 94.9 |
| P4 | 2 | 0.1 | 0.1 |
| P5 | 2,011 | 98.8 | 93.8 |
| P6 | 2,006 | 98.5 | 93.5 |
| P7 | 2 | 0.1 | 0.1 |
| P8 | 0 | 0.0 | 0.0 |
| P9 | 17 | 0.8 | 0.8 |

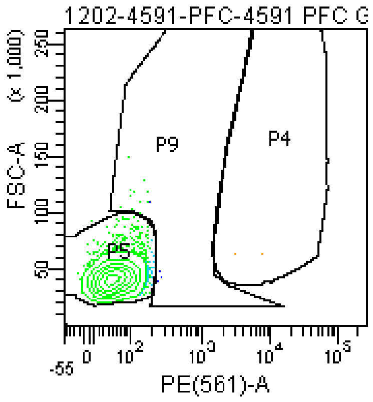


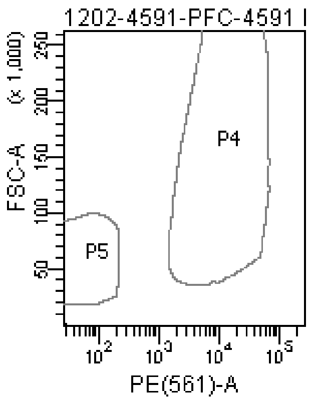

### FACSDiva Version 6.1.3

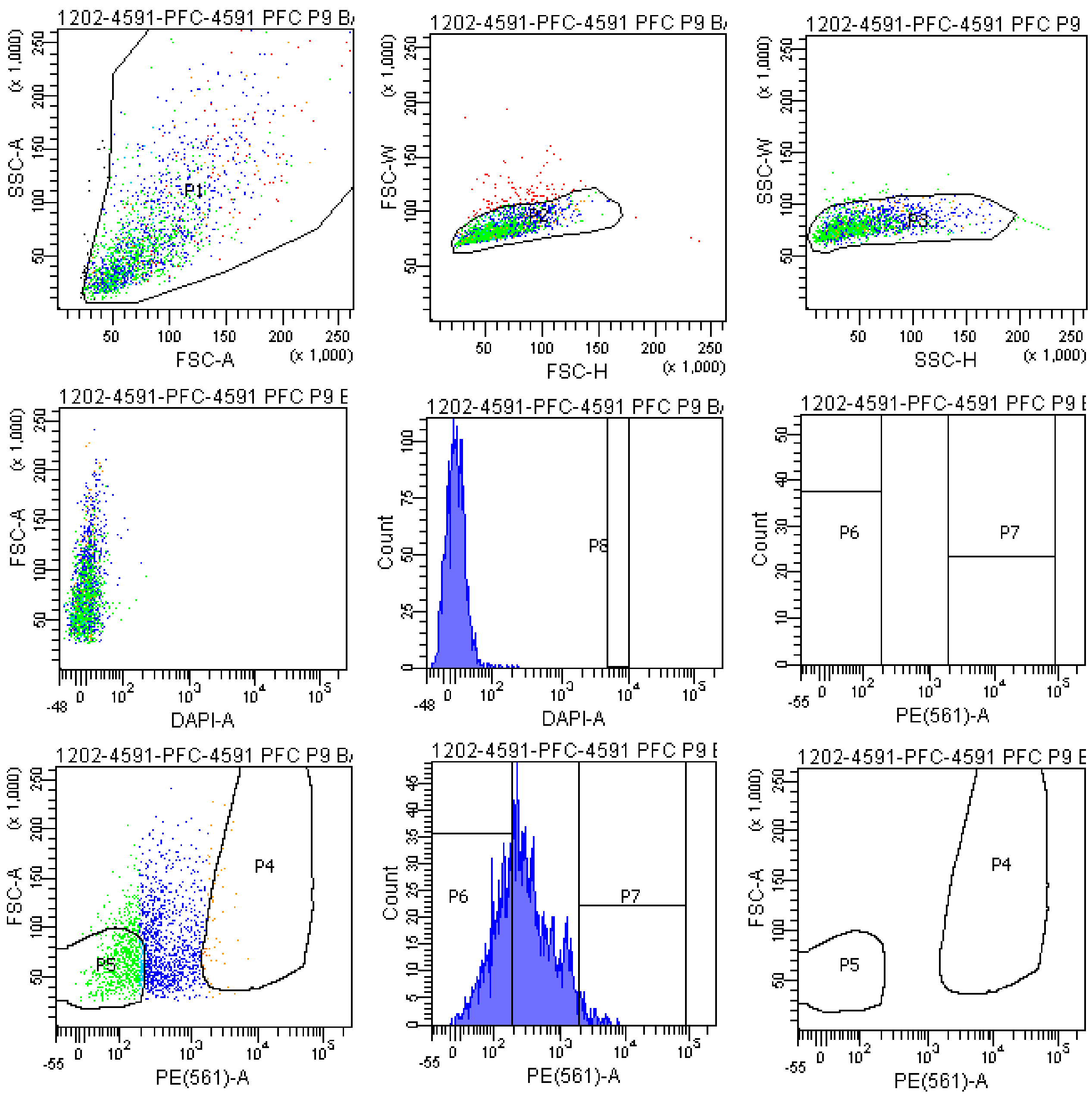


Tube: 4591 PFC P9 BACK

| Population | #Events | %Parent | %Total |
|---|---|---|---|
| All Events | 2,106 | #### | 100.0 |
| P1 | 2,069 | 98.2 | 98.2 |
| P2 | 1,962 | 94.8 | 93.2 |
| P3 | 1,920 | 97.9 | 91.2 |
| P4 | 47 | 2.4 | 2.2 |
| P5 | 682 | 35.5 | 32.4 |
| P6 | 688 | 35.8 | 32.7 |
| P7 | 50 | 2.6 | 2.4 |
| P8 | 0 | 0.0 | 0.0 |
| P9 | 1,173 | 61.1 | 55.7 |

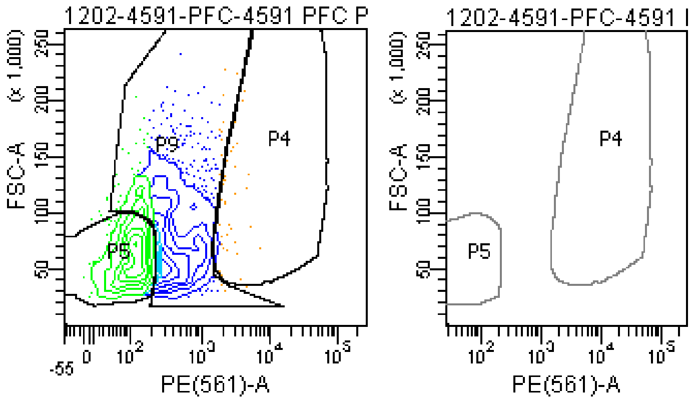

### FACSDiva Version 6.1.3

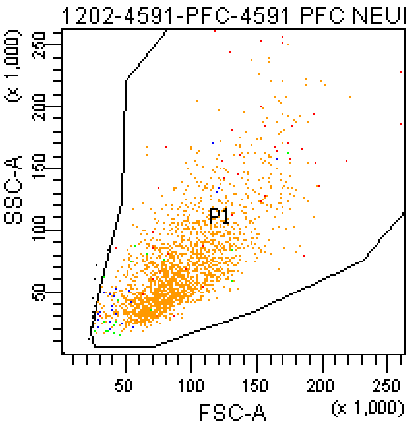


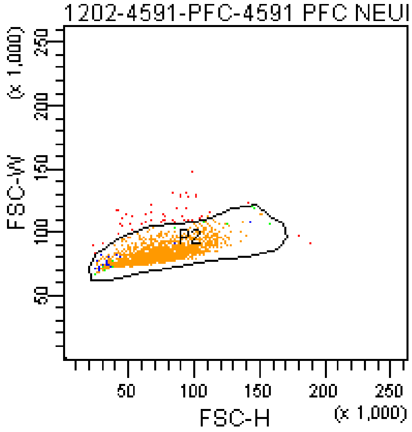


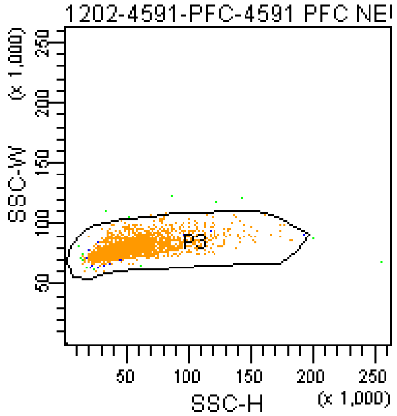


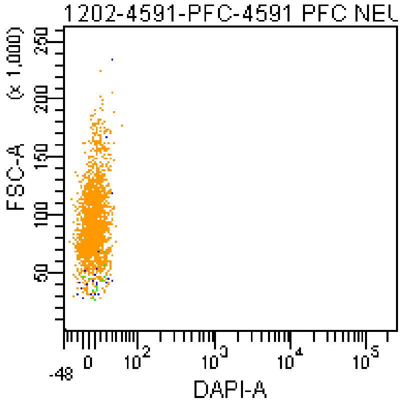


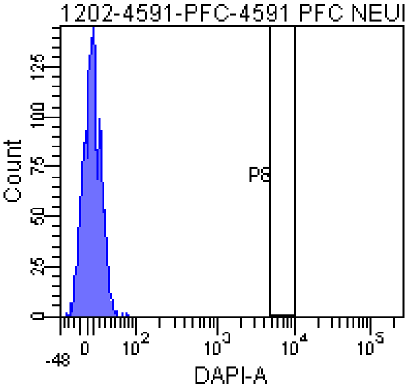


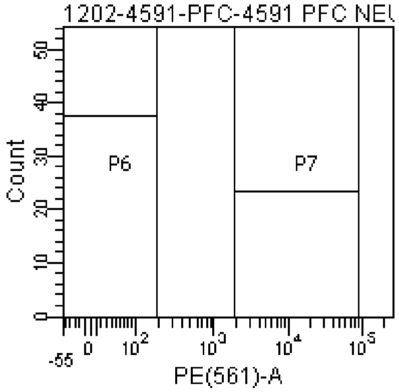


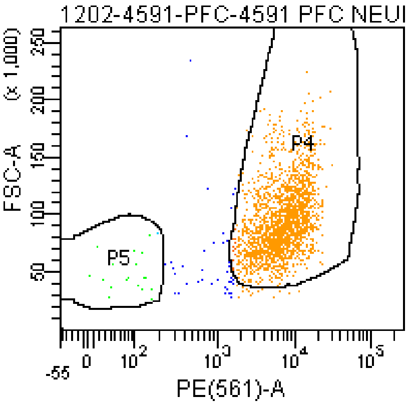


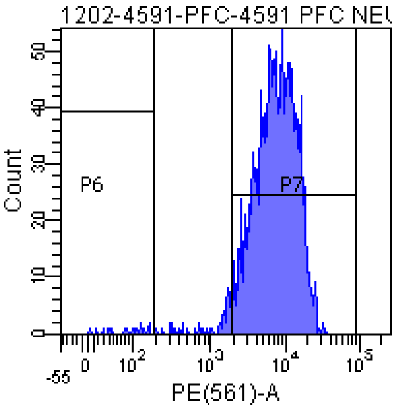


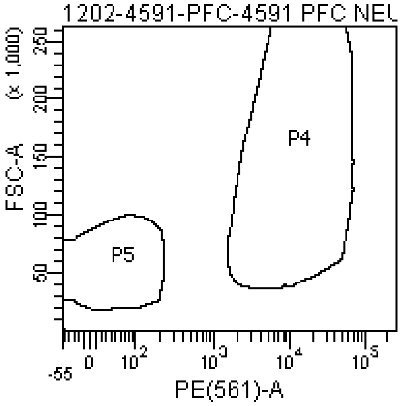


Tube: 4591 PFC NEURON BACK

| Population | #Events | %Parent | %Total |
|---|---|---|---|
| All Events | 2,127 | #### | 100.0 |
| P1 | 2,114 | 99.4 | 99.4 |
| P2 | 2,065 | 97.7 | 97.1 |
| P3 | 2,058 | 99.7 | 96.8 |
| P4 | 1,973 | 95.9 | 92.8 |
| P5 | 20 | 1.0 | 0.9 |
| P6 | 19 | 0.9 | 0.9 |
| P7 | 1,980 | 96.2 | 93.1 |
| P8 | 0 | 0.0 | 0.0 |
| P9 | 56 | 2.7 | 2.6 |

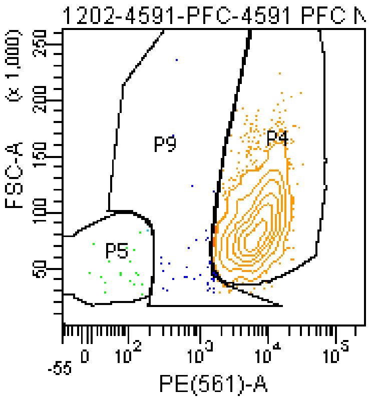


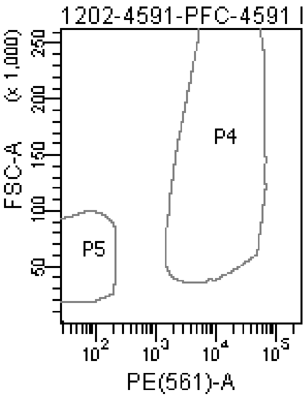

**The BD Arial II FACS sorting reports of UMB#1420**

Sample: prefrontal cortex

Category: Rett patient

### FACSDiva Version 6.1.3

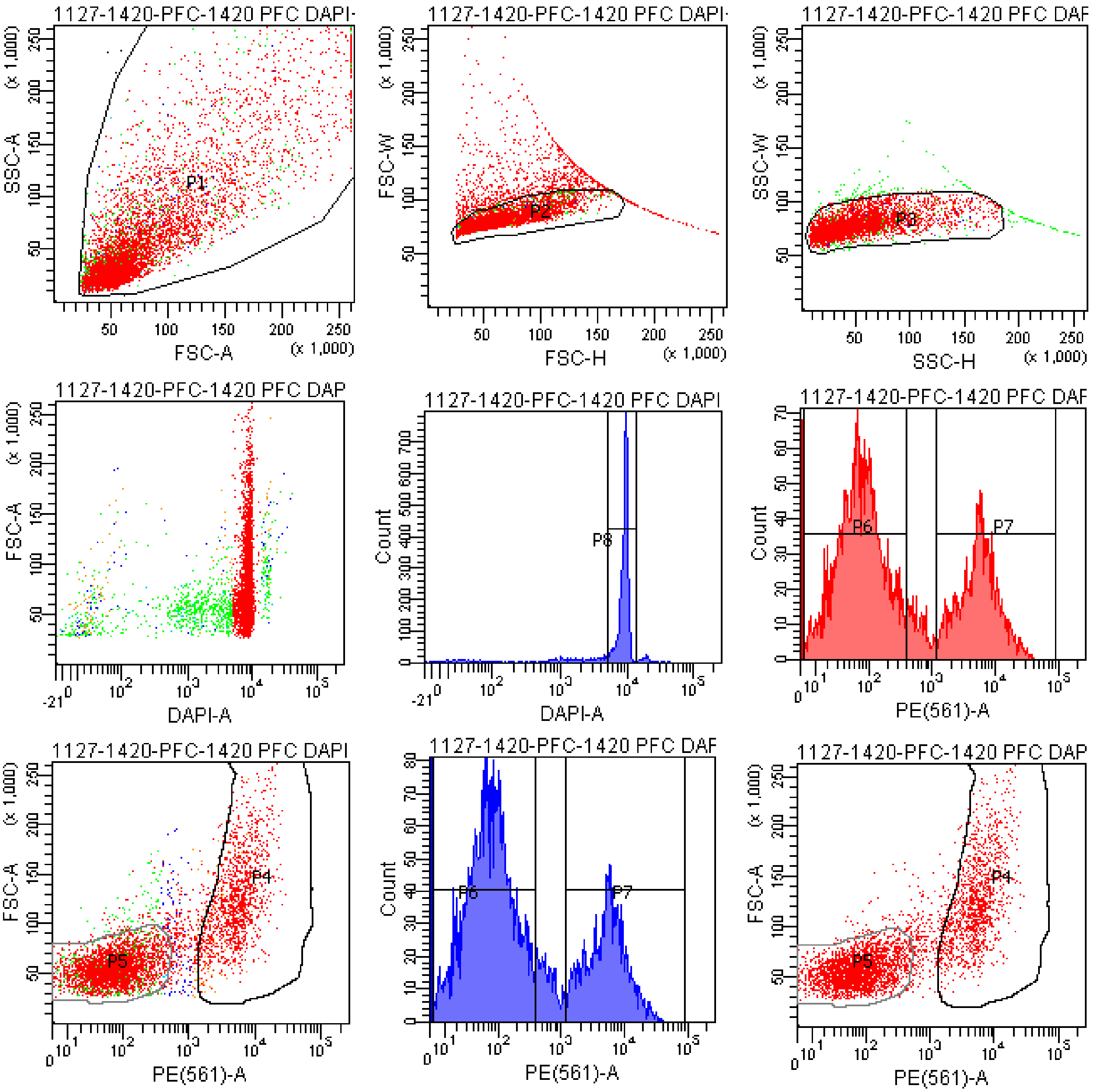


Tube: 1420 PFC DAPI+PE 457V

| Population | #Events | %Parent | %Total |
|---|---|---|---|
| All Events | 6,578 | #### | 100.0 |
| P1 | 6,564 | 99.8 | 99.8 |
| P2 | 5,553 | 84.6 | 84.4 |
| P3 | 5,379 | 96.9 | 81.8 |
| P4 | 1,392 | 25.9 | 21.2 |
| P5 | 3,514 | 65.3 | 53.4 |
| P6 | 3,418 | 63.5 | 52.0 |
| P7 | 1,482 | 27.6 | 22.5 |
| P8 | 4,411 | 82.0 | 67.1 |
| P9 | 400 | 7.4 | 6.1 |

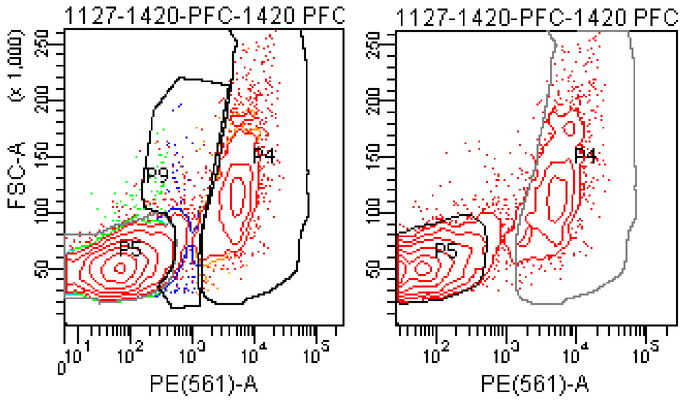

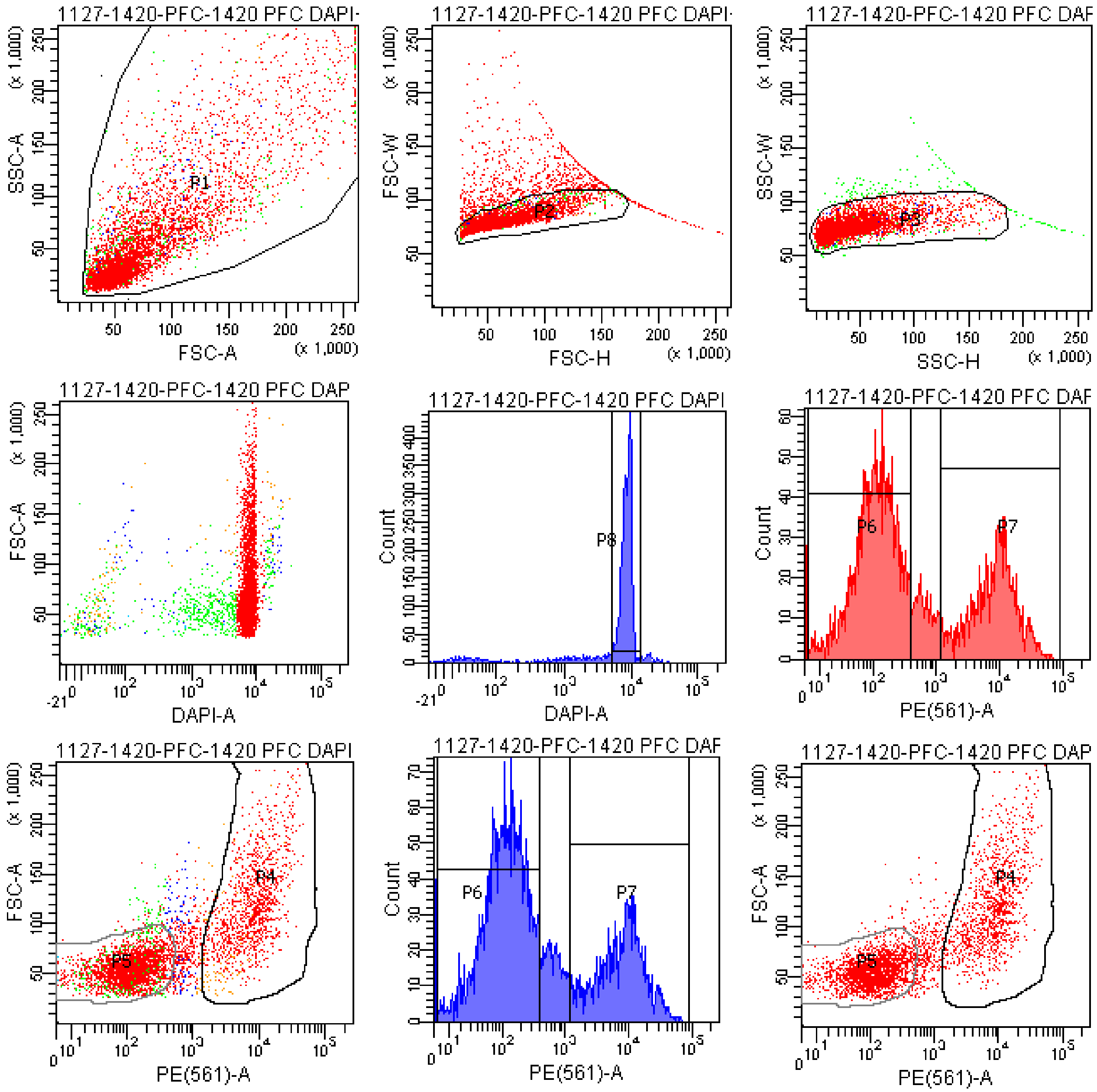


Tube: 1420 PFC DAPI+PE 500V

| Population | #Events | %Parent | %Total |
|---|---|---|---|
| All Events | 5,528 | #### | 100.0 |
| P1 | 5,519 | 99.8 | 99.8 |
| P2 | 4,628 | 83.9 | 83.7 |
| P3 | 4,465 | 96.5 | 80.8 |
| P4 | 1,248 | 28.0 | 22.6 |
| P5 | 2,697 | 60.4 | 48.8 |
| P6 | 2,640 | 59.1 | 47.8 |
| P7 | 1,311 | 29.4 | 23.7 |
| P8 | 3,665 | 82.1 | 66.3 |
| P9 | 496 | 11.1 | 9.0 |

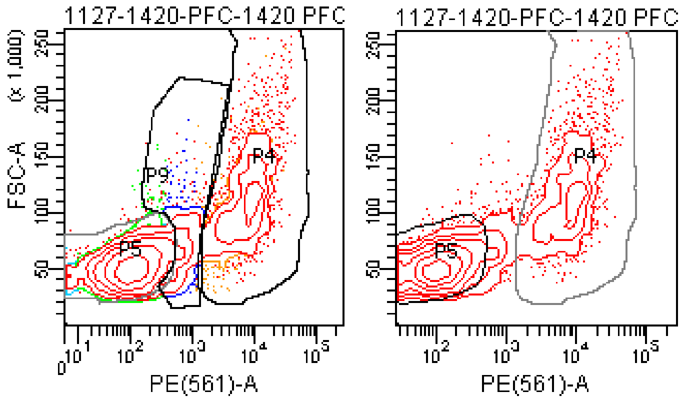

### FACSDiva Version 6.1.3

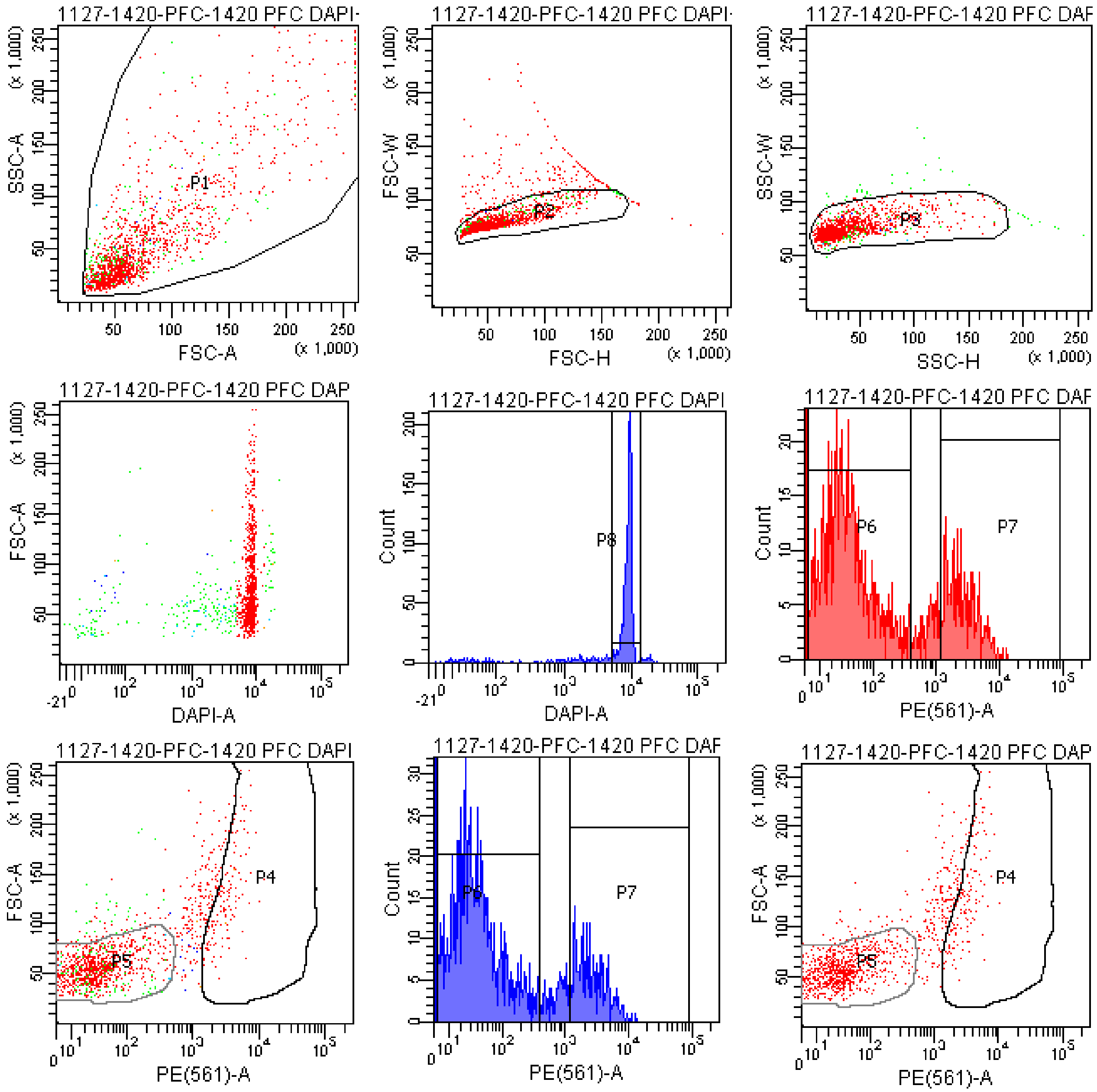


Tube: 1420 PFC DAPI+PE 400V

| Population | #Events | %Parent | %Total |
|---|---|---|---|
| All Events | 1,614 | #### | 100.0 |
| P1 | 1,611 | 99.8 | 99.8 |
| P2 | 1,403 | 87.1 | 86.9 |
| P3 | 1,360 | 96.9 | 84.3 |
| P4 | 143 | 10.5 | 8.9 |
| P5 | 981 | 72.1 | 60.8 |
| P6 | 903 | 66.4 | 55.9 |
| P7 | 239 | 17.6 | 14.8 |
| P8 | 1,134 | 83.4 | 70.3 |
| P9 | 176 | 12.9 | 10.9 |

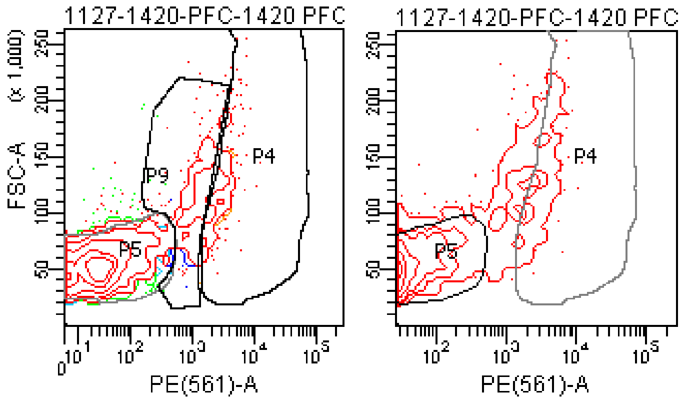

### FACSDiva Version 6.1.3

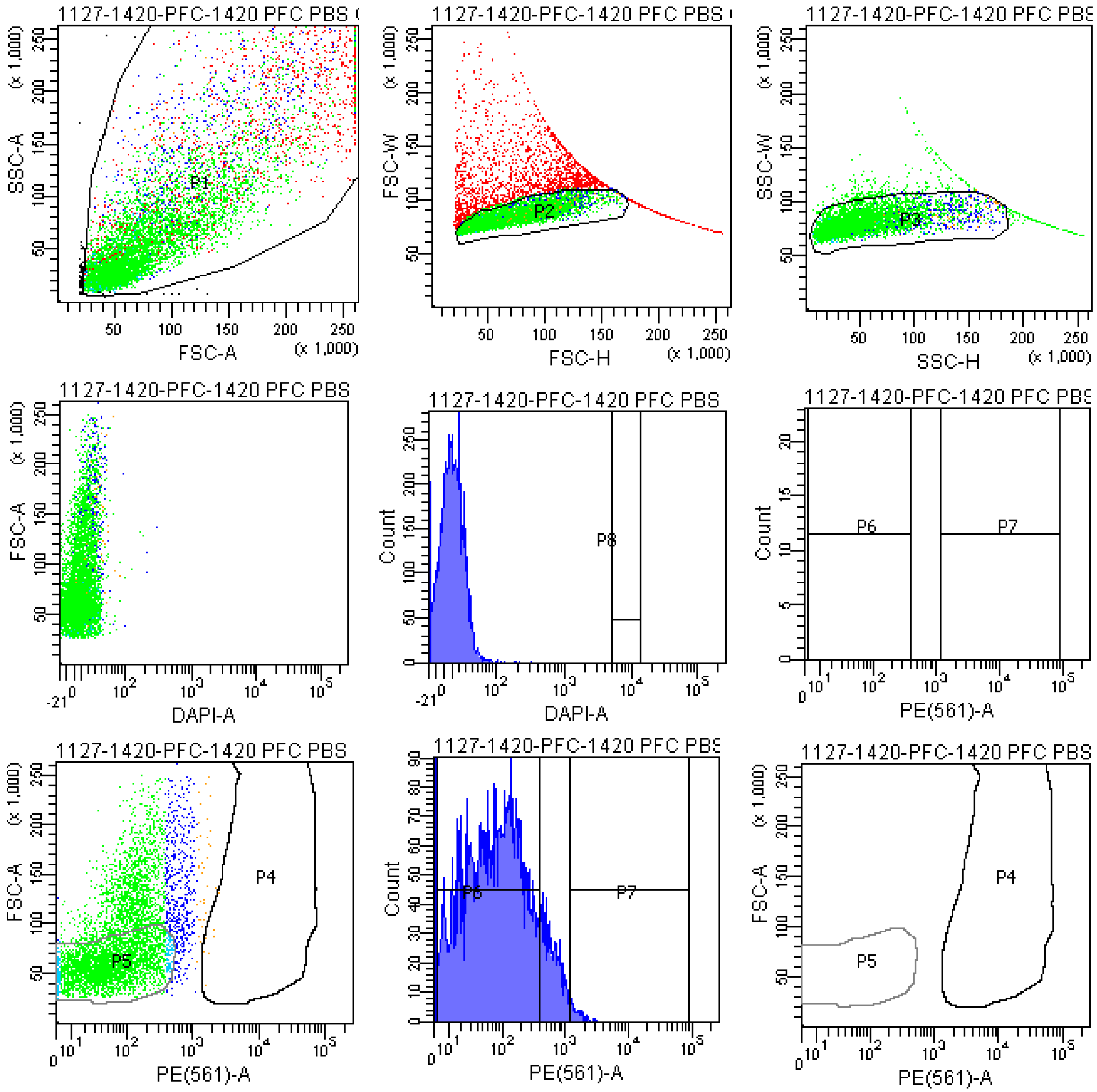


Tube: 1420 PFC PBS CONTROL

| Population | #Events | %Parent | %Total |
|---|---|---|---|
| All Events | 9,017 | #### | 100.0 |
| P1 | 8,925 | 99.0 | 99.0 |
| P2 | 6,391 | 71.6 | 70.9 |
| P3 | 5,975 | 93.5 | 66.3 |
| P4 | 8 | 0.1 | 0.1 |
| P5 | 3,877 | 64.9 | 43.0 |
| P6 | 4,690 | 78.5 | 52.0 |
| P7 | 50 | 0.8 | 0.6 |
| P8 | 0 | 0.0 | 0.0 |
| P9 | 898 | 15.0 | 10.0 |

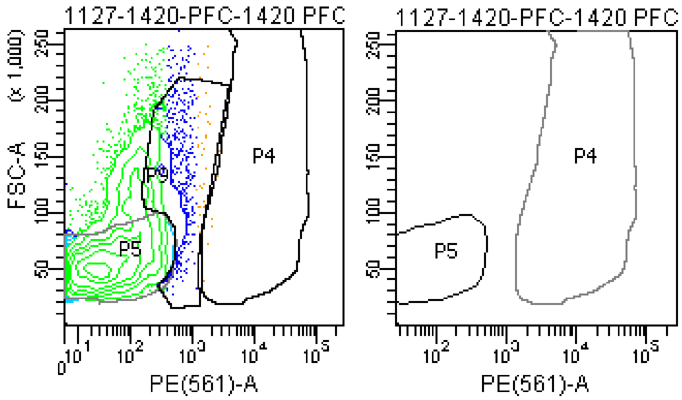

### FACSDiva Version 6.1.3

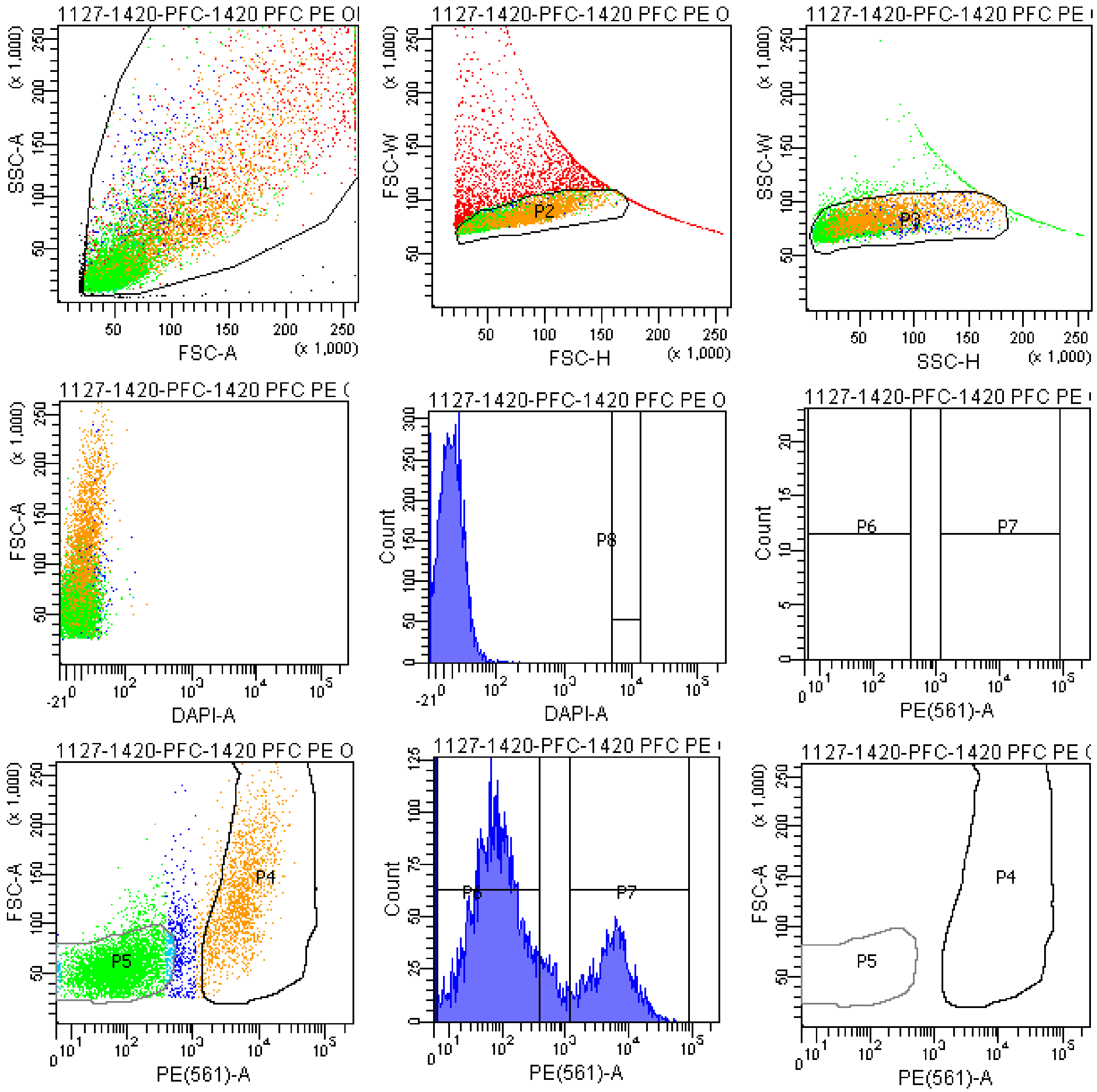


Tube: 1420 PFC PE ONLY SAMPLE

| Population | #Events | %Parent | %Total |
|---|---|---|---|
| All Events | 10,168 | #### | 100.0 |
| P1 | 10,000 | 98.3 | 98.3 |
| P2 | 7,642 | 76.4 | 75.2 |
| P3 | 7,300 | 95.5 | 71.8 |
| P4 | 1,626 | 22.3 | 16.0 |
| P5 | 4,939 | 67.7 | 48.6 |
| P6 | 4,824 | 66.1 | 47.4 |
| P7 | 1,744 | 23.9 | 17.2 |
| P8 | 0 | 0.0 | 0.0 |
| P9 | 613 | 8.4 | 6.0 |

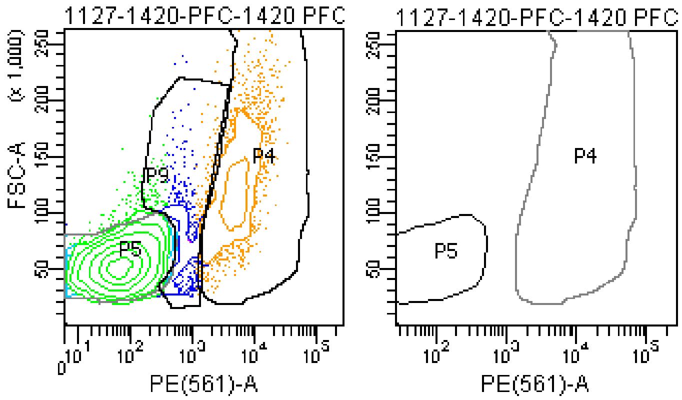

### FACSDiva Version 6.1.3

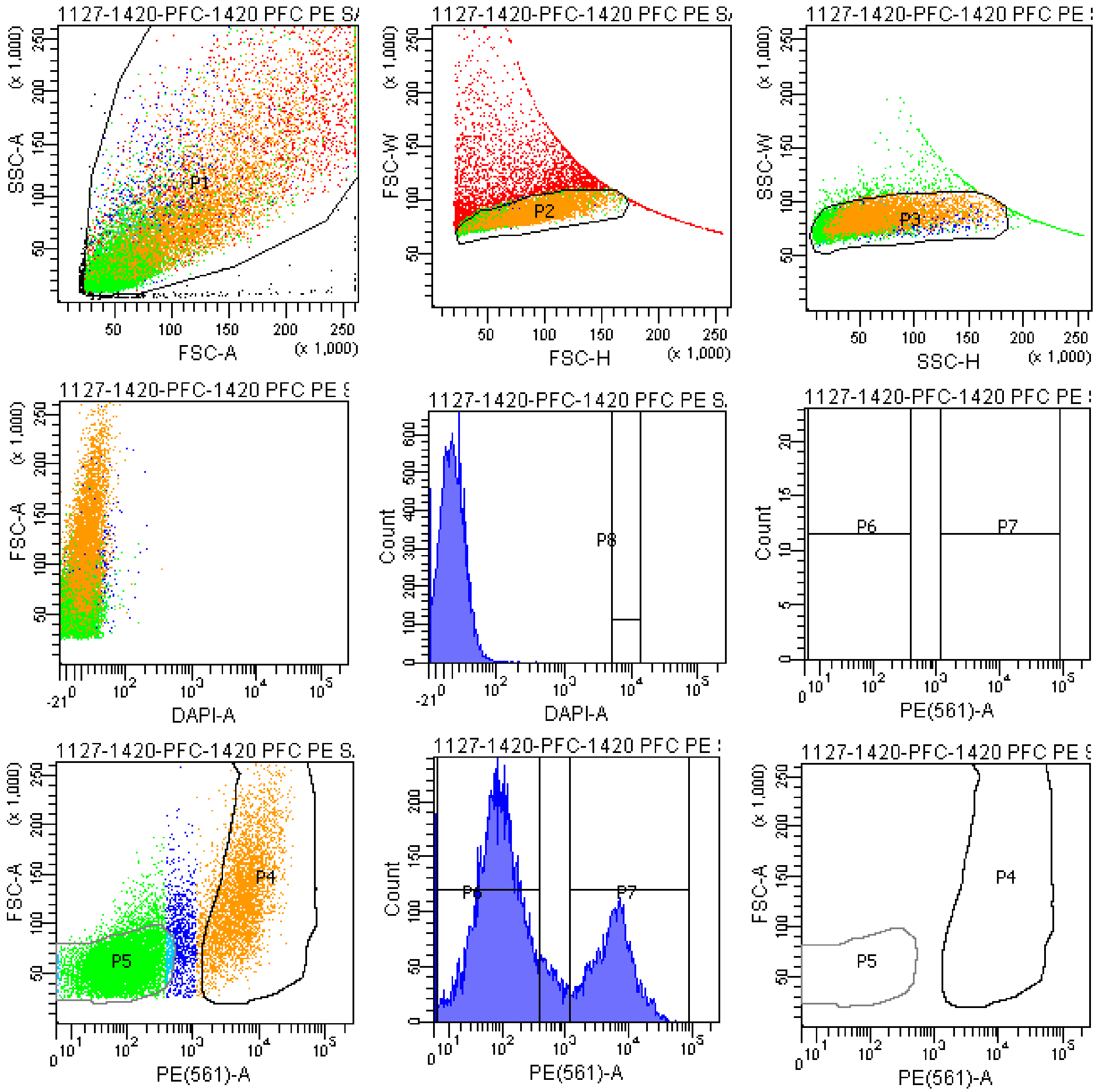


Tube: 1420 PFC PE SAMPLE

| Population | #Events | %Parent | %Total |
|---|---|---|---|
| All Events | 20,378 | #### | 100.0 |
| P1 | 20,000 | 98.1 | 98.1 |
| P2 | 15,820 | 79.1 | 77.6 |
| P3 | 15,235 | 96.3 | 74.8 |
| P4 | 3,911 | 25.7 | 19.2 |
| P5 | 9,697 | 63.6 | 47.6 |
| P6 | 9,709 | 63.7 | 47.6 |
| P7 | 4,181 | 27.4 | 20.5 |
| P8 | 0 | 0.0 | 0.0 |
| P9 | 1,308 | 8.6 | 6.4 |

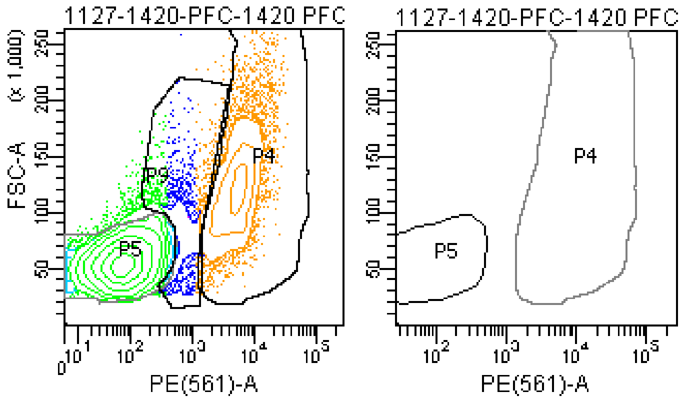

### FACSDiva Version 6.1.3

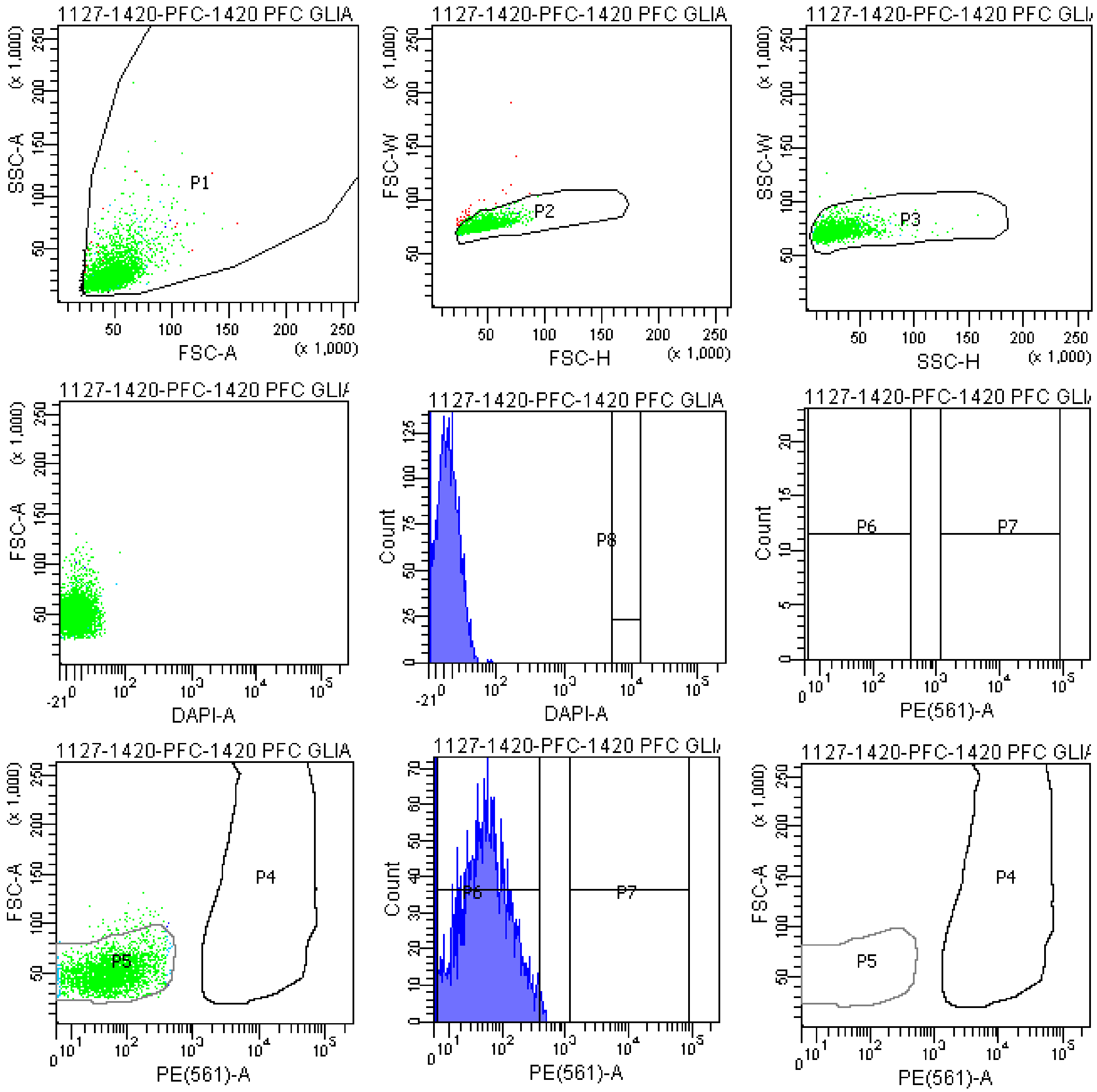


Tube: 1420 PFC GLIA BACK

| Population | #Events | %Parent | %Total |
|---|---|---|---|
| All Events | 3,174 | #### | 100.0 |
| P1 | 3,128 | 98.6 | 98.6 |
| P2 | 3,084 | 98.6 | 97.2 |
| P3 | 3,079 | 99.8 | 97.0 |
| P4 | 0 | 0.0 | 0.0 |
| P5 | 3,020 | 98.1 | 95.1 |
| P6 | 2,865 | 93.0 | 90.3 |
| P7 | 0 | 0.0 | 0.0 |
| P8 | 0 | 0.0 | 0.0 |
| P9 | 27 | 0.9 | 0.9 |

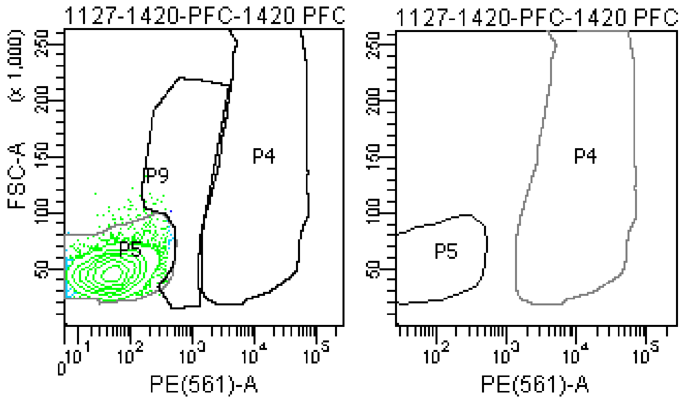

### FACSDiva Version 6.1.3

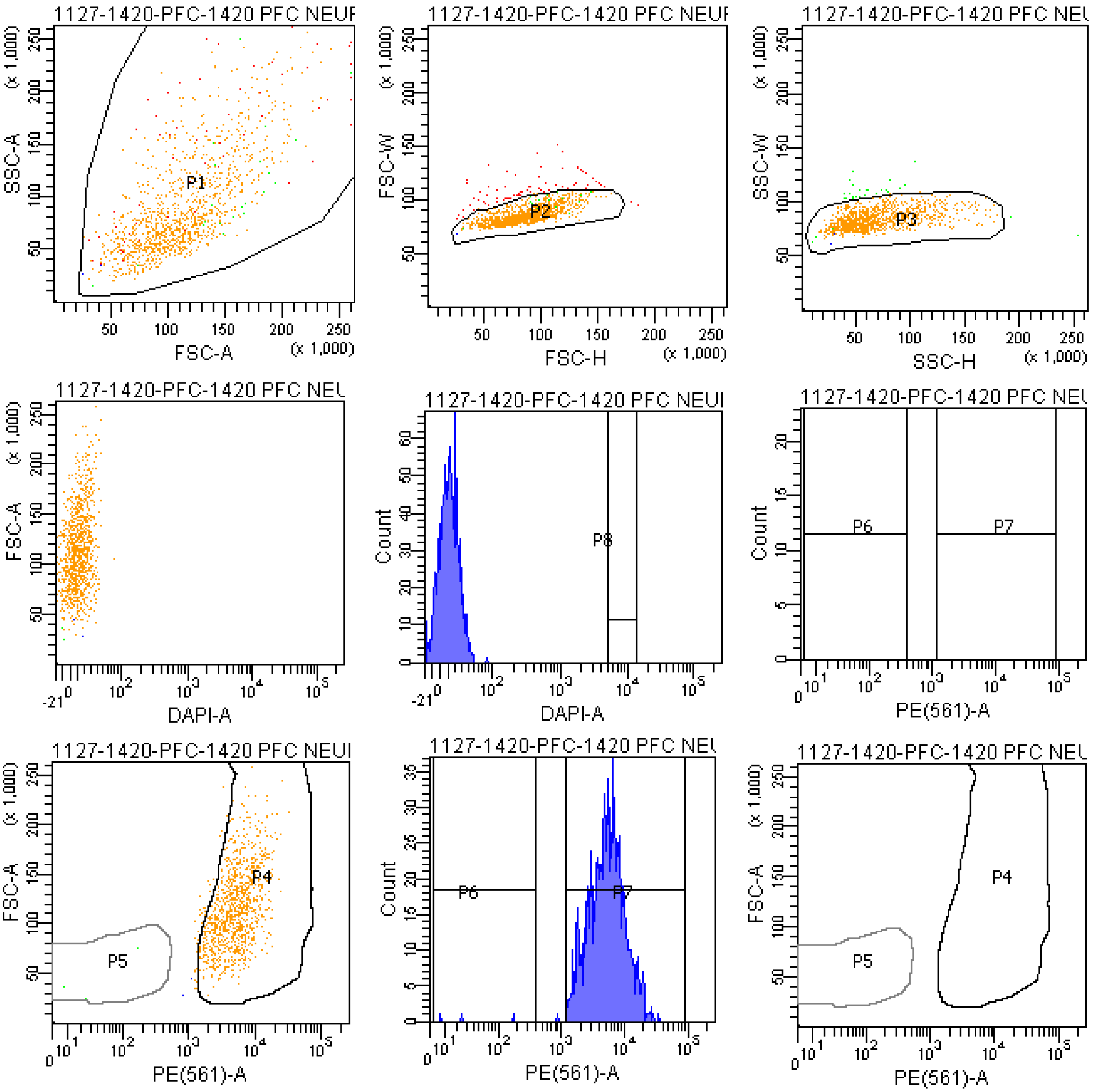


Tube: 1420 PFC NEURON BACK

| Population | #Events | %Parent | %Total |
|---|---|---|---|
| All Events | 1,156 | #### | 100.0 |
| P1 | 1,154 | 99.8 | 99.8 |
| P2 | 1,081 | 93.7 | 93.5 |
| P3 | 1,056 | 97.7 | 91.3 |
| P4 | 1,019 | 96.5 | 88.1 |
| P5 | 3 | 0.3 | 0.3 |
| P6 | 3 | 0.3 | 0.3 |
| P7 | 1,051 | 99.5 | 90.9 |
| P8 | 0 | 0.0 | 0.0 |
| P9 | 21 | 2.0 | 1.8 |

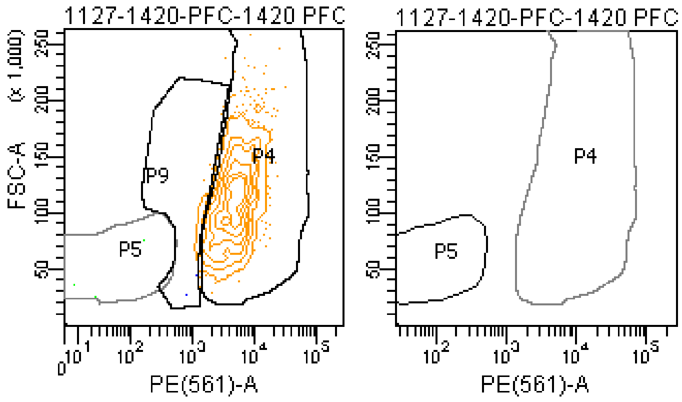

**The BD Arial II FACS sorting reports of UMB#1455**

Sample: prefrontal cortex

Category: healthy control

### FACSDiva Version 6.1.3

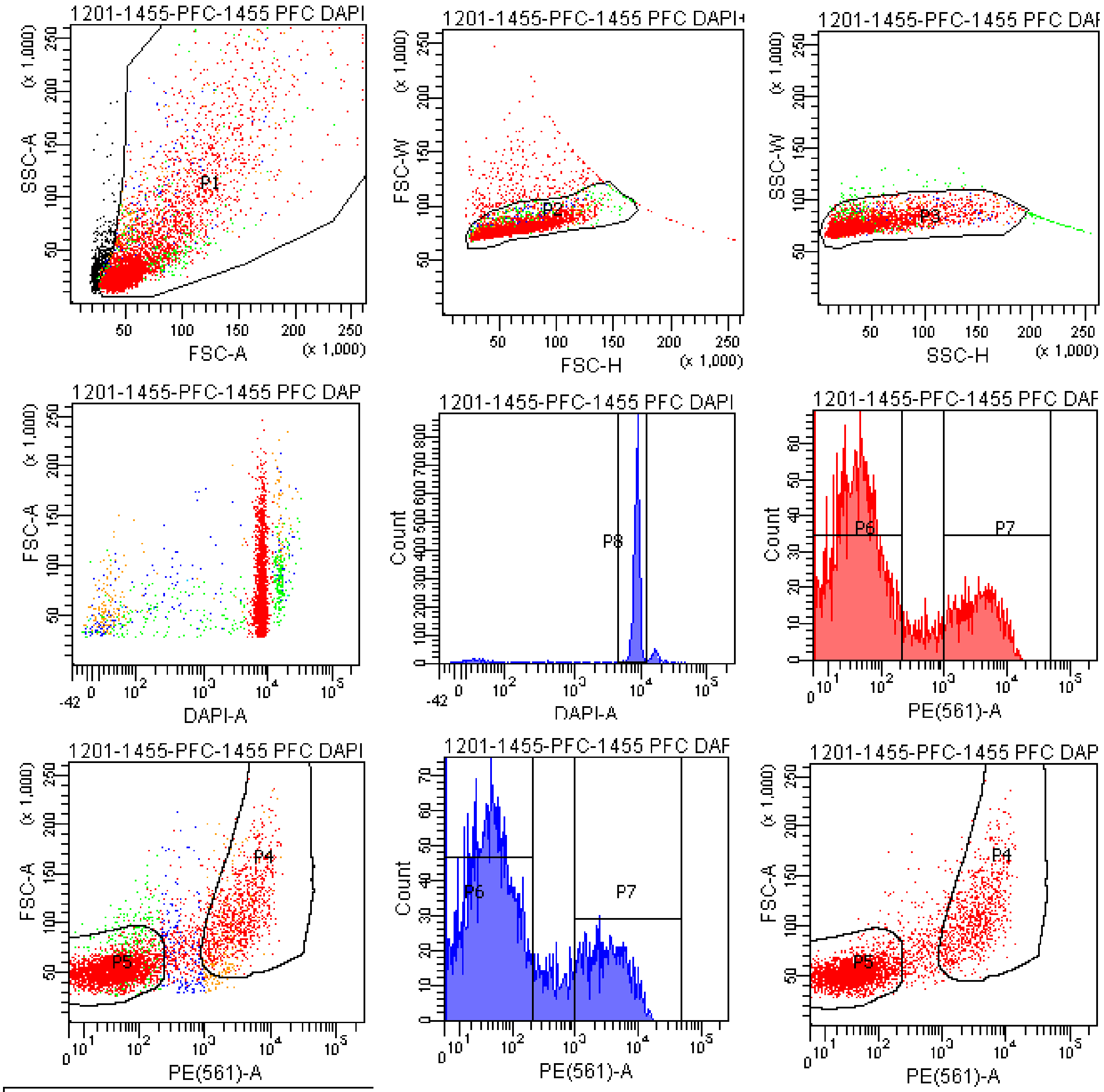


Tube: 1455 PFC DAPI+PE

| Population | #Events | %Parent | %Total |
|---|---|---|---|
| All Events | 5,852 | #### | 100.0 |
| P1 | 5,347 | 91.4 | 91.4 |
| P2 | 5,011 | 93.7 | 85.6 |
| P3 | 4,888 | 97.5 | 83.5 |
| P4 | 1,004 | 20.5 | 17.2 |
| P5 | 3,248 | 66.4 | 55.5 |
| P6 | 3,322 | 68.0 | 56.8 |
| P7 | 1,080 | 22.1 | 18.5 |
| P8 | 4,161 | 85.1 | 71.1 |
| P9 | 576 | 11.8 | 9.8 |

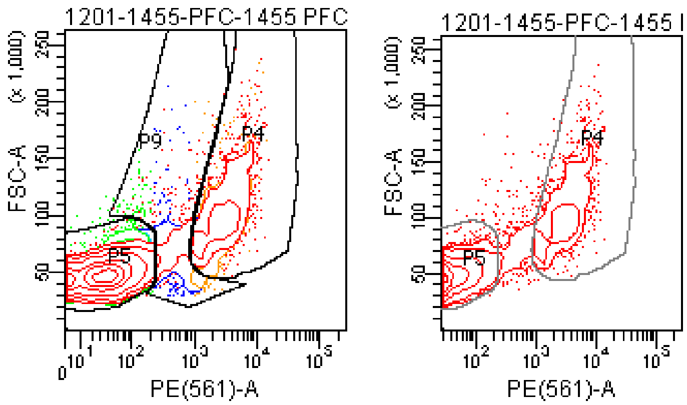

### FACSDiva Version 6.1.3

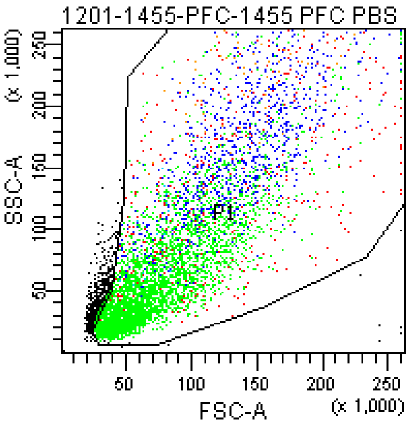


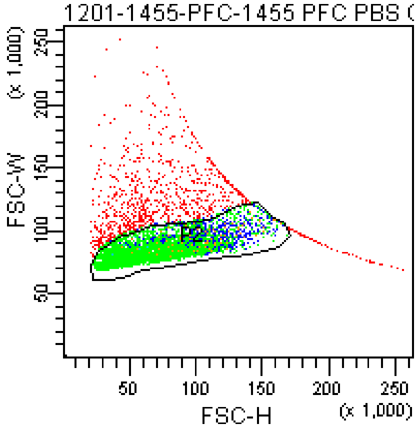


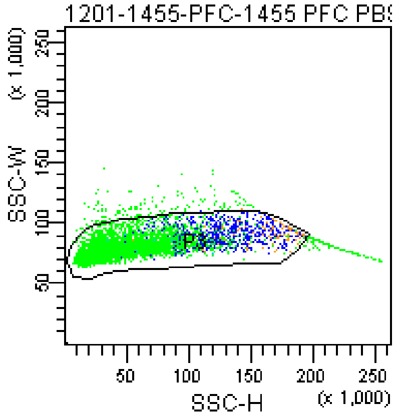


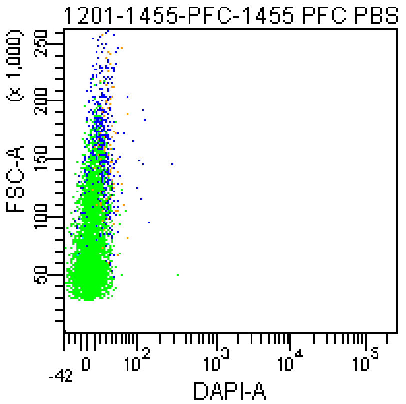


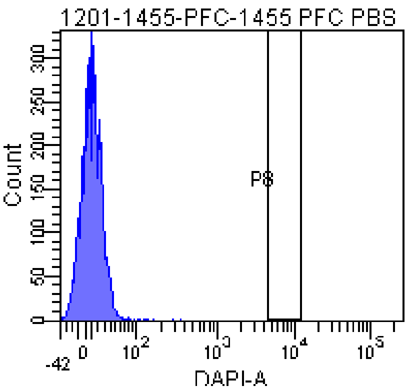


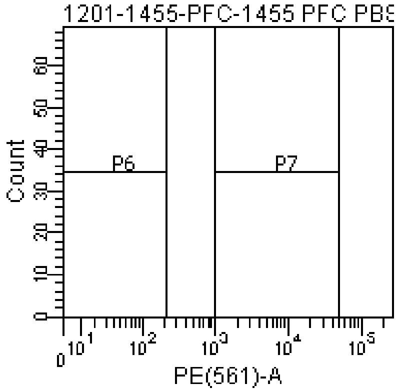


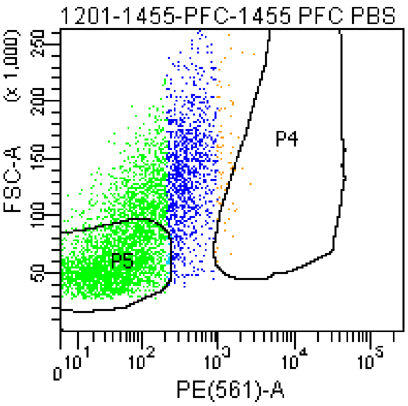


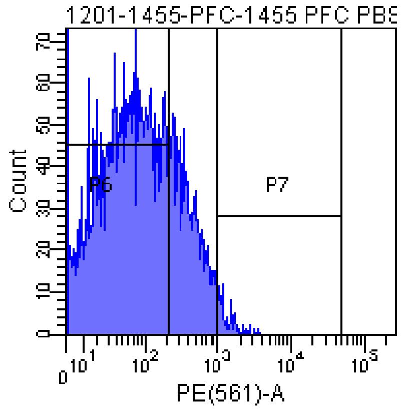


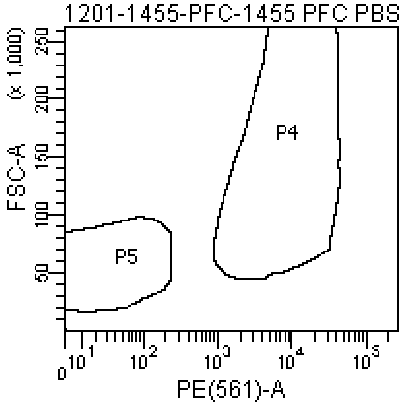


Tube: 1455 PFC PBS CONTROL

| Population | #Events | %Parent | %Total |
|---|---|---|---|
| All Events | 6,671 | #### | 100.0 |
| P1 | 6,051 | 90.7 | 90.7 |
| P2 | 5,205 | 86.0 | 78.0 |
| P3 | 4,854 | 93.3 | 72.8 |
| P4 | 19 | 0.4 | 0.3 |
| P5 | 2,842 | 58.5 | 42.6 |
| P6 | 3,699 | 76.2 | 55.4 |
| P7 | 77 | 1.6 | 1.2 |
| P8 | 0 | 0.0 | 0.0 |
| P9 | 1,694 | 34.9 | 25.4 |

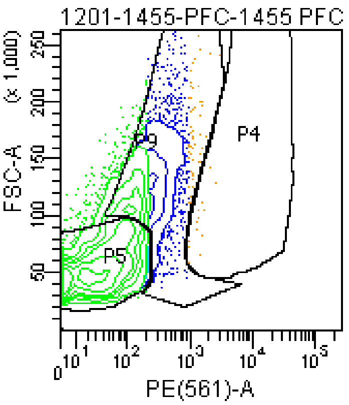


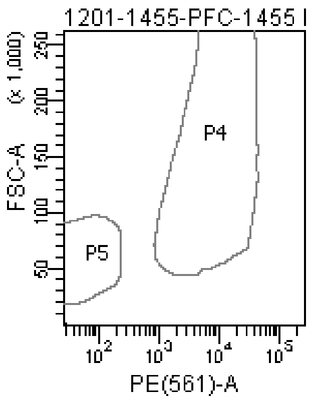

### FACSDiva Version 6.1.3

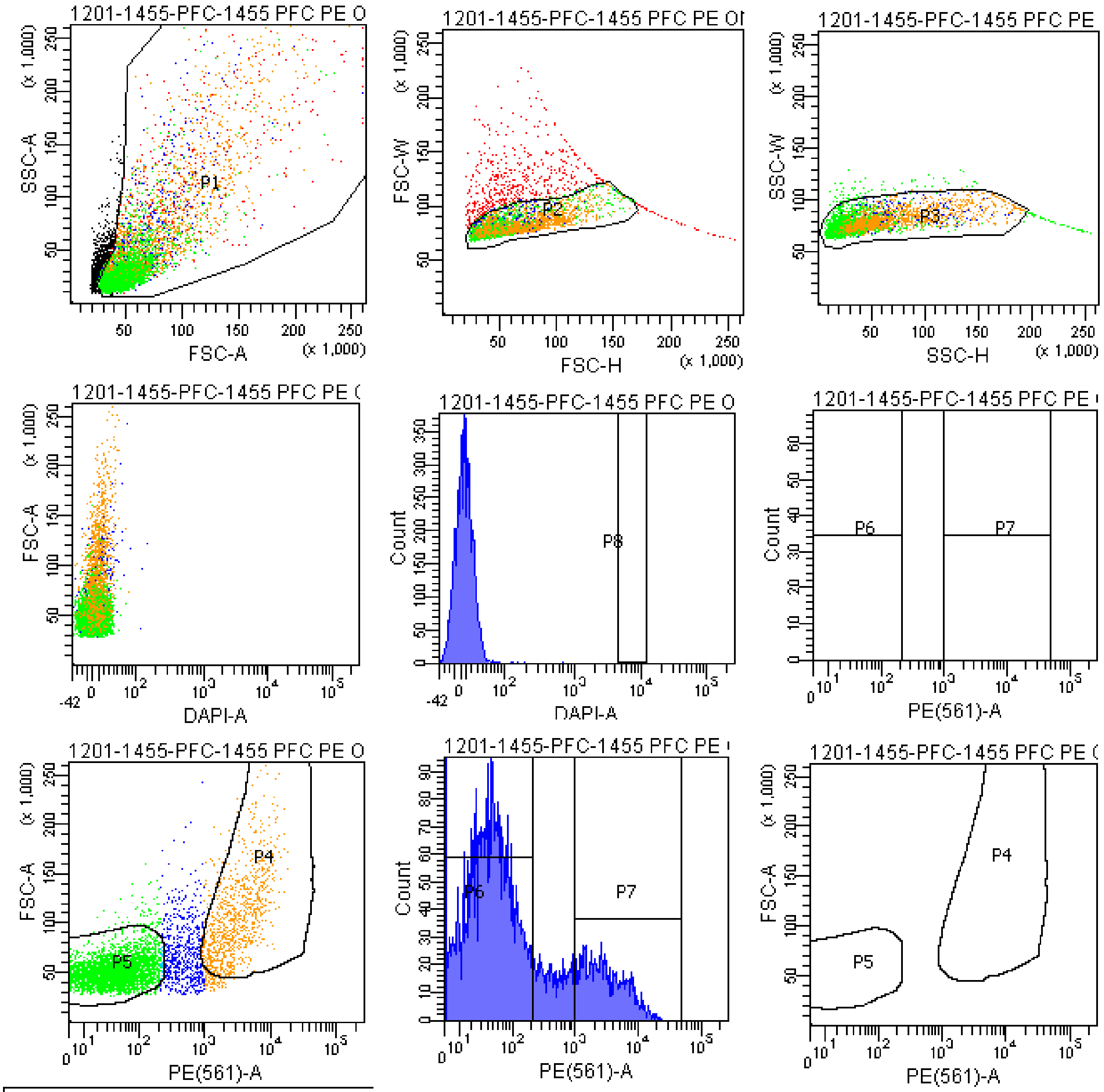


Tube: 1455 PFC PE ONLY CONTROL

| Population | #Events | %Parent | %Total |
|---|---|---|---|
| All Events | 7,367 | #### | 100.0 |
| P1 | 6,513 | 88.4 | 88.4 |
| P2 | 5,945 | 91.3 | 80.7 |
| P3 | 5,765 | 97.0 | 78.3 |
| P4 | 886 | 15.4 | 12.0 |
| P5 | 4,101 | 71.1 | 55.7 |
| P6 | 4,152 | 72.0 | 56.4 |
| P7 | 980 | 17.0 | 13.3 |
| P8 | 0 | 0.0 | 0.0 |
| P9 | 718 | 12.5 | 9.7 |

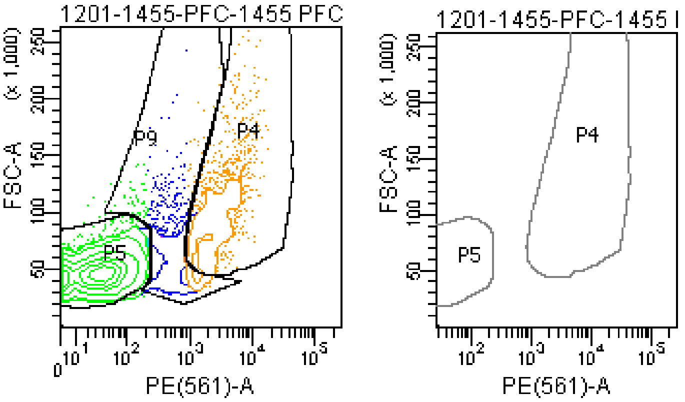

### FACSDiva Version 6.1.3

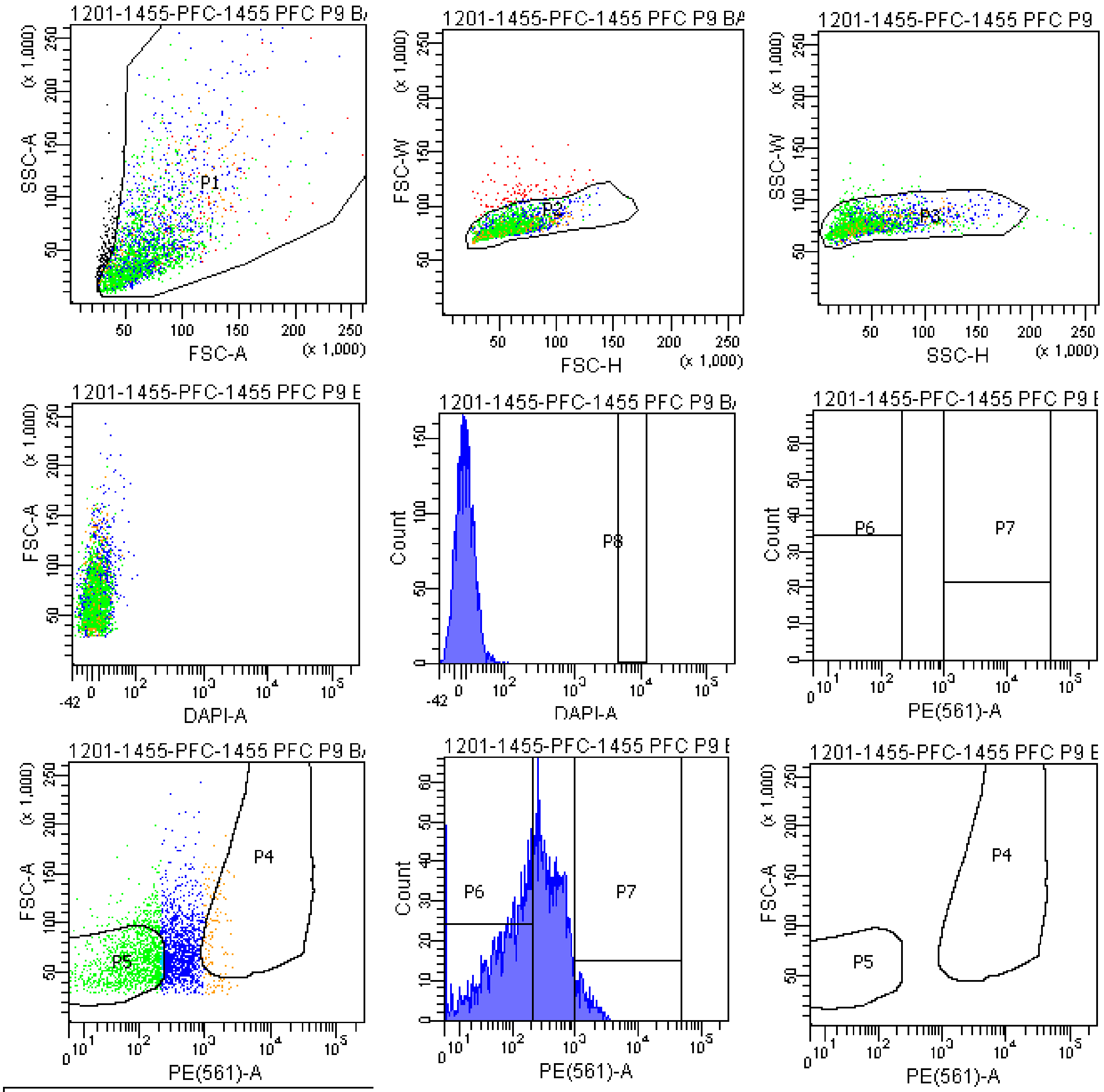


Tube: 1455 PFC P9 BACK

| Population | #Events | %Parent | %Total |
|---|---|---|---|
| All Events | 3,092 | #### | 100.0 |
| P1 | 2,914 | 94.2 | 94.2 |
| P2 | 2,811 | 96.5 | 90.9 |
| P3 | 2,769 | 98.5 | 89.6 |
| P4 | 88 | 3.2 | 2.8 |
| P5 | 1,229 | 44.4 | 39.7 |
| P6 | 1,321 | 47.7 | 42.7 |
| P7 | 137 | 4.9 | 4.4 |
| P8 | 0 | 0.0 | 0.0 |
| P9 | 1,363 | 49.2 | 44.1 |

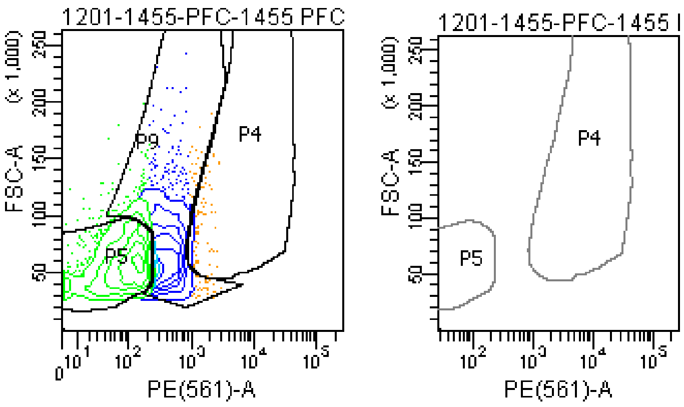

### FACSDiva Version 6.1.3

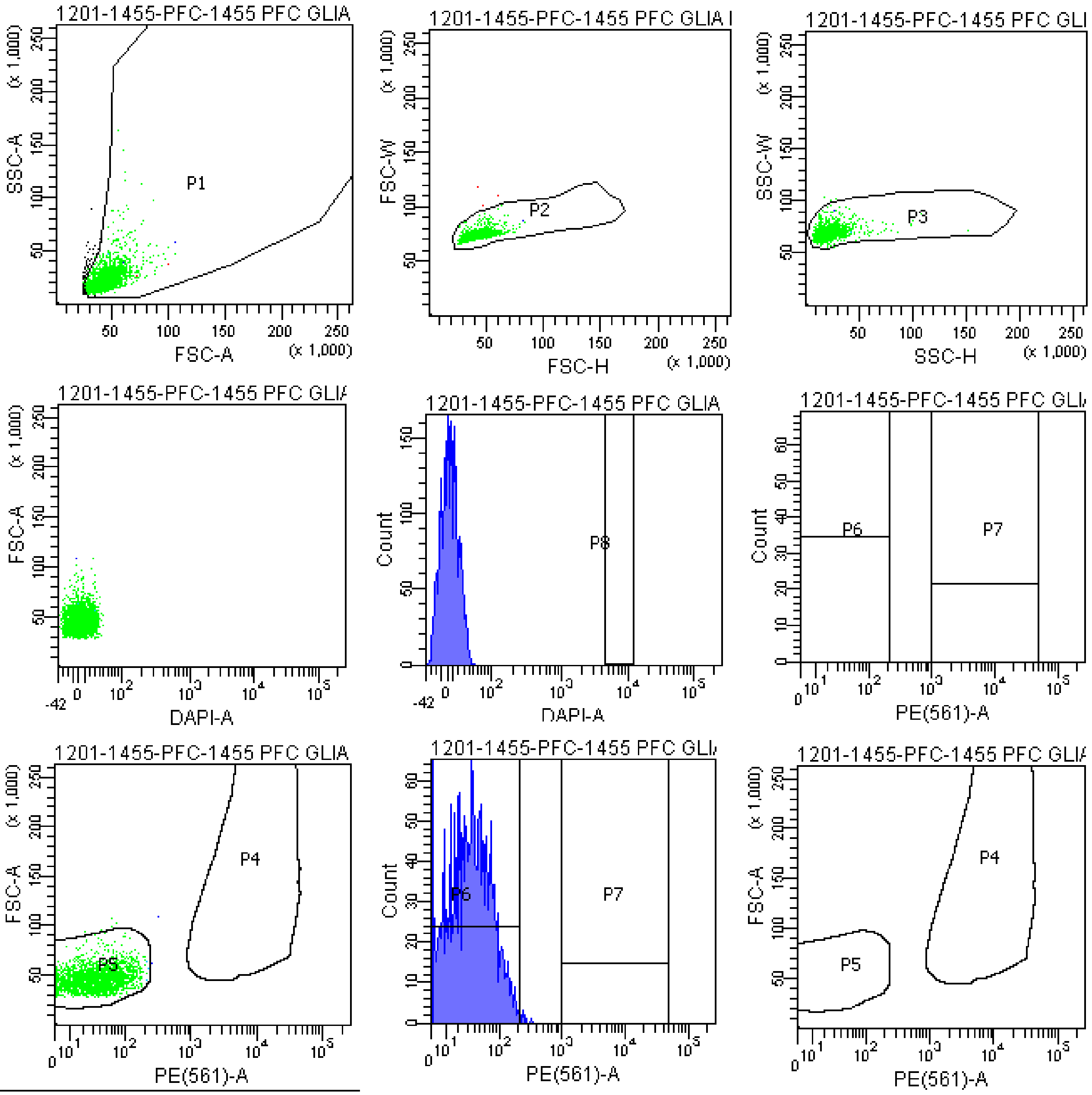


Tube: 1455 PFC GLIA BACK

| Population | #Events | %Parent | %Total |
|---|---|---|---|
| All Events | 2,833 | #### | 100.0 |
| P1 | 2,742 | 96.8 | 96.8 |
| P2 | 2,739 | 99.9 | 96.7 |
| P3 | 2,733 | 99.8 | 96.5 |
| P4 | 0 | 0.0 | 0.0 |
| P5 | 2,724 | 99.7 | 96.2 |
| P6 | 2,725 | 99.7 | 96.2 |
| P7 | 0 | 0.0 | 0.0 |
| P8 | 0 | 0.0 | 0.0 |
| P9 | 5 | 0.2 | 0.2 |

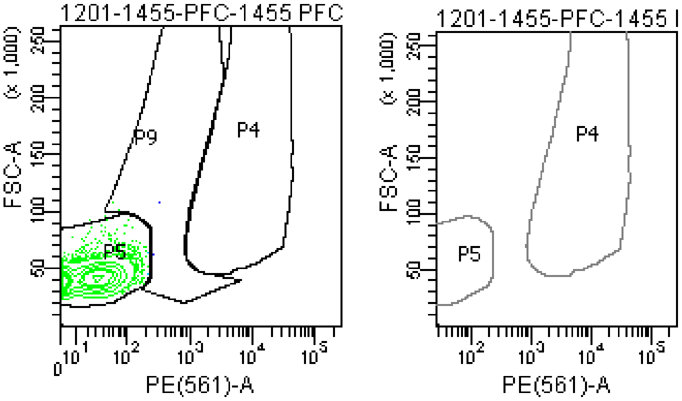

### FACSDiva Version 6.1.3

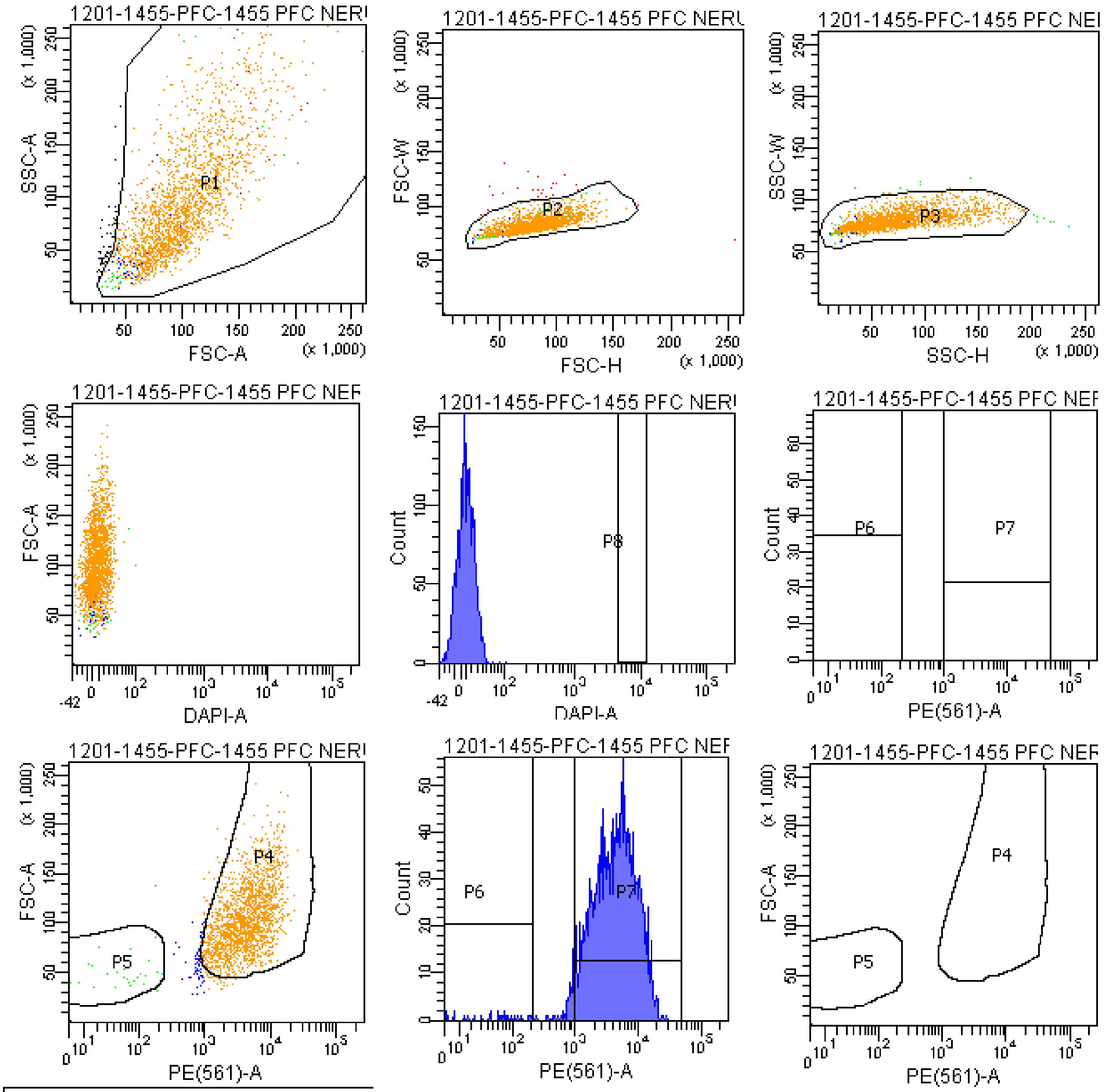


Tube: 1455 PFC NERUON BACK

| Population | #Events | %Parent | %Total |
|---|---|---|---|
| All Events | 2,232 | #### | 100.0 |
| P1 | 2,176 | 97.5 | 97.5 |
| P2 | 2,150 | 98.8 | 96.3 |
| P3 | 2,135 | 99.3 | 95.7 |
| P4 | 1,991 | 93.3 | 89.2 |
| P5 | 28 | 1.3 | 1.3 |
| P6 | 29 | 1.4 | 1.3 |
| P7 | 2,035 | 95.3 | 91.2 |
| P8 | 0 | 0.0 | 0.0 |
| P9 | 101 | 4.7 | 4.5 |

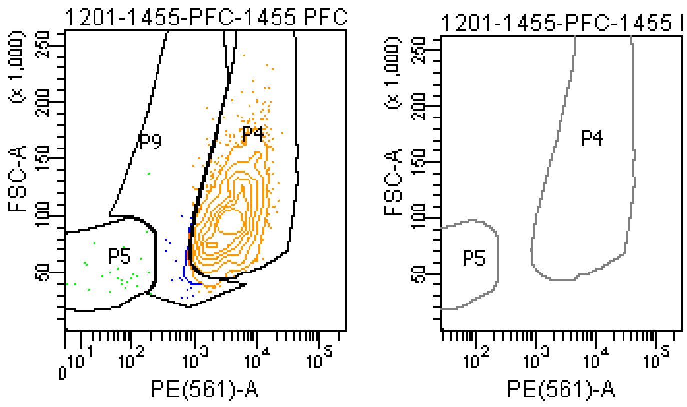
