## Appendix 2. Identification of ACC1-specific insertions and their zygosity for "Somatic LINE-1 retrotransposition in cortical neurons and non-brain tissues of Rett patients and healthy individuals"

To identify ACC1-specific insertions, at the first step, we identified 172 ACC1 non-reference germline L1Hs insertions using HAT-seq. Next, 64 of them were confirmed to be ACC1-specific by 3' junction PCR (3' PCR) analysis of gDNA from ACC1 and ACC2. At last, we performed full-length PCR to determine the zygosity of ACC1-specific insertions.

### Identification of ACC1-specific insertions

Validation primer sequences used for each candidate insertion and the predicted target amplicon sizes of 3' junction PCR products could be found in S2 Table. The IDs of ACC1-specific insertions were labeled at the top of agarose gel image. A: ACC1 gDNA; Z: ACC2 gDNA; M: 100 bp Plus DNA ladder; NTC: water. For details, see sections in the Materials and Methods "Positive control experiments" and "L1 3' PCR and full-length PCR validation".

### Zygosity of ACC1-specific insertions

Validation primer sequences used for each ACC1-specific insertion and the predicted empty site sizes of full-length PCR products could be found in S2 Table. The IDs of ACC1-specific insertions were labeled at the top of agarose gel image. M: 100 bp Plus DNA ladder; NTC: water. For details, see Materials and Methods "L1 3' PCR and full-length PCR validation". Several sites were tested by full-length PCR for multiple times to robustly determine their zygosity.

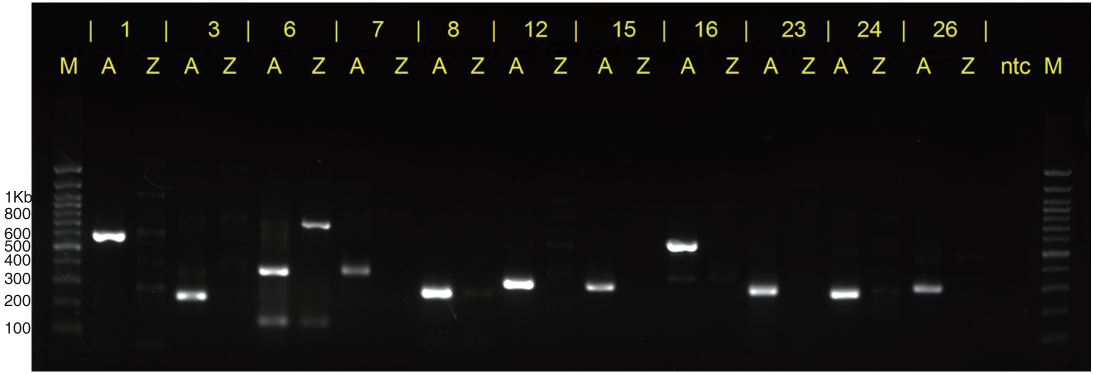

Identification of ACC1-specific insertions agarose gel image 1

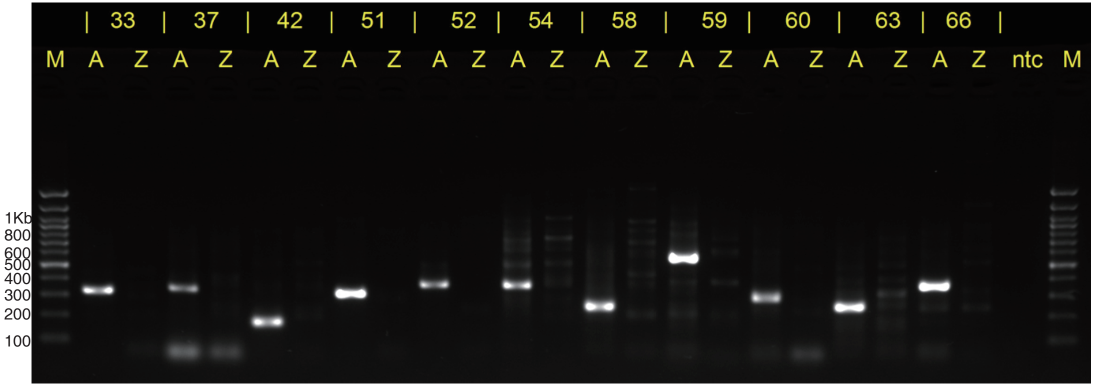

Identification of ACC1-specific insertions agarose gel image 2

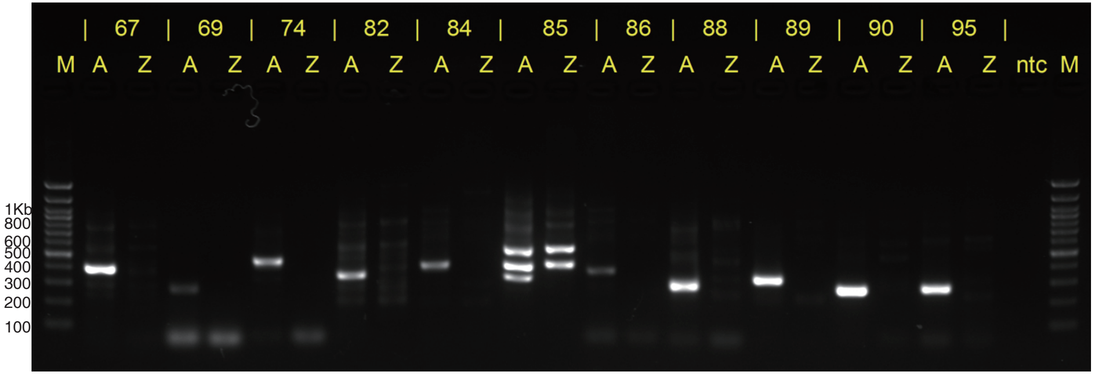

Identification of ACC1-specific insertions agarose gel image 3

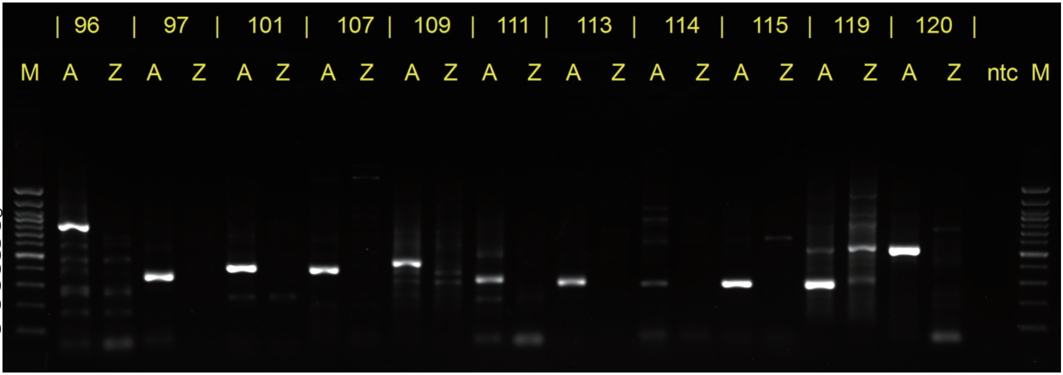

Identification of ACC1-specific insertions agarose gel image 4

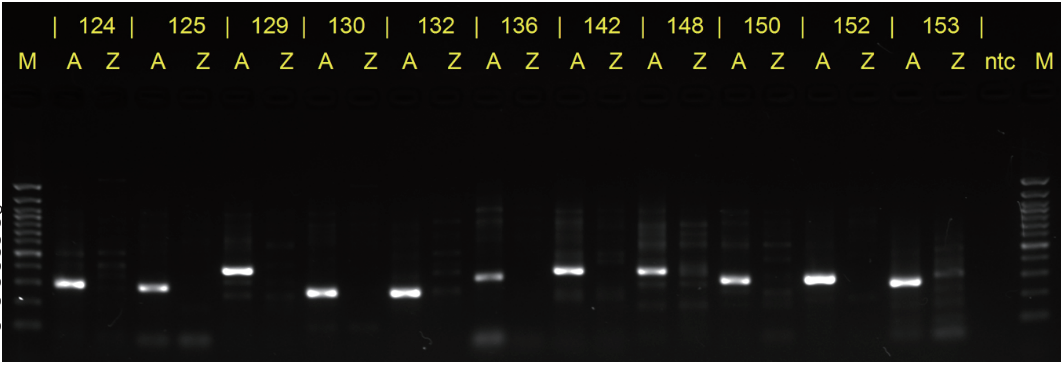

Identification of ACC1-specific insertions agarose gel image 5

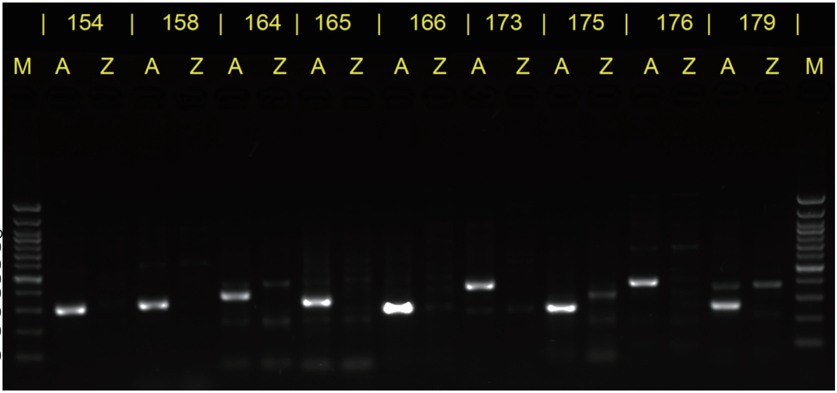

Identification of ACC1-specific insertions agarose gel image 6

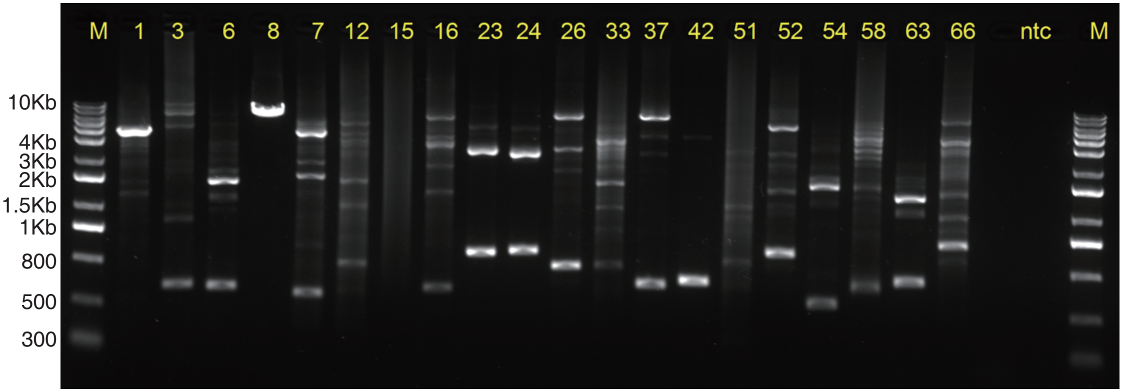


Zygosity of ACC1-specific insertions agarose gel image 1

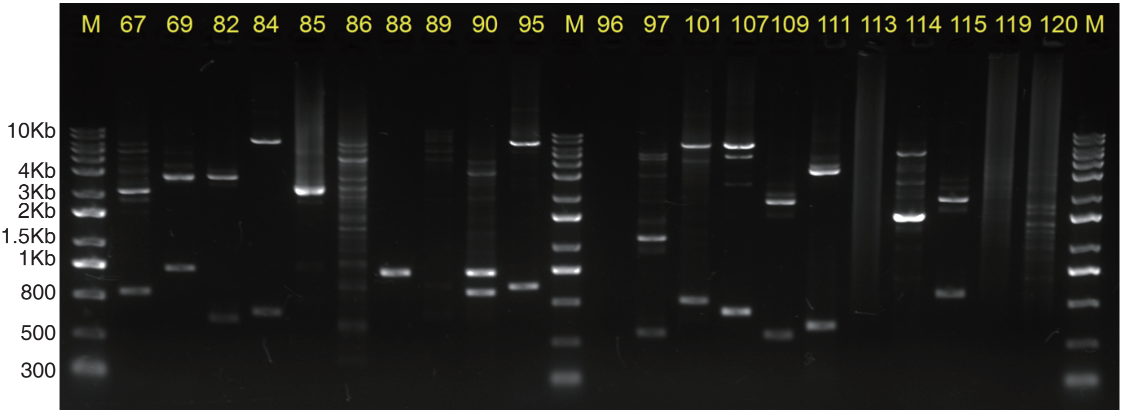


Zygosity of ACC1-specific insertions agarose gel image 2

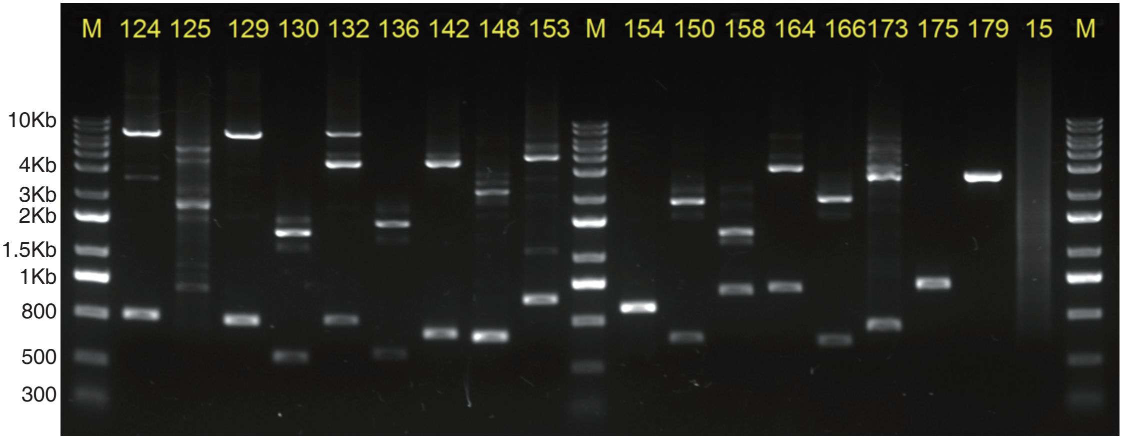


Zygosity of ACC1-specific insertions agarose gel image 3

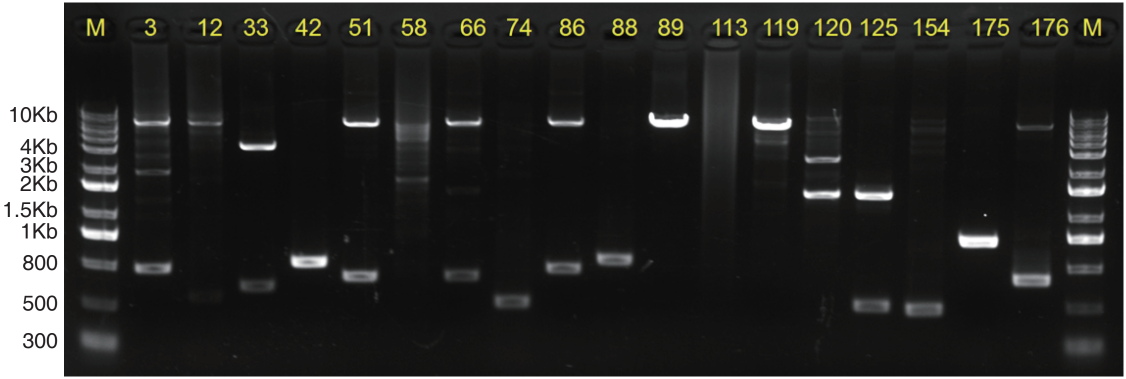


Zygosity of ACC1-specific insertions agarose gel image 4

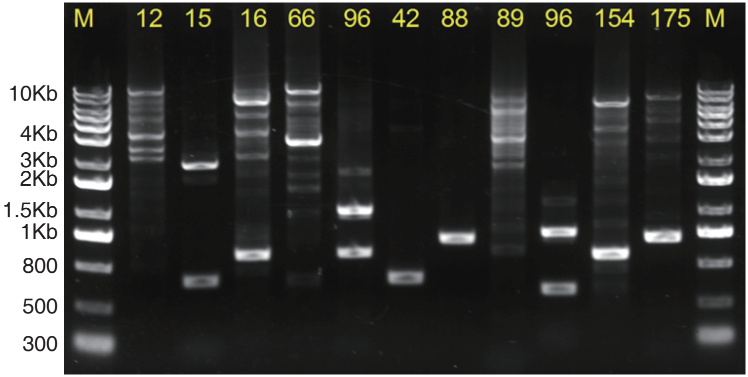


Zygosity of ACC1-specific insertions agarose gel image 5

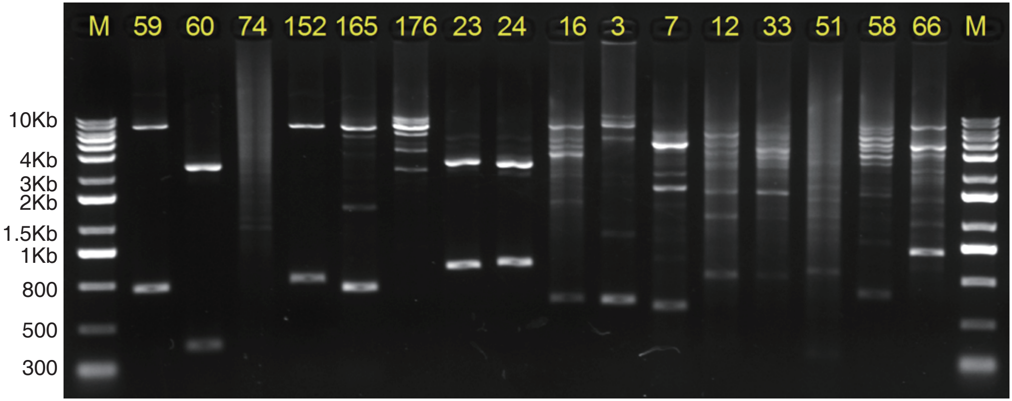


Zygosity of ACC1-specific insertions agarose gel image 6
