## Appendix 3. Detection of somatic insertions in positive control experiments for "Somatic LINE-1 retrotransposition in cortical neurons and non-brain tissues of Rett patients and healthy individuals"

**Appendix 3. Detection of somatic L1Hs insertions at low levels in spike-in HAT-seq libraries.**

To benchmark the performance of HAT-seq for detecting somatic L1Hs insertions, we experimentally generated a series of positive control samples with insertions at different frequencies by mixing the genomic DNA (gDNA) extracted from the blood samples of two unrelated adults, ACC1 and ACC2. 64 ACC1-specific insertions were identified by 3' junction PCR and served as positive controls. Three HAT-seq libraries were generated from samples consisting of ACC2 gDNA spiked with 1%, 0.1%, or 0.01% of ACC1 gDNA. Notably, in 0.01% spike-in library, 10 out of 64 insertions were detected. Our data showed that, with about 3,000 cells as input, HAT-seq was able to detect somatic insertion events present in a single cell.

<u>IGV images of low-frequency ACC1-specific insertions in spike-in libraries</u>

Each IGV image showed both merged contigs and unassembled reads that were uniquely mapped to an ACC1-specific insertion. The IDs of ACC1-specific insertions were labeled at the top of images. Supporting signal counts for each insertion were indicated by black arrows and calculated by counting the unique start positions. The depth of each signal in spike-in libraries (the number of PCR duplicates for each signal with unique start position) could be found in S2 Table. The depth of coverage for insertions were shown by gray histogram tracks. Gene symbols and transcription orientation were shown in the RefSeq annotation track at the bottom of image. Mismatches and small indels were labeled in distinct colors. 1%, 0.1%, and 0.01% contig: poly-T trimmed merged contigs in spike-in libraries. 1%, 01%, and 0.01% PE: poly-T trimmed unassembled reads (Read 1) in spike-in libraries.

An overview of ten ACC1-specific insertions that were detected in 0.01 spike-in HAT-seq library.

| ID | Chr | Start | End | Peak Strand | Merged contigs | | | Unassembled read pairs | | |
|---|---|---|---|---|---|---|---|---|---|---|
| | | | | | 1% spike-in library | 0.1% spike-in library | 0.01% spike-in library | 1% spike-in library | 0.1% spike-in library | 0.01% spike-in library |
| 66 | 4 | 54368037 | 54368301 | + | 4,1,1 | " | 5,1 | 4,3,2,1 | 5,2 | 7 |
| 69 | 4 | 98312367 | 98312640 | + | 4,3,3 | " | " | 5,3 | 1 | 3 |
| 89 | 7 | 90331640 | 90331920 | + | 6,4,2,2,2,1,1,1,1,1 | 10,7 | " | 6,4,4,2,2,2,2,1,1 | 4,1 | 5 |
| 115 | 13 | 61734481 | 61734810 | - | 9,4,4,2,1,1 | " | 4 | 3,2 | " | " |
| 124 | 2 | 63165217 | 63165513 | - | 4,2,2,2,2 | 2 | " | 4,2,1,1,1,1 | " | 1 |
| 129 | 2 | 164288500 | 164288750 | - | 7,6,4,4,3,3,2,1 | " | 7 | 5,3 | 7 | " |
| 132 | 21 | 29069165 | 29069552 | - | 5,3,2,1,1 | 6,2 | 2 | 5,4,2,1,1 | 11 | " |
| 153 | 5 | 155619860 | 155620166 | - | 4,4,4,3,3,2 | 4 | 10 | 11,8,7,6,5,5,5,4,3,3,2 | 6,1,1 | " |
| 154 | 6 | 13191011 | 13191296 | - | 8,6,5,4,2,2,1,1 | 13,1,1,1,1,1,1 | 3,3,2,1 | 3,1 | " | " |
| 158 | 6 | 66731906 | 66732173 | - | 2 | " | 10 | 4,2,1 | 2 | " |

Footnotes:

The numbers separated by comma were the depth of each signal in spike-in libraries (the number of PCR duplicates for each signal with unique start position).

### ACC1-specific insertion (ACC1_66 at chr4:54368301) in spike-in libraries

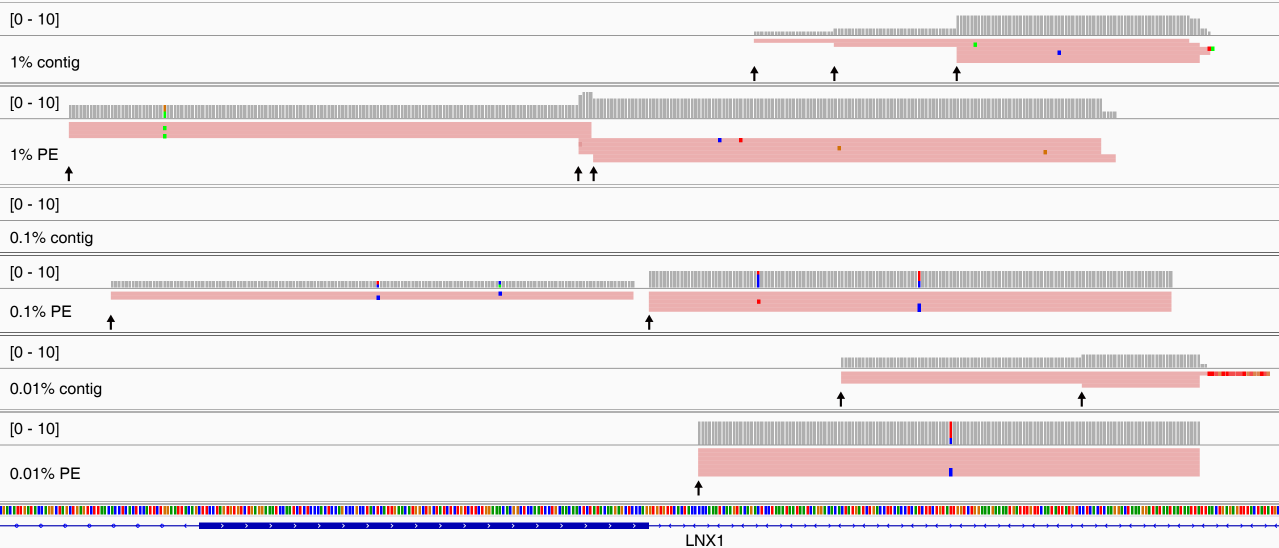

### ACC1-specific insertion (ACC1_69 at chr4:98312640) in spike-in libraries

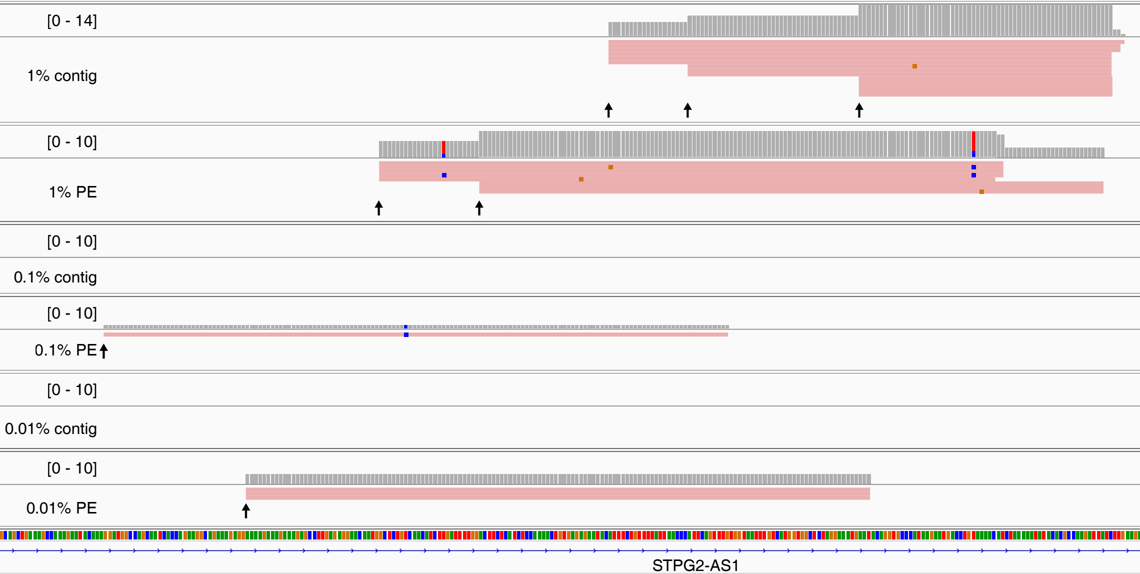

### ACC1-specific insertion (ACC1_89 at chr7:90331920) in spike-in libraries

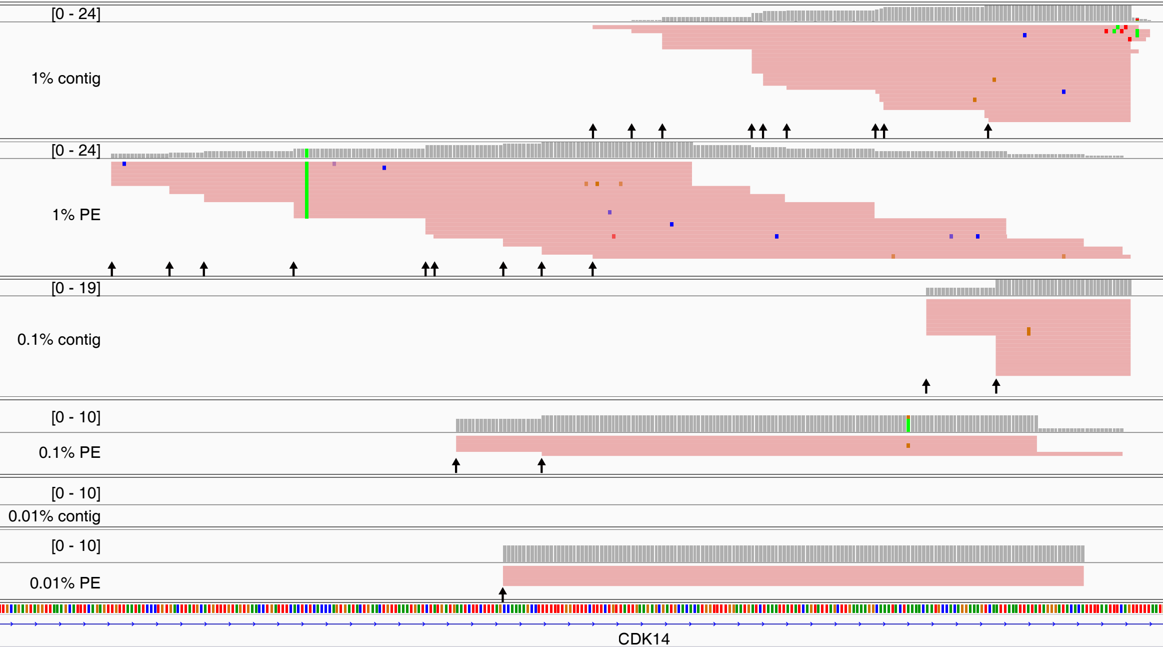

### ACC1-specific insertion (ACC1_115 at chr13:61734481) in spike-in libraries

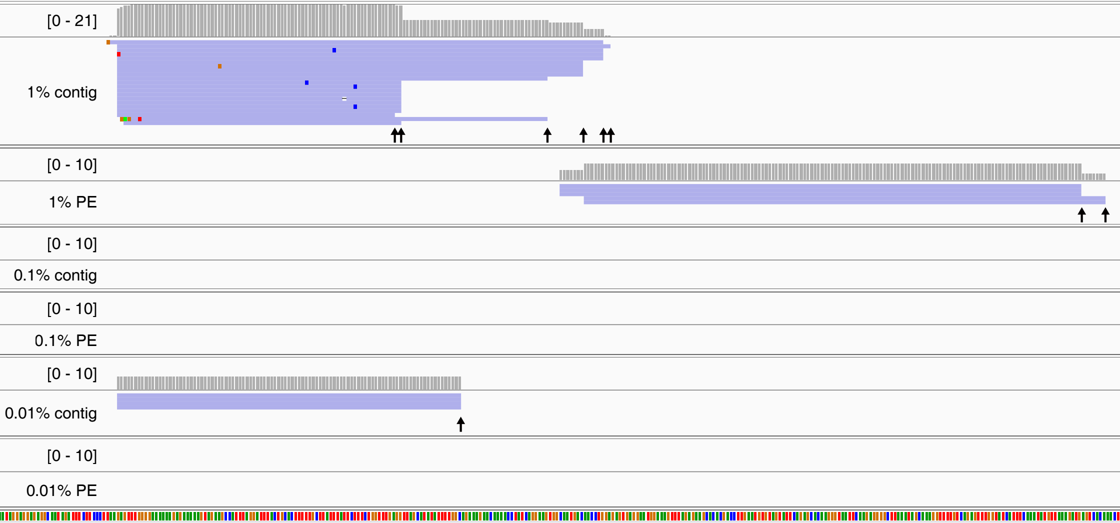

### ACC1-specific insertion (ACC1_124 at chr2:63165217) in spike-in libraries

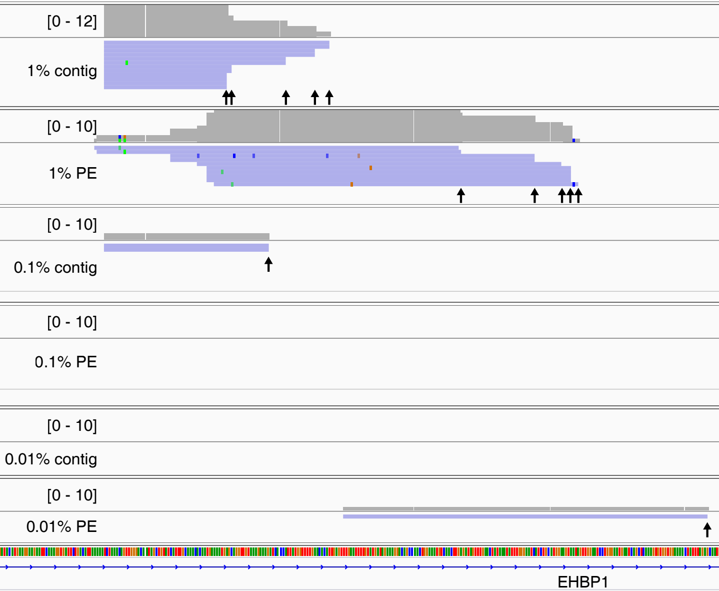

ACC1-specific insertion (ACC1_129 at chr2:164288500) in spike-in libraries

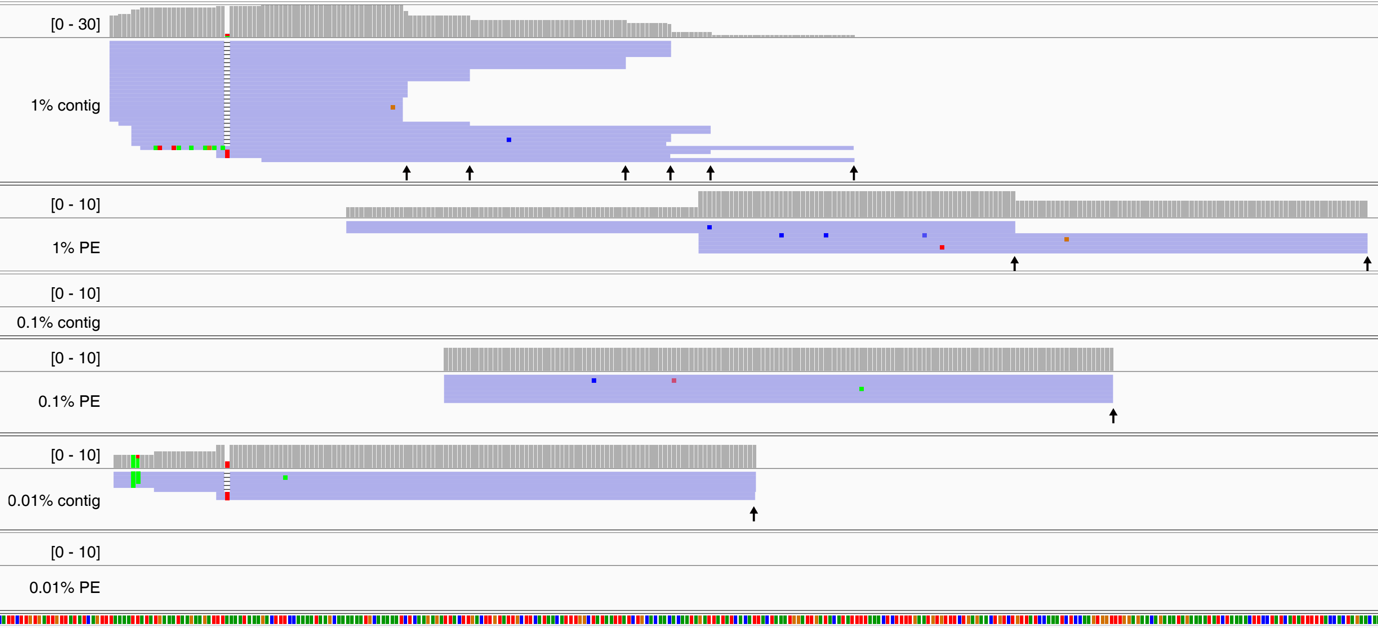

### ACC1-specific insertion (ACC1_132 at chr21:29069165) in spike-in libraries

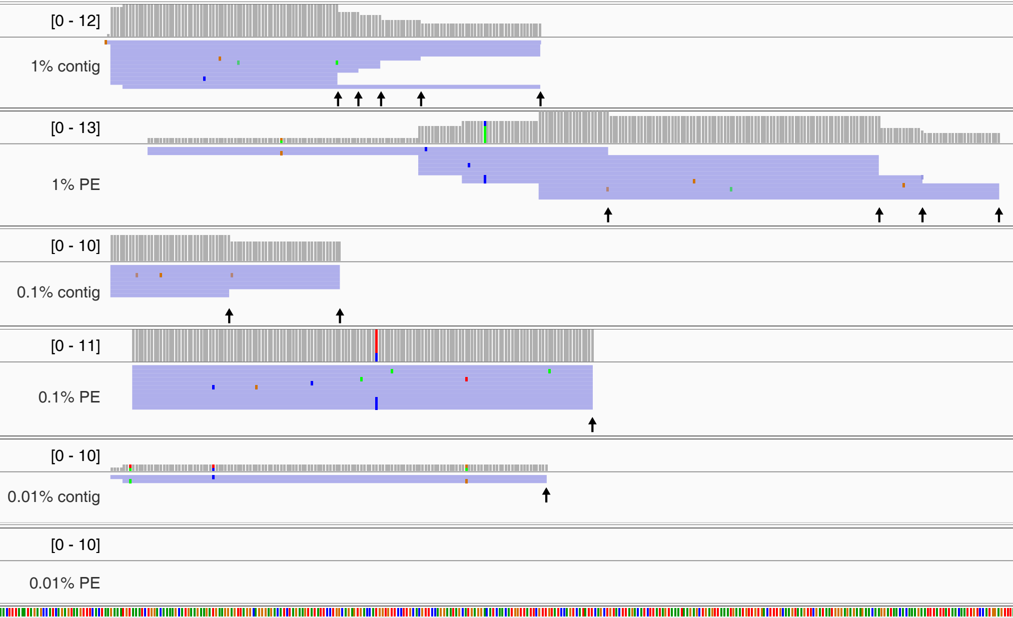

### ACC1-specific insertion (ACC1_153 at chr5:155619860) in spike-in libraries

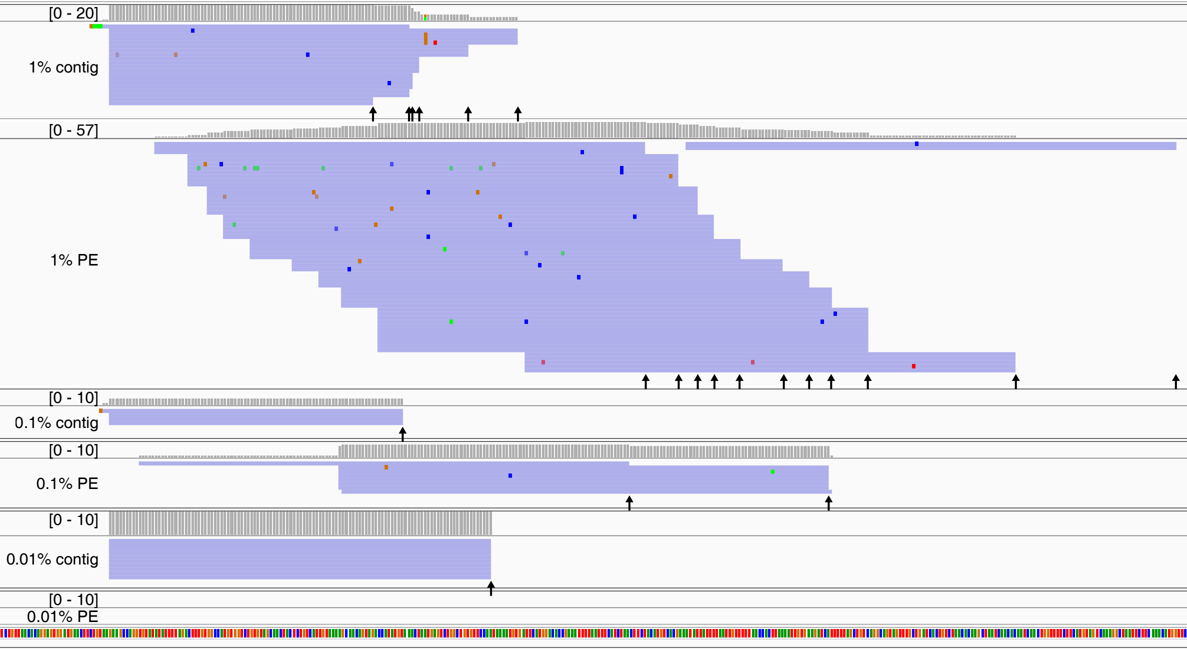

### ACC1-specific insertion (ACC1_154 at chr6:13191011) in spike-in libraries

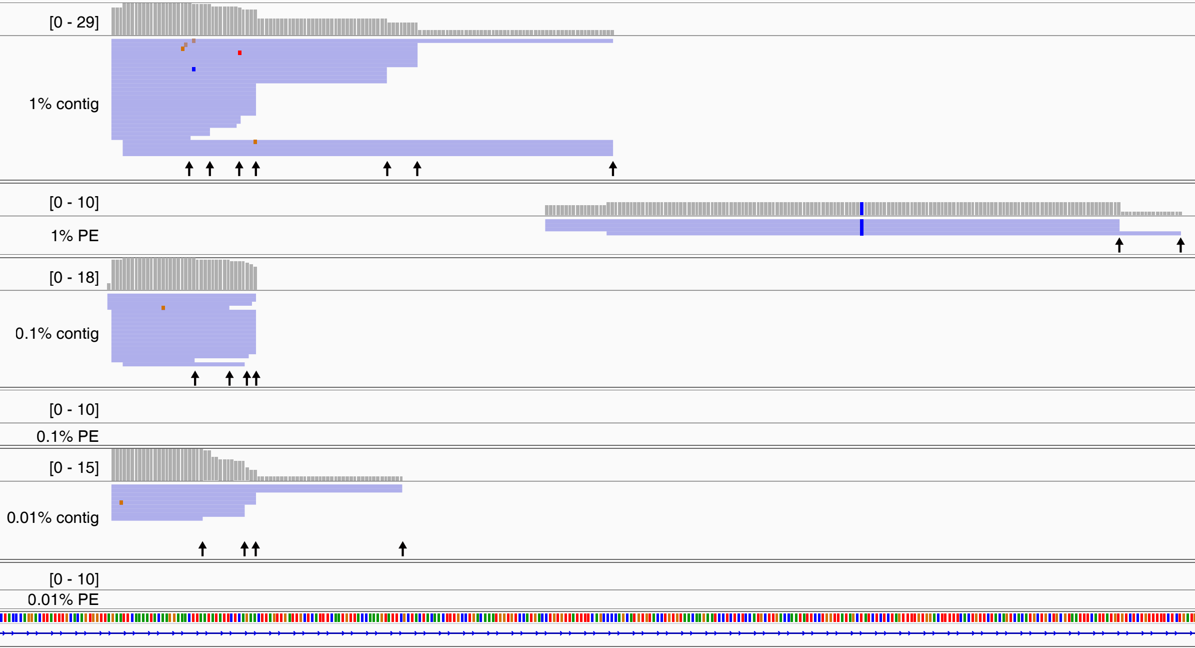

ACC1-specific insertion (ACC1_158 at chr6:66731906) in spike-in libraries

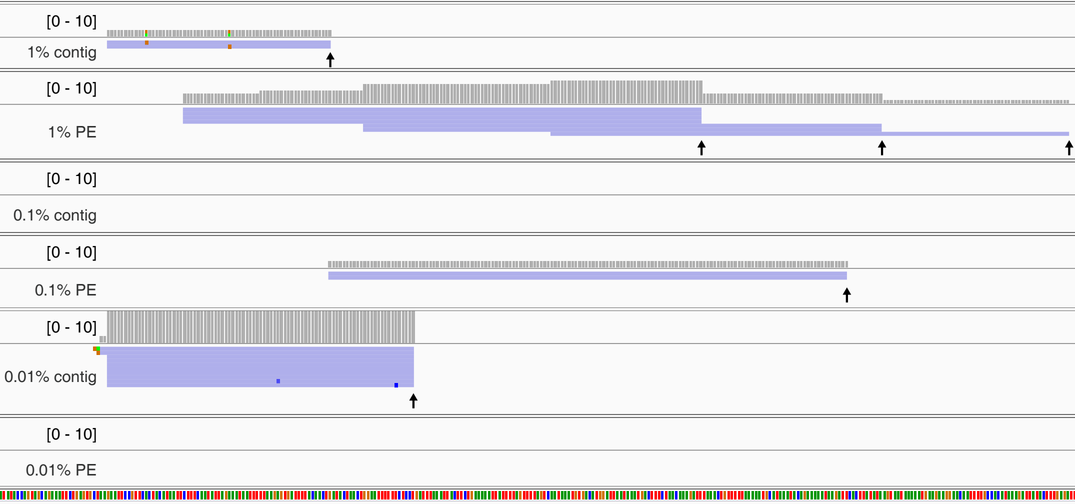
