## Appendix 4. Benchmarking PCR validation assays for low-frequency somatic insertions for "Somatic LINE-1 retrotransposition in cortical neurons and non-brain tissues of Rett patients and healthy individuals"

To fully characterized the TPRT hallmarks of putative somatic L1Hs insertion, Sanger sequencing of both 3' and 5' L1-genome junctions were required to precisely resolve the terminal site duplicate (TSD), the L1 endonuclease cleavage site (L1 EN motif), and poly-A tail length, yielding a high-confidence somatic L1Hs insertion. However, validating such low-frequency somatic insertions, in particular when one of the primers was complementary to numerous homologous sequences in the genome was challenging.

We systematically assessed the detection limit of several PCR methods, including full-length PCR, 3' junction PCR, 3' junction nested PCR, 5' junction PCR and 5' junction nested PCR using a series of 10%, 1%, 0.1%, and 0.01% spike-in DNA. Our positive control results showed that both 3' and 5' junction nested PCR could detect and yield visible amplicon bands for rare somatic L1Hs insertions whose mosaicism > 0.1%, whereas other conventional PCR assays could detect insertions whose mosaicism > 1%.

Detection limit of full-length PCR

As shown in the figure below, the detection limit of full-length PCR was 1% mosaicism. Because most of somatic retrotransposition events were 5' highly truncated, we selected three heterozygous, truncated ACC1-specific L1Hs insertions to evaluate. The range of L1Hs insertion length was between 800 bp (ACC1_97) and 1,500 bp (ACC1_15). A mixed-DNA series containing 10%, 1%, and 0.1% ACC1 gDNA were prepared using ACC1 and ACC2 (labeled as "ZBX") gDNA. ACC1 gDNA, ACC2 gDNA, and mixed DNA series were used as the template for full-length PCR. The upper band in full-length PCR of the heterozygous insertion was the allele with the L1Hs insertion (filled site), and the lower band was the allele with no insertion (empty site). The IDs of ACC1-specific insertions were labeled at the top of agarose gel image. ZBX: ACC2 gDNA; ACC1: ACC1 gDNA; 100M: 100 bp Plus DNA ladder; NTC: water. For details, see sections in the Materials and Methods "Positive control experiments" and "L1 3' PCR and full-length PCR validation".

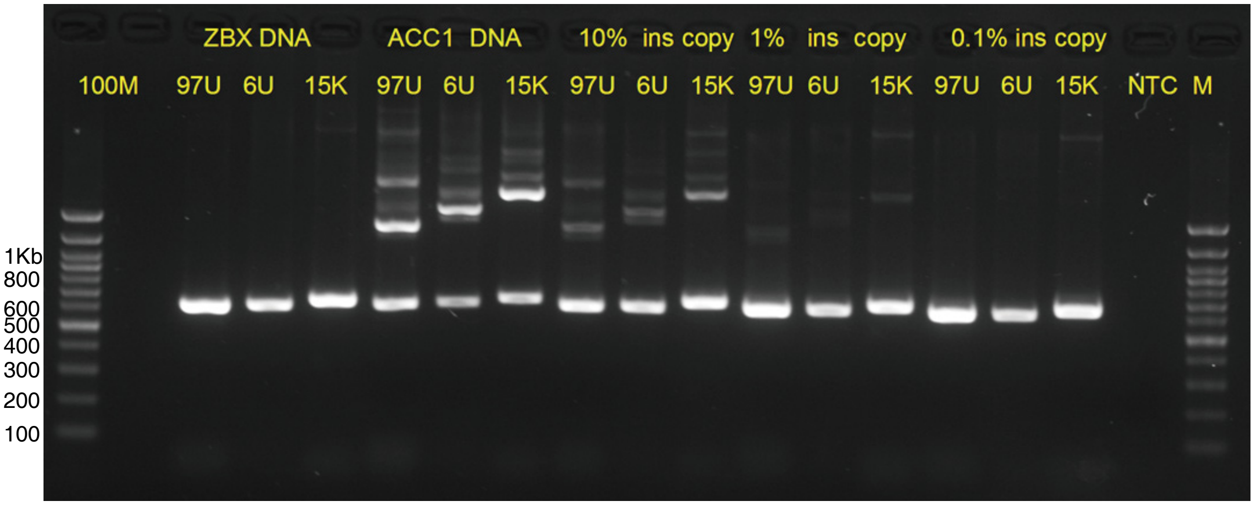

The detection limit of full-length PCR

Detection limit of 3' junction PCR and 3' junction nested PCR

As shown in the figures in the following pages, our results showed that 3' nested PCR could detect and yield visible amplicon bands for rare somatic L1Hs insertions whose mosaicism (percentage of cells) > 0.1%, whereas the conventional 3' PCR assay could detect insertions whose mosaicism > 1%. In addition, two previous studies demonstrated that 3' junction nested PCR could detect low-frequency insertions. Ewing et al. reported that they detected 1 copy in 10 cells (very faint band) with conventional PCR, and 1 copy per 1,000 cells with semi-nested PCR (Ewing et al., 2015). Using digital nested 3' PCR, Evrony et al. successfully amplified the 3' junction and measured the length of poly-A tail of somatic insertion whose mosaicism was ~0.5% (Evrony et al., 2015). Based on these results, we performed 3' junction nested PCR by adapting a modified version of digital nested 3' PCR (Evrony et al., 2015) that focused exclusively on clonal somatic insertions with three or more supporting signals, whose mosaicism were at least 0.1%.

First, we selected three heterozygous ACC1-specific L1Hs insertions to evaluate the detection limit of conventional 3' junction PCR. A mixed-DNA series containing 10%, 1%, and 0.1% ACC1 gDNA were prepared using ACC1 and ACC2 (labeled as "ZBX") gDNA. The IDs of ACC1-specific insertions were labeled at the top of agarose gel image. ZBX: ACC2 gDNA; ACC1: ACC1 gDNA; 100M: 100 bp Plus DNA ladder; NTC: water. For details, see sections in the Materials and Methods "Positive control experiments" and "L1 3' PCR and full-length PCR validation".

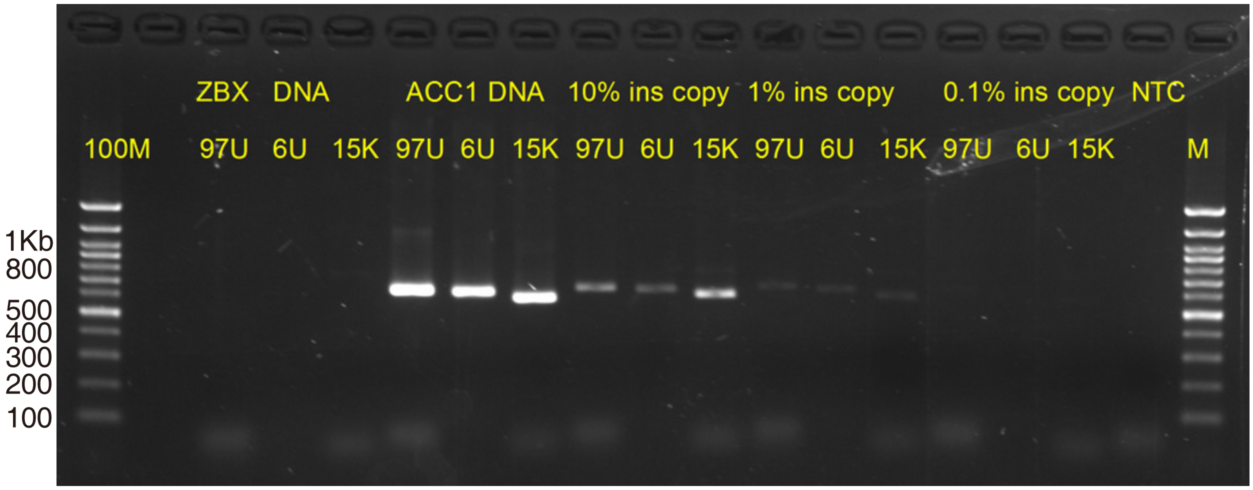

The detection limit of 3’ junction PCR using KAPA2G Robust HotStart ReadyMix

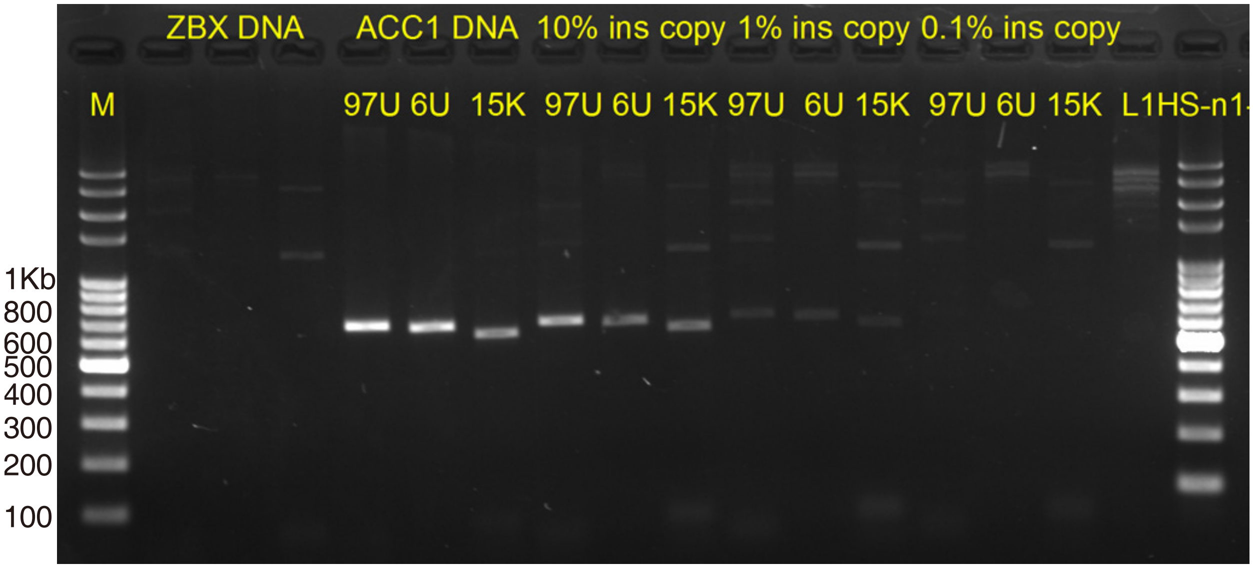

The detection limit of 3’ junction PCR using PrimeSTAR GXL DNA polymerase

Our positive spike-in experiments showed that the 3’ junction PCR can detect and yield visible amplicon bands for somatic L1Hs insertions whose mosaicism (percentage of cells) > 1%.

Second, we keep using these three heterozygous ACC1-specific L1Hs insertions to evaluate the detection limit of 3' junction nested PCR. A mixed-DNA series containing 1% and 0.1% ACC1 gDNA were prepared using ACC1 and ACC2 gDNA. The IDs of ACC1-specific insertions were labeled at the top of agarose gel image. The three lanes on the left were amplified using 1% spike-in gDNA, with 0.1% spike-in gDNA for three lanes on the right. M: 100 bp Plus DNA ladder. For details, see sections in the Materials and Methods "Positive control experiments" and "L1 3' digital nested PCR validation".

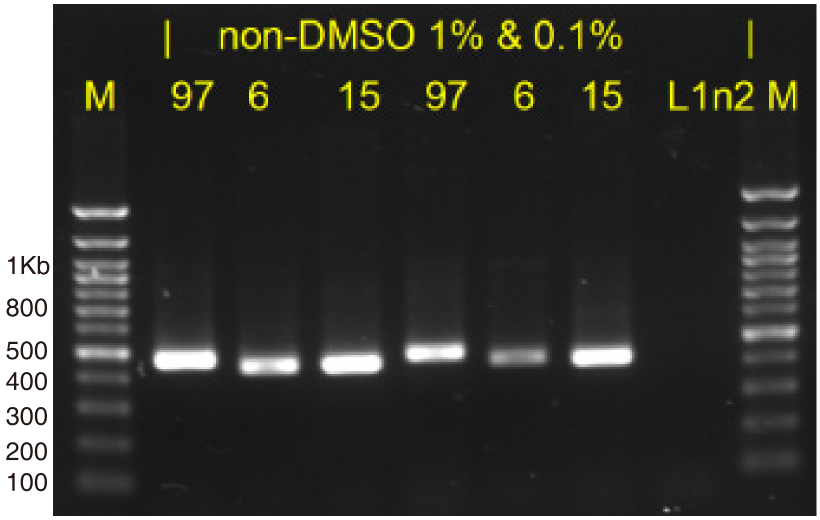

The detection limit of 3' junction nested PCR using 1% and 0.1% spike-in ACC1 gDNA

Our positive spike-in experiments showed that the 3' junction nested PCR can robustly detect and yield visible amplicon bands for rare somatic L1Hs insertions whose mosaicism (percentage of cells) > 0.1%. We then asked if the detection limit of 3' junction nested PCR could reach to 0.01% mosaicism.

A mixed-DNA series containing 0.01% ACC1 gDNA were prepared using ACC1 and ACC2 gDNA. The IDs of ACC1-specific insertions were labeled at the top of agarose gel image. The PCR assay for each insertion was repeated twice. R1: PCR products from the Round 1 PCR. Yellow arrow indicated the amplicon band with the targeted size. M: 100 bp Plus DNA ladder.

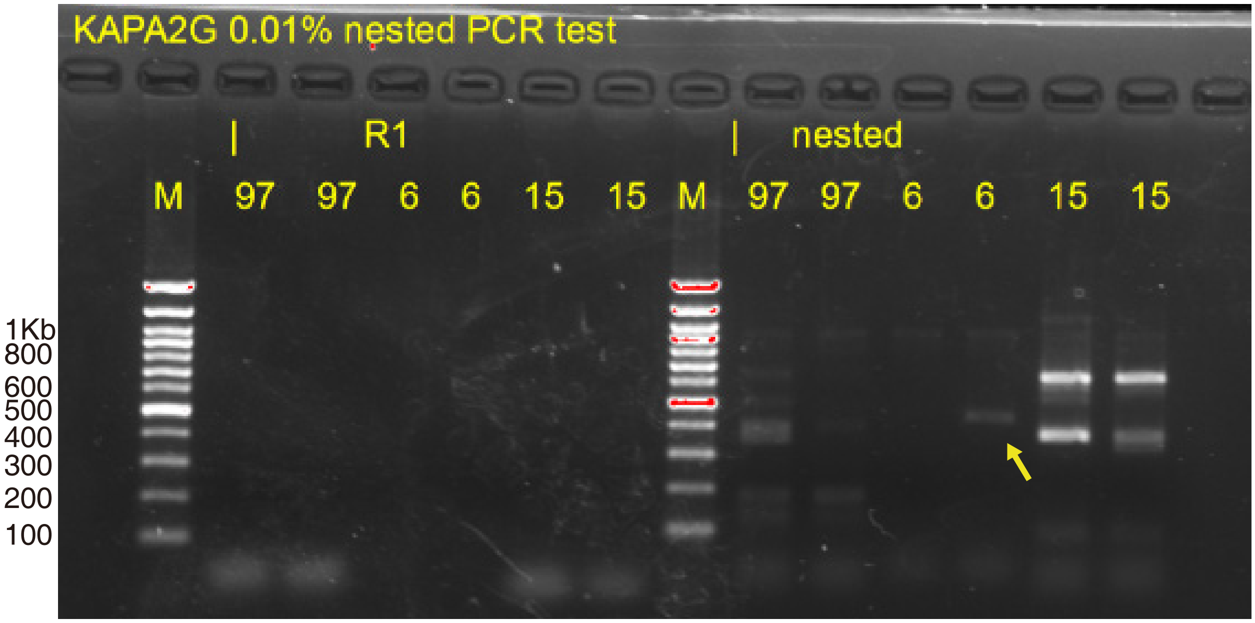

The detection limit of 3' junction conventional and nested PCR using 0.01% spike-in ACC1

Our results showed that the detection limit of 3' junction nested PCR could reach to 0.01% mosaicism. However, due to the low-frequency of such insertions, only some of them could be re-sampled by chance and amplified with a faint band. In summary, we concluded that 3' junction nested PCR could robustly amplify insertions with mosaicism > 0.1%. In this study, our validation focused exclusively on clonal somatic insertions whose mosaicism were at least 0.1%.

L1 5’ junction nested PCR validation

Sequence information of L1-genome 3’ junction enable us to determine the genomic location, insertional orientation, and poly-A tail of somatic insertions. Further leveraging the 5’ junction sequence information enabled us to precisely resolve additional hallmark features, including the TSD and the EN motif of integration event. However, there were more technical challenges for amplification of 5’ junction than 3’ junction PCR assays. First, many somatic L1Hs insertions harbor 5’ inversion, which making the 5’ junction PCR assay more complicated. Previous full-length PCR validation on MDA amplified single cell genomic DNA showed that 2 out of 3 somatic L1Hs insertions contained an 5’ inversion (Evrony et al., 2015). Second, unlike the primers for 3’ junction PCR that specifically target to the diagnostic motifs of L1Hs subfamily, most of the step-wise primers for 5’ junction PCR could anneal to the homologous sequences of other L1 subfamilies and lead to non-specific amplifications. This non-specific annealing could reduce the detection sensitivity of 5’ junction nested PCR.

Because most of somatic L1Hs insertions were 5’ truncated with varied lengths, to maximum the sensitivity and specificity of 5’ junction PCR, we screened and selected 22 high-quality step-wise primers covering the full-length L1Hs elements to capture their 5’ junction. Because the amplicon with long size was typically hard to be amplified, we selected several 5’ truncated L1Hs insertions with known insert size to verify the step-wise primers. These 5’ truncated L1Hs insertions include: ACC1_97 (~800 bp), ACC1_15 (~1,500 bp), ACC1_60 (~2,000 bp), ACC1_111 (~3,000 bp), ACC1_99 (~4,500–5,000 bp), ACC1_20 (~6,000 bp), and ACC1_16 (~6,000 bp).

Step-wise 5' junction PCR primers spanning L1Hs elements

| Primer Name | Verified by ACC1_N1 | Verified by ACC1_N2 |
|---|---|---|
| L1Hs_5p_86 | 20 | 16 |
| L1Hs_5p_283 | 20 | 16 |
| L1Hs_5p_485 | 20 | 16 |
| L1Hs_5p_1086 | 20 | 16 |
| L1Hs_5p_1484 | 99 | 16 |
| L1Hs_5p_1635 | 99 | 16 |
| L1Hs_5p_2359 | 99 | 16 |
| L1Hs_5p_2486 | 99 | 16 |
| L1Hs_5p_2867 | 99 | |
| L1Hs_5p_3205 | 111 | |
| L1Hs_5p_3448 | 111 | |
| L1Hs_5p_3548 | 111 | |
| L1Hs_5p_3993 | 111 | |
| L1Hs_5p_4325 | 111 | 60 |
| L1Hs_5p_4451 | 111 | 60 |
| L1Hs_5p_4568 | 111 | 60 |
| L1Hs_5p_4962 | 111 | 60 |
| L1Hs_5p_5380 | 97 | 15 |
| L1Hs_5p_5767 | 97 | 15 |
| L1Hs_5p_5810 | 97 | 15 |
| L1Hs_5p_5946 | 97 | 15 |
| L1Hs_5p_6011 | | |

ACC1 gDNA were used as template for 5' junction PCR. The IDs of ACC1-specific insertions and the names of step-wise primer were labeled at the top of agarose gel image. M: 100 bp Plus DNA ladder.

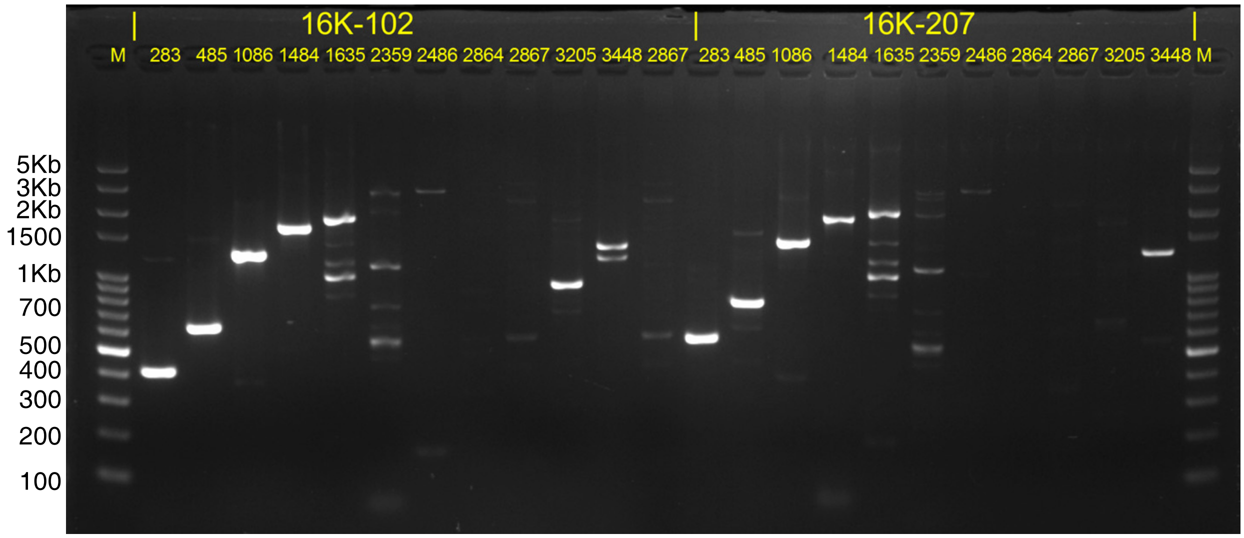

Stair-step 5' junction bands of ACC1_16 (~6,000 bp)

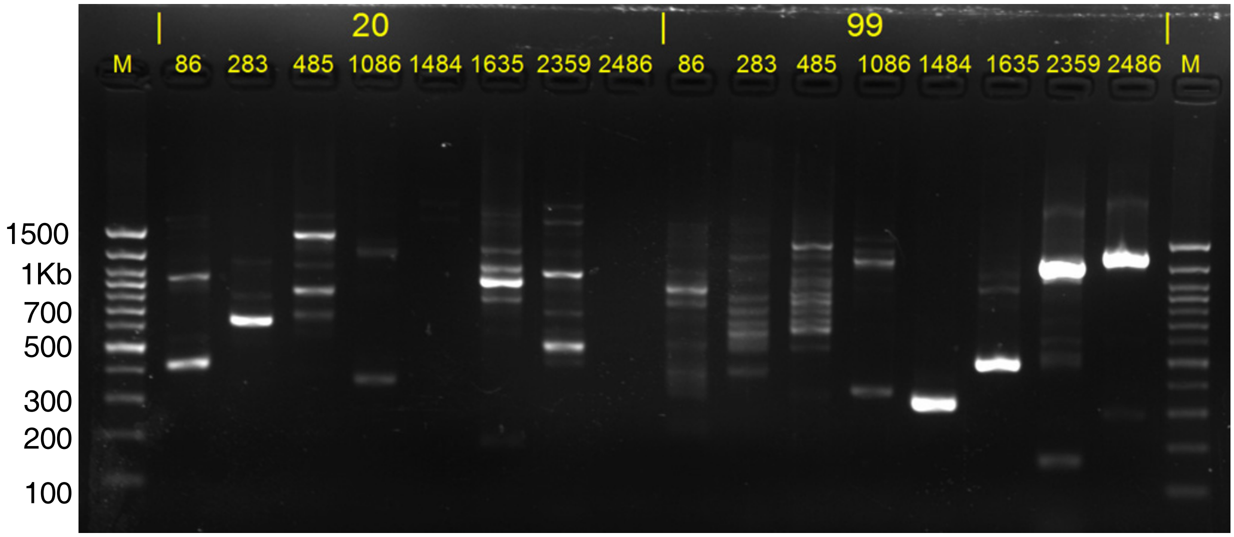

Stair-step 5' junction bands of ACC1_20 (~6,000 bp) and ACC1_99 (4,500–5,000 bp)

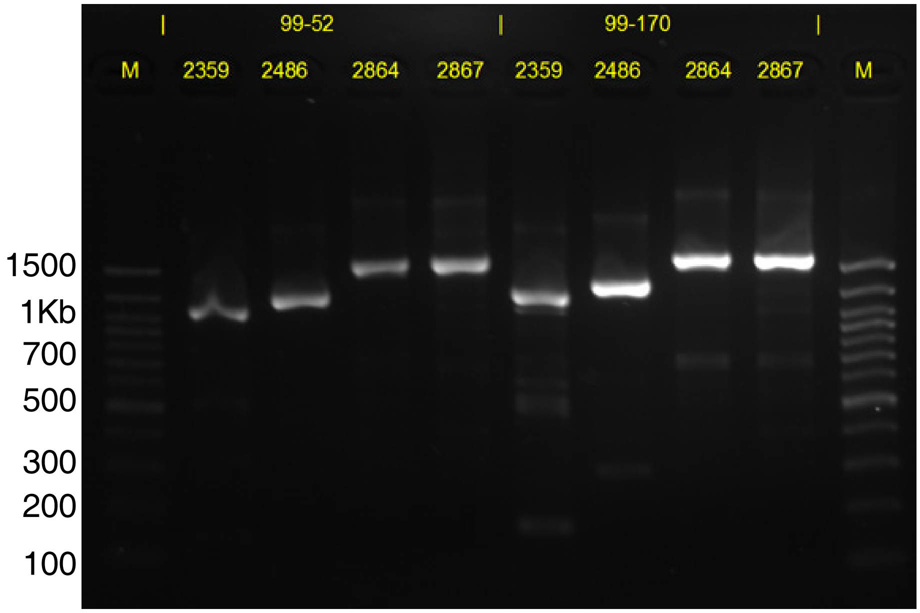

Stair-step 5’ junction bands of ACC1_99 (4,500–5,000 bp)

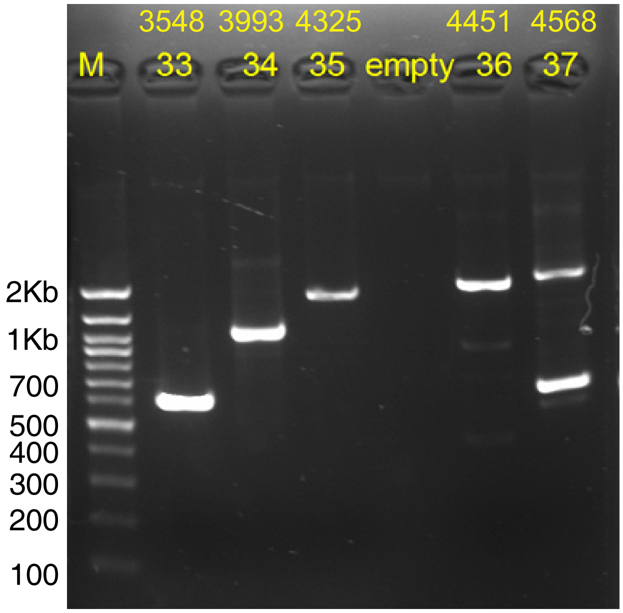

Stair-step 5’ junction bands of ACC1_111 (3,000 bp)

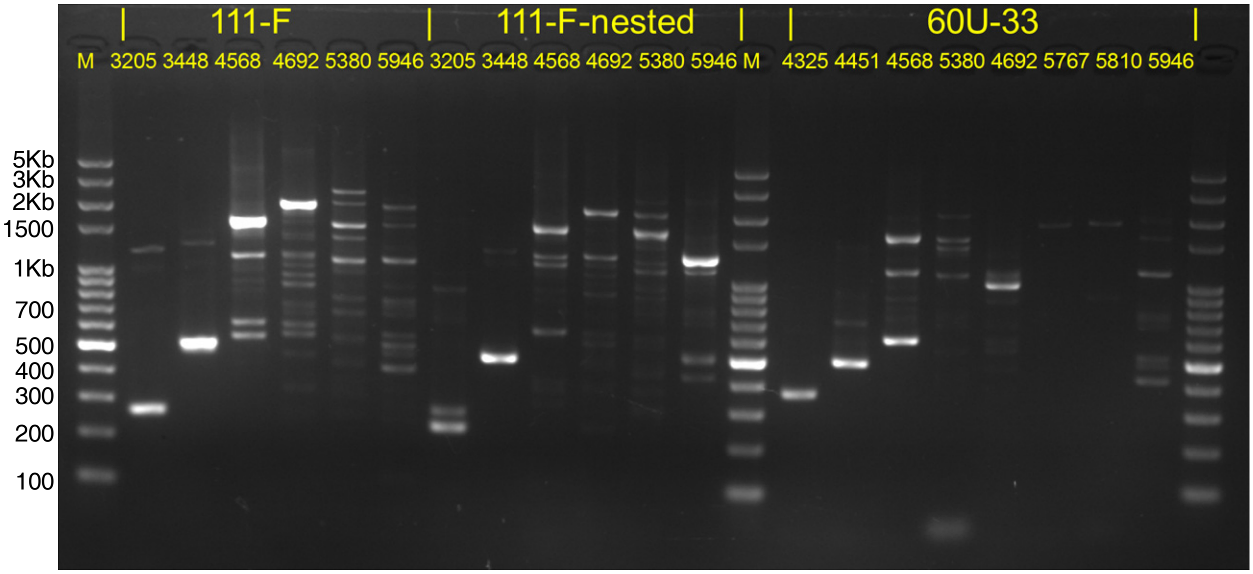

Stair-step 5' junction bands of ACC1_111 (3,000 bp) and ACC1_60 (2,000 bp)

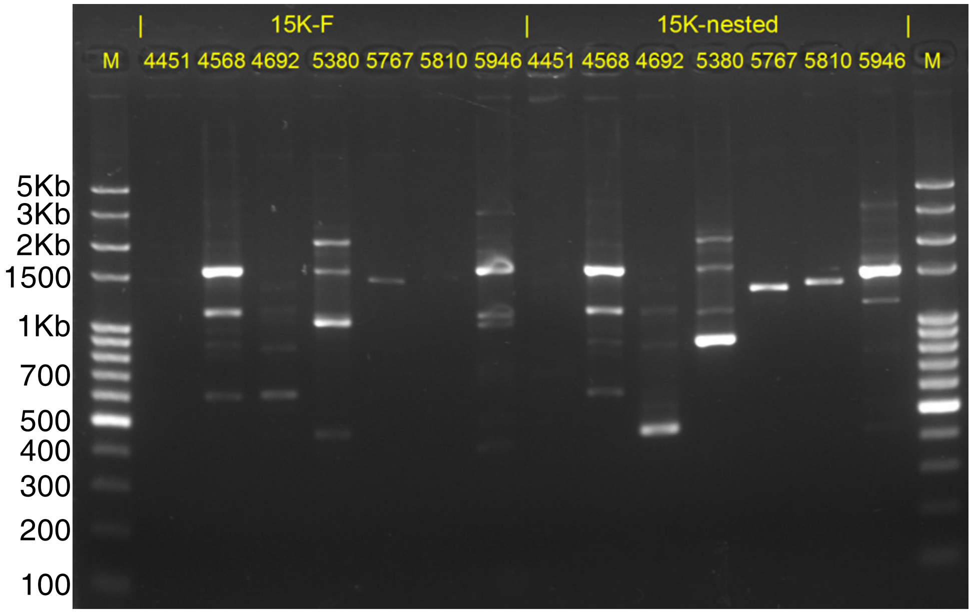

Stair-step 5' junction bands of ACC1_15 (1,500 bp)

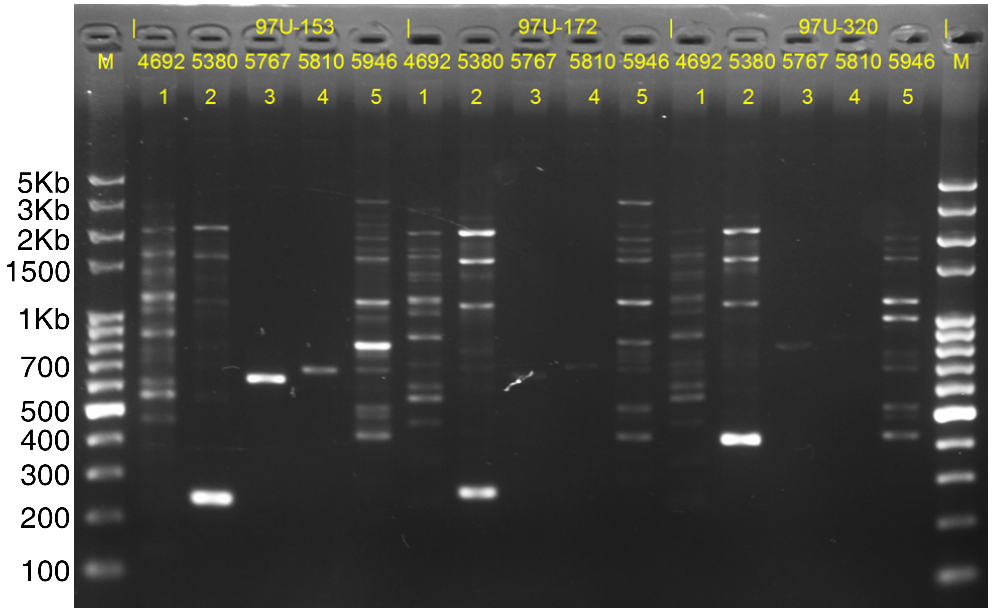

Stair-step 5' junction bands of ACC1_97 (800 bp)

In summary, our results showed that the 22 high-quality step-wise primers covering the full-length L1Hs elements could capture the 5' junction of L1Hs insertion including both full-length insertion and those 5' truncated ones with varied lengths.

Detection limit of 5' junction PCR and 5' junction nested PCR

As shown in the figures in the following pages, our results showed that 5' nested PCR could detect and yield visible amplicon bands for rare somatic L1Hs insertions whose mosaicism (percentage of cells) > 0.1%, whereas the conventional 5' PCR assay could detect insertions whose mosaicism > 10%. Because most of somatic L1Hs insertions were 5' truncated with varied lengths, to get an unbiased evaluation of the sensitivity of 5' junction PCR, we selected and tested on four 5' truncated L1Hs insertions with known insert size. These 5' truncated L1Hs insertions include: ACC1_97 (~800 bp), ACC1_60 (~2,000 bp), ACC1_111 (~3,000 bp) and ACC1_16 (~6,000 bp).

First, we evaluate the detection limit of conventional 5' junction PCR. A mixed-DNA series containing 50%, 10%, 1%, 0.1%, and 0.01 ACC1 gDNA were prepared using ACC1 and ACC2 gDNA. The IDs of ACC1-specific insertions and spike-in percentages were labeled at the top of agarose gel image. Yellow arrows indicated the band with target size. Red arrow indicated the non-specific band. M: 100 bp Plus DNA ladder. For details, see sections in the Materials and Methods "Positive control experiments" and "L1 5' junction nested PCR validation".

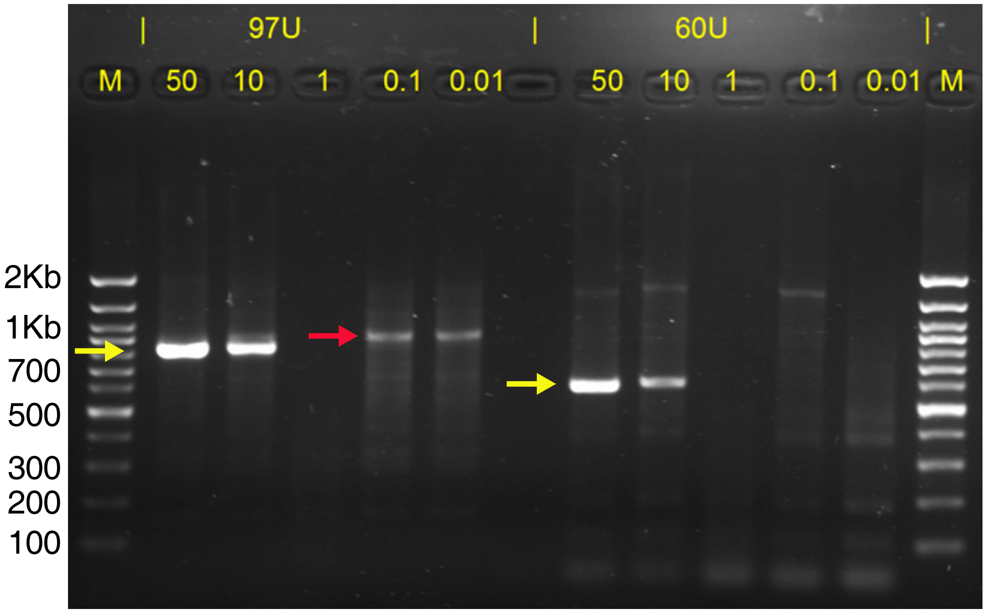

The detection limit of 5’ junction PCR for ACC1_97 and ACC1_60

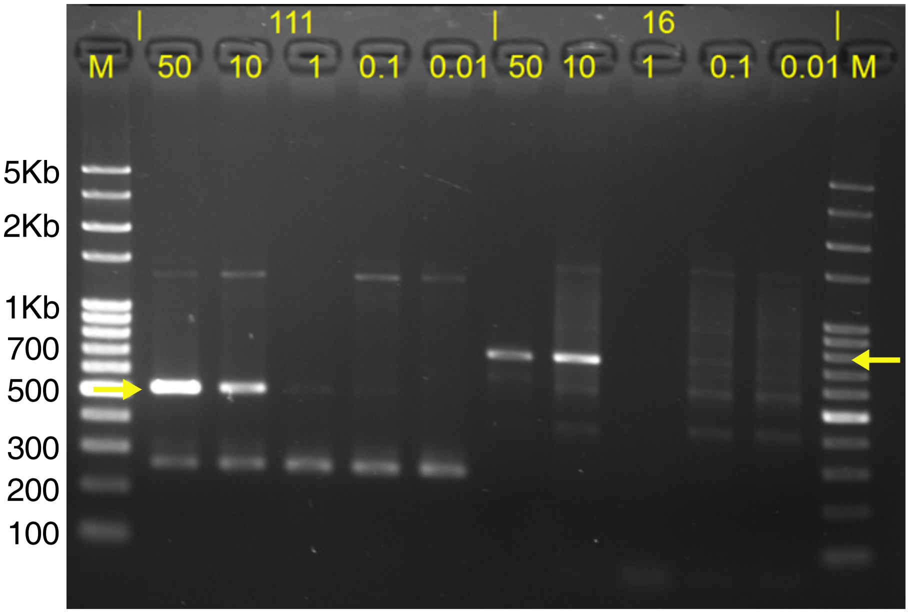

The detection limit of 5’ junction PCR for ACC1_111 and ACC1_16

Our positive spike-in experiments showed that the 5’ junction PCR can detect and yield visible amplicon bands for somatic L1Hs insertions whose mosaicism (percentage of cells) > 10%.

As shown in the figure below, we also noticed that 5' junction PCR could have higher sensitivity when deceasing the annealing temperature (Ta). The detection limit could reach to spike-in with 1% mosaicism. At the same time, lower Ta would lead to more non-specific amplification.

A mixed-DNA series containing 50%, 1%, and 0.1% ACC1 gDNA were prepared using ACC1 and ACC2 gDNA. The spike-in percentages and annealing temperatures were labeled at the top of agarose gel image. Yellow arrows indicated the band with target size. M: 100 bp Plus DNA ladder.

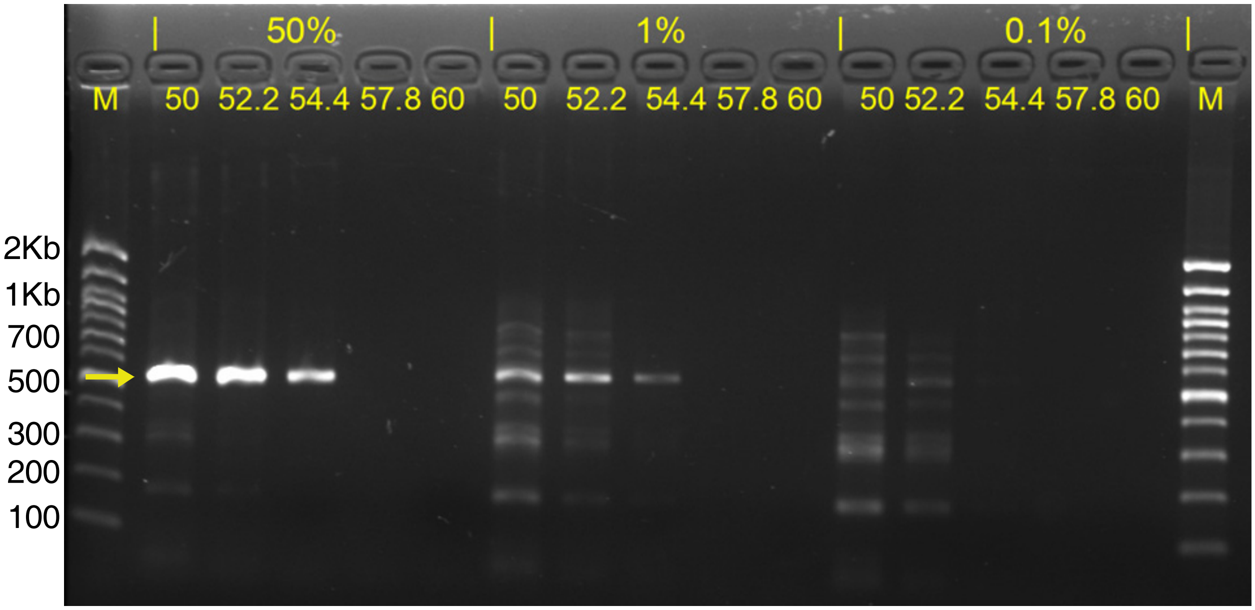

5' junction PCR for ACC1_111 (3,000 bp) insertion with low Ta

Second, we keep using these four ACC1-specific L1Hs insertions with varied lengths to evaluate the detection limit of 5' junction nested PCR. A mixed-DNA series containing 50%, 10%, 1%, 0.1% and 0.01% ACC1 gDNA were prepared using ACC1 and ACC2 gDNA. The IDs of ACC1-specific insertions and spike-in percentages were labeled at the top of agarose gel image. Ctrl: 5 ng ACC1 gDNA as template. M: 100 bp Plus DNA ladder. For details, see sections in the Materials and Methods "Positive control experiments" and "L1 5' junction nested PCR validation".

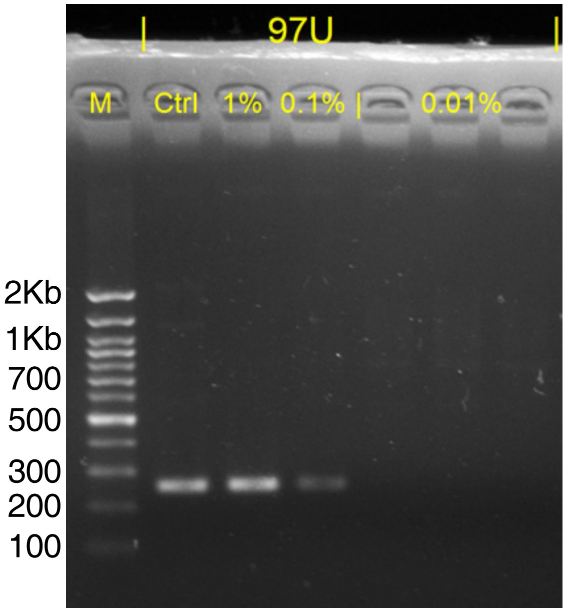

The detection limit of 5' junction nested PCR for ACC1_97

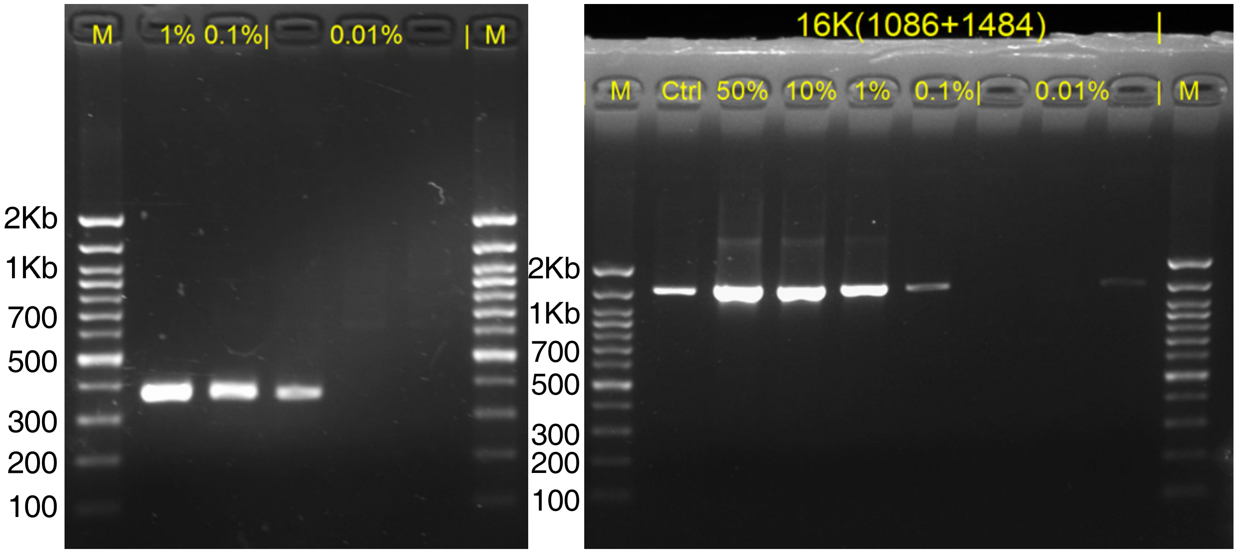

The detection limit of 5’ junction nested PCR for ACC1_16

using different step-wise primer pairs

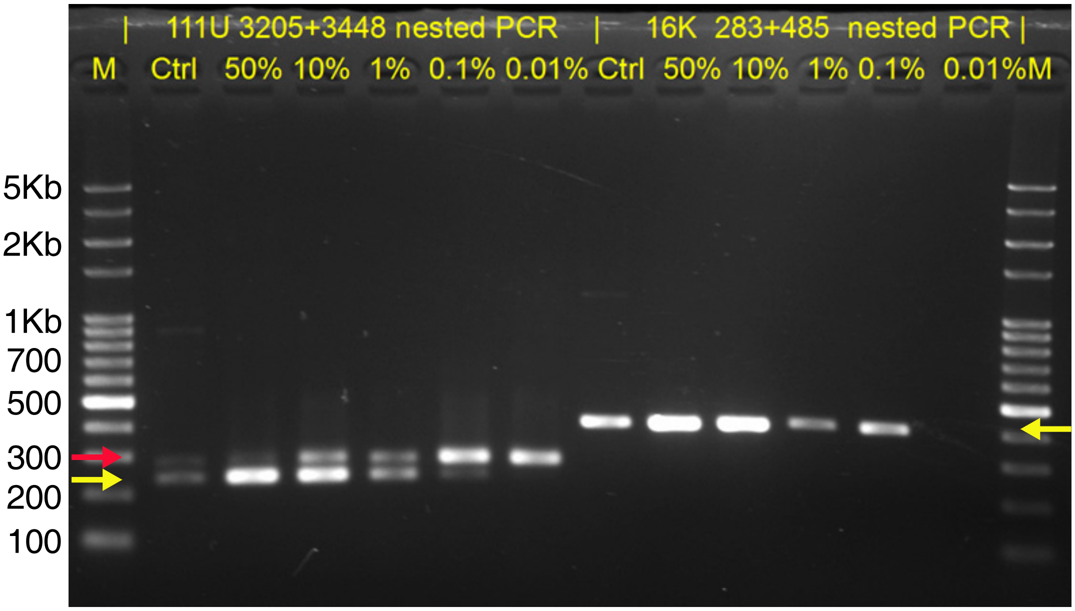

The detection limit of 5’ junction nested PCR for ACC1_111 and ACC1_16

Our results showed that the detection limit of 5’ junction nested PCR could reach to 0.01% mosaicism. However, due to the low-frequency of such insertions, only some of them could be re-sampled by chance and amplified with a faint band. In summary, we concluded that 5' junction nested PCR could robustly amplify insertions with mosaicism > 0.1%. Therefore, in this study, our validation focused exclusively on clonal somatic insertions whose mosaicism were at least 0.1%.
