## Appendix 5. Experimental validation of polymorphic germline L1Hs insertions for "Somatic LINE-1 retrotransposition in cortical neurons and non-brain tissues of Rett patients and healthy individuals"

To experimentally validate the HAT-seq predicted germline insertions, we performed 3' junction PCR validation on a random subset of polymorphic insertions from among the ten individuals, including 8 sites out of 160 polymorphic KRs, 20 sites out of 451 KNRs, and 2 sites out of 48 UNKs (S8 Table [3' PCR NOTE]).

3' PCR validation of polymorphic germline L1Hs insertions

Validation primer sequences used for each candidate insertion, the predicted target amplicon sizes of 3' junction PCR products, and statistics on 3' PCR validation results could be found in S16 Table. The IDs of ten individuals were labeled at the top of each page and agarose gel image. M: 100 bp Plus DNA ladder. For details, see sections in the Materials and Methods "L1 3' PCR and full-length PCR validation".

### Germline L1Hs KR insertions validation using 3' junction PCR

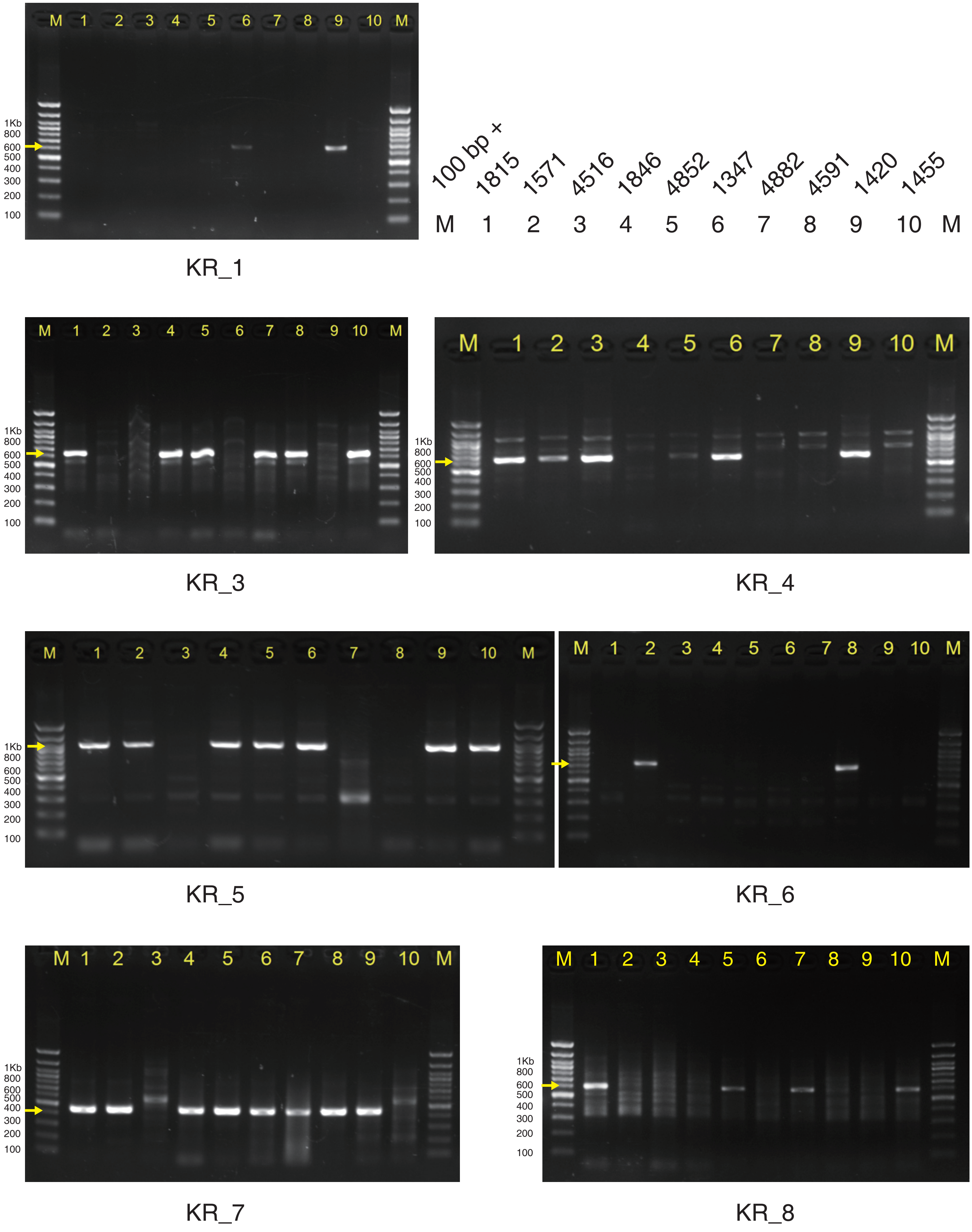

### Germline L1Hs KNR insertions validation using 3' junction PCR

| M | 1 | 2 | 3 | 4 | 5 | 6 | 7 | 8 | 9 | 10 | M |
|---|---|---|---|---|---|---|---|---|---|---|---|
| 100 bp + | 1815 | 1571 | 4516 | 1846 | 4852 | 1347 | 4882 | 4591 | 1420 | 1455 | |

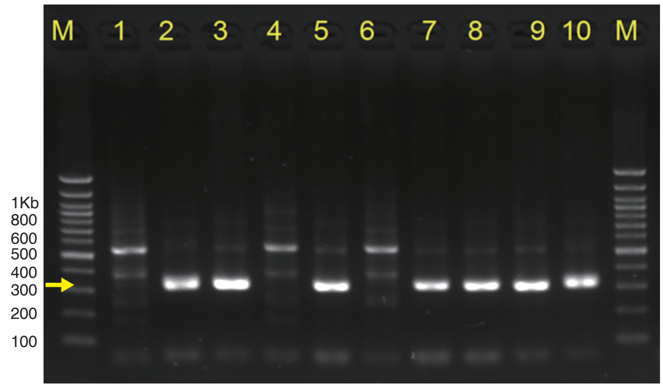


KNR_1

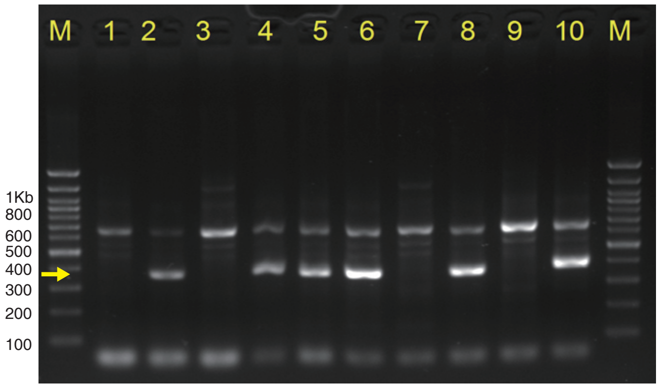


KNR_2

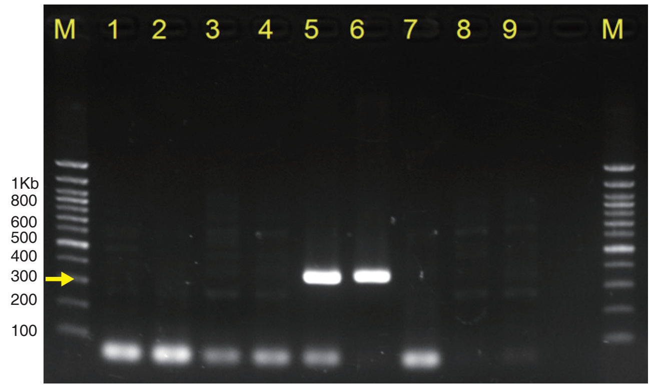


KNR_3

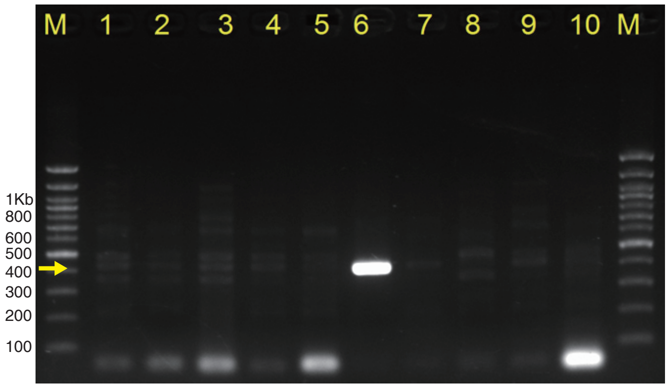


KNR_4

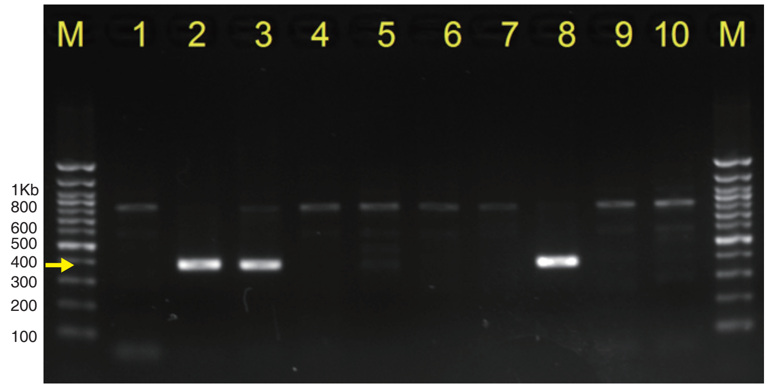


KNR_5

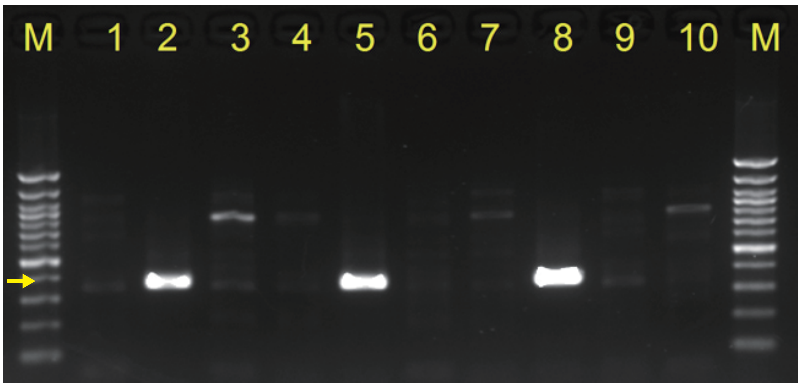


KNR_6

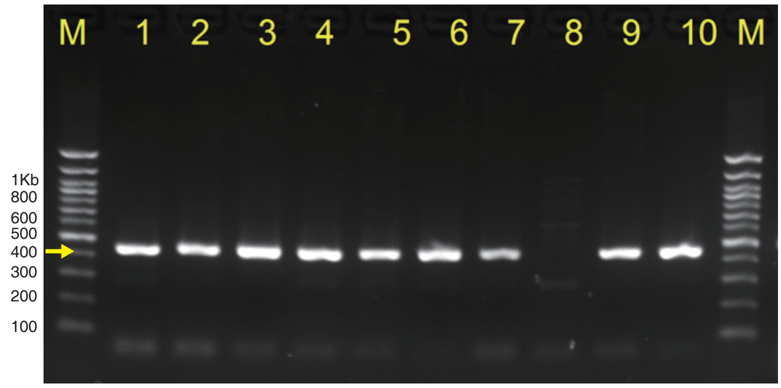


KNR_7

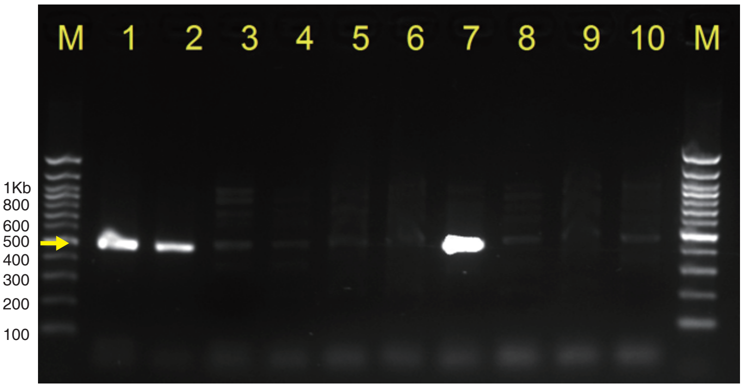


KNR_8

### Germline L1Hs KNR insertions validation using 3' junction PCR

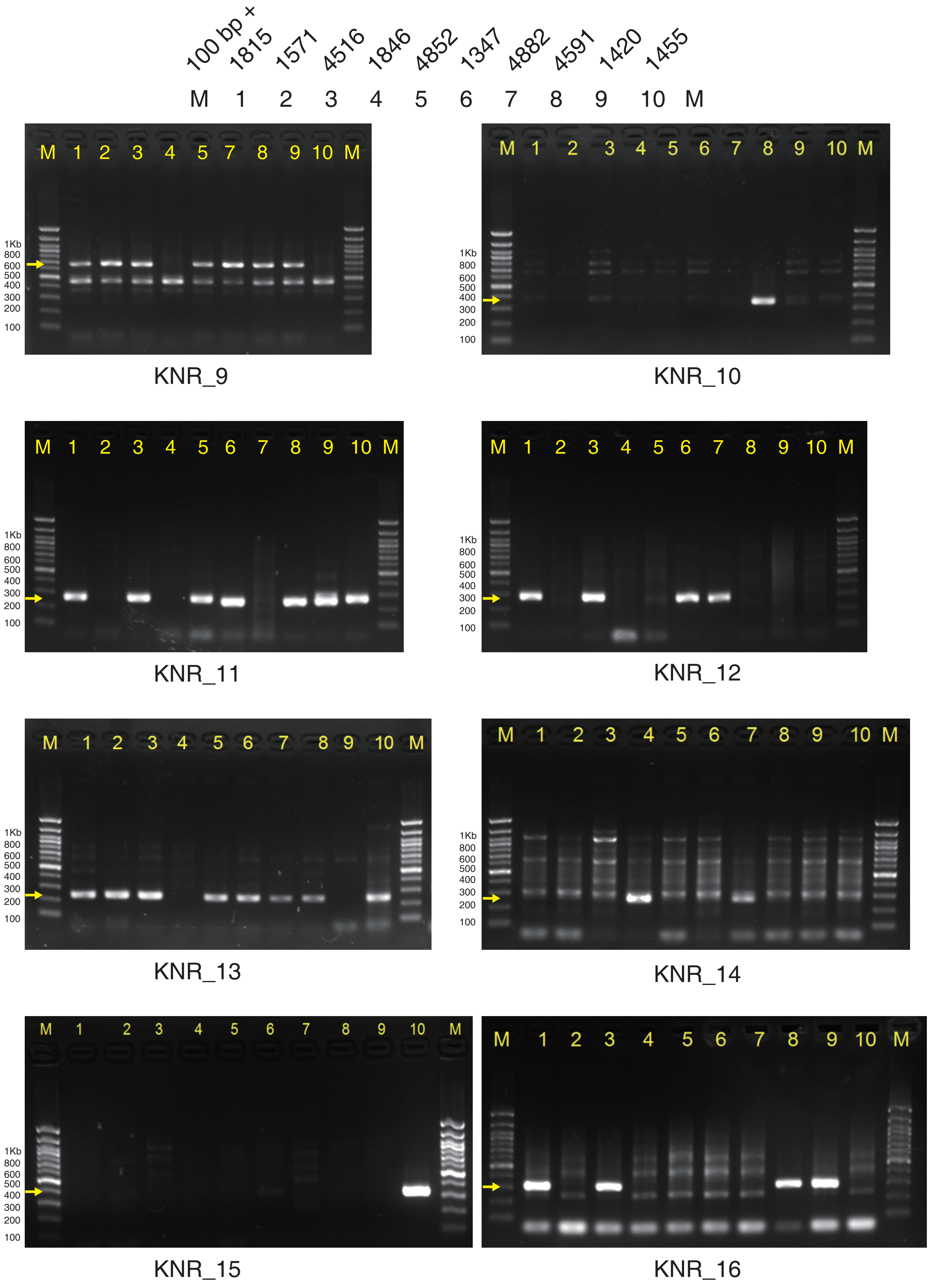

### Germline L1Hs KNR insertions validation using 3’ junction PCR

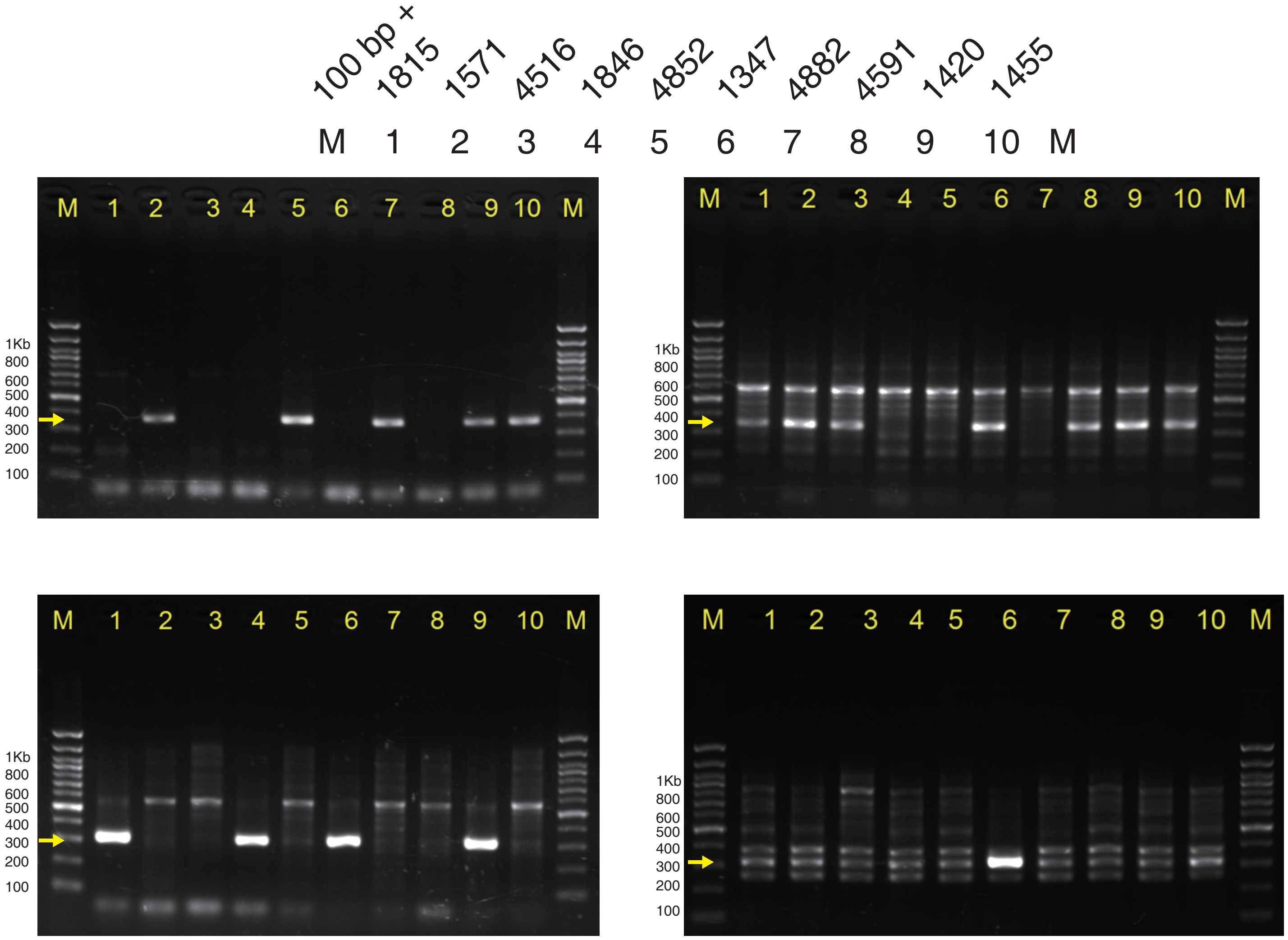


### Germline L1Hs UNK insertions validation using 3’ junction PCR

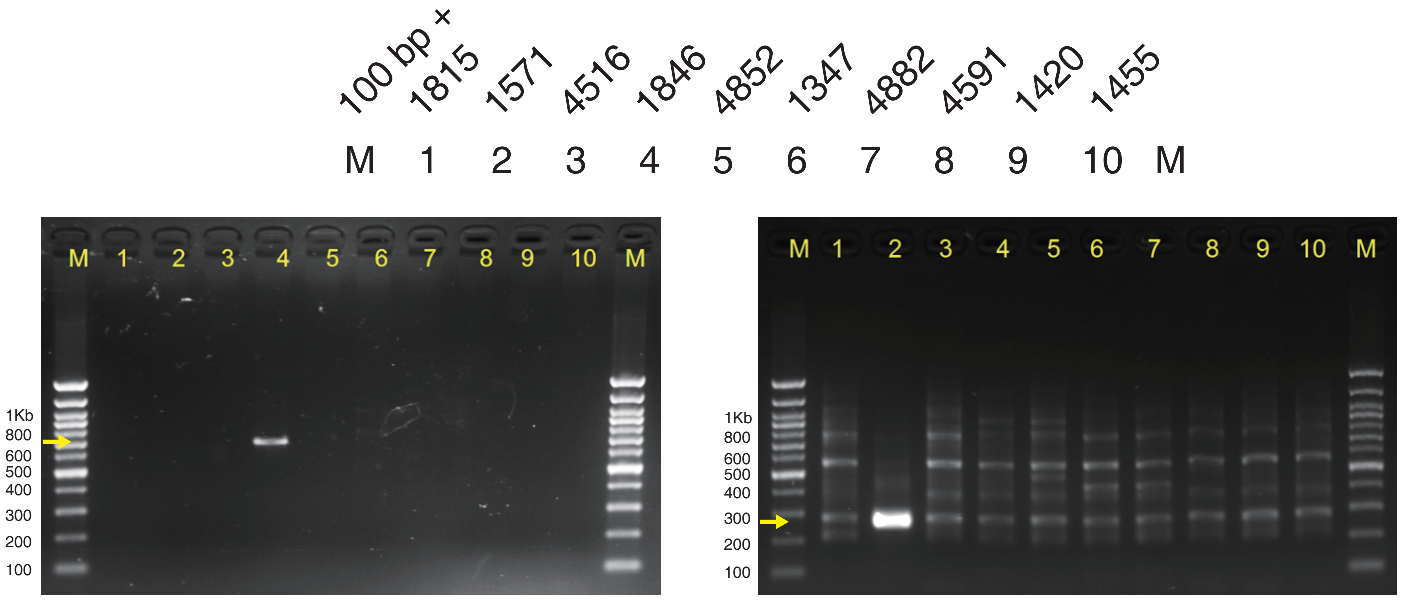
